# Modeling of possible quadruplexes and i-motifs formed during DNA contacts: strategy, classification, most probable shapes, origami based on quadruplexes

**DOI:** 10.1101/2022.10.10.511558

**Authors:** Vladimir B. Tsvetkov

**Affiliations:** Federal Research and Clinical Center of Physical-Chemical Medicine, Moscow 119435, Russia; Institute of Biodesign and Complex System Modeling, I.M. Sechenov First Moscow State Medical University, Trubetskaya Str. 8-2, 119991 Moscow, Russia

## Abstract

A strategy for creating 3D models of non-canonical forms of DNA and RNA is proposed in this research. Through this strategy, models of all possible forms of non-canonical structures created by contacts of single-stranded DNA with sequence (G_3_T)_n_G_3_ and/or duplexes with (G_3_T)_n_G_3_ and (C_3_A)_n_C_3_ fragments were built and tested by MD. In particular, models of stacks formed from right and left twisted quadruplexes; the structure resembling the Holliday structure, with the hole, on the border of which the quadruplex (G4) and the i-motif (IM) were located opposite each other; layers formed from G4s grids were constructed. The most probable of these non-canonical forms arising from contacts of duplexes with (G_3_T)_n_G_3_ for n=1,3,5 were determined by estimating the contributions to the free energy.

## INTRODUCTION

It is known that DNA and RNA regions containing G-repeats can form quadruplexes (G4s) [1], while those containing C-repeats can form i-motifs (IMs) [2], respectively. The basis of IMs structure is formed by C-pairs arranged crosswise one above the other, due to which the variety of IMs forms is due only to the variation in the number of nucleotides in the loops and the number of C-pairs themselves. The world of possible G4s forms is much larger and described in sufficient detail [3]. The possibility of forming tetramers through the formation of G4 by single-stranded DNA or RNA bearing G-repeats was shown [4]. At the same time, the question of whether the formation of such forms is possible at the contacts of DNA and/or RNA duplexes has remained open until recently. Our previous article [5] described data suggesting the existence of such forms. The AFM images presented in [5] made it possible to register the fact of contact between duplexes having such repeats and melted in the region of their localization. However, due to the insufficient resolution of AFM images, it is not possible to answer the question, which particular non-canonical forms were formed. In this study, using molecular modeling, we studied the possible variants of both themselves G4s and IMs at the contact of duplexes having G and C repeats, as well as possible variations in their localization relative to each other. The conducted research demonstrated a significantly larger number of possible non-canonical forms arising in the contacts in comparison to just using the ability to draw diagrams. The topology of these shapes can be quite complex and varied. Perhaps the probable structures discovered in this research will help clarify the still unclear issues in the processes of replication, recombination and repair.

## RESULTS AND DISCUSSION

### Construction and testing of 3D models of the complexes

When, for one reason or another, it is not possible to reproduce the spatial model of the molecular structure under study by means of X-ray crystallography and NMR, and the resolution capabilities of atomic force, electron, and cryogenic-electron microscopy do not allow a detailed “atomic” study of the geometry and topology of molecular structures due to the existing physical and technical limitations can be helped by computer molecular modeling (MM). In this research MM was used to create structural full-atom models of complexes arising from the contacts of single-stranded DNA with sequence (G_3_T)_n_G_3_ or duplexes with (G_3_T)_n_G_3_ and (C_3_A)_n_C_3_ fragments resulting in formation of non-canonical structures.

The following strategy was used to create the models. At the first stage, a hypothesis was made about possible non-canonical structure, or structures and it’s or their topology. If there were several of these structures, then an assumption was made about their geometry of localization relative to each other. At the second step, a search was made for homologues for these non-canonical forms for which the existing databases contain structures obtained using XRD and NMR. In this case homologues mean the sequences having the same number of G and/or C repeats in their structures, separated by the same number of nucleotides as in the sequence under study. If a homologue was found, then its structure was taken as the basis of the model, in which mismatched nucleotides in the loops were replaced with the necessary ones using software (for example: Sybyl X, ICM-Pro) with the appropriate options. The resulting model was optimized in force fields containing the necessary parameters for describing interatomic interactions. Details are provided in the experimental section.

In the absence of a homologue assumptions based on the sequence analysis were made about the possible topology of the G4 core, the number of tetrads in the G4 case or C-pairs in the IM case, the number of nucleotides in the loops, and the type of loops. In the G4 case, the choice of the type of loop with the required number of nucleotides was implemented taking into account the topology and the number of the tetrads. To create the starting model of the non-canonical form based on the accepted assumptions, a search of structures was made in the databases containing the structures with required topology types of the core and number of tetrads in the G4 case and/or the required number of C-pairs in the IM case, and containing loops with the required number of the nucleotides and the required type.

If the databases contained only the cores of the G4 or the IM, then the cores were taken from them, and then the loops were created taking into account the required type of number of nucleotides for subsequent attachment of created loops to the core. If the core was found with a mismatched number of tetrads or C-C pairs, then in the case of a larger number than necessary, the extra tetrads or pairs were removed, and in the case of a missing one, they were completed to the required number. The completion was carried out taking into account the rotation of tetrads in the G4 or the C-C pairs in the IM relative to each other, as well as the distance at which stacking occurs between the nucleotide bases of neighboring tetrads in the G4 or the C-C pairs in the IM. At the next stage, the pre-made loops were attached to the created structures of the core of the G4 or IM. At the final stage, the created G4s and IMs fragments were placed relative to each other according to the hypothesis adopted at the first stage, and already prepared models of duplexes were attached to the ends of these structures. The assembled models of the complexes were optimized and their stability was tested by the method of molecular dynamics, the protocol of which is presented in the experimental section. Note that the above strategy can also be used to assemble similar constructs for RNA basing on the G4s and IMs DNA structures with substitutions of 2-deoxyribose by the alternative pentose sugar ribose and thymine for uracil, and subsequent optimization of the structure obtained after the substitutions. However, it should be noted that the replacement of thymine in the loops by uracil can induce a change in the core topology.

To analyze the evolution of the structural deformation of the studied non-canonical forms in the process of MD calculations, the following parameters were used:

1. to score the planar deformation of tetrads in G4s - the distances from the centers of mass of the guanine bases to the center of mass (COM) of their containing tetrad and the angles between the normals to the guanine bases and the vector connecting the COMs of the boundary tetrads
2. to make an evaluation of the twisting deformation in G4s - the angles of rotation of one tetrad with respect to another
3. to estimate the structural deformation of cytosine pairs in IMs - the distances between the COMs of the cytosine bases and the angles between the normals to the bases of the cytosines forming the pair.

Changes in the geometry of localization of the G4s relative to each other were evaluated by:

1. distances between the COMs of G4s, if they did not have stacking tetrads, and distances between the COMs of stacking boundary tetrads otherwise.
2. twist angles of G4s relative to each other, i.e. rotation angles between stacking tetrads (in the case of stacking between boundary tetrads)
3. the angle between the axes of the G4s - straight lines passing through the COMs of the boundary tetrads.

In case of 4 or more duplex contacts:

1. in the case of the formation of G4s located in the same plane, the angles between the axis of each of the G4s and the normal to the plane, in which the G4s are located, were estimated
2. when G4s are packed into a stack or stacks parallel to each other, the angles between the axis of each of the G4s and the axis of the stack (a straight line, passing through the COMs of the boundary tetrads of the boundary G4s) were estimated
3. angles between the axes of the IMs (straight lines passing through the COMs of the boundary cytosine pairs) and the normal to the plane in which the G4s are located, or the axis of the stack.

The purpose of this work is not only to propose possible variants of the non-canonical forms arising from the contact of duplexes that have G-repeats in their structure, but also to test their stability. As a result, on the graphs of the evolution of the parameters described above and characterizing the stability of the G4s and the IMs, not the values themselves are presented, but their deviation from the average of these values along the trajectory. Such an approach to presenting data, when it is G4s and IMs stability that is being studied, and the interest is not so much the values themselves, but the stability of these variables values, seems to be very reasonable. In contrast to the parameters characterizing the stability of G4s and IMs, in the case of describing the position of duplexes relative to each other, on the graphs of the angles of the axes passing through the duplexes, the values of these angles were used.

A few explanatory notes to the figures, which show the models themselves, their schemes, graphs of the evolution of parameter values that describe the geometry of the arrangement of nucleotides in non-canonical forms, the localization of non-canonical forms relative to each other:

1. In the schemes of the studied models, all nucleotides in (G_3_T)_n_G_3_ and (C_3_A)_n_C_3_ involved in the formation of non-canonical forms are numbered.
2. The legends for the graphs of parameters describing the arrangement of nucleotides in non-canonical forms indicate the name of the nucleotide by means of the first letter of its name and its number.
3. In the description of the behavior of entire tetrads in G4s, in the case when the tetrads are from different G4s, the G4’s number is indicated in the legend for each tetrad. If the curve depicts the behavior of tetrads of the same G4 relative to each other, then the number of the G4 to which both tetrads belong is indicated at the beginning of the legend, then the numbers of the tetrads themselves.
4. The numbers of G4s and IMs are marked with Roman numerals, and the numbers of tetrads in G4s are marked with Arabic.
5. Hydrogen bonds between nucleotides are marked with black dotted lines.
6. The duplexes on the drawings and diagrams are colored and named as follows: the first symbol is a capital letter of the color of this duplex on the drawing (for instance R[ed]) and the second symbol is its number in Roman numeral.

Let’s designate strands containing fragment (G_3_T)_n_G_3_ as G-strands, and complementary to them as C-strands, similarly, the motives (G_3_T)_n_G_3_ and (C_3_A)_n_C_3_ themselves as G-motive and C-motive.

To create the models 5’-d(TATCTGA(C_3_A)_n_C_3_ACAGATA) sequence from our previous research [5] was used, in this case n had values from 1 to 5 respectively. Note that the obtained results about the stability of the non-canonical forms do not depend on the choice of the nucleotide sequence in the duplexes flanking these non-canonical structures. Three variants of contacts were considered in the study: without a strand transitions from one duplex to another, with and without a transition, but with mutual girth of one strand by another. The first variant is the most probable one and occurs in the case of melting of duplexes in the regions containing G and C repeats. The case of strands transition may arise during homologous recombination. The latter option can happen in the process of DNA repair. All three variants can also arise in a hypothetical situation, when a solution containing DNA sequences is heated to the point where duplexes break up into separate strands. And then, as a result of cooling, formation of duplexes from complementary strands occurs. In this situation, variants of contacts and non-complementary strands are also possible. In this case, if the contact occurs with C-motive, then it will be a tetrameric i-motif with A side bulges, while in the case of G-motive contact, it will be the stack of three-tetrad G4s with T side bulges. In view of the obvious geometry of tetramers, they are not considered further in this study. The possibility of forming G4s and IMs appears already for n=2. The variety of their possible arrangements relative to each other increases with the growth of n.

### Possible variants of non-canonical structures in the case of n=1

Obviously, in the case of n=1, the formation of any G4s or IMs within one strand is impossible. Therefore, their formation occurs only when two strands of contacting duplexes come into contact. When n=1, both simple variants with the formation of only G4 or IM are possible, as well as complex ones, in which both strands of each of the duplexes participate in the formation of non-canonical structures. In **Fig. 1.A-B** and **Fig. 1.A-B.S.** simple variants are shown, only one of the complementary strands of the contacting duplexes is involved in the formation of the non-canonical form. Further such options will not be considered in view of their obviousness. In **Fig.1.C-D** and **Fig. 1.C-D.S.** data are presented for the cases when the G-strands of the contacting duplexes form the dimeric G4, and complementary strands form the dimeric IM.

**Figure 1.**
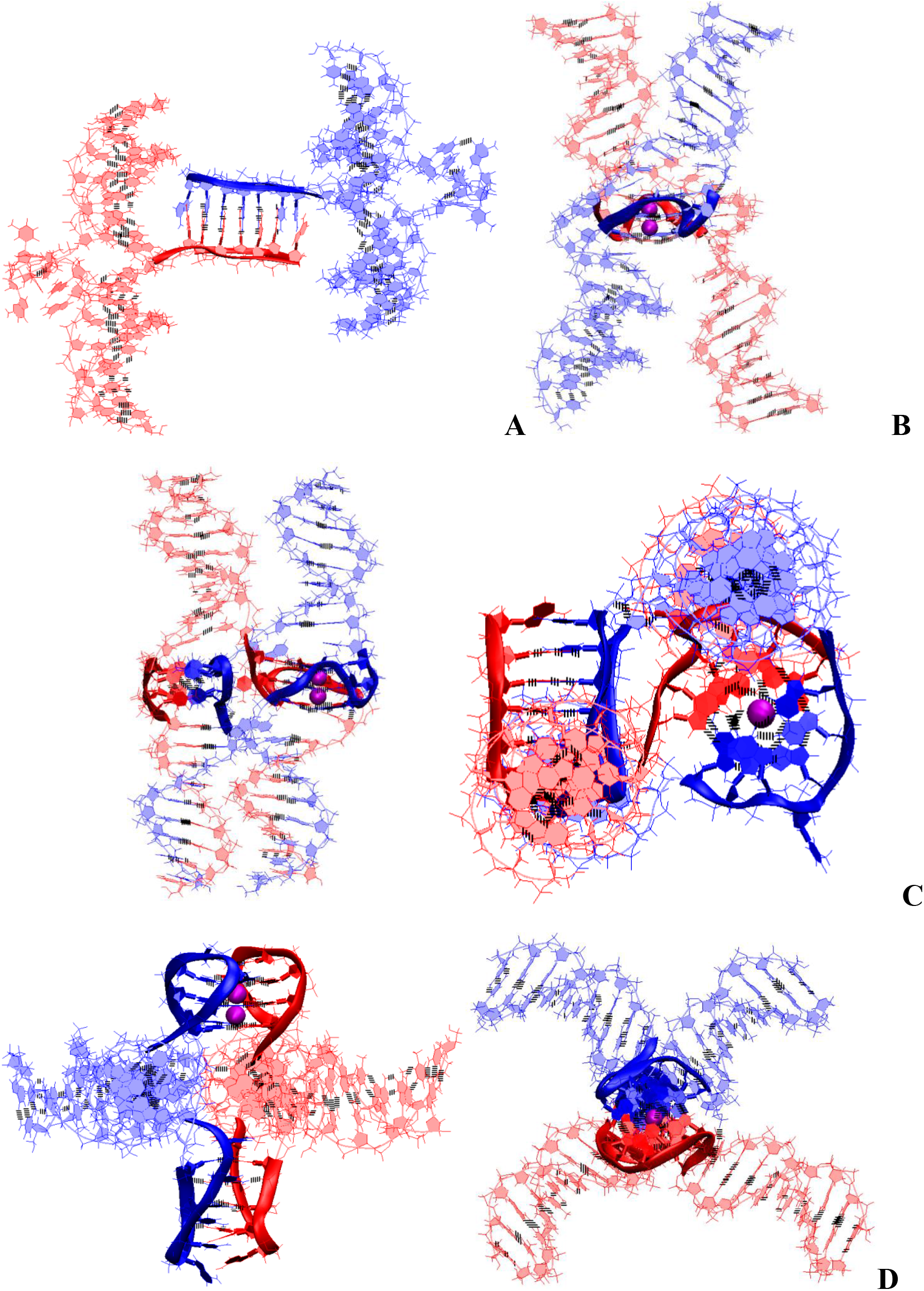
Bimolecular complex of DNA duplexes containing G_3_TG_3_ and C_3_AC_3_ fragments (starting conformations). **A** – with only dimeric head-to-tail IM; **B** – with only dimeric parallel G4; **C** – with dimeric parallel G4 and head-to-tail IM; **D** – with dimeric antiparallel G4 and head-to-head IM.

The case of simultaneous formation of both G4 and IM without the strands transfer from one duplex to another, in the case of n=1, the only one is possible, and it is shown in **Fig.1.D** and **Fig.1.D.S.** In this case, the G4 has the antiparallel topology with the lateral loops and the dimeric IM is twisted at one of the points of contact with the duplex. The IM is of head-to-head type. The location of the parts of the duplexes in the complex, involved in the formation of the G4 and the IM, resembles the Holiday structure. But the hole in the Holliday structure has non-complicated structure, while in the case of contact through non-canonical forms, the complexity of organizing the hole increases significantly. From the graphs describing the positions of pairs in the IM, see **Fig.1.D.S.F-G,** it can be seen that, in whole, this IM variant is stable, except for the behavior of only one boundary pair. The cytosines included in it, as a result of thermal fluctuations, sometimes move away from each other, while the angle between the normals to the planes containing the bases of cytosines do not change so much as to lose the possibility of forming hydrogen bonds. The terminal guanines in the lateral loops form a tetrad via Hoogsteen pairs, adding another tetrad to the G4. Should be noted indeed that the bases of these boundary guanines, in the process of stability testing by the MD method, were more often located at a slightly acute angle to the plane of the neighboring tetrad, see **Fig.1.D.S.D-E**. This is due to the fact that the presence of only one nucleotide in the lateral loop can create insignificant stresses in the sugar-phosphate backbone in case when the bases of boundary guanines form tetrad plane. The halves of the duplexes are arranged like rays coming out of one point at an angle of 90 degrees._ Comparative analysis of the images of the initial and obtained at the last step of the MD calculations of conformations, as well as the behavior of the curves on the graphs of **Fig.1.D.S.C**, showing the evolution of the angles between the halves of the duplexes, indicates that such an arrangement of duplexes relative to each other stays unchanged throughout the entire calculated trajectory.

The variant of forming the parallel G4s by G-strands under the condition of simultaneous formation of the IM by the complementary strands is possible only in the case when the strands pass from one duplex to another through non-canonical structures, see **Fig.1.C** and **Fig.1.C.S**. In this case, the IM has a head-to-tail type, and the configuration of the IM relative to the G4 is such that the straight line passing through the COMs of cytosine pairs is perpendicular to the axis connecting the tetrads’ COMs.

Analysis of the evolution of the values of the parameters characterizing the geometry of the IM and the G4, as well as the conformations initial and obtained at the last step of the trajectory, shown in **Fig.1** and **Fig.1.S,** allows to see that all variants of non-canonical forms are stable.

In **Fig.1.S.E.** graphs of evolution of the values of contributions to free energy are presented in the case of variants of complexes with the simultaneous formation of G4 and IM. The analysis of the data presented in the graphs indicates that the variant with the formation of an antiparallel G4 is more energetically favorable. The stress energy is practically the same. The electrostatic and Van der Waals contributions in the case of the variant with the formation of an antiparallel G4 are significantly lower, which determines that the internal energy is significantly lower. On the other hand, the polar component of the solvation energy is significantly lower in the case of the formation of the parallel G4, which ultimately leads to the fact that the difference between the sums of contributions to the free energy for these two variants is insignificant compared to the difference in the values of internal energy.

In the cases where the local concentration of duplexes is high, it is possible that the strand of one duplex containing G_3_TG_3_ forms the parallel G4 with a similar strand of the second duplex, while the complementary strand forms the IM with the corresponding strand of the third duplex. The embodiment of this option is shown in **Fig.2.A** and **Fig.2.A.S.**

**Figure 2.**
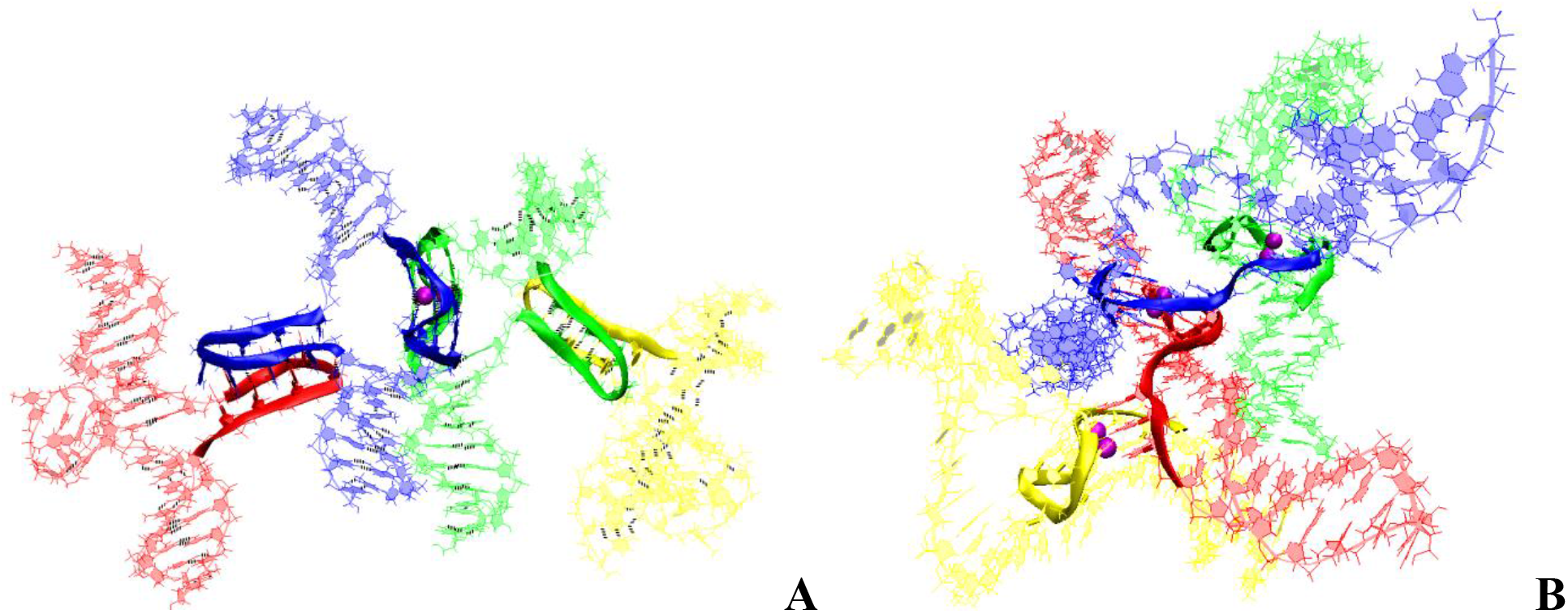
Tetrameric complexes complex of DNA duplexes containing (G_3_T)_n_G_3_ and (C_3_A)_n_C_3_ fragments (starting conformations). **A** – n = 1 with dimeric two head-to-tail IM and parallel G4; **B** – n = 2 with three dimeric parallel G4.

### Possible variants of the non-canonical structures in the case of n=2

At n=2, in the case of the formation of only one IM upon contact of the duplexes, there are no qualitative changes compared to n=1, due to the fact that the number of possible cytosine pairs arranged crosswise with respect to each other is also equal to 5. With the formation of the only one parallel dimeric G4, the number of nucleotides in the propeller loops will increase to four. Variants with simultaneous formation of the G4 and the IM are possible only the same as in the case of n=1. In contrast to the previous variants, for n=2 in the case of contacts of four duplexes it is possible to form 3 parallel G4s, see **Fig.2.B.** From **Fig.2.B** and **Fig.2.B.S.1** it can be seen that the geometry of the location of the G4s is such that the projections of the COMs of the G4s onto the plane form an equilateral triangle with acute angles, the value of which is 60 degrees. In this case, the central G4-dimer is formed with an equal contribution of the strands that form it. The boundary dimeric G4 are formed in the ratio of 3/4:1/4, i.e. 3 out of 4 G-tracts forming the G4 belong to one strand, and the rest of the other. From the evolution of the behavior of the parameter values characterizing the stability of the geometry of the G4s during the MD calculations, see **Fig. 2.B.S.2-3**, it is obvious that the G4 is stable.

### Possible variants of the non-canonical structures, in the case of n=3

Of course, as with n < 3, in this case, variants with the formation of only G4 or IM forms are possible. In view of the obviousness of the geometry of the arrangement of nucleotides in these variants, they are not considered further. With an increase n to 3, the diversity of forms with the simultaneous formation of G4s and IMs forms arising from contacts of duplexes containing (G_3_T)_n_G_3_, cardinally increases. All the variety possible geometries of the location of non-canonical forms relative to each other in the case of contacts of 2 duplexes can be divided into two groups. The first group includes variants in which G-strands form G4s located one above the other.

1. **“1,3 hitch and two IM-monomers” (1.3h-2mi)** is presented in **Fig.3.1.A** and **Fig.3.1.A.S.1**. In this option, G-strands form a stack consisting of two G4s with side bulges of T. Each of these G4s is formed in the ratio of 3/4:1/4, i.e. 3/4 of each G-fragment takes part in formation one G4, and the rest 1/4 of the same G-fragment belongs to another G4. It should be noted that while folding the G4’s form the G-strand from one duplex girths the similar G-strand of another duplex. Each of the C-strands forms the monomeric IM consisting of six cytosine pairs. MD showed, see **Fig.3.1.A.S.2-3**, that the G4s and their location relative to each other are stable, but the penultimate pairs in the IMs were not stable.
2. **“1,3 hitch and head-to-tail IM-dimer” (1.3h-hti)** is shown in **Fig.3.1.B** and **Fig.3.1.B.S.1**. In contrast to the previous case, the C-strands form the head-to-tail dimeric i-motif containing 10 cytosine pairs, divided in two equal halves by a fragment, containing adenines and cytosines, separated from each other the way that they become unable to form hydrogen bonds, see diagram **Fig.3.1.B.S.1.B**. Analysis of the values of the parameters presented in **Fig.3.1.B.S.2-3** showed that during MD there were no changes in the G4s structures, the IMs and their geometry of location relative to each other.
3. **“1,2 hitch and two IM-monomers” (1,2h-2mi)** is shown in **Fig.3.1.C** and **Fig.3.1.C.S.1**. As in the previous two cases, the G-strands with mutual girth of each other form a stack of two dimeric parallel G4s, whose boundary tetrads form a stacking interaction. Each G4 is formed in the ratio of 2/4:2/4, i.e. 2 of the 4 *G-tracts*, that form the G4, belong to one strand, and the other two to another one. In this case, the *G-tracts* formed by one strand are located on opposite sides of the stack. As in the first case considered above, each of the C-strands forms IM-monomer consisting of six cytosine pairs. From **Fig.3.1.C.S.2-3**, it can be seen that the G4s and they localizations relative to each other are stable, and the penultimate pairs in the IMs, as in case **1,** were deformed during the MD.
4. **“Stacking and two IM-monomers” (st-2mi)** is presented in **Fig.3.2.A** and **Fig.3.2.A.S.1**. Earlier [6,7] it was shown that in the duplex, containing GGGT repeats and melted in the area of their localization, the simultaneous existence of the G4 in the strand containing G-repeats and in the IM in the complementary strand, is impossible due to their steric overlap. However, the author constructed 3D model of the DNA duplex corresponding to this case. The stability of the G4 and the IM located opposite to each other was tested by the MD method. The analysis of the calculated trajectory has demonstrated the stability of non-canonical forms in this model. As a result, the possibility of contact was considered, in which the lower tetrad of the G4 formed by the G-strand of one duplex is located above the upper tetrad of the G4 formed by the similar strand of another duplex. In this variant, as in cases **1** and **3**, each of the C-strands forms the monomeric IM consisting of six cytosine pairs. During the MD calculations, the G4s themselves did not undergo significant changes. As can be seen from the graphs on **Fig.3.2.A.S.2**, the distance between the G4s increased by 0.5 angstroms, and one G4 rotated relative to the other by an angle equal to 30 degrees during the modeling. According to **Fig. 3.2.A.S.3** as well as in cases **1** and **3**, the penultimate pairs in the IMs were deformed.
5. **“Stacking and head-to-tail IM-dimer” (st-hti)** is shown in **Fig.3.2.B** and **Fig.3.2.B.S.1**. In this case, the G-strands form the G4s located relative to each other as in the previous variant **4**. In its turn, the C-strands form the head-to-tail dimeric IM, as in variant **2**. During the dynamics, the G4s remained stable. In contrast to previous case **4**, the distance between the G4s and the angle of rotation between neighboring tetrads did not change significantly during the calculation, see **Fig.3.2.B.S.2**. In the IM, only the boundary pairs were deformed, see **Fig. 3.2.B.S.3**.
6. **“Flip-flop and two IM-monomers” (ff-2mi)** is shown in **Fig.3.2.C** and **Fig.3.2.C.S.1**. In this option, G-strands form the two-tetrad G4s located one above the other. But unlike the cases considered earlier, the axes, passing through the COMs of tetrads, are directed to the opposite sides. In this case, the boundary stacking tetrads consist of 3 guanines from one strand and 1 guanine from the other. The second tetrads are joined by the tetrads formed from 2 guanines and 2 thymines, see the scheme **Fig. 3.2.C.S.1.B**. An analysis of the values of the quantities, characterizing the geometry of the arrangement of guanines and thymines in tetrads, showed that tetrads, composed exclusively of guanines, are stable, see **Fig. 3.2.C.S.2**. And the tetrads, formed from two guanines and two thymines, slightly spread. The thymines, included in the tetrads, slightly move away from their original location relative to the tetrad’ COM, while the thymine bases remain in the tetrad’ plane. Each of the C-strands forms the monomeric IM consisting of six cytosine pairs, as in previous cases **1**, **3** and **4**. The geometry of the location of the G4s relative to each other has not changed during the MD, see **Fig. 3.2.C.S.2**. As follows from **Fig. 3.2.C.S.3**, in the IMs, in contrast to cases **1**, **3** and **4**, during the calculation, not the penultimate, but the last cytosine pairs were subjected to deformations.
7. **“Stacking of right and left handed parallel G4-dimers, and two IM-monomers” (rl-2mi)** is demonstrated on **Fig.3.3.A** and **Fig.3.3.A.S.1**. In this option, the G-strands form two parallel G4s located one above the other, wherein the lower one is right-handed, and the upper one is left-handed. Each of them is formed by half of one strand, half of the other. As in case **6**, the axes, passing through the tetrads’ COMs, are directed to the opposite sides. The C-strands form monomeric the IMs, consisting of six cytosine pairs. Analysis of the evolution of the parameter values presented **in Fig. 3.3.A.S.2-3**, showed that the structures of both right-handed and left-handed G4, as well as their localization relative to each other, did not change during the calculations, and in the IM, as in previous cases **1**, **3** and **4**, the penultimate pairs underwent deformation.
8. **“Antiparallel G4-dimer and head-to-head IM-dimer” (aq-hhi)** is shown in **Fig.3.3.B** and **Fig.3.3.B.S.1**. This option has already been considered for n=1. As n increases to 3, the number of all tetrads in the antiparallel G4s increases to 5. Thymines, which are part of G-motive and are not included in the lateral loops, form side bulges. The guanines in lateral loops compared to the previously considered variant with n=1, during MD calculations showed a stronger deviation from the original location, see **Fig. 1.D.S.D-E** and **Fig.3.3.B.S.2**. The C-strands form the head-to-head dimeric IM, which, in contrast to cases **2** and **5**, contains 12 cytosine pairs, divided in half by the fragment containing adenines. In this variant of the IM, all cytosines form the C-C pairs. From the analysis of the data presented in **Fig. 3.3.B.S.3**, it can be concluded that the IM structure demonstrated stability during the dynamics. The next group of possible options of the geometry of arrangement of non-canonical forms relative to each other in the variant of contacts of 2 duplexes is formed by cases in which G-strands do not form stacks of G4s.
9. **“Head-to-tail IM-dimer between two parallel G4-monomers” (mq-hti-mq)** is demonstrated on **Fig.3.3.C** and **Fig.3.3.C.S.1**. In this option, as in previous cases **2** and **5**, the C-strands form the head-to-tail dimeric IM, but unlike the mentioned cases, it is located localized between the parallel G4s formed by the G-strands. In this case, the COMs of all non-canonical structures are located on a straight line passing through these COMs. Also, as in cases **2** and **5**, the IM contains 10 cytosine pairs, divided in half by the area containing adenines. Comparison of the initial conformation and the one obtained at the last step of the trajectory, as well as the analysis of the behavior of the curve **QIQII**, see **Fig.3.3.C.S.2.C**, indicates that the G4s were located in the starting conformation at 90 degrees angle tend to be located parallel to each other during the calculations. Analysis of the parameter values given in **Fig.3.3.C.S.2-3**, showed that, the G4s and the IM did not undergo deformations during the calculations.
10. **“Head-to-head IM-dimer between two parallel G4-monomers” (mq-hhi-mq)** is demonstrated on **Fig.3.4.A** and **Fig.3.4.A.S.1**. In contrast to the previous case, in this one as in case **8** the G-strands form the dimeric head-to-head IM is localized between the parallel G4s, formed by G-strands. The G4s arrangement in the starting conformation is similar to the previous case, see **Fig.3.3.C** and **Fig.3.4.A**. However, the analysis of the parameters describing the geometry arrangement of the G4s relative to each other: the angle between the axes passing through the tetrads’ COMs (curve **QIQII** in **Fig.3.4.A.S.2.C**), the distance between the COMs of internal tetrads (curve **Iq2IIq2** in **Fig.3.4.A.S.2.E**); indicated that the G4s converged on each other during the calculations and the axes passing through the tetrads’ COMs turned to become parallel to each other in the final conformation, see **Fig.3.4.A.S.1**. **T51** and **T113** thymines belonging to propeller loops can form a stacking interaction. The COMs of all non-canonical structures, as in the previous case, are located in the same plane. Analysis of the data presented in **Fig.3.4.A.S.2-3**, showed the stability of the structure of the G4s and the IM during MD calculations.
11. **“Head-to-head IM-dimer between two parallel G4-monomers in case of strands exchange” (mq-hhi-mq-ex)** is demonstrated on **Fig.3.4.B** and **Fig.3.4.B.S.1**. The localization of non-canonical structures, their geometry, and behavior in the process of MD calculations are identical to case **10** just described. However, unlike case **10**, in this variant, the strands forming the IM are exchanged in the duplexes. As in the previous case, in the process of the MD calculations, the structures of the G4s and the IM demonstrated stability, see **Fig.3.4.B.S.2-3**.
12. **“Two parallel G4-dimers in the same plane and head-to-head IM-dimer between two duplexes with mutual girth of the strands” (2dq-hhi)** is demonstrated on **Fig.3.4.C** and **Fig.3.4.C.S.1**. In this option, the parallel G4s formed by G-strands are located in the same plane, and each strand forms G4’s half. During the transition of the strands from one G4 to another, the sugar-phosphate backbone turns in such a way that the side formed by the strand in one G4 is opposite to the side formed by the same strand in the neighboring G4. As a result, the strands mutualy girth each other, and the angle between the axes passing through the tetrads’ COMs is 180 degrees, and remains so throughout the entire MD trajectory (**QIQII** curve in **Fig.3.4.C.S.2.C**). The head-to-head IM, formed by the C-strands, is the same as in case **8** and **10** and is located parallel to the tetrads of the G4s located in the same plane. As follows from **Fig.3.4.C.S.3**, only two cytosine pairs were subjected to deformation during the MD.
13. **“Two parallel G4-dimers in the same plane and two IM-monomers clamped duplexes with mutual girth of the strands” (2dq-2mi)** is demonstrated on **Fig.3.5.A** and **Fig.3.5.A.S.1**. In this option, the G4s topology and the G4s arrangement relative to each other are the same as in case **12**. But unlike the previous case, the C-strands form monomeric IMs consisting of six cytosine pairs as in previous cases **1**, **3**, **5** – **7**. The IMs are localized between parts of the duplexes, the strands of which are involved in the formation of these IMs. Data analysis in **Fig.3.5.A.S.2-3** indicates that the G4s did not undergo significant changes during the MD, and only one cytosine pair in the IMs turned to be unstable.
14. **“Two parallel G4-dimers in the same plane and head-to-tail IM-dimer between the duplexes with exchange and mutual girth of the strands” (2dq-hti-ex)** is demonstrated on **Fig.3.5.B** and **Fig.3.5.B.S.1**. In this option, the localization and the topology of the G4s formed by G-strands are similar to one in previous cases **12** and **13**. But in contrast to cases **12** and **13** the C-strand form the head-to-tail dimeric IM, similar to those that were formed in cases **2**, **5** and **9**. In addition to mutual girth of the G-strands, in this variant the exchange of the strands in the duplexes also exist. During the MD process, all non-canonical structures showed stability, see **Fig.3.5.B.S.2-3**.
15. **“Two parallel G4-dimers in the same plane with head-to-tail IM-dimer between two duplexes and the strands exchange” (2dq-hti)** is demonstrated on **Fig.3.5.C** and **Fig.3.5.C.S.1**. In this option, in contrast to the cases **12**, **13**, and **14** considered earlier, the halves of the G4s formed by G-strands are localized on one side of the plane containing the vectors connecting the tetrads’ COMs in the G4s. As a result, the strands forming G4s do not mutually girth each other. The value of the angle between the axes passing through the COMs of the tetrads of the G4s fluctuates during the trajectory near the value equal to 90 degrees. C-strands form the dimeric head-to-tail IM, similar to those formed in cases **2**, **5**, **9** and **14**. In this option as in the previous one, the strands exchange take place. As in previous case 14, no deformations occurred in the non-canonical structures during the MD, see **Fig. 3.5.C.S.2-3**.

**Figure 3.1.**
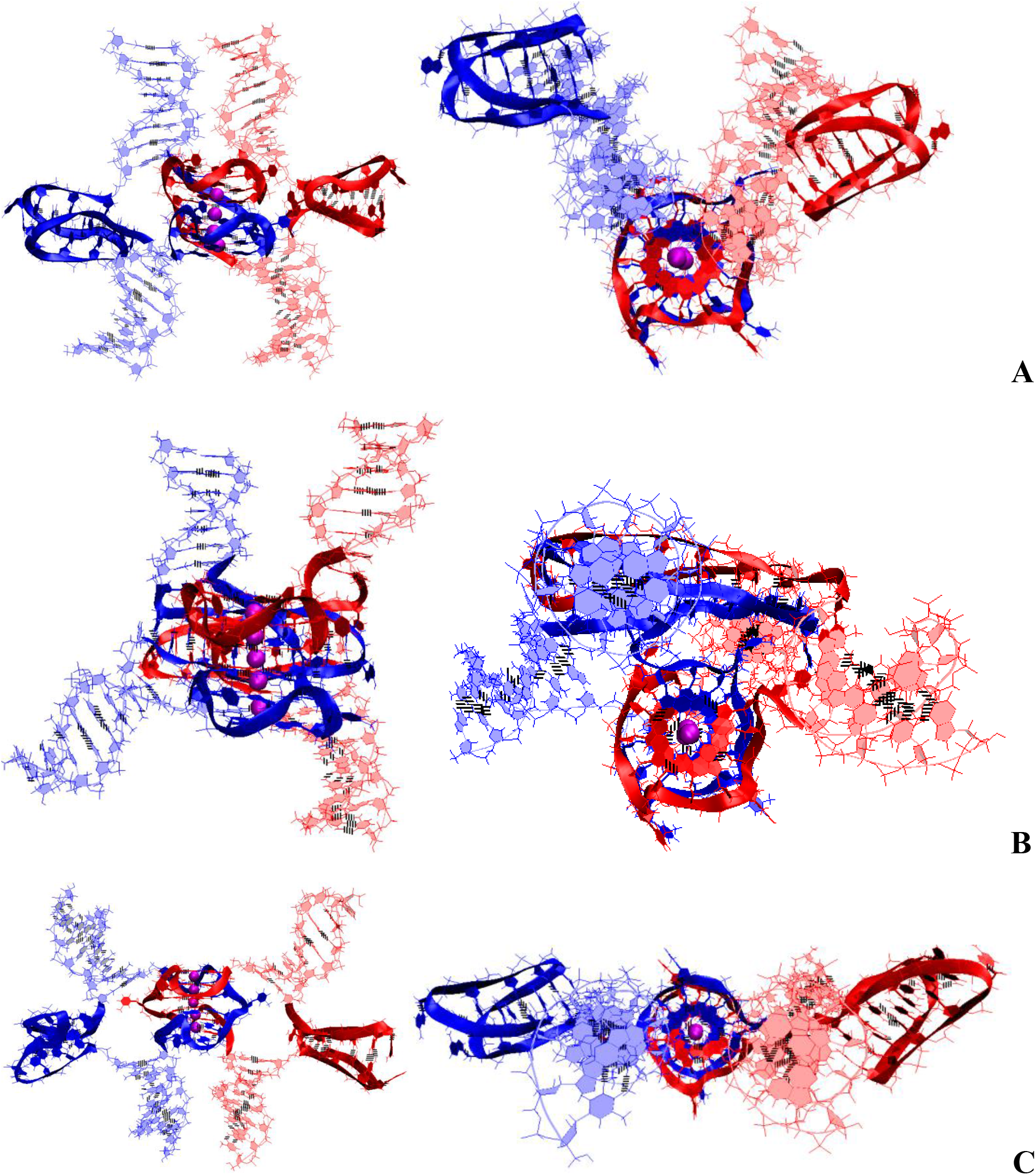
Bimolecular complex of DNA duplexes containing (G_3_T)_3_G_3_ and (C_3_A)_3_C_3_ fragments (starting conformations). **A** – case **1 “1,3 hitch and two IM-monomers”**; **B** – case **2 “1,3 hitch and head-to-tail IM-dimer”**; **C** – case **3 “1,2 hitch and two head-to-head IM-monomers with mutual girth of the strands”**.

**Figure 3.2.**
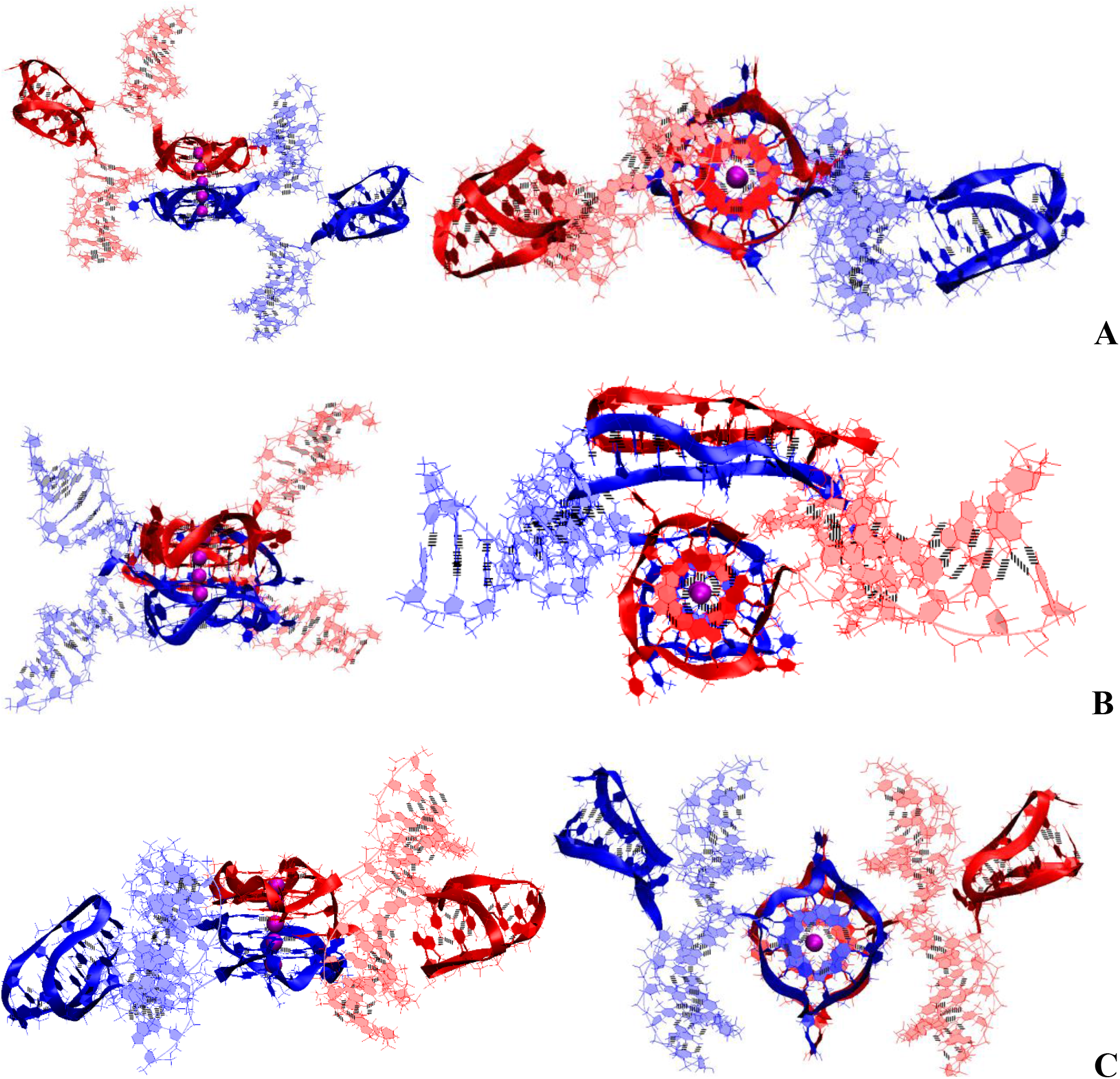
Bimolecular complex of DNA duplexes containing (G_3_T)_3_G_3_ and (C_3_A)_3_C_3_ fragments (starting conformations). **A** – case **4 “Stacking with 2 IM-monomers”**; **B** – case **5 “Stacking and head-to-tail IM**-**dimer”**; **C** – case **6 “Flip-flop interlock and two IM-monomers”**.

**Figure 3.3.**
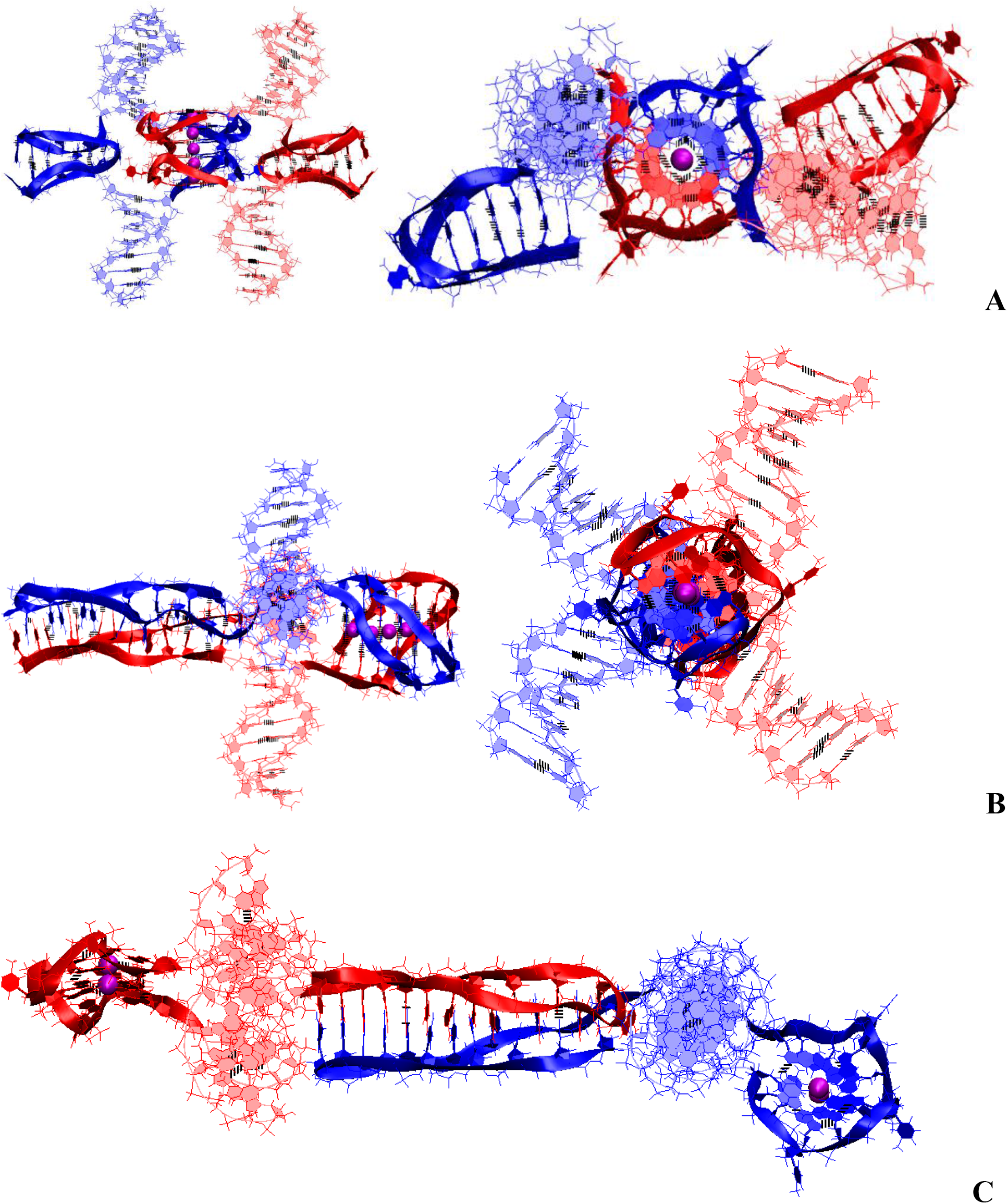
Bimolecular complex of DNA duplexes containing (G_3_T)_3_G_3_ and (C_3_A)_3_C_3_ fragments (starting conformations). **A** – case **7 “Stacking of right and left handed parallel G4-dimers, and two IM-monomers”**; **B** – case **8 “Antiparallel G4-dimer and head-to-head IM-dimer”**; **C** – case **9 “Head-to-tail IM-dimer between two parallel G4-monomers”**.

**Figure 3.4.**
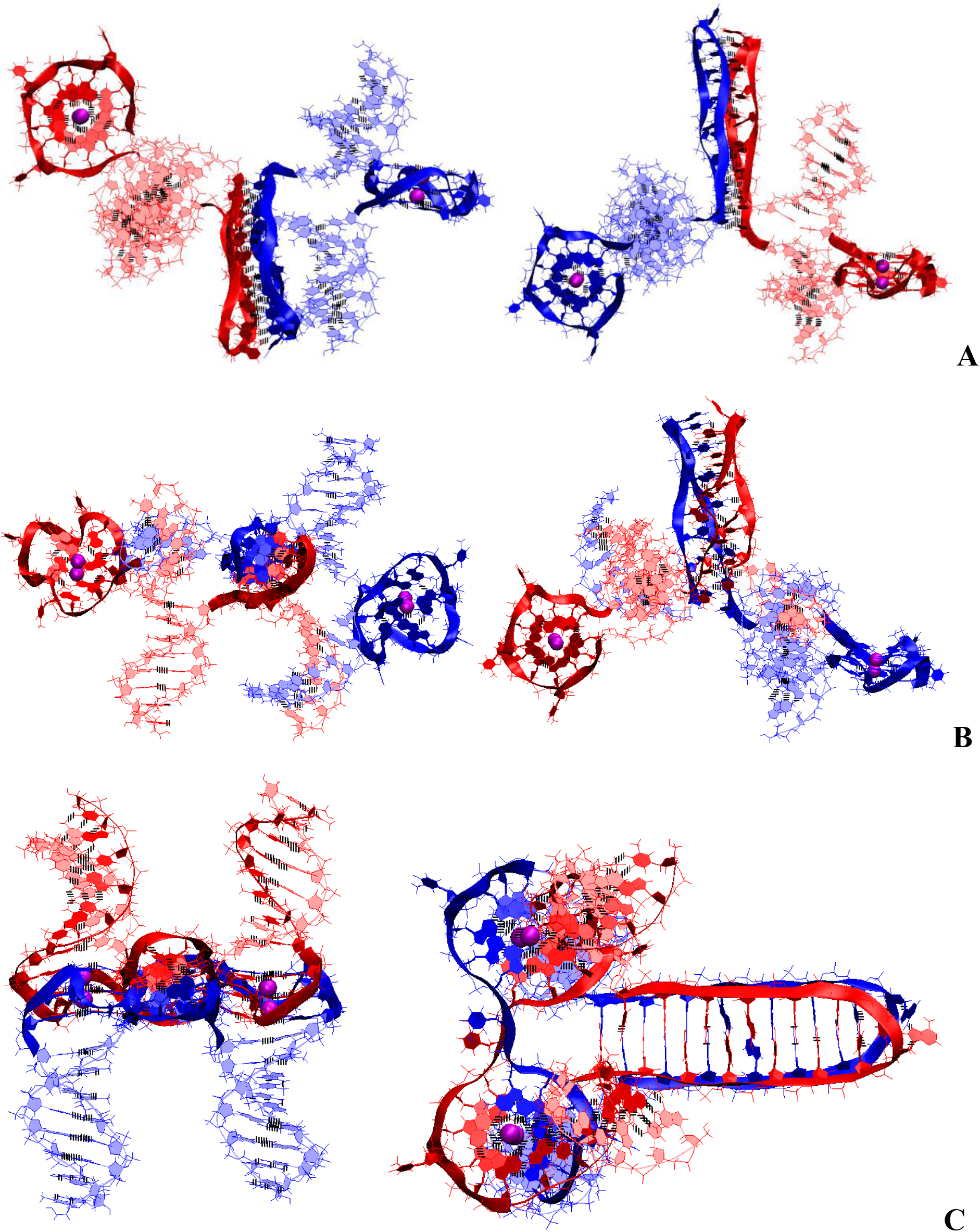
Bimolecular complex of DNA duplexes containing (G_3_T)_3_G_3_ and (C_3_A)_3_C_3_ fragments (starting conformations). **A** – case **10 “Head-to-head IM-dimer between two parallel G4-monomers”**; **B** – case **11 “Head- to-head IM-dimer between two parallel G4-monomers in case of the strands exchange”**; **C** – case **12 “Two parallel G4-dimer in the same plane and head-to-head IM-dimer between two unmelted fragments of duplexes with mutual girth of the strands”**.

**Figure 3.5.**
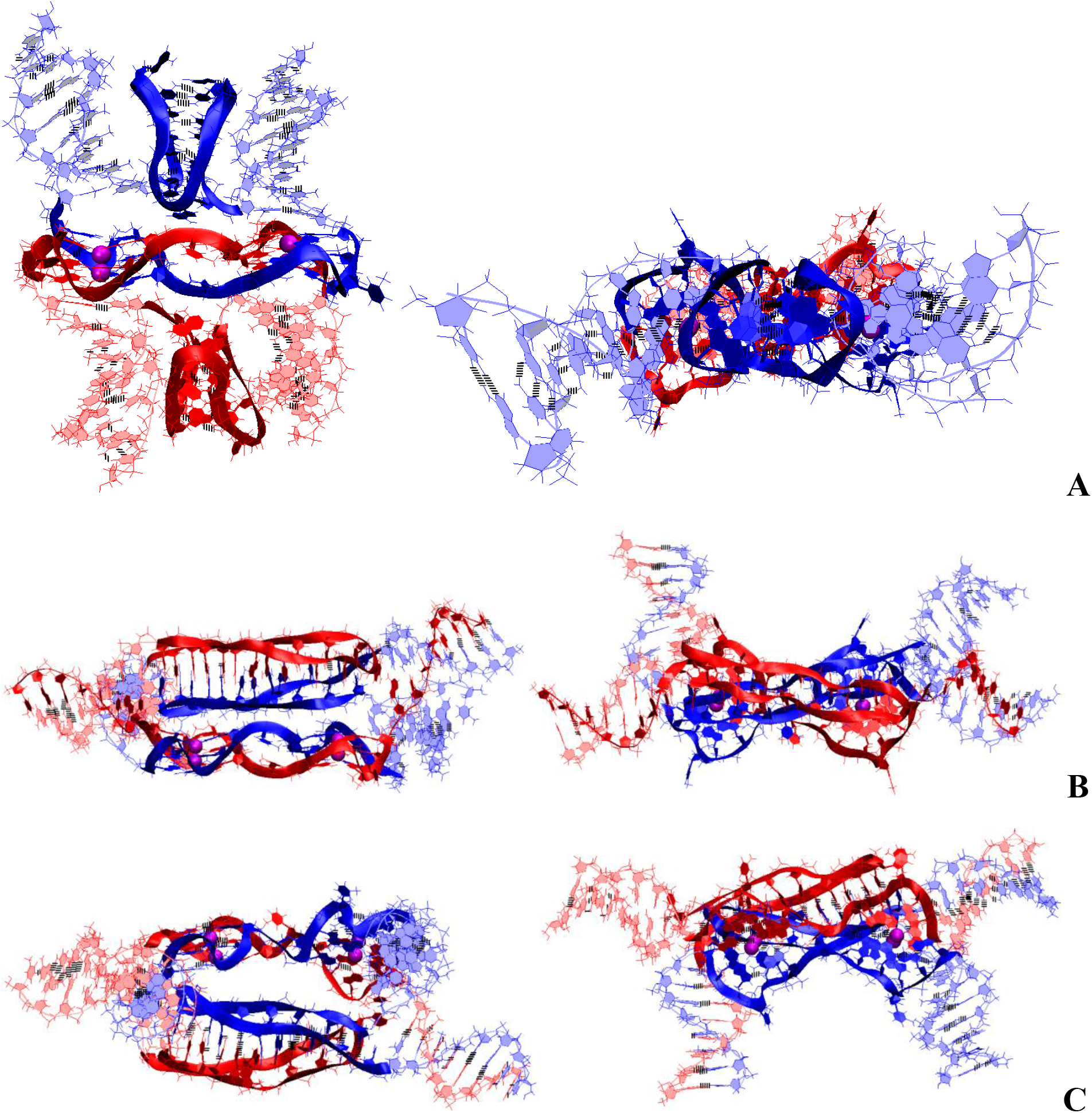
Bimolecular complex of DNA duplexes containing (G_3_T)_3_G_3_ and (C_3_A)_3_C_3_ fragments (starting conformations). **A** – case **13 “Two parallel G4-dimers in the same plane and two IM-monomers clamped unmelted fragments of duplexes with mutual girth of the strands”**; **B** – case **14 “Two parallel G4-dimers in the same plane and head-to-tail IM-dimer between two unmelted fragments of duplexes with exchange and mutual girth of the strands”**; **C** – case **15 “Two parallel G4-dimers in the same plane and head-to-tail IM-dimer between two unmelted fragments of duplexes with the strands exchange”**.

On **Fig.3.S.E** plots of the evolution of the contributions to the free energy for the cases described above for contacts of duplexes containing (G_3_T)_3_G_3_ and (C_3_A)_3_C_3_ are presented. An analysis of the values of the contributions presented in **Fig. 3.S.E** showed that the most significant contribution to the free energy was made by the electrostatic contribution and the polar component of the solvation energy. The lowest value of the sum of these contributions is for case **9**, see on **Fig.3.3.C**, results in the lowest sum of all contributions to free energy for this option. Case **10**, presented in **Fig.3.4.A**, has the lowest value of internal energy. The same variant also has the lowest stress energy; for the other cases, the stress energy values deviate from case **10** by no more than 0.5%. The least contribution to the free energy is made by the non-polar component of the solvation energy, which has small positive values, and the difference in its values in the considered variants is localized within a few kcal/mol. The Van der Waals contribution to the free energy is also insignificant, and the largest one is for case **12** shown in **Fig.3.4.C**.

At a high concentration of DNA, contacts of more than 2 duplexes can occur simultaneously. Below are the possible geometries of the complexes for the case of simultaneous contact of 4 duplexes.

1. **“Four parallel G4-dimers in the same plane and four IM-monomers between the duplexes with exchange and mutual girth of the strands” (4tq-4mi)** is demonstrated on **Fig.4.1** and **Fig.4.1.S.1**. In this option, G-strands form the parallel G4s laying in the same plane. Top view of the G4 part shows that the COMs of the G4s form the square. Each G4 is formed by three strands in the ratio of 2/4:1/4:1/4. Each of the strands that form quadruplexes is arranged in them as follows: forms two of the four G-tracts in the first, then forms one in the next to the first, and then another in the next to the second one. If the G4s’ COMs are located at the vertices of the diagonals, then the G4 axes are unidirectional, and if the G4s’ COMs are located at the vertices of the sides of the square, then the axes are directed oppositely. The C-strands form the monomeric IMs consisting of six cytosine pairs as in the previously considered cases for the contacts of 2 duplexes. The IMs are localized between the parts of the duplexes. In addition at the start the vectors, connecting the COMs of the first and last pair of cytosines of each IMs, are located perpendicular to the plane of the square, formed by the vertices in which the G4s’ COMs are located. During the dynamics, the angles between these vectors and the plane can change so that one of them can become parallel to the plane. Geometry of the IMs localization is such that their heads are directed away from the plane of the square, at the same time the heads of two IMs are oriented in one direction, and the other two are in the other. If the IMs’ COMs are connected by the line segments, then the square will be formed, which will be located in a plane perpendicular to the plane in which the G4s’ COMs are located. The G4s and the IMs are arranged in accordance with the rule: each IM formed by the C-strand is located the closest to the last quadruplex formed by the complementary G-strand. Analysis of the MD results showed, see **Fig.4.1.S.3-4**, that the all G4s are stable, in three IMs one pair has undergone deformation, namely, in two of these three cytosines in the penultimate pair have diverged, and in third IM the last cytosine pair turned to be unstable. The starting localization of the G4s relative to each other during the MD did not undergo significant changes. For two IMs, their positions relative to the G4s has changed. More precisely the slope of the vector, connecting the COMs of the first and last pair of cytosines, has changed so that for one IM the angle between this vector and the normal to the plane, in which the G4s COMS are located, has changed from 0 to 90 degrees, and for the other IM from 180 to 120, see **Fig. 4.1.S.2.E**.
2. **“Four parallel G4-dimers in two stacks and four IM-monomers with exchange and mutual girth of the strands” (4dq-4mi)** is demonstrated on **Fig.4.2** and **Fig.4.2.S.1**. This case differs from the previous one in that G-strands form the dimeric parallel G4s packed in stacks parallel to each other. In this case, the geometry of the stacks is such that the axes passing through the G4s’ COMs included in the stacks are directed oppositely. Each G4 is formed by two strands in the ratio of 2/4:2/4. In this case, one of the strands involved in the formation of the G4 is involved in the formation of the neighboring G4 laying with the first G4 in the same plane, and the other strand is also involved in the formation of another neighboring quadruplex, which is located either under or above the first G4. Thus, two of the strands are involved in the formation of the G4s laying in the same plane, and the remaining two are involved in the formation of the G4s located one above the other. As in the previously considered case **1**, C-strands form the monomeric IMs consisting of six cytosine pairs. In this case, 2 IMs are located above and below the planes in which the G4s’ COMs are located, and the other two are located on the sides of the stacks with the G4s. Those IMs that are located on the sides are formed by strands that are complementary to the strands involved in the formation of the G4s laying in the same plane. Both options are possible only through mutual girthing of the strands, forming G4s, and transition of strands from one duplex to another. For this case the analysis of the MD results showed (see **Fig.4.2.S.3-4**) that the G4s and their geometry relative to each other have not changed. The IMs location in contrast to case **1** did not change significantly during the MD. However, as follows from **Fig.4.2.S.5**, two cytosine pairs in two IMs, and one in the other two, were deformed in the course of calculations.
3. **“Stacking of four parallel G4-dimers with four IM-monomers” (st-4mi)** is demonstrated on **Fig.4.3** and **Fig.4.3.S.1**. In contrast to cases **1** and **2** described above, in this option, when forming the tetrameric complex by sequential stacking of the G4s formed by G-strands, neither mutual girthing of the strands nor the transition them from one duplex to another occur. The formation of the G4s stacking occurs according to the following algorithm. Initially, the complexes described in option **4** for the case with two duplexes are formed. In these complexes, the IMs are opposite each other. Then the pairs of duplexes in these complexes are localized relative to each other so that the line passing through the COMs of IMs included in one pair form a right angle with the same line for the other pair. In this case, the lower G4 of one pair of the duplexes is stacked with the upper G4 of the other. As follow from the analysis of the MD results, see **Fig.4.3.S.3-4**, the G4s, their localization relative to each other have not changed. The localization of the IMs relative to the Stack of the G4s also did not qualitatively change during the MD. However, as follows from **Fig.4.3.S.5**, during the calculations, two IMs were stable, and in each of the other two remaining IMs one cytosine pair was deformed.
4. **“Two parallel stack with right and left handed G4-dimers and two head-to-head IM-dimers” (st-2hhi)**, demonstrated on **Fig.4.4** and **Fig.4.4.S.1**, is one more possible option for the contact of 4 duplexes with the formation of non-canonical forms, in which the strands do not mutually girth and are not transited from one duplex to another. As in case **2,** the G-strands form dimeric the G4s with the ratio of 2/4:2/4, which are packed in stacks arranged parallel to each other. As in case **7** for the two duplexes described above, each stack consists of two parallel dimeric G4s located one above the other, formed half by one strand, half by the other, wherein the lower G4 is right-handed, and the upper one is left-handed. In this option, the axes passing through the tetrads’ COMs are oriented in the opposite directions. The C4-strands form 2 head-to-head dimeric IM. The geometry of the location of the G4s and the IMs relative to each other is as follows: their COMs are located in the same plane and form a rhombus, in which the COMs of stacks of G4s are located at the vertices placed at the ends of the small diagonal, and the IMs’ COMs are located at the vertices of the large diagonal. As in options **8**, **10**, **11**, and **12** described earlier for the case of two duplexes, each of the dimeric IMs is formed from two blocks consisting of 6 cytosine pairs separated by adenine inserts. The location of the IMs relative to each other is such that the vectors connecting the COMs of the first and the twelfth pairs of cytosines are oriented in the opposite directions. From the analysis of the MD results presented in **Fig.4.4.S.3-4**, it can be concluded that the G4s, their localization relative to each other, the geometry of the location of the IMs relative to the stack of the G4s did not change during the MD. As follows from **Fig. 4.4.S.5**, only one cytosine pair in each of the IMs was deformed in the course of calculations.

**Figure 4.1.**
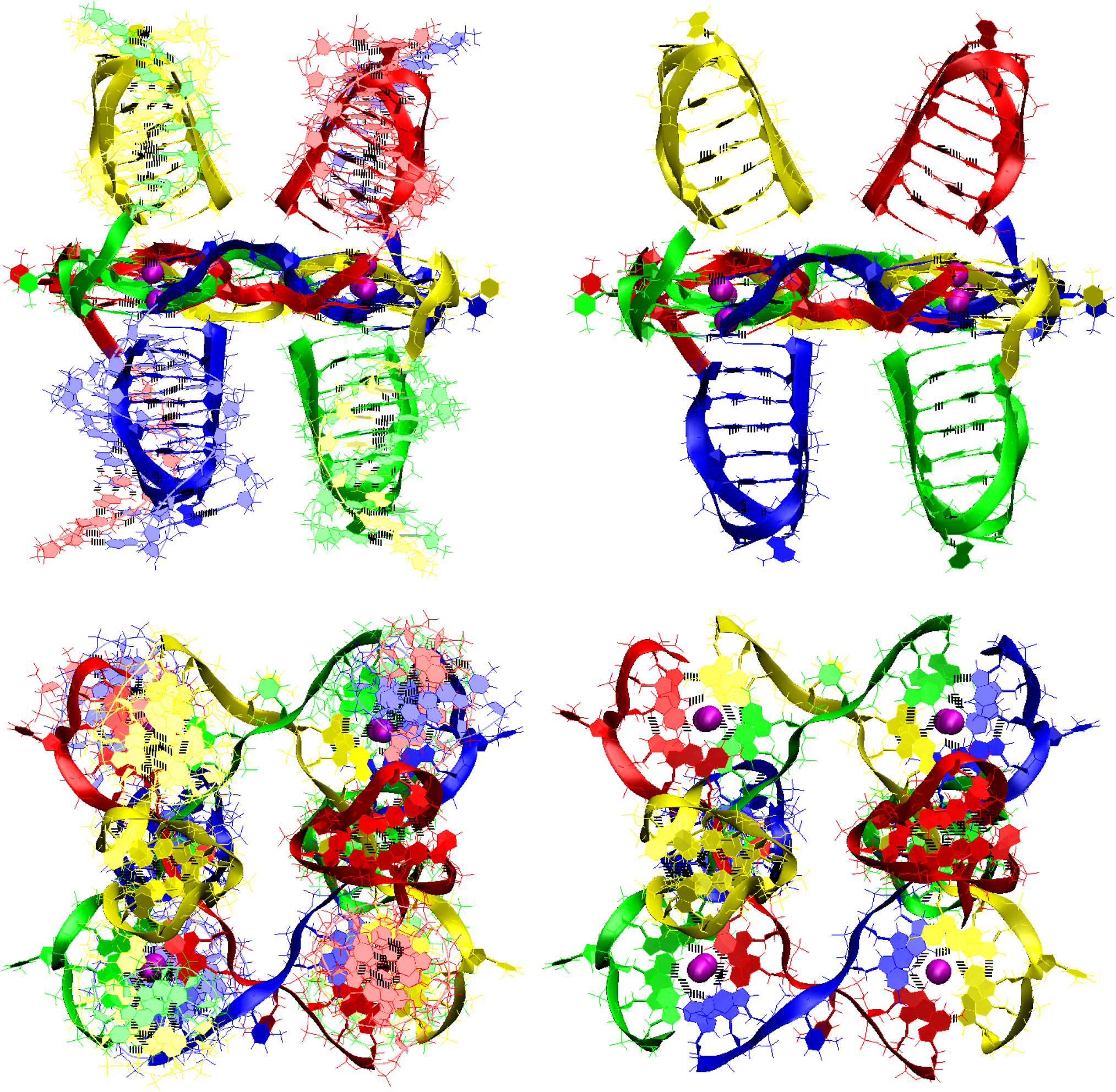
Tetrameric complex of DNA duplexes containing (G_3_T)_3_G_3_ and (C_3_A)_3_C_3_ fragments (starting conformations). Case **1 “Four parallel G4-dimers in the same plane and four IM-monomers between the unmelted fragments of duplexes with exchange and mutual girth of the strands”**. Top row is the side view, bottom row is the top view. On the left in both rows are views with unmelted fragments of the duplexes, on the right there are only G4s and IMs.

**Figure 4.2.**
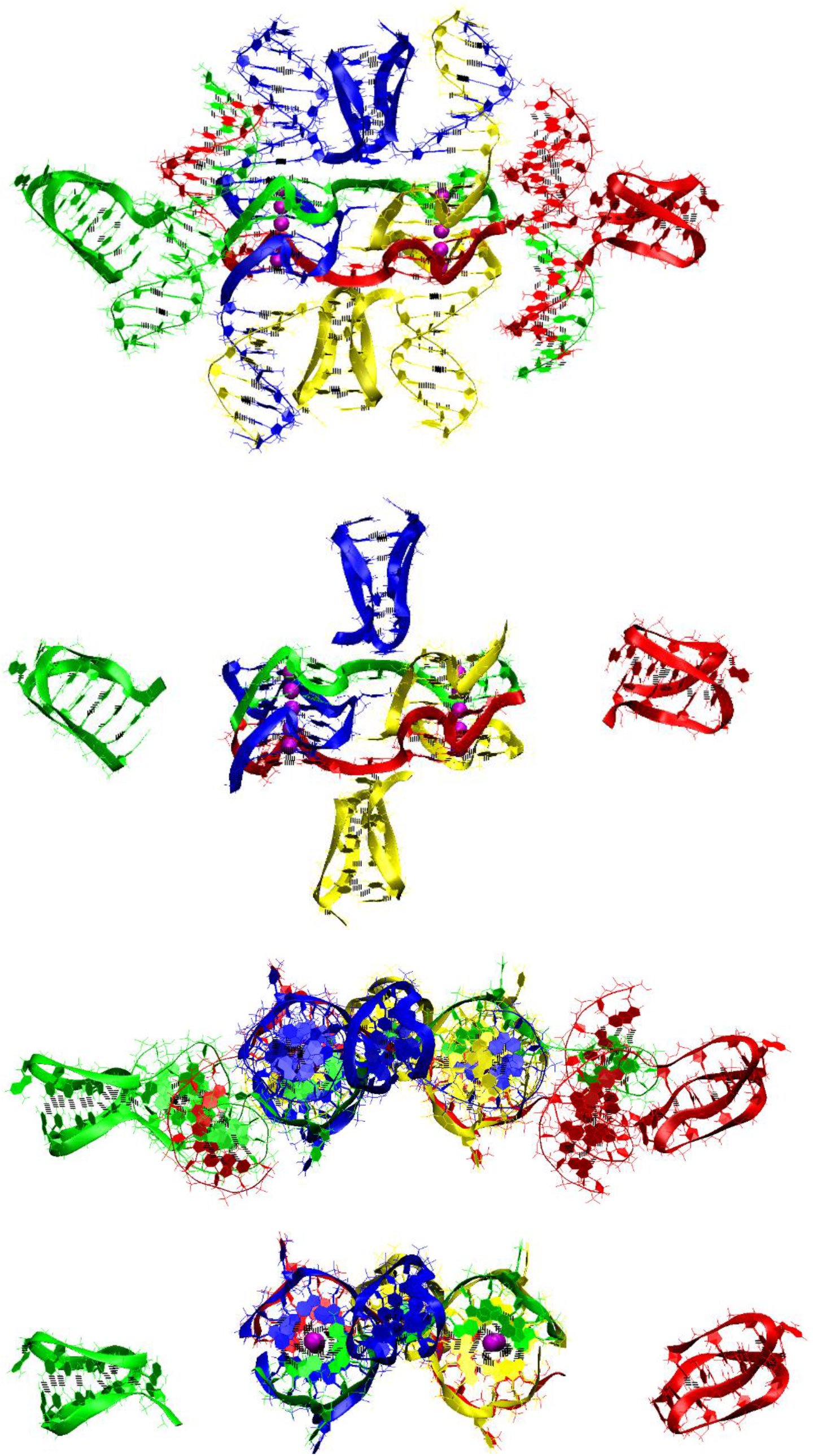
Tetrameric complex of DNA duplexes containing (G_3_T)_3_G_3_ and (C_3_A)_3_C_3_ fragments (starting conformations). Case **2: “Four parallel G4-dimers in two stacks and four IM-monomers with exchange and mutual girth of the strands”:** The first row is the side view with unmelted fragments of the duplexes, the second row is the side view with only G4s and IMs, the third row is top view with unmelted fragments of the duplexes, and the fourth row is top view with only G4s and IMs.

**Figure 4.3.**
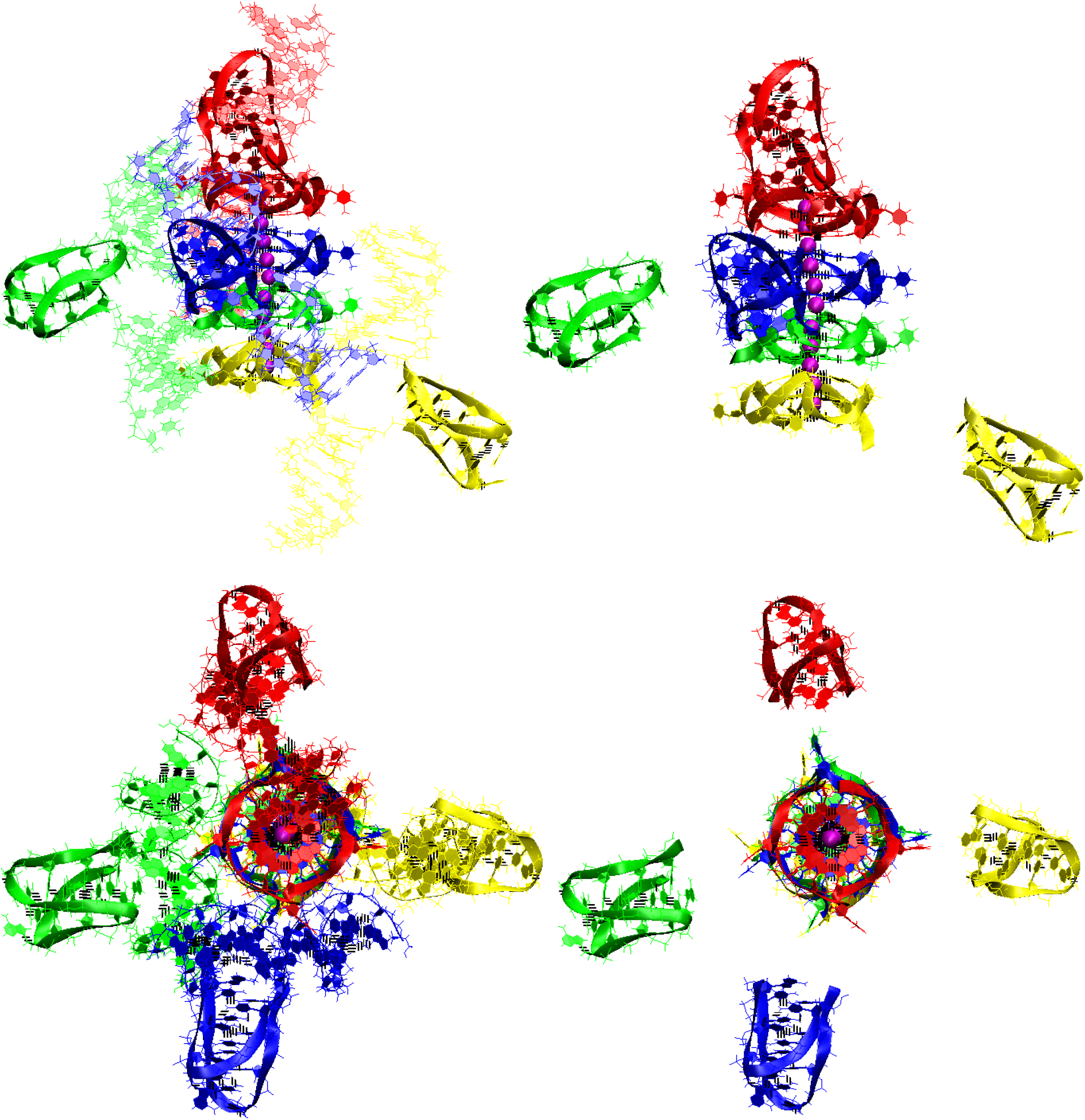
Tetrameric complex of DNA duplexes containing (G_3_T)_3_G_3_ and (C_3_A)_3_C_3_ fragments (starting conformations). Case **3: “Stacking of four parallel G4-dimers and four IM-monomers”: A-** the starting conformation. Top row is the side view, bottom row is the top view. On the left in both rows are views with unmelted fragments of the duplexes, on the right there are only G4s and IMs.

**Figure 4.4.**
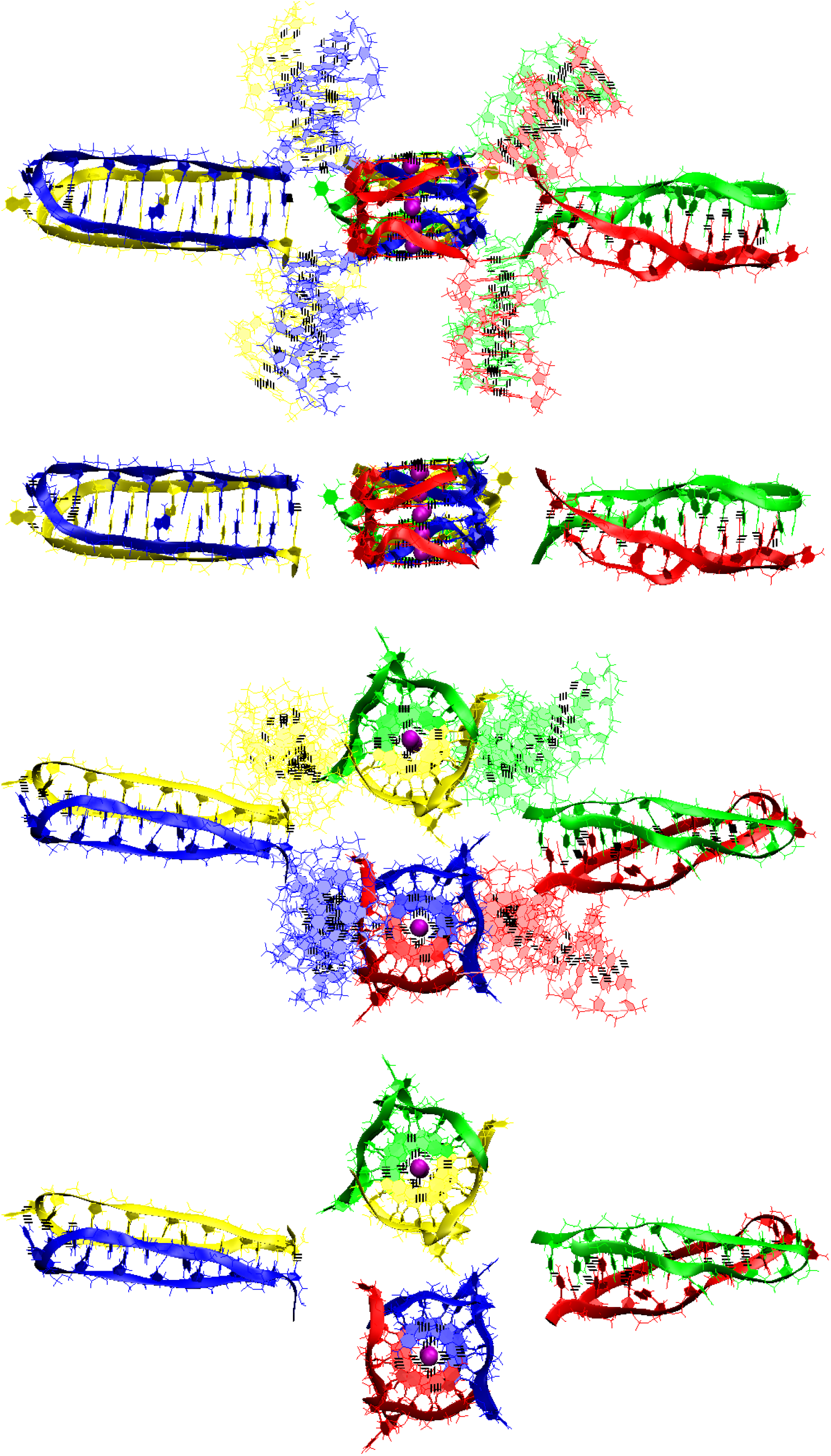
Tetrameric complex of DNA duplexes containing (G_3_T)_3_G_3_ and (C_3_A)_3_C_3_ fragments (starting conformations). Case **4: “Two parallel stack with right and left handed G4-dimers, and two head-to-head IM-dimers”: A-** the starting conformation. The first row is the side view with unmelted fragments of the duplexes, the second row is the side view with only G4s and IMs, the third row is top view with unmelted fragments of the duplexes, and the fourth row is top view with only G4s and IMs.

In **Fig.4.S.E** the graphs of the evolution of the contributions to the free energy are presented in the case of contacts of 4 duplexes containing (G_3_T)_3_G_3_ and (C_3_A)_3_C_3_ fragments. From the data presented in **Fig.4.S.E**, it follows that the lowest value of the sum of contributions to free energy is in option **4**, presented in **Fig.4.4**. This is due to the fact that this option has the lowest value of the sum of the electrostatic contribution and the polar component of the solvation energy, and it is this sum that makes the main contribution to the free energy. Thus, it is this case of the simultaneous formation of G4 and IM forms is most probable upon contact of 4 duplexes containing (G_3_T)_3_G_3_ and (C_3_A)_3_C_3_.

The same variant of the geometry and topology of the G4s and the IMs can be emboded for the contact of 8 duplexes. The embodiment of the 3D model for such the case is shown in **Fig.5** and **Fig.5.S.1**. In this option, already 4 stacks of the dimeric G4s and 4 dimeric head-to-head IMs are formed. In this case, the stacks’ COMs are located at the vertices of the rectangle, and the IMs’ COMs are located at the vertices of the rhombus. These rhombus and square lay in the same plane. The stacks are located between the IMs, and vice versa. During the MD, as follows from **Fig. 5.S.3-6**, the G4s and stacking them were not deformed. Some of the stacks got so close that the thymines from the side loops were able to form a stacking interaction, see **Fig.5.S.1.** From **Fig. 5.S.7-8** it follows that the localization of the IMs did not change significantly, only in two IMs the boundary cytosine pairs were deformed.

**Figure 5.**
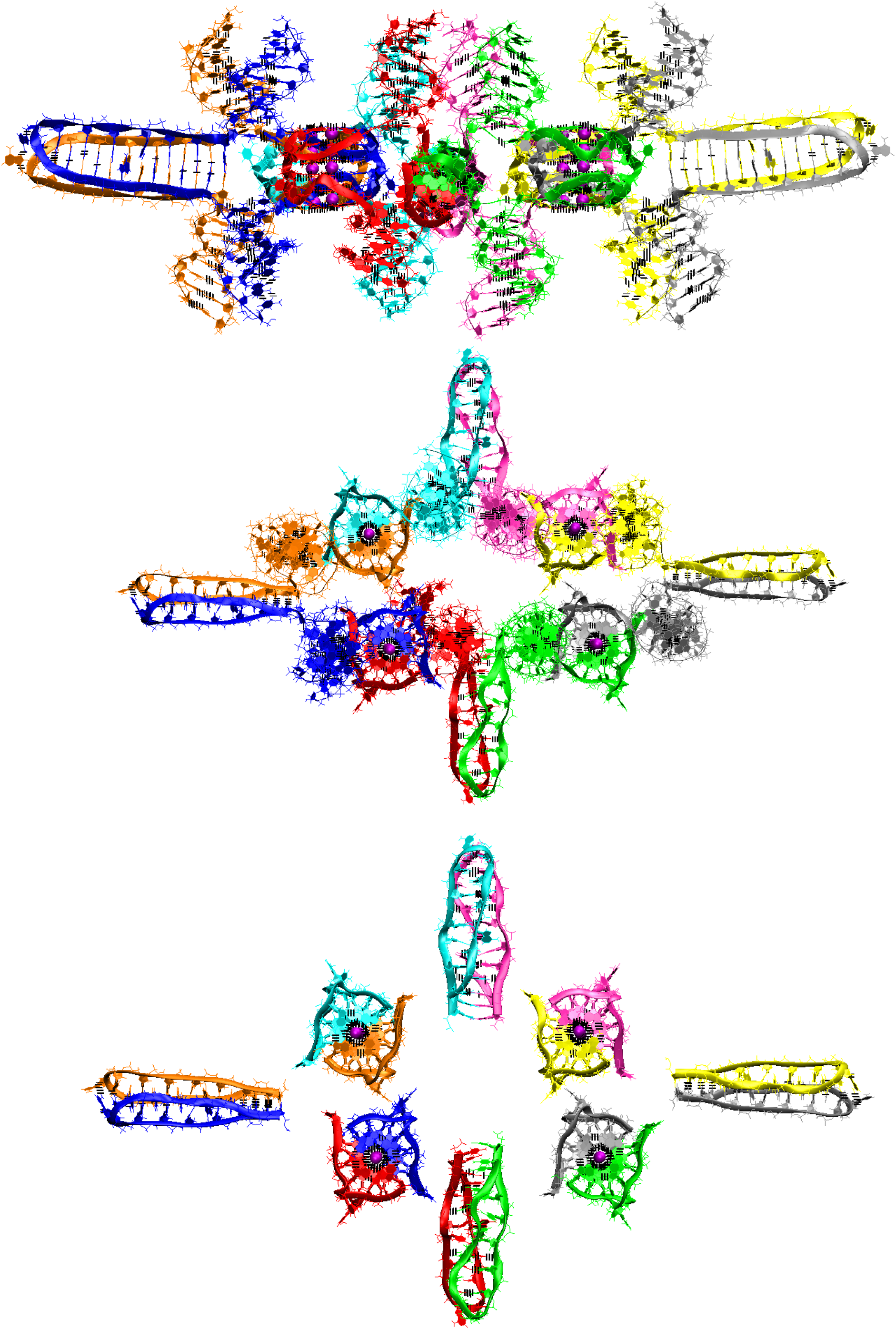
Octameric complex of DNA duplexes containing (G_3_T)_3_G_3_ and (C_3_A)_3_C_3_ fragments (starting conformations). Side view, top view, view with only G4s and IMs.

### Possible variants of the non-canonical structures, in the case of n=4

For n=4, the options considered earlier for n=3 are possible. At the same time, additional GGGT and CCCA motifs are either terminal and are not included in the structures of the G4s and the IMs, or together with the boundary T and A from neighboring GGGT and CCCA form 5 nucleotide loops, where in the G4s case, these have a propeller type. The G4s stack and 2 monomeric head-to-head IMs presented in **Fig.6** and **Fig.6.S.1**. In this option, 5-nucleotide loops both in the G4s and in the IMs are located opposite each other. Thymines in single-nucleotide loops can stack with thymines in 5-nucleotide loops, see **Fig.6.S.1.**B. During the MD, as follows from **Fig.6.S.2-4,** neither of the G4s, their stacking and the IMs were deformed.

**Figure 6.**
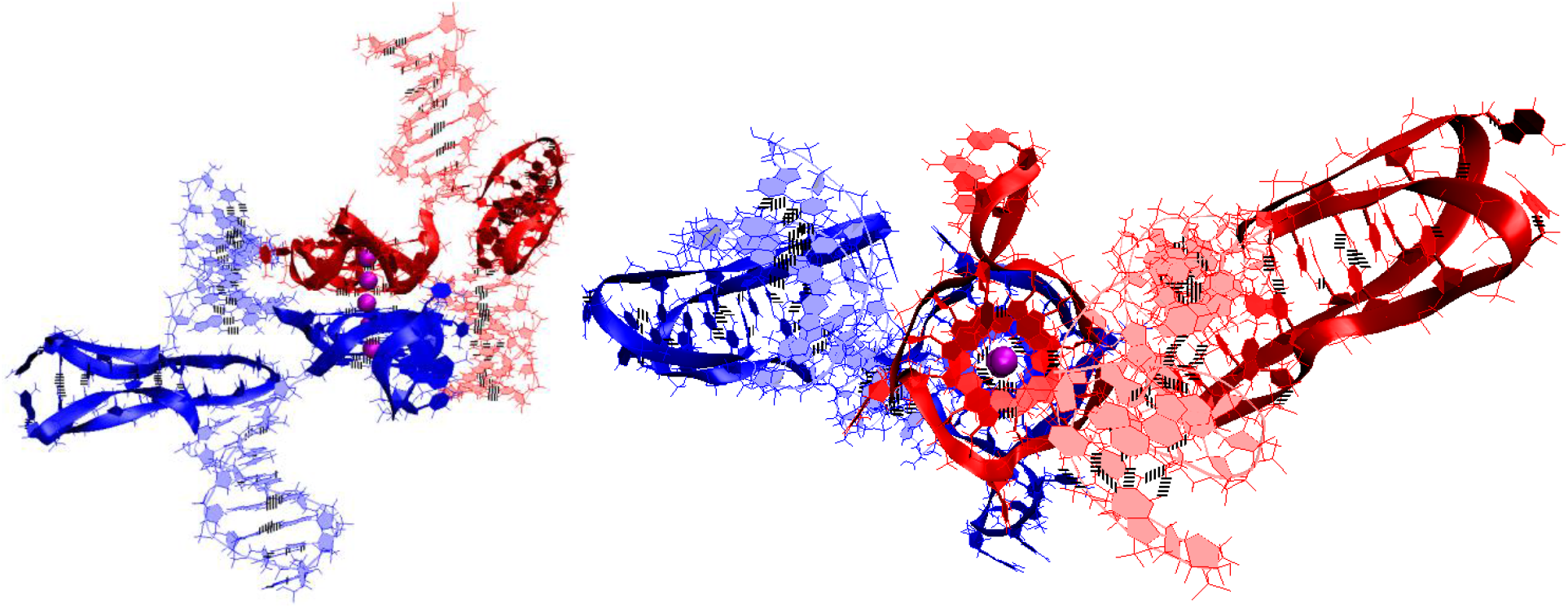
Bimolecular complex of DNA duplexes containing (G_3_T)4G3 and (C3A)4C3 fragments (starting conformations). Stacking with 2 IM-monomers.

### Possible variants of the non-canonical structures, in the case of n=5

1. Let’s start considering possible options with the case **“stacking with two IM-monomers” (st-2mi(6))** which is presented in **Fig.7.1.A** and **Fig.7.1.A.S.1**, and similar to the one discussed above for n=4. As in the case considered for n=4, the monomeric parallel G4s are arranged in the stack, while the lower tetrad of the upper G4 is stacked with the upper tetrad of the lower G4. Each of the G4s has two propeller 5-nucleotide loops, between which there is the propeller single-nucleotide loop. At the same time, 5-nucleotide loops, located one above another, diverge during the MD calculations, and these loops, if you look at the G4s stack from above, look like petals located at an angle of 90 degrees. The IMs located opposite each other are separated from the duplexes by CCCA fragments at each end. During the MD, as follows from **Fig.7.1.A.S.3-4**, the G4s, their stack and the IMs were not deformed.
2. **“1,2 hitch with two IM-monomers with two mini-duplexes” (1,2h-2mi(6))** is presented in **Fig.7.1.B** and **Fig.7.1.B.S.1**. As in case **3** for n=3, the G-strands form the stack consisting of the dimeric parallel G4s. The geometry of the strands forming the G4s is similar to the one considered in option **3** for n=3, with the only difference that two of the propeller loops are 5-nucleotide. But in contrast to the above option **1**, 5-nucleotides are neighboring. As in the considered case **1**, the IMs are monomeric head-to-head, but unlike variant 1, the IMs border on the duplexes at one end, and are separated by C_3_AC_3_ fragment from their other end. At the same time, the cytosines of the CCCA fragment, the closest to the IM, form a mini-duplex with guanines in the 5-nucleotide loop, the closest to the IM. **Table 1 (**Appendix to **Fig.7.1.B.S.4)** presents data of the percentage of MD trajectory snapshots during which the presence of hydrogen bonds, formed by G-C pairs in the mini-duplexes, was observed, and **Fig.7.1.B.S.4.K** shows the evolution of the number of hydrogen bonds in the mini-duplexes. Analysis of the data, presented in Table 1 and in **Fig.7.1.B.S.4.K,** shows that both mini-duplexes were preserved during the first 10 ns, then one of them demonstrated instability as a result of thermal fluctuations. During the MD, as follows from **Fig.7.1.B.S.3-4**, the G4s’ structures were unchanged, the angle of rotation of one G4 relative to the other changed by 15 degrees, and boundary pairs in the IMs were undergone deformations.
3. **“Stacking of right and left handed parallel G4-dimers with two IM-monomers and 4 mini-duplexes” (rl-2mi(6))** is presented in **Fig.7.2.A** and **Fig.7.2.A.S.1**. In this option, the geometry of the location of the G4s relative to each other and their topology is the same as in the previously considered case 7 for n=3 with 2 duplexes in contact. Namely, G-strands form with a ratio of 2/4:2/4 two parallel dimeric G4s located one above another, one of which is right-handed at the bottom and the other is left-handed at the top. At the same time, G-repeats **2** and **5** from (G_3_T)_5_G_3_ fragments with neighboring thymines form the 5-nucleotide propeller loops. The C-strands form the monomeric IMs located opposite each other, consisting of six cytosine pairs. As in option **1**, the IMs are separated from duplexes by CCCA fragments at each end. These CCCA fragments form the mini-duplexes with the 5-nucleotide propeller loops. **Table 2** (Appendix to **Fig.7.2.A.S.4**) presents data of the percentage of the MD trajectory snapshots, during which the presence of hydrogen bonds, formed by G-C pairs in the mini-duplexes, was observed, and the evolution of the number of hydrogen bonds in the mini-duplexes is shown on **Fig.7.2.A.S.4.K-L**. Analysis of the data presented in **Table 2** and in **Fig.7.2.A.S.4.K-L** shows that all mini-duplexes are preserved throughout the entire computation time. During the calculations, as follows from **Fig. 7.2.A.S.4.I-J**, only one cytosine pair from each IM was changed.
4. **“Three parallel G4-dimers in the same plane with head-to-tail IM-dimer with the strands exchange” (3dq-hti)** is presented in **Fig.7.2.B** and **Fig.7.2.B.S.1**. As in the previously described case **15** for n=3, the G-strands form the parallel dimeric G4, the COMS of which lay on the same straight line. Each G4 in this case is formed by two strands in the ratio of 2/4:2/4. The analysis of the angles between neighboring vectors connecting the tetrads’ COMs, see curves **QIQII and QIIQIII** on **Fig.7.2.B.S.2.C**, shows that during the MD calculations the values of these angles fluctuated mainly around 45 degrees. The structures of the first and the third G4s were stable, and in the second G4 the guanine of one of the boundary tetrads moved slightly away from the tetrad’s COM, see **Fig.7.2.B.S.3** and **Fig.7.2.B.S.4.I-J**. The C-strands form the head-to-tail dimeric IM containing 3 fragments with 5 cytosine pairs each, separated by the parts containing adenines and cytosines, located so far away from each other that they cannot form hydrogen bonds, see diagram **Fig.7.2.B.S.2.B**. This variant is possible only on the condition of exchange the strands in the duplexes. As follows from **Fig.7.2.B.S.4.K-L**, during the MD, the IM structure remained unchanged.
5. **“Stacking of three parallel G4-dimers” (st-3dq)** is presented in **Fig.7.3.A** and **Fig.7.3.A.S.1**. The G-strands form three parallel dimeric G4s located one above another with the ratio of 2/4:2/4. The vectors connecting the tetrads’ COMs are unidirectional for all G4s. The C-strands wind round the stack formed by the G4s. During the MD, no changes occurred in the G4 stack, see **Fig.7.3.A.S.3-4**.
6. **“Stacking of two parallel G4-monomers and G4-dimer, and two IM-monomers” (mq-dq-mq-2mi)** is presented in **Fig.7.3.B** and **Fig.7.3.B.S.1**. As in case **5**, the G-strands form the stack of the parallel G4s located one above another. But unlike the previous variant, the boundary G4s are monomers, and the middle one is formed from two strands in the ratio 2/4:2/4. As in variant **1** considered above, C-strand form two monomeric IMs separated from duplexes by CCCA segments at each end. During the MD, as follows from **Fig.7.3.B.S.3-4**, the G4s’ stack did not change, one cytosine pair was subjected to deformation in each of the IMs.
7. **“Stacking of three parallel G4-dimer and two IM-monomers with mutual girth of the strands” (st-3dq-2mi)** is presented in **Fig.7.4.A** and **Fig.7.4.A.S.1**. As in case **5**, the G-strands form three parallel dimeric G4s located one above another with the strands’ ratio of 2/4:2/4. However, in contrast to case **5**, C-strands form the monomeric IMs as in cases **1** and **6** considered above. This option is possible only in the case of the G-strands mutual girthing during the G4s formation. As well as in case **6**, as follows from **Fig. 7.4.A.S.3-4**, during MD only 2 cytosine pairs from the IMs turned out to be unstable.
8. **“Three parallel G4-dimers in the same plane and two IM-monomers with mutual girth of the strands” (3dq-2mi)** is presented in **Fig.7.4.B** and **Fig.7.4.B.S.1**. In this option, the G-strands form the dimeric parallel G4s, the COMs of which lay in the same plane. The G4s’ COMs are located at the vertices of the triangle. Two of the three G4s are formed in the ratio of 3/4:1/4, i.e. 3 of 4 the G-tracts that form the G4 belong to one strand, and the rest of another one. The third G4 is formed half by one strand, half by the other. The vectors connecting COMs of the tetrads of the G4s formed with the ratio 3/4:1/4 are oriented in one direction, and the third G4 in the opposite direction. During the MD, the G4s structures demonstrated stability, and the localization of the G4s relative to each other did not change qualitatively, only the distances between the G4s changed, see **Fig.7.4.B.S.3** and **Fig.7.4.B.S.4.I-J.** The C-strands form the monomeric IMs as in case **1**, **6**, and **7** considered above. The IMs are placed on the opposite sides of the plane in which the G4s’ COMs are located. At the same time, their end parts are directed towards each other. In each of the IMs, one cytosine pair was deformed during the MD calculations, see **Fig.7.4.B.S.4.K-L**.

**Figure 7.1.**
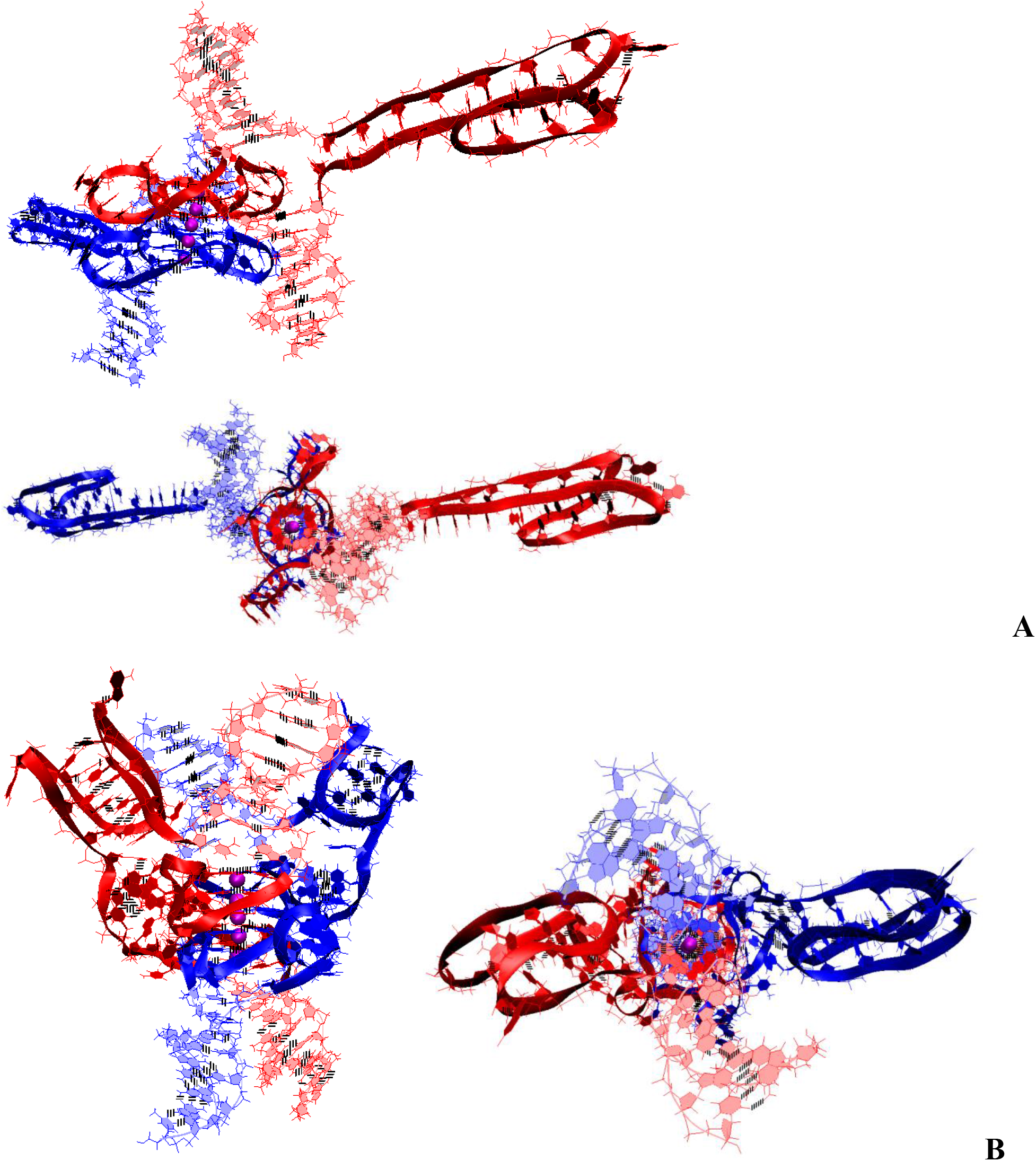
Bimolecular complex of DNA duplexes containing (G_3_T)_5_G_3_ and (C_3_A)_5_C_3_ fragments (starting conformations). **A** – case **1 “Stacking and two IM-monomers”**; **B** – case **2 “1,2 hitch and two IM-monomers with two mini-duplex”**.

**Figure 7.2.**
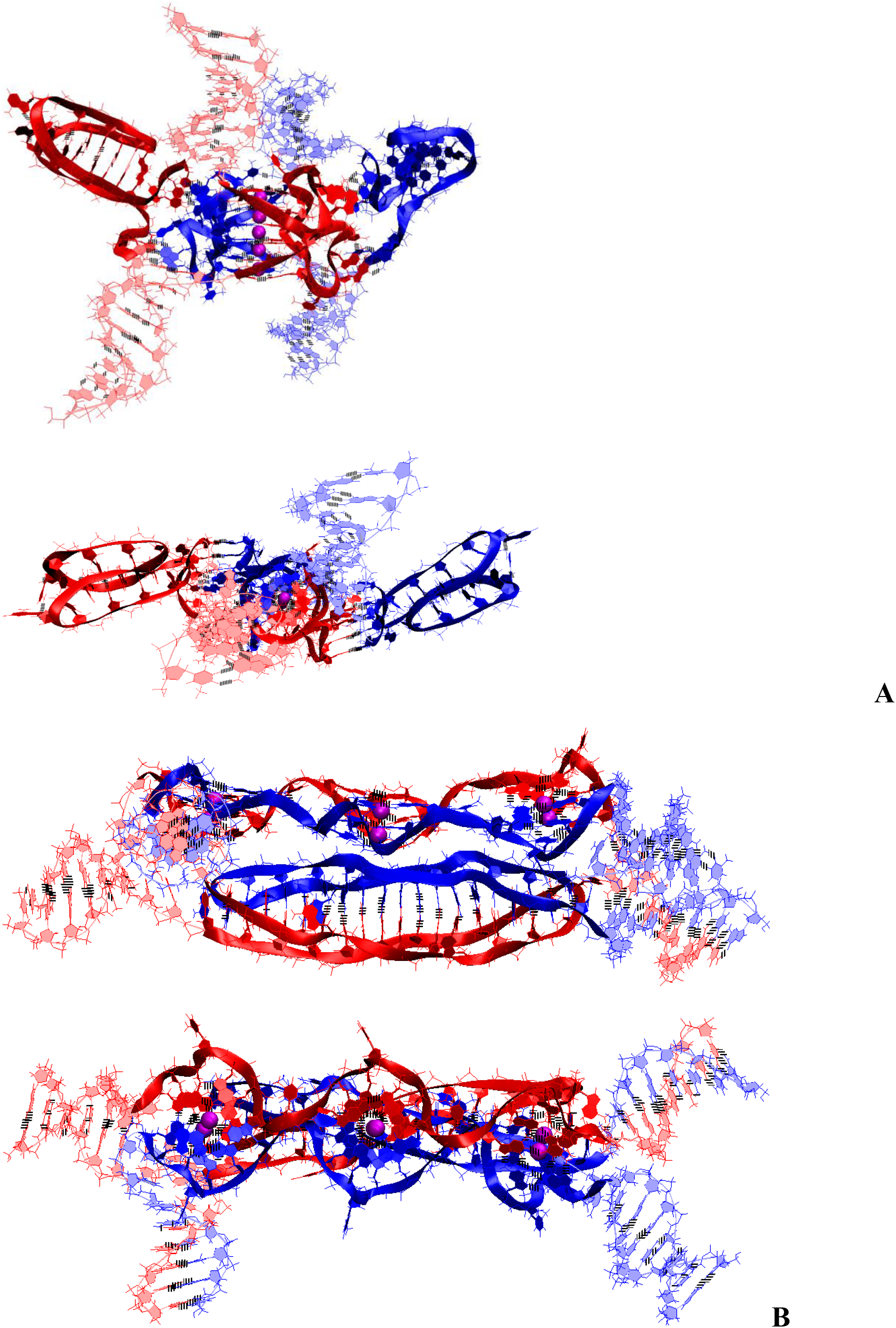
Bimolecular complex of DNA duplexes containing (G_3_T)_5_G_3_ and (C_3_A)_5_C_3_ fragments (starting conformations). **A** – case **3 “Stacking of right and left handed parallel G4-dimers, and two IM-monomers with 4 mini-unmelted fragments of duplexes”**; **B** – case **4 “Three parallel G4-dimers in the same plane and head-to- tail IM-dimer with the strands exchange”**.

**Figure 7.3.**
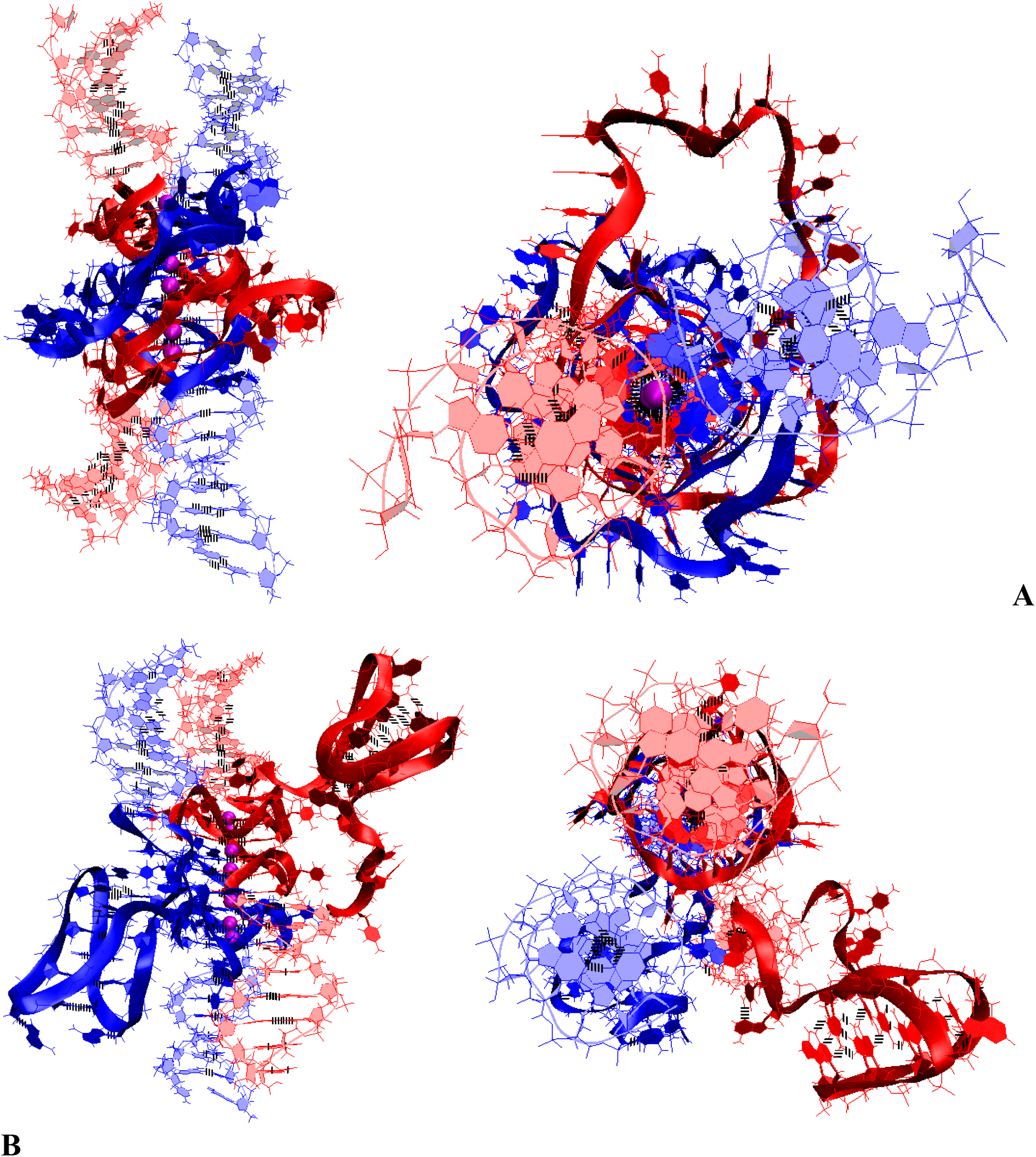
Bimolecular complex of DNA duplexes containing (G_3_T)_5_G_3_ and (C_3_A)_5_C_3_ fragments (starting conformations). **A** – case **5 “Stacking of three parallel G4-dimers”**; **B** – case **6 “Stacking of two parallel G4-monomers and G4-dimer, and two IM-monomers”**.

**Figure 7.4.**
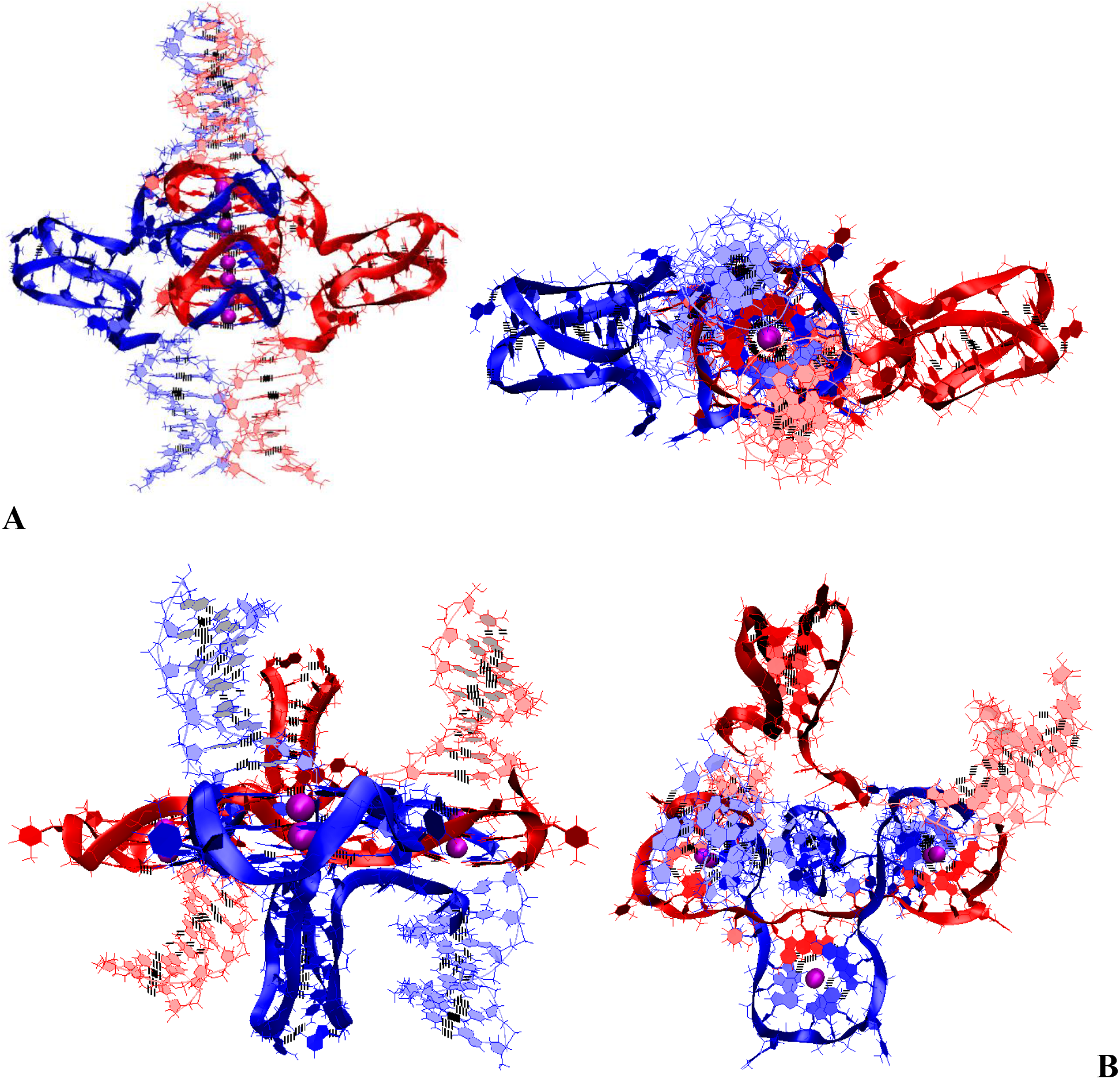
Bimolecular complex of DNA duplexes containing (G_3_T)_5_G_3_ and (C_3_A)_5_C_3_ fragments (starting conformations). **A** – case **7 “Stacking of three parallel G4-dimer and two IM-monomers with mutual girth of the strands”**; **B** – case **8 “Three parallel G4-dimers in the same plane with and two IM-monomers with mutual girth of the strands”**.

On **Fig.7.S.E** plots of the evolution of the contributions to the free energy for the options described above in case of contacts of 2 duplexes, containing (G_3_T)_5_G_3_ and (C_3_A)_5_C_3_, are presented. From the data presented in **Fig.7.S.E**, it follows that the lowest sum of all contributions to free energy in case **5** presented on **Fig.7.3.A**. In this option, C-strands do not form IM structures. From all the cases containing both non-canonical forms, as follows from **Fig.7.S.E**, the most probable are cases **4** and **8** presented on **Fig.7.2.B** and **Fig.7.4.B** respectively. In these cases, the G4s are located in the same plane. In case 4, the strands are exchanged, in case 8, chains are mutually girthed. Amongst the cases without exchange and mutual girthing, the lowest value of the sum of contributions to free energy was observed in case **1**, presented in **Fig.7.1.A**. Without taking into account solvation, case **4** has the lowest internal energy. The same case also has the lowest stress energy.

### Possible packing of single-stranded DNA with the sequence (G_3_T)_n_G_3_ into G4 forms in case of contact of several strands

The options of the formation of G4s configurations in cases of contacts of (G_3_T)_n_G_3_ fragments from G-strands with n taking values from 1 to 5 have already been considered. It follows from the above that the formation of G4s’ configurations upon contacts of single-stranded DNA with G repeats is possible through the G4s, already formed before the contact, and/or through the G4s formation during the contact. In the first variant, the geometry of the G4s arrangement relative to each other is represented by a stack formed by the monomeric G4s whose boundary tetrads are stacked. In the second variant, dimeric, trimeric, and tetrameric G4s can be formed during the contact of the G-strands. As a result, the localization of G4s relative to each other becomes more diverse due to the fact that the strands can move from one G4 to another. As a result, it is possible to form not only a stack, but also horizontal localization relative to each other, in which the G4s are located in the same plane. An example of a flat shape is a G4s’ tape. If in the stack the G4s’ COMs lay on a straight line that is perpendicular to the planes to which the tetrads locate, in it’ s turn in the tape this straight line is parallel to these planes. The increase in the variety of G4 forms also causes that the G4s can be packed into parallel stacks and tapes linked to each other. Linkage is created by the strands involved in the formation of the G4s from neighboring stacks and/or tapes. **Fig.8** shows examples of tetramers in the form of a stack, interconnected parallel stacks and tapes. The variety also increases due to the fact that when stacked, both right-handed and left-handed G4s can be formed, see **Fig.9**. The possible stacking geometries of G4s with n greater than 6 will represent various combinations based on the options discussed above. For example, **Fig.9** shows a variant of the folding of a dimer and trimer with right-handed and left-handed G4s in the case of contact of single-stranded DNA (GGGT)_7_GGG.

**Figure 8.**
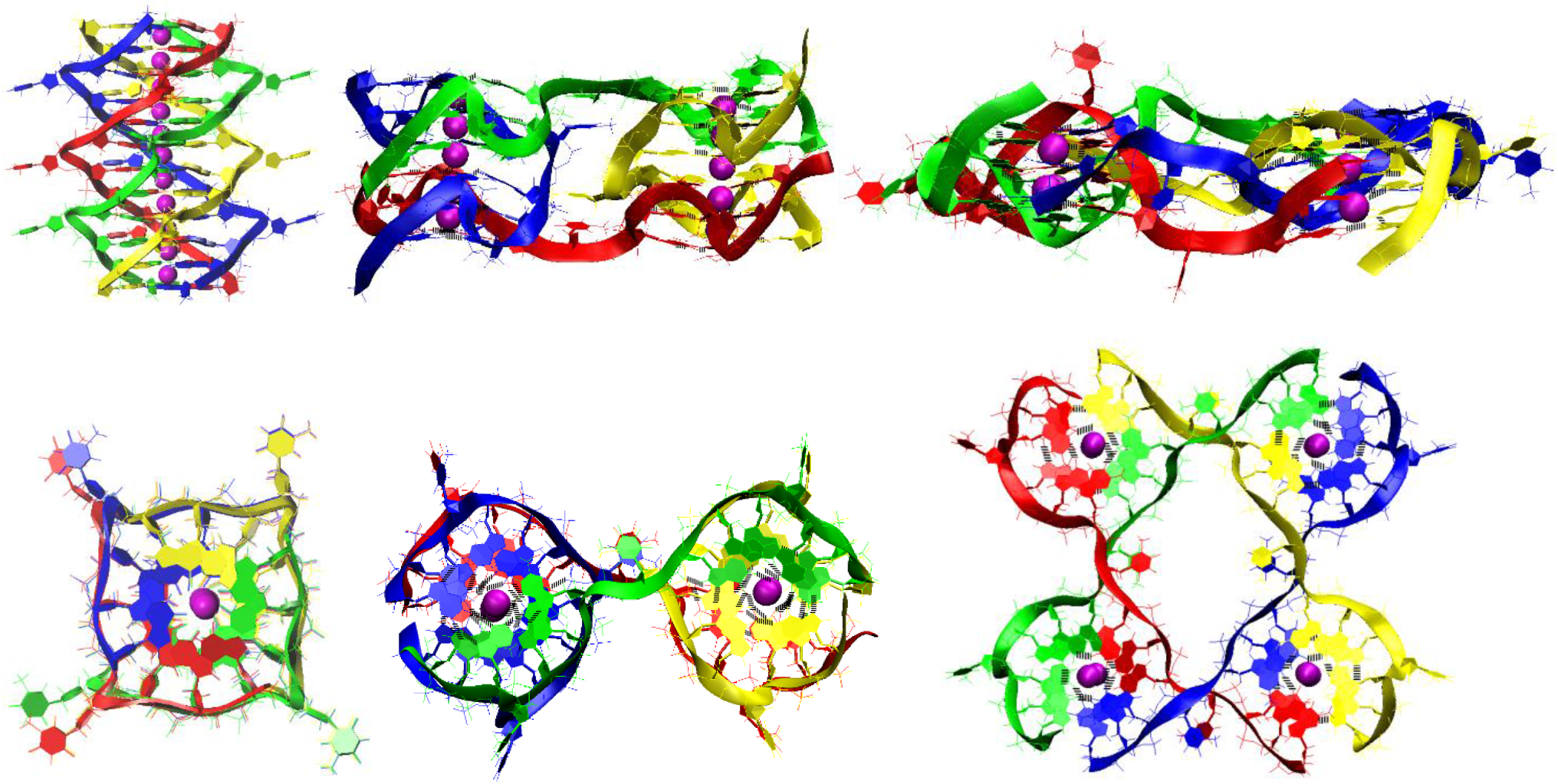
Potential G4 forms formed by DNA strands (G_3_T)_3_G_3_. The top row shows side views; the bottom row shows top view.

**Figure 9.**
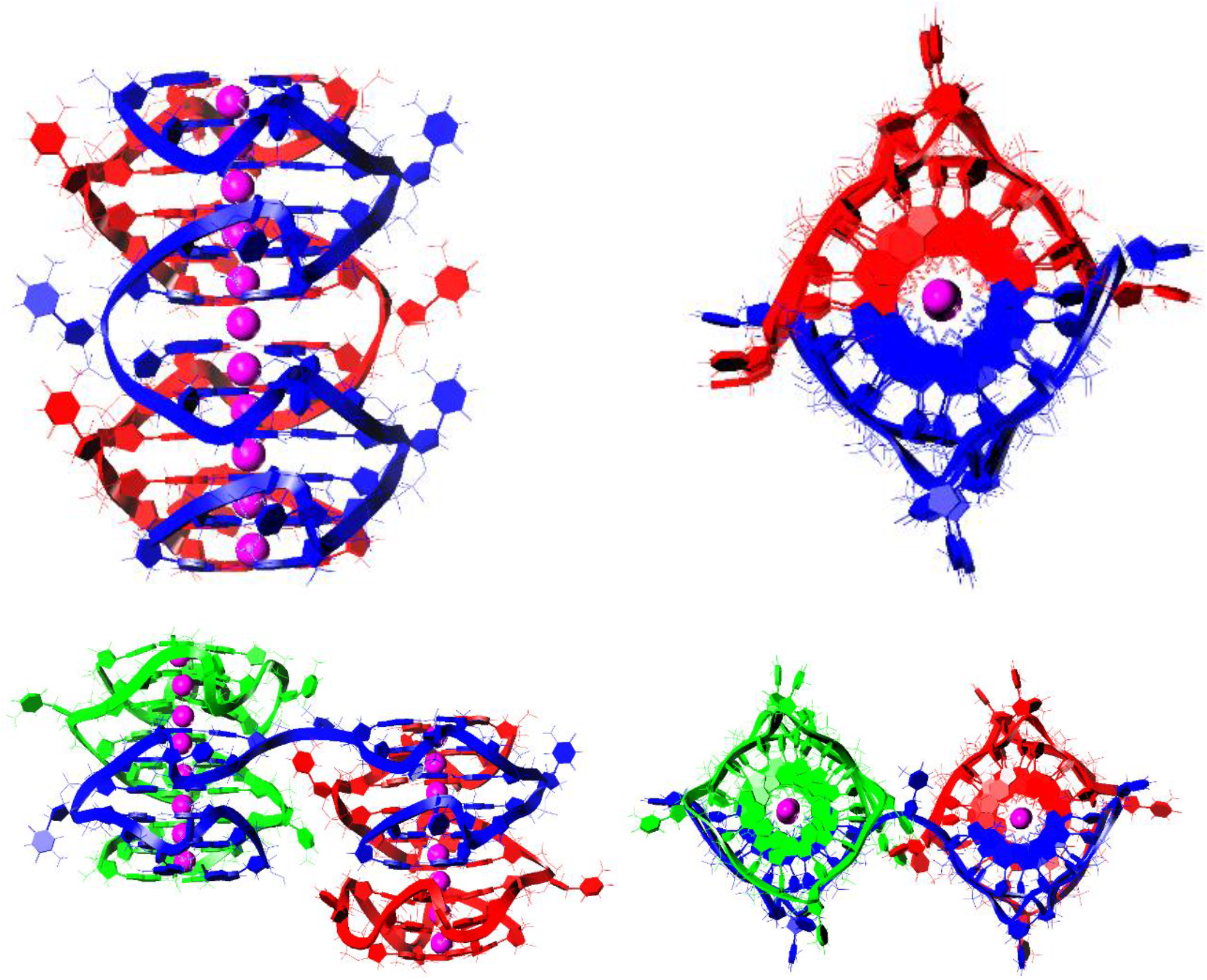
Possible dimer (top) and trimer (botton) formed by DNA strands (G_3_T)_7_G_3_ through the simultaneous formation of right-handed and left-handed G4s. Left - side view, right - top view.

Away from the hydrophobic flat surface, which has a strong affinity to the guanine bases, upon contact of single-stranded DNA with the (GGGT)n sequence, stacks will be formed, since it is the latter that have the smallest number of boundary tetrads, and, as a result, the smallest hydrophobic area available to the polar solvent - water. On the contrary, near the flat surface with a strong affinity to guanine bases, as a result of strands contacts, a G4s’ network can be formed with a higher probability, see Fig. 42, laying on this surface, since it is in this variant the area of contact with the surface will be the largest. A G4s’ network, formed on the surface, can induce the formation of the same on itself, then the process can be repeated over and over again, and eventually a layer can form, see **Fig.10**.

**Figure 10.**
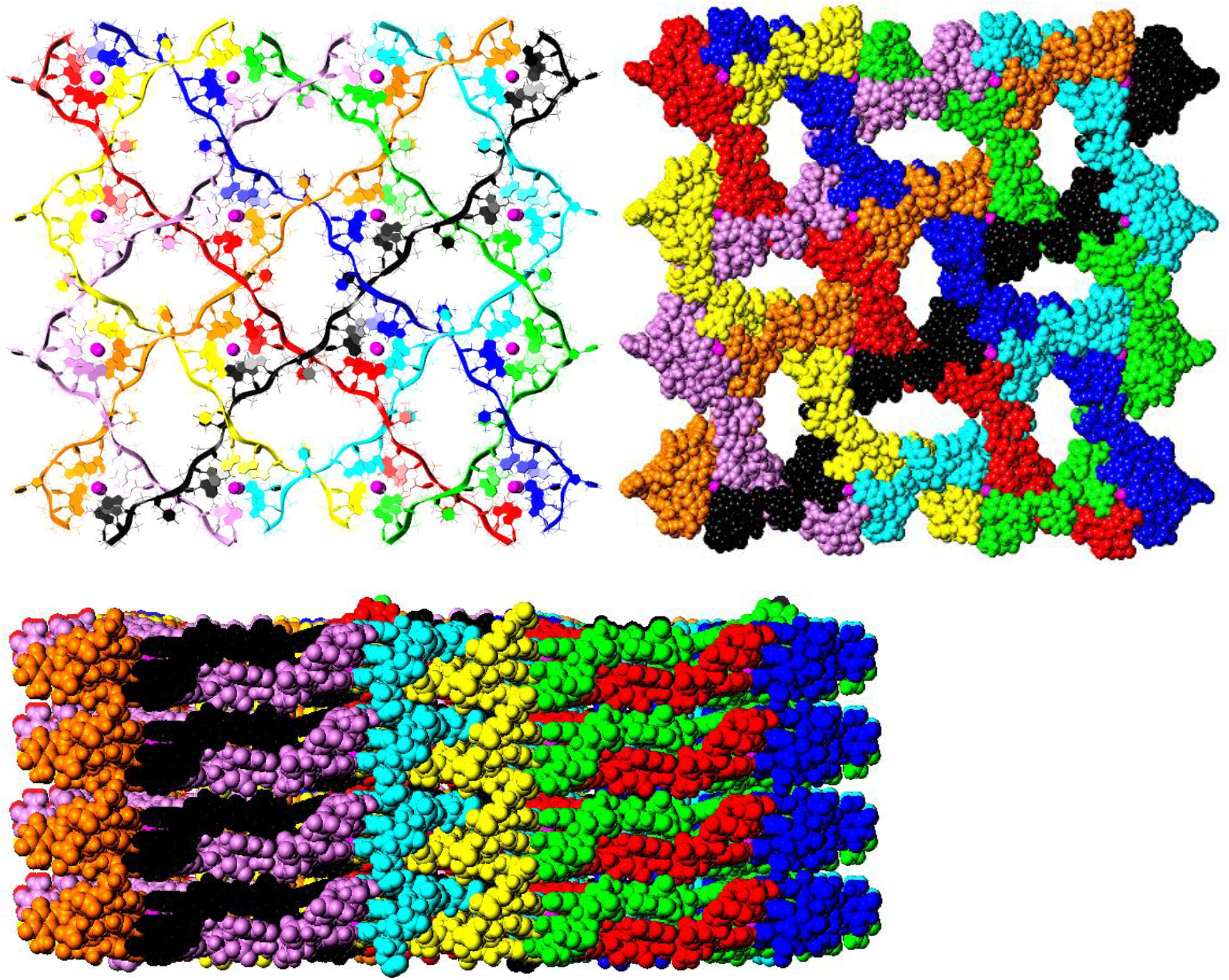
Possible the grid of G4s formed by DNA strands (G_3_T)_7_G_3_ and the layer of the overlaid grids. The top row shows top views; the bottom row shows side view.

## CONCLUSION

The use of the strategy described above for creating 3D models of G4s and IMs forms of DNA and RNA allowed the author to suggest and construct 3D models of possible variants of G4s and IMs forms that can occur during contacts of both single-stranded DNA with sequence (G_3_T)_n_G_3_ and duplexes having (G_3_T)_n_G_3_ and (C_3_A)_n_C_3_ fragments. Testing of the created models using MD and subsequent analysis by means of the parameters described above to evaluate the geometry of the G4s and the IMs showed the absence of deformations resulting in qualitative structural rearrangement of the non-canonical forms. Said above in its turn suggests the possibility of the existence in reality of the forms proposed by the author, in particular: the formation of stacks from right and left-handed G4s, a cruciform structure with antiparallel G4 and IM locating opposite each other - an analogue of the Holiday structure, and others discussed above. Comparison of contributions to free energy made it possible to determine the most probable variants of the simultaneous existence of G4 and IM forms arising at contacts of duplexes having G_3_T and C3A repeats. The proposed model of a layer of G4s’ grids may be a possible DNA origami form construed from G4s. The strategy proposed by the author can allow researchers who are proficient in computer modeling methods to create G4 and IM forms models of DNA and RNA of any complexity in situations where it is necessary.

## EXPERIMENTAL

All 3D models of the studied structures were built using the molecular graphics software package Sybyl-X software (Certara; USA) using the following strategy. Initially, models of the required duplexes, G4s, and IMs were created. Further, the created models were located relative to each other in the required geometry and connected. At each stage, molecular mechanical optimization was performed to eliminate the van der Waals overlap, which could occur during a certain step. The molecular mechanical optimizations were performed using Sybyl-X and Powell’s method with the following settings: parameters for intermolecular interactions and the values of partial charges: taken from force field amber7ff99, non-bonded cut-off distance: 8 Ǻ, effect of the medium: dielectric constant of 4, the number of iterations: 1000, simplex method for initial optimization, and 0.05 kcal*mol^-1^*Å^-1^ energy gradient convergence criterion. The stability of the created models was tested by molecular dynamics using Amber 20 software [8]. The MD simulations in the production phase were performed using constant temperature (T=300 K) and pressure (p=1 atm) over 50 ns. To control the temperature, a Langevin thermostat was used with 1 ps^-1^ collision frequency. Influence of the solvent simulated with the application model of water molecules OPC3 [9]. K+ ions were used to neutralize the negative charge of the DNA backbone. The parameters needed for the interatomic energy calculation were taken from the force fields OL15 [10, 11].

The free energy was calculated as the sum of the electrostatic energies (E_q_), Van der Waals energies (E_VDW_), energy of solvation energy of solvation and deformation energy of valence bonds, valence and dihedral angles (U). The energy of solvation was calculated as the sum of the polar and nonpolar contributions. The polar contribution (E_GB_) was computed using the Generalized Born (GB) method and the algorithm developed by Onufriev et al. for calculating the effective Born radii [12]. The non-polar contribution to the solvation energy (E_surf_), which includes solute-solvent van der Waals interactions and the free energy of cavity formation in solvent, was estimated from a solvent-accessible surface area (SASA).

## ACKNOWLRDGEMENT

The author is grateful to Lilia Yadkarovna Alkhanova for her immeasurable assistance in the work.

## SUPPLEMENTARY MATERIALS

In supplementary, for each considered option in case duplexes, the conformations obtained at the last step of the MD trajectory, the complexes’ schemes, evolution of parameter values for stability scoring of G4s and IMs described in this article, evolution of the contributions to free energy are presented.

