## Supplement for "Modeling of possible quadruplexes and i-motifs formed during DNA contacts: strategy, classification, most probable shapes, origami based on quadruplexes"

**Vladimir B. Tsvetkov** [a, b, *]

[a] Federal Research and Clinical Center of Physical-Chemical Medicine, Moscow 119435, Russia

[b] Institute of Biodesign and Complex System Modeling, I.M. Sechenov First Moscow State Medical University, Trubetskaya Str. 8-2, 119991 Moscow, Russia

| | |
|---|---|
| **Figure 1.A.S. Bimolecular complex with head-to-tail IM-dimer: A** – conformation obtained at the last step of the MD trajectory; **B** – the complex scheme; **C** – angles between unmelted fragments of the duplexes; **D -** distances between COMs of the cytosines' bases; **E -** angles between normals to the cytosines' bases. | 12 |
| **Figure 1.B.S. Bimolecular complex with parallel G4-dimer**: **A** – conformation obtained at the last step of the MD trajectory; **B** – the complex scheme; **C** - distances from COMs of the guanines' bases to COMs of their containing tetrads; **D** – angles between normals to the guanines' bases and vectors connecting COMs of the boundary tetrads; **E** – angles of rotation of the tetrads relative to each other; **F** – angles between unmelted fragments of the duplexes. | 13 |
| **Figure 1.C.S.1. Bimolecular complex with parallel G4-dimer and head-to-tail IM-dimer in case of the strands exchange: A** – conformation obtained at the last step of the MD trajectory (side and top view); **B** – the complex scheme. | 14 |
| **Figure 1.C.S.2. Bimolecular complex with parallel G4-dimer and head-to-tail IM-dimer in case of the strands exchange: C** – angles between unmelted fragments of the duplexes; **D** - evolution of the values of distances from COMs of the guanines' bases to COMs of their containing tetrads, **E** - angles between normals to the guanines' bases and vectors connecting COMs of the boundary tetrads; **F** - distances between COMs of the cytosines' bases; **G** - angles between normals to the cytosines' bases; **H** – angles of rotation of the tetrads relative to each other. | 15 |
| **Figure 1.D.S.1. Bimolecular complex with antiparallel G4-dimer and head-to-head IM-dimer: A** – conformation obtained at the last step of the MD trajectory (side and top view); **B** – the complex scheme. | 16 |
| **Figure 1.D.S.2. Bimolecular complex with antiparallel G4-dimer and head-to-head IM-dimer: C** – angles between unmelted fragments of the duplexes; **D** - distances from COMs of the guanines' bases to COMs of their containing tetrads; **E** - angles between normals to the guanines' bases and vectors connecting COMs of the boundary tetrads; **F** - distances between COMs of the cytosines' bases; **G** - angles between normals to the cytosines' bases; **H** – angles of rotation of the tetrads relative to each other. | 17 |
| **Figure 1.S.E. The contributions to free energy for variants shown on Figure 2 and Figure 3 during MD calculations.** $E_{eq}$ – electrostatic, $E_{vdw}$ - Van der Waals, $E_{GB}$ - polar energy of solvation, $E_{surf}$ - non-polar energy of solvation due to the hydrophobic surface available to the solvent, U = $E_{bond}$ + $E_{angle}$+ $E_{tor}$ , e.g. $E_{bond}$, $E_{angle}$ and $E_{tor}$ –bond, angle and torsion stress energies. The energy plots were smoothed using moving average method (span = 5). Average energy values are indicated in the figure legends. | 18 |
| **Figure 2.A.S.1. Tetrameric complex with parallel G4-dimer and two head-to-tail IM-dimers: A** – conformation obtained at the last step of the MD trajectory (side and top view); **B** – the complex scheme; **C** – angles of rotation of the tetrads relative to each other. | 19 |

**Figure 2.A.S.2. Tetrameric complex with parallel G4-dimer and two head-to-tail IM-dimers: Tetrameric complex with parallel G4-dimer and two head-to-tail IM-dimers: D**– angles between unmelted fragments of the duplexes; **E** - distances from COMs of the guanines' bases to COMs of their containing tetrads; **F** - angles between normals to the guanines' bases and vectors connecting COMs of the boundary tetrads; **G** - distances between COMs of the cytosines' bases; **H** - angles between normals to the cytosines' bases. 20

**Figure 2.B.S.1. Tetrameric complex with three parallel G4-dimers: A** – conformation obtained at the last step of the MD trajectory (side and top view); **B** – the complex scheme. 21

**Figure 2.B.S.2. Tetrameric complex with three parallel G4-dimers: C** – angles between unmelted fragments of the duplexes; **D, F** - distances from COMs of the guanines' bases to COMs of their containing tetrads; **E, G** - angles between normals to the guanines' bases and vectors connecting COMs of the boundary tetrads. 22

**Figure 2.B.S.3. Tetrameric complex with three parallel G4-dimers: H** - distances from COMs of the guanines' bases to COMs of their containing tetrads; **I** - angles between normals to the guanines' bases and vectors connecting COMs of the boundary tetrads; **J** - angles of rotation of the tetrads relative to each other. 23

**Figure 3.1.A.S.1. Case 1 "1,3 hitch and two IM-monomers": A** – conformation obtained at the last step of the MD trajectory (side and top view); **B** – the complex scheme. 24

**Figure 3.1.A.S.2. Case 1 "1,3 hitch and two IM-monomers": C** – angles between unmelted fragments of the duplexes, angle between axes passing through COMs of the boundary tetrads (**QIQII**); **D**– angles of rotation of the tetrads relative to each other; **E, G** - distances from COMs of the guanines' bases to COMs of their containing tetrads, distance between COMs of the boundary tetrads (**Iq3 IIq1**); **F, H** - angles between normals to the guanines' bases and vectors connecting COMs of the boundary tetrads. 25

**Figure 3.1.A.S.3. Case 1 "1,3 hitch and two IM-monomers": I -** distances between COMs of the cytosines' bases; **J** - angles between normals to the cytosines' bases. 26

**Figure 3.1.B.S.1. Case 2: "1,3 hitch and head-to-tail IM-dimer"**: **A** – conformation obtained at the last step of the MD trajectory (side and top view); **B** – the complex scheme. 27

**Figure 3.1.B.S.2. Case 2: "1,3 hitch and head-to-tail IM-dimer"**: **C** – angles between unmelted fragments of the duplexes, angle between axes passing through COMs of the boundary tetrads (**QIQII**); **D**– angles of rotation of the tetrads relative to each other; **E, G** - distances from COMs of the guanines' bases to COMs of their containing tetrads, distance between COMs of the boundary tetrads (**Iq3 IIq1**); **F, H** - angles between normals to the guanines' bases and vectors connecting COMs of the boundary tetrads. 28

**Figure 3.1.B.S.3. Case 2: "1,3 hitch and head-to-tail IM-dimer": I -** distances between COMs of the cytosines' bases; **J** - angles between normals to the cytosines' bases. 29

**Figure 3.1.C.S.1. Case 3: "1,2 hitch and two head-to-head IM-monomers with mutual girth of the strands"**: **A** – conformation obtained at the last step of the MD trajectory (side and top view); **B** – the complex scheme. 30

**Figure 3.1.C.S.2. Case 3: "1,2 hitch and two head-to-head IM-monomers with mutual girth of the strands"**: **C** – angles between unmelted fragments of the duplexes, angle between axes passing through COMs of the boundary tetrads (**QIQII**); **D**– angles of rotation of the tetrads relative to each other; **E, G** - distances from COMs of the guanines' bases to COMs of their containing tetrads, distance between COMs of the boundary tetrads (**Iq3 IIq1**); **F, H** - angles between normals to the guanines' bases and vectors connecting COMs of the boundary tetrads. 31

**Figure 3.1.C.S.3. Case 3: "1,2 hitch and two head-to-head IM-monomers with mutual girth of the strands"**: **I -** distances between COMs of the cytosines' bases; **J** - angles between normals to the cytosines' bases. 32

| | |
|---|---|
| **Figure 3.2.A.S.1. Case 4: “Stacking with 2 IM-monomers”**: **A** – conformation obtained at the last step of the MD trajectory (side and top view); **B** – the complex scheme. | 33 |
| **Figure 3.2.A.S.2. Case 4: “Stacking with 2 IM-monomers”**: **C** – angles between unmelted fragments of the duplexes, angle between axes passing through COMs of the boundary tetrads (**QIQII**); **D**– angles of rotation of the tetrads relative to each other; **E, G** - distances from COMs of the guanines’ bases to COMs of their containing tetrads, distance between COMs of the boundary tetrads (**Iq3 IIq1**); **F, H** - angles between normals to the guanines’ bases and vectors connecting COMs of the boundary tetrads. | 34 |
| **Figure 3.2.A.S.3. Case 4: “Stacking with 2 IM-monomers”**: **I -** distances between COMs of the cytosines’ bases; **J** - angles between normals to the cytosines’ bases. | 35 |
| **Figure 3.2.B.S.1. Case 5: “Stacking and head-to-tail IM-dimer”**: **A** – conformation obtained at the last step of the MD trajectory (side and top view); **B** – the complex scheme. | 36 |
| **Figure 3.2.B.S.2. Case 5: “Stacking and head-to-tail IM-dimer”**: **C** – angles between unmelted fragments of the duplexes, angle between axes passing through COMs of the boundary tetrads (**QIQII**); **D**– angles of rotation of the tetrads relative to each other; **E, G** - distances from COMs of the guanines’ bases to COMs of their containing tetrads, distance between COMs of the boundary tetrads (**Iq3 IIq1**); **F, H** - angles between normals to the guanines’ bases and vectors connecting COMs of the boundary tetrads. | 37 |
| **Figure 3.2.B.S.3. Case 5: “Stacking and head-to-tail IM-dimer”**: **I -** distances between COMs of the cytosines’ bases; **J** - angles between normals to the cytosines’ bases. | 38 |
| **Figure 3.2.C.S.1. Case 6: “Flip-flop interlock and two IM-monomers”**: **A** – conformation obtained at the last step of the MD trajectory (side and top view); **B** – the complex scheme. | 39 |
| **Figure 3.2.C.S.2. Case 6: “Flip-flop interlock and two IM-monomers”**: **D -** angles between unmelted fragments of the duplexes, angle between axes passing through COMs of the boundary tetrads (**QIQII**); **E**– angles of rotation of the tetrads relative to each other; **F, H** - distances from COMs of the guanines’ bases to COMs of their containing tetrads, distance between COMs of the boundary tetrads (**Iq1 IIq1**); **G, I** - angles between normals to the guanines’ bases and vectors connecting COMs of the boundary tetrads. | 40 |
| **Figure 3.2.C.S.3. Case 6: “Flip-flop interlock and two IM-monomers”**: **I -** distances between COMs of the cytosines’ bases; **J** - angles between normals to the cytosines’ bases. | 41 |
| **Figure 3.3.A.S.1. Case 7: “Stacking of right and left handed parallel G4-dimers, and two IM-monomers”**: **A** – conformation obtained at the last step of the MD trajectory (side and top view); **B** – the complex scheme. | 42 |
| **Figure 3.3.A.S.2. Case 7: “Stacking of right and left handed parallel G4-dimers, and two IM-monomers”**: **C** – angles between unmelted fragments of the duplexes, angle between axes passing through COMs of the boundary tetrads (**QIQII**); **D**– angles of rotation of the tetrads relative to each other; **E, G** - distances from COMs of the guanines’ bases to COMs of their containing tetrads, distance between COMs of the boundary tetrads (**Iq3 IIq1**); **F, H** - angles between normals to the guanines’ bases and vectors connecting COMs of the boundary tetrads. | 43 |
| **Figure 3.3.A.S.3. Case 7: “Stacking of right and left handed parallel G4-dimers, and two IM-monomers”**: **I -** distances between COMs of the cytosines’ bases; **J** - angles between normals to the cytosines’ bases. | 44 |
| **Figure 3.3.B.S.1. Case 8: “Antiparallel G4-dimer and head-to-head IM-dimer”**: **A** – conformation obtained at the last step of the MD trajectory (side and top view); **B** – the complex scheme. | 45 |
| **Figure 3.3.B.S.2. Case 8: “Antiparallel G4-dimer and head-to-head IM-dimer: C** – angles between unmelted fragments of the duplexes, angle between axes passing through COMs of the boundary tetrads (**QIQII**); **D**– angles of rotation of the tetrads relative to each other; **E, G** - distances from COMs of the guanines’ bases to COMs of their containing tetrads, distance between COMs of | 46 |

the boundary tetrads (**Iq3 IIq1**); **F, H** - angles between normals to the guanines' bases and vectors connecting COMs of the boundary tetrads.

**Figure 3.3.B.S.3. Case 8: "Antiparallel G4-dimer and head-to-head IM-dimer"**: **I -** distances between COMs of the cytosines' bases; **J** - angles between normals to the cytosines' bases. 47

**Figure 3.3.C.S.1. Case 9: "Head-to-tail IM-dimer between two parallel G4-monomers"**: **A** – conformation obtained at the last step of the MD trajectory (side and top view); **B** – the complex scheme. 48

**Figure 3.3.C.S.2. Case 9: "Head-to-tail IM-dimer between two parallel G4-monomers"**: **C** – angles between unmelted fragments of the duplexes, angle between axes passing through COMs of the boundary tetrads (**QIQII**); **D** – angles of rotation of the tetrads relative to each other; **E, G** - distances from COMs of the guanines' bases to COMs of their containing tetrads; **F, H** - angles between normals to the guanines' bases and vectors connecting COMs of the boundary tetrads. 49

**Figure 3.3.C.S.3. Case 9: "Head-to-tail IM-dimer between two parallel G4-monomers"**: **I** - distances between COMs of the cytosines' bases; **J** - angles between normals to the cytosines' bases. 50

**Figure 3.4.A.S.1. Case 10: "Head-to-head IM-dimer between two parallel G4-monomers"**: **A** – conformation obtained at the last step of the MD trajectory (side and top view); **B** – the complex scheme. 51

**Figure 3.4.A.S.2. Case 10: "Head-to-head IM-dimer between two parallel G4-monomers"**: **C** – angles between unmelted fragments of the duplexes, angle between axes passing through COMs of the boundary tetrads (**QIQII**); **D** – angles of rotation of the tetrads relative to each other; **E** – evolution of values of distances between COMs of the G4s and distances between COMs of bases of **T51** and **T113**; **F, H** - distances from COMs of the guanines' bases to COMs of their containing tetrad;, **G,I** - angles between normals to the guanines' bases and vectors connecting COMs of the boundary tetrads. 52

**Figure 3.4.A.S.3. Case 10: "Head-to-head IM-dimer between two parallel G4-monomers"**: **J** - distances between COMs of the cytosines' bases; **K** - angles between normals to the cytosines' bases. 53

**Figure 3.4.B.S.1. Case 11: "Head-to-head IM-dimer between two parallel G4-monomers in case of the strands exchange"**: **A** – conformation obtained at the last step of the MD trajectory (side and top view); **B** – the complex scheme. 54

**Figure 3.4.B.S.2. Case 11: "Head-to-head IM-dimer between two parallel G4-monomers in case of the strands exchange"**: **C** – angles between unmelted fragments of the duplexes, angle between axes passing through COMs of the boundary tetrads (**QIQII**); **D** – angles of rotation of the tetrads relative to each other; **E** – evolution of values of distances between COMs of the G4s and distances between COMs of bases of **T51** and **T113**; **F, H** - distances from COMs of the guanines' bases to COMs of their containing tetrad; **G, I** - angles between normals to the guanines' bases and vectors connecting COMs of the boundary tetrads. 55

**Figure 3.4.B.S.3. Case 11: "Head-to-head IM-dimer between two parallel G4-monomers in case of the strands exchange"**: **J** - distances between COMs of the cytosines' bases; **K** - angles between normals to the cytosines' bases. 56

**Figure 3.4.C.S.1. Case 12: "Two parallel G4-dimer in the same plane and head-to-head IM-dimer between two unmelted fragments of duplexes with mutual girth of the strands"**: **A** – conformation obtained at the last step of the MD trajectory (side and top view); **B** – the complex scheme. 57

**Figure 3.4.C.S.2. Case 12: "Two parallel G4-dimer in the same plane and head-to-head IM-dimer between two unmelted fragments of duplexes with mutual girth of the strands"**: **C** – angles between unmelted fragments of the duplexes, angle between axes passing through COMs of the boundary tetrads (**QIQII**); **D** – angles of rotation of the tetrads relative to each other; **E, G** - between normals to the guanines' bases and vectors connecting COMs of the boundary tetrads, distances from COMs of the guanines' bases to COMs of their containing tetrad; **F, H** - angles between normals to the guanines' bases and vectors connecting COMs of the boundary tetrads. 58

| | |
|---|---|
| **Figure 3.4.C.S.3. Case 12: “Two parallel G4-dimer in the same plane and head-to-head IM-dimer between two unmelted fragments of duplexes with mutual girth of the strands”**: **I** - distances between COMs of the cytosines’ bases; **J** - angles between normals to the cytosines’ bases. | 59 |
| **Figure 3.5.A.S.1. Case 13: “Two parallel G4-dimers in the same plane and two IM-monomers clamped unmelted fragments of duplexes with mutual girth of the strands”: A** – conformation obtained at the last step of the MD trajectory (side and top view); **B** – the complex scheme. | 60 |
| **Figure 3.5.A.S.2. Case 13: “Two parallel G4-dimers in the same plane and two IM-monomers clamped unmelted fragments of duplexes with mutual girth of the strands”: C** – angles between unmelted fragments of the duplexes, angle between axes passing through COMs of the boundary tetrads (**QIQII**); **D**– angles of rotation of the tetrads relative to each other; **E, G** - distances from COMs of the guanines’ bases to COMs of their containing tetrads; **F, H** - angles between normals to the guanines’ bases and vectors connecting COMs of the boundary tetrads. | 61 |
| **Figure 3.5.A.S.3. Case 13: “Two parallel G4-dimers in the same plane and two IM-monomers clamped unmelted fragments of duplexes with mutual girth of the strands”: I** - distances between COMs of the cytosines’ bases; **J** - angles between normals to the cytosines’ bases. | 62 |
| **Figure 3.5.B.S.1. Case 14: “Two parallel G4-dimers in the same plane and head-to-tail IM-dimer between two unmelted fragments of duplexes with exchange and mutual girth of the strands”: A** – conformation obtained at the last step of the MD trajectory (side and top view); **B** – the complex scheme. | 63 |
| **Figure 3.5.B.S.2. Case 14: “Two parallel G4-dimers in the same plane and head-to-tail IM-dimer between two unmelted fragments of duplexes with exchange and mutual girth of the strands”: C** - angles between unmelted fragments of the duplexes, angle between axes passing through COMs of the boundary tetrads (**QIQII**); **D** - angles of rotation of the tetrads relative to each other; **E, G** - distances from COMs of the guanines’ bases to COMs of their containing tetrads; **F, H** - angles between normals to the guanines’ bases and vectors connecting COMs of the boundary tetrads. | 64 |
| **Figure 3.5.B.S.3. Case 14: “Two parallel G4-dimers in the same plane and head-to-tail IM-dimer between two unmelted fragments of duplexes with exchange and mutual girth of the strands”: I** - distances between COMs of the cytosines’ bases; **J** - angles between normals to the cytosines’ bases. | 65 |
| **Figure 3.5.C.S.1. Case 15: “Two parallel G4-dimers in the same plane and head-to-tail IM-dimer between two unmelted fragments of duplexes with the strands exchange”: A** – conformation obtained at the last step of the MD trajectory (side and top view); **B** – the complex scheme. | 66 |
| **Figure 3.5.C.S.2. Case 15: “Two parallel G4-dimers in the same plane and head-to-tail IM-dimer between two unmelted fragments of duplexes with the strands exchange” : C** – angles between unmelted fragments of the duplexes, angle between axes passing through COMs of the boundary tetrads (**QIQII**); **D**– angles of rotation of the tetrads relative to each other; **E, G -** distances from COMs of the guanines’ bases to COMs of their containing tetrads; **F, H** - angles between normals to the guanines’ bases and vectors connecting COMs of the boundary tetrads. | 67 |
| **Figure 3.5.C.S.3. Case 15: “Two parallel G4-dimers in the same plane and head-to-tail IM-dimer between two unmelted fragments of duplexes with the strands exchange”: I** - distances between COMs of the cytosines’ bases; **J** - angles between normals to the cytosines’ bases. | 68 |
| **Figure 3.S.E.1. The contributions to free energy during MD calculations for the variants of bimolecular complexes of unmelted fragments of duplexes containing $(G_3T)_3G_3$ and $(C_3A)_3C_3$ sequences with G4/IM in cases from 1 to 5**. $E_{eq}$ – electrostatic, $E_{vdw}$ - Van der Waals, $E_{GB}$ - polar energy of solvation, $E_{surf}$ - non-polar energy of solvation due to the hydrophobic surface available to the solvent, $U = E_{bond} + E_{angle} + E_{tor}$ , e.g. $E_{bond}$, $E_{angle}$ and $E_{tor}$ –bond, angle and torsion stress energies. The energy plots were smoothed using moving average method (span = 5). Average energy values are indicated in the figure legends. | 69 |

**Figure 3.S.E.2. The contributions to free energy during MD calculations for the variants of bimolecular complexes of unmelted fragments of duplexes containing $(G_3T)_3G_3$ and $(C_3A)_3C_3$ sequences with G4/IM in cases from 6 to 10**. $E_{eq}$ – electrostatic, $E_{vdw}$ - Van der Waals, $E_{GB}$ - polar energy of solvation, $E_{surf}$ - non-polar energy of solvation due to the hydrophobic surface available to the solvent, U = $E_{bond}$+ $E_{angle}$+ $E_{tor}$ , e.g. $E_{bond}$, $E_{angle}$ and $E_{tor}$ –bond, angle and torsion stress energies. The energy plots were smoothed using moving average method (span = 5). Average energy values are indicated in the figure legends. 70

**Figure 3.S.E.3. The contributions to free energy during MD calculations for the variants of bimolecular complexes of unmelted fragments of duplexes containing $(G_3T)_3G_3$ and $(C_3A)_3C_3$ sequences with G4/IM in cases from 11 to 15**. $E_{eq}$ – electrostatic, $E_{vdw}$ - Van der Waals, $E_{GB}$ - polar energy of solvation, $E_{surf}$ - non-polar energy of solvation due to the hydrophobic surface available to the solvent, U = $E_{bond}$+ $E_{angle}$+ $E_{tor}$ , e.g. $E_{bond}$, $E_{angle}$ and $E_{tor}$ –bond, angle and torsion stress energies. The energy plots were smoothed using moving average method (span = 5). Average energy values are indicated in the figure legends. 71

**Figure 4.1.S.1. Case 1: “Four parallel G4-dimers in the same plane and four IM-monomers between the unmelted fragments of duplexes with exchange and mutual girth of the strands”: A -** conformation obtained at the last step of the MD trajectory. Top row is the side view, bottom row is the top view. On the left in both rows are views with unmelted fragments of the duplexes, on the right there are only G4s and IMs. 72

**Figure 4.1.S.2. Case 1: “Four parallel G4-dimers in the same plane and four IM-monomers between the unmelted fragments of duplexes with exchange and mutual girth of the strands”: B** – the complex scheme; **C** – angles between straight lines, passing through the COMs of the first and the last complementary pairs of unmelted fragments of duplexes, and planes, containing COMs of the G4s tetrads; **D**– angles of rotation of the tetrads relative to each other; **E** - angles between the straight line, passing through the COM of all upper tetrads and the COM of all lower tetrads, and the straight lines, passing through the COMs of the upper and lower tetrads, in the G4s’ case and straight lines, passing through the COMs of the boundary cytosine pairs, in the IMs’ case. 73

**Figure 4.1.S.3. Case 1: “Four parallel G4-dimers in the same plane and four IM-monomers between the unmelted fragments of duplexes with exchange and mutual girth of the strands”:** distances from COMs of the guanines’ bases to COMs of their containing tetrad. 74

**Figure 4.1.S.4. Case 1: “Four parallel G4-dimers in the same plane and four IM-monomers between the unmelted fragments of duplexes with exchange and mutual girth of the strands”:** angles between normals to the guanine’ bases and vectors connecting COMs of the boundary tetrads. 75

**Figure 4.1.S.5. Case 1: “Four parallel G4-dimers in the same plane and four IM-monomers between the unmelted fragments of duplexes with exchange and mutual girth of the strands”: F, H** - distances between COMs of the cytosines’ bases; **G, I** - angles between normals to the cytosines’ bases. 76

**Figure 4.2.S.1. Case 2: “Four parallel G4-dimers in two stacks and four IM-monomers with exchange and mutual girth of the strands”: A-** conformation obtained at the last step of the MD trajectory. The first row is the side view with unmelted fragments of the duplexes, the second row is the side view with only G4s and IMs, the third row is top view with unmelted fragments of the duplexes, and the fourth row is top view with only G4s and IMs. 77

**Figure 4.2.S.2. Case 2: “Four parallel G4-dimers in two stack and four IM-monomers with exchange and mutual girth of the strands”: B** – the complex scheme; **C** – angles between straight lines, passing through COMs of the first and the last complementary pairs of unmelted fragments of the duplexes, and straight line, passing through COMs of boundary tetrads of the G4s; **D**– angles of rotation of the tetrads relative to each other; **E** - angles between the straight line, passing through the COM of all upper tetrads and the COM of all lower tetrads, and the straight lines, passing through the COMs of the upper and lower tetrads, in the G4s’ case and straight lines, passing through the COMs of the boundary cytosine pairs, in the IMs’ case. 78

**Figure 4.2.S.3. Case 2: “Four parallel G4-dimers in two stack and four IM-monomers with exchange and mutual girth of the strands”:** distances from COMs of the guanines’ bases to COMs of their containing tetrad. 79

| | |
|---|---|
| **Figure 4.2.S.4. Case 2: “Four parallel G4-dimers in two stack and four IM-monomers with exchange and mutual girth of the strands”:** angles between normals to the guanine’ bases and vectors connecting COMs of the boundary tetrads. | 80 |
| **Figure 4.2.S.5. Case 2: “Four parallel G4-dimers in two stack and four IM-monomers with exchange and mutual girth of the strands”: F, H** - distances between COMs of the cytosines’ bases; **G, I** - angles between normals to the cytosines’ bases. | 81 |
| **Figure 4.3.S.1. Case 3: “Stacking of four parallel G4-dimers and four IM-monomers”: A**-conformation obtained at the last step of the MD trajectory. Top row is the side view, bottom row is the top view. On the left in both rows are views with unmelted fragments of the duplexes, on the right there are only G4s and IMs. | 82 |
| **Figure 4.3.S.2. Case 3: “Stacking of four parallel G4-dimers and four IM-monomers”: B** – the complex scheme; **C** – angles between straight lines, passing through COMs of the first and the last complementary pairs of unmelted fragments of the duplexes, and straight line, passing through COMs of boundary tetrads of the G4s; **D**– angles of rotation of the tetrads relative to each other; **E** - angles between the straight line, passing through the COM of all upper tetrads and the COM of all lower tetrads, and the straight lines, passing through the COMs of the upper and lower tetrads, in the G4s’ case and straight lines, passing through the COMs of the boundary cytosine pairs, in the IMs’ case. | 83 |
| **Figure 4.3.S.3. Case 3: “Stacking of four parallel G4-dimers and four IM-monomers”:** distances from COMs of the guanines’ bases to COMs of their containing tetrad, distance between COMs of the boundary tetrads (**Iq3 IIq1, IIq3 IIIq1, IIIq3 IVq1**). | 84 |
| **Figure 4.3.S.4. Case 3: “Stacking of four parallel G4-dimers and four IM-monomers”:** angles between normals to the guanine’ bases and vectors connecting COMs of the boundary tetrads. | 85 |
| **Figure 4.3.S.5. Case 3: “Stacking of four parallel G4-dimers and four IM-monomers”: F, H** - distances between COMs of the cytosines’ bases; **G, I** - angles between normals to the cytosines’ bases. | 86 |
| **Figure 4.4.S.1. Case 4: “Two parallel stack with right and left handed G4-dimers, and two head-to-head IM-dimers”: A -** conformation obtained at the last step of the MD trajectory. The first row is the side view with unmelted fragments of the duplexes, the second row is the side view with only G4s and IMs, the third row is top view with unmelted fragments of the duplexes, and the fourth row is top view with only G4s and IMs. | 87 |
| **Figure 4.4.S.2. Case 4: “Two parallel stack with right and left handed G4-dimers, and two head-to-head IM-dimers”: B -** the complex scheme; **C -** angles between the straight lines, passing through the COMs of the first and the last complementary pairs unmelted fragments of duplexes, and the straight line passing through the COMs of boundary tetrads of the G4s; **D -** angles of rotation of the tetrads relative to each other; **E** - angles between the straight line, passing through the COM of all upper tetrads and the COM of all lower tetrads, and the straight lines, passing through the COMs of the upper and lower tetrads, in the G4s’ case and straight lines, passing through the COMs of the boundary cytosine pairs, in the IMs’ case. | 88 |
| **Figure 4.4.S.3. Case 4: “Two parallel stack with right and left handed G4-dimers, and two head-to-head IM-dimers”:** distances from COMs of the guanines’ bases to COMs of their containing tetrad, distance between COMs of the boundary tetrads (**Iq3 IIq1, IIIq3 IVq1**). | 89 |
| **Figure 4.4.S.4. Case 4: “Two parallel stack with right and left handed G4-dimers, and two head-to-head IM-dimers”:** angles between normals to the guanine’ bases and vectors connecting COMs of the boundary tetrads. | 90 |
| **Figure 4.4.S.5. Case 4: “Two parallel stack with right and left handed G4-dimers, and two head-to-head IM-dimers”: F, H** - distances between COMs of the cytosines’ bases; **G, I** - angles between normals to the cytosines’ bases. | 91 |

| | |
|---|---|
| **Figure 4.S.E. The contributions to free energy during MD calculations for the variants of tetrameric complex of unmelted fragments of duplexes containing $(G_3T)_3G_3$ and $(C_3A)_3C_3$ fragments.** $E_{eq}$ – electrostatic, $E_{vdw}$ - Van der Waals, $E_{GB}$ - polar energy of solvation, $E_{surf}$ - non-polar energy of solvation due to the hydrophobic surface available to the solvent, $U = E_{bond} + E_{angle} + E_{tor}$ , e.g. $E_{bond}$, $E_{angle}$ and $E_{tor}$ –bond, angle and torsion stress energies. The energy plots were smoothed using moving average method (span = 5). Average energy values are indicated in the figure legends. | 92 |
| **Figure 5.S.1. Octameric complex with four parallel stack with right and left handed G4-dimers and four head-to-head IM-dimers: A** – conformation obtained at the last step of the MD trajectory (side view, top view, view with only G4s and IMs). | 93 |
| **Figure 5.S.2. Octameric complex with four parallel stack with right and left handed G4-dimers and four head-to-head IM-dimers: B** – the complex scheme; **C** – angles between the straight lines, passing through the COMs of the first and the last complementary pairs unmelted fragments of duplexes, and the straight line passing through the COMs of boundary tetrads of the G4s; **D**– angles of rotation of the tetrads relative to each other. | 94 |
| **Figure 5.S.3. Octameric complex with four parallel stack with right and left handed G4-dimers and four head-to-head IM-dimers: :** distances from COMs of the guanines' bases to COMs of their containing tetrad, distance between COMs of the boundary tetrads (**Iq3 IIq1, IIIq3 IVq1**). | 95 |
| **Figure 5.S.4. Octameric complex with four parallel stack with right and left handed G4-dimers and four head-to-head IM-dimers: :** angles between normals to the guanines' bases and vectors connecting COMs of the boundary tetrads. | 96 |
| **Figure 5.S.5. Octameric complex with four parallel stack with right and left handed G4-dimers and four head-to-head IM-dimers:** distances from COMs of the guanines' bases to COMs of their containing tetrad, distance between COMs of the boundary tetrads (**Vq3 VIq1, VIIq3 VIIIq1**). | 97 |
| **Figure 5.S.6. Octameric complex with four parallel stack with right and left handed G4-dimers and four head-to-head IM-dimers: :** angles between normals to the guanines' bases and vectors connecting COMs of the boundary tetrads. | 98 |
| **Figure 5.S.7. Octameric complex with four parallel stack with right and left handed G4-dimers and four head-to-head IM-dimers: : E, G** - distances between COMs of the cytosines' bases; **F, H** - angles between normals to the cytosines' bases. | 99 |
| **Figure 5.S.8. Octameric complex with four parallel stack with right and left handed G4-dimers and four head-to-head IM-dimers: I** - distances between COMs of the cytosines' bases; **J** - angles between normals to the cytosines' bases; **K** - evolution of angle values between straight line, passing through COM of all upper tetrads and COM of all lower tetrads, and straight lines, passing through the COMs of the upper and lower quarters, in the case of G4s, and straight lines, passing through the COMs of the middle pairs of cytosines, in the case of the IMs. | 100 |
| **Figure 6.S.1. Stacking and two IM-monomers**: **A, B** – conformation obtained at the last step of the MD trajectory (**A** -side and top view, **B** - enlarged view of quadruplexes). | 101 |
| **Figure 6.S.2. Stacking and two IM-monomers**: **B** – the complex scheme; **C** – angles between unmelted fragments of the duplexes and axes passing through COMs of the tetrads, angle between axes passing through COMs of the boundary tetrads (**QIQII**); **D** – angles of rotation of the tetrads relative to each other. | 102 |
| **Figure 6.S.3. Stacking and two IM-monomers**: **E, G** - distances from COMs of the guanines' bases to COMs of their containing tetrads, distance between COMs of the boundary tetrads (**Iq3 IIq1**); **F, H** - angles between normals to the guanines' bases and vectors connecting COMs of the boundary tetrads. | 103 |
| **Figure 6.S.4. Stacking and two IM-monomers**: **J -** distances between COMs of the cytosines' bases; **K** - angles between normals to the cytosines' bases. | 104 |

| | |
|---|---|
| **Figure 7.1.A.S.1. Case 1: “Stacking and two IM-monomers”**: **A** – conformation obtained at the last step of the MD trajectory (side and top view). | 105 |
| **Figure 7.1.A.S.2. Case 1: “Stacking and two IM-monomers”**: **B** – the complex scheme; **C** – angles between unmelted fragments of the duplexes and axes passing through COMs of the tetrads, angle between axes passing through COMs of the boundary tetrads (**QIQII**); **D**– angles of rotation of the tetrads relative to each other. | 106 |
| **Figure 7.1.A.S.3. Case 1: “Stacking and two IM-monomers: E, G -** distances from COMs of the guanines’ bases to COMs of their containing tetrads, distance between COMs of the boundary tetrads **(Iq3 IIq1); F, H** - angles between normals to the guanines’ bases and vectors connecting COMs of the boundary tetrads. | 107 |
| **Figure 7.1.A.S.4. Case 1: “Stacking and two IM-monomers”**: **I** - distances between COMs of the cytosines’ bases; **J** - angles between normals to the cytosines’ bases. | 108 |
| **Figure 7.1.B.S.1. Case 2: “1,2 hitch and two IM-monomers with two mini-duplex”**: **A** – conformations obtained at the last step of the MD trajectory (side and top view). | 109 |
| **Figure 7.1.B.S.2. Case 2: “1,2 hitch and two IM-monomers with two mini-duplex”**: **B** – the complex scheme; **C** – angles between unmelted fragments of the duplexes and axes passing through COMs of the tetrads, angle between axes passing through COMs of the boundary tetrads (**QIQII**); **D** – angles of rotation of the tetrads relative to each other. | 110 |
| **Figure 7.1.B.S.3. Case 2: “1,2 hitch and two IM-monomers with two mini-duplex”**: **E, G -** distances from COMs of the guanines’ bases to COMs of their containing tetrads, distance between COMs of the boundary tetrads **(Iq3 IIq1); F, H** - angles between normals to the guanines’ bases and vectors connecting COMs of the boundary tetrads. | 111 |
| **Figure 7.1.B.S.4. Case 2: “1,2 hitch and two IM-monomers with two mini-duplexes”**: **I** - distances between COMs of the cytosines’ bases; **J** - angles between normals to the cytosines’ bases ; **K** - number of hydrogen bonds in mini-duplexes. | 112 |
| **Table .1(Appendix to Figure 7.1.B.S.4. Case 2)** Percentage of snapshots with hydrogen bonds generated by pairs in the mini-duplexes. | 113 |
| **Figure 7.2.A.S.1. Case 3: “Stacking of right and left handed parallel G4-dimers, and two IM-monomers with 4 mini-unmelted fragments of duplexes”**: **A** – conformation obtained at the last step of the MD trajectory (side and top view). | 114 |
| **Figure 7.2.A.S.2. Case 3: “Stacking of right and left handed parallel G4-dimers, and two IM-monomers with 4 mini-unmelted fragments of duplexes”**: **B** – the complex scheme; **C** – angles between unmelted fragments of the duplexes and axes passing through COMs of the tetrads, angle between axes passing through COMs of the boundary tetrads (**QIQII**); **D**– angles of rotation of the tetrads relative to each other. | 115 |
| **Figure 7.2.A.S.3. Case 3: “Stacking of right and left handed parallel G4-dimers, and two IM-monomers with 4 mini-unmelted fragments of duplexes”**: **E, G -** distances from COMs of the guanines’ bases to COMs of their containing tetrads, distance between COMs of the boundary tetrads (**Iq3 IIq1**); **F, H** - angles between normals to the guanines’ bases and vectors connecting COMs of the boundary tetrads. | 116 |
| **Figure 7.2.A.S.4. Case 3: “Stacking of right and left handed parallel G4-dimers, and two IM-monomers with 4 mini-unmelted fragments of duplexes”**: **I** - distances between COMs of the cytosines’ bases; **J** - angles between normals to the cytosines’ bases; **K**, **L** - number of hydrogen bonds in mini-duplexes. | 117 |
| **Table .2. (Appendix to Figure 7.2.A.S.4. Case 3) Percentage of snapshots with hydrogen bonds generated by pairs in the mini-duplexes.** | 118 |

| | |
|---|---|
| **Figure 7.2.B.S.1. Case 4: “Three parallel G4-dimers in the same plane and head-to-tail IM-dimer with the strands exchange”**: **A** – conformation obtained at the last step of the MD trajectory (side and top view). | 119 |
| **Figure 7.2.B.S.2. Case 4: “Three parallel G4-dimers in the same plane and head-to-tail IM-dimer with the strands exchange”**: **B** – the complex scheme; **C** – angles between unmelted fragments of the duplexes and axes passing through COMs of the tetrads, angle between axes passing through COMs of the boundary tetrads (**QIQII, QIIQIII**); **D** – angles of rotation of the tetrads relative to each other. | 120 |
| **Figure 7.2.B.S.3. Case 4: “Three parallel G4-dimers in the same plane and head-to-tail IM-dimer with the strands exchange”**: **E, G** - distances from COMs of the guanines’ bases to COMs of their containing tetrad; **F, H** - angles between normals to the guanines’ bases and vectors connecting COMs of the boundary tetrads. | 121 |
| **Figure 7.2.B.S.4. Case 4: “Three parallel G4-dimers in the same plane and head-to-tail IM-dimer with the strands exchange”**: **I** - distances from COMs of the guanines’ bases to COMs of their containing tetrad, **J** - angles between normals to the guanines’ bases and vectors connecting COMs of the boundary tetrads; **K** - distances between COMs of the cytosines’ bases; **L** - angles between normals to the cytosines’ bases. | 122 |
| **Figure 7.3.A.S.1. Case 5: “Stacking of three parallel G4-dimers”**: **A** – conformation obtained at the last step of the MD trajectory (side and top view). | 123 |
| **Figure 7.3.A.S.2. Case 5: “Stacking of three parallel G4-dimers”**: **B** – the complex scheme; **C** – angles between unmelted fragments of the duplexes and axes passing through COMs of the tetrads, angle between axes passing through COMs of the boundary tetrads (**QIQII, QIIQIII**); **D** – angles of rotation of the tetrads relative to each other. | 124 |
| **Figure 7.3.A.S.3. Case 5: “Stacking of three parallel G4-dimers”**: **E, G** - distances from COMs of the guanines’ bases to COMs of their containing tetrad, distance between COMs of the boundary tetrads (**Iq3 IIq1**); **F, H** - angles between normals to the guanines’ bases and vectors connecting COMs of the boundary tetrads. | 125 |
| **Figure 7.3.A.S.4. Case 5: “Stacking of three parallel G4-dimers”**: **I** - distances from COMs of the guanines’ bases to COMs of their containing tetrad, distance between COMs of the boundary tetrads (**IIq3 IIIq1**); **J** - angles between normals to the guanines’ bases and vectors connecting COMs of the boundary tetrads. | 126 |
| **Figure 7.3.B.S.1. Case 6: “Stacking of two parallel G4-monomers and G4-dimer, and two IM-monomers”**: **A** – conformation obtained at the last step of the MD trajectory (side and top view). | 127 |
| **Figure 7.3.B.S.2. Case 6: “Stacking of two parallel G4-monomers and G4-dimer, and two IM-monomers”**: **B** – the complex scheme; **C** – angles between unmelted fragments of the duplexes, angles between axes passing through the COMs of boundary tetrads of the G4s (**QIQII**, **QIIQIII**); **D** – angles of rotation of the tetrads relative to each other. | 128 |
| **Figure 7.3.B.S.3. Case 6: “Stacking of two parallel G4-monomers and G4-dimer, and two IM-monomers”**: **E, G** - distances from COMs of the guanines’ bases to COMs of their containing tetrad, distance between COMs of the boundary tetrads (**Iq3 IIq1**); **F, H** - angles between normals to the guanines’ bases and vectors connecting COMs of the boundary tetrads. | 129 |
| **Figure 7.3.B.S.4. Case 6: “Stacking of two parallel G4-monomers and G4-dimer, and two IM-monomers”**: **I** - distances from COMs of the guanines’ bases to COMs of their containing tetrad, distance between COMs of the boundary tetrads (**IIq3 IIIq1**); **J** - angles between normals to the guanines’ bases and vectors connecting COMs of the boundary tetrads; **K** - distances between COMs of the cytosines’ bases; **L** - angles between normals to the cytosines’ bases. | 130 |
| **Figure 7.4.A.S.1. Case 7: “Stacking of three parallel G4-dimer and two IM-monomers with mutual girth of the strands”: A** – conformation obtained at the last step of the MD trajectory (side and top view). | 131 |

**Figure 7.4.A.S.2. Case 7: “Stacking of three parallel G4-dimer and two IM-monomers with mutual girth of the strands”: B** – the complex scheme; **C** – angles between unmelted fragments of the duplexes, angles between axes passing through the COMs of boundary tetrads of the G4s (**QIQII**, **QIIQIII**); **D**– angles of rotation of the tetrads relative to each other. 132

**Figure 7.4.A.S.3. Case 7: “Stacking of three parallel G4-dimer and two IM-monomers with mutual girth of the strands”: E, G** - distances from COMs of the guanines’ bases to COMs of their containing tetrad, distance between COMs of the boundary tetrads (**Iq3 IIq1**); **F, H** - angles between normals to the guanines’ bases and vectors connecting COMs of the boundary tetrads. 133

**Figure 7.4.A.S.3. Case 7: “Stacking of three parallel G4-dimer and two IM-monomers with mutual girth of the strands”: I** - distances from COMs of the guanines’ bases to COMs of their containing tetrad, distance between COMs of the boundary tetrads (**IIq3 IIIq1**); **J** - angles between normals to the guanines’ bases and vectors connecting COMs of the boundary tetrads; **K**- distances between COMs of the cytosines’ bases; **L** - angles between normals to the cytosines’ bases. 134

**Figure 7.4.B.S.1. Case 8: “Three parallel G4-dimers in the same plane with and two IM-monomers with mutual girth of the strands”**: **A** – conformation obtained at the last step of the MD trajectory (side and top view). 135

**Figure 7.4.B.S.2. Case 8: “Three parallel G4-dimers in the same plane with and two IM-monomers with mutual girth of the strands”**: **B** – the complex scheme; **C** – angles between unmelted fragments of the duplexes and the axes passing through the COMs of the tetrads, the angle values between the G4s (**QIQII**, **QIIQIII**); **D** – angles of rotation of the tetrads relative to each other. 136

**Figure 7.4.B.S.3. Case 8: “Three parallel G4-dimers in the same plane with and two IM-monomers with mutual girth of the strands”**: **E, G** - distances from COMs of the guanines’ bases to COMs of their containing tetrad, distance between COMs of the G4s (**Iq2 Iiq2**); **F, H** - angles between normals to the guanines’ bases and vectors connecting COMs of the boundary tetrads. 137

**Figure 7.4.B.S.4. Case 8: “Three parallel G4-dimers in the same plane with and two IM-monomers with mutual girth of the strands”**: **I** - distances from COMs of the guanines’ bases to COMs of their containing tetrad, distance between COMs of the boundary tetrads (**Iiq2 IIIq2**); **J** - angles between normals to the guanines’ bases and vectors connecting COMs of the boundary tetrads; **K** - distances between COMs of the cytosines’ bases; **L** - angles between normals to the cytosines’ bases. 138

**Figure 7.S.E.1. The contributions to free energy during MD calculations for the variants of bimolecular complex of duplexes containing $(G_3T)_5G_3$ and $(C_3A)_5C_3$ sequences with G4/IM in cases from 1 to 4.** $E_{eq}$ – electrostatic, $E_{vdw}$ - Van der Waals, $E_{GB}$ - polar energy of solvation, $E_{surf}$ - non-polar energy of solvation due to the hydrophobic surface available to the solvent, U = $E_{bond}$+ $E_{angle}$+ $E_{tor}$ , e.g. $E_{bond}$, $E_{angle}$ and $E_{tor}$ –bond, angle and torsion stress energies. The energy plots were smoothed using moving average method (span = 5). Average energy values are indicated in the figure legends. 139

**Figure 7.S.E.2. The contributions to free energy during MD calculations for the variants of bimolecular complex of duplexes containing $(G_3T)_5G_3$ and $(C_3A)_5C_3$ sequences with G4/IM in cases from 5 to 8.** $E_{eq}$ – electrostatic, $E_{vdw}$ - Van der Waals, $E_{GB}$ - polar energy of solvation, $E_{surf}$ - non-polar energy of solvation due to the hydrophobic surface available to the solvent, U = $E_{bond}$+ $E_{angle}$+ $E_{tor}$ , e.g. $E_{bond}$, $E_{angle}$ and $E_{tor}$ –bond, angle and torsion stress energies. The energy plots were smoothed using moving average method (span = 5). Average energy values are indicated in the figure legends. 140

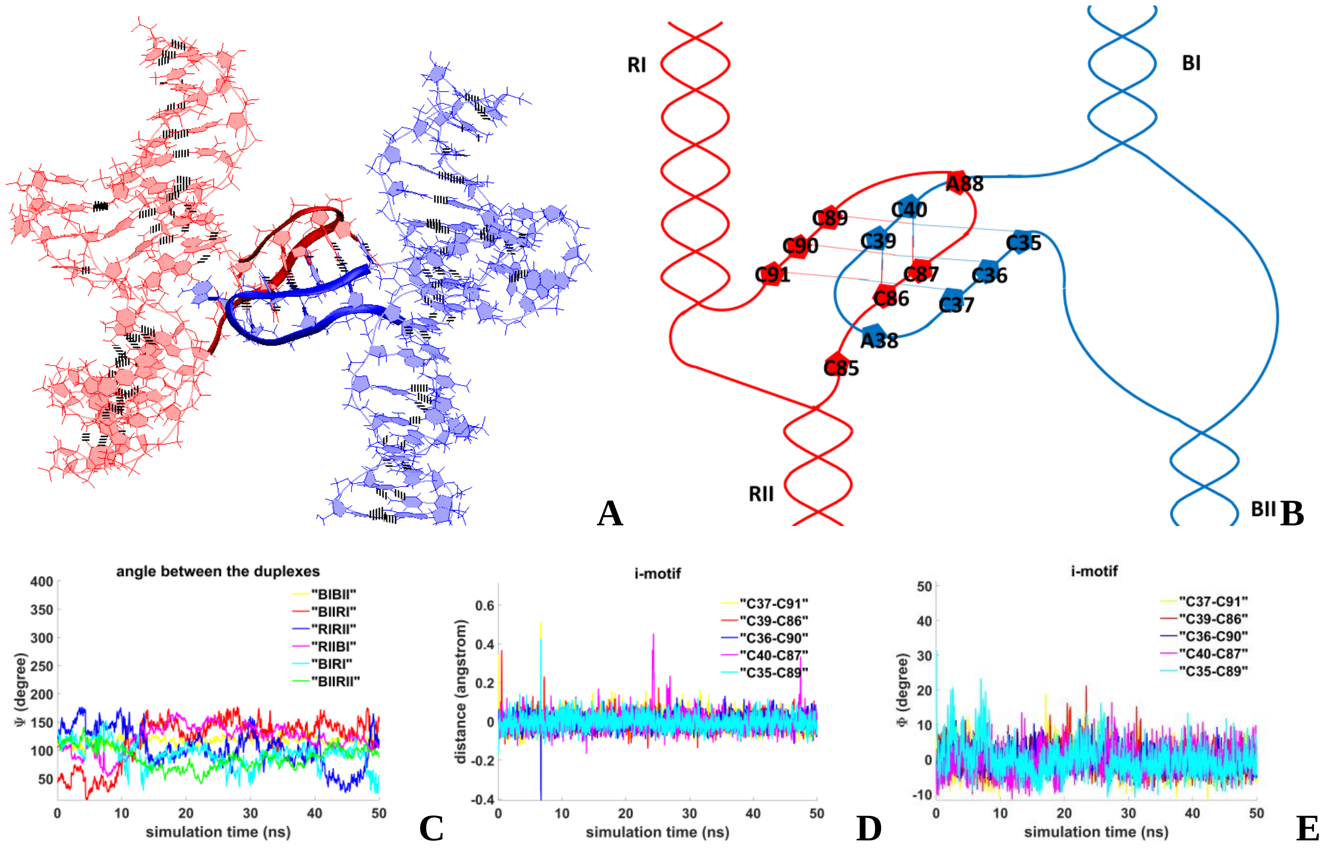


**Figure 1.A.S. Bimolecular complex with head-to-tail IM-dimer: A** – conformation obtained at the last step of the MD trajectory; **B** – the complex scheme; **C** – angles between unmelted fragments of the duplexes; **D -** distances between COMs of the cytosines' bases; **E -** angles between normals to the cytosines' bases.

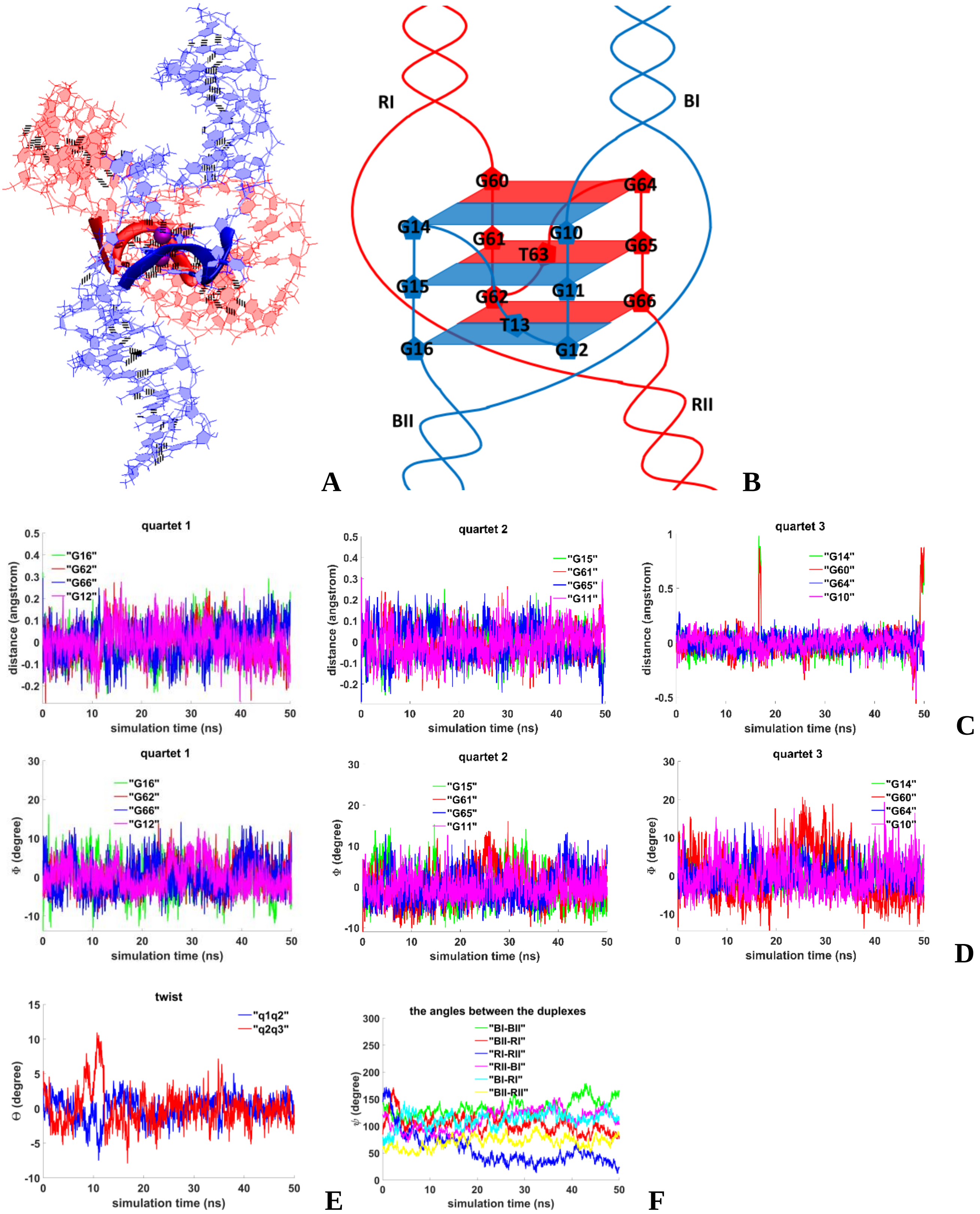


**Figure 1.B.S. Bimolecular complex with parallel G4-dimer**: **A** – conformation obtained at the last step of the MD trajectory; **B** – the complex scheme; **C** - distances from COMs of the guanines' bases to COMs of their containing tetrads; **D** – angles between normals to the guanines' bases and vectors connecting COMs of the boundary tetrads; **E** – angles of rotation of the tetrads relative to each other; **F** – angles between unmelted fragments of the duplexes.

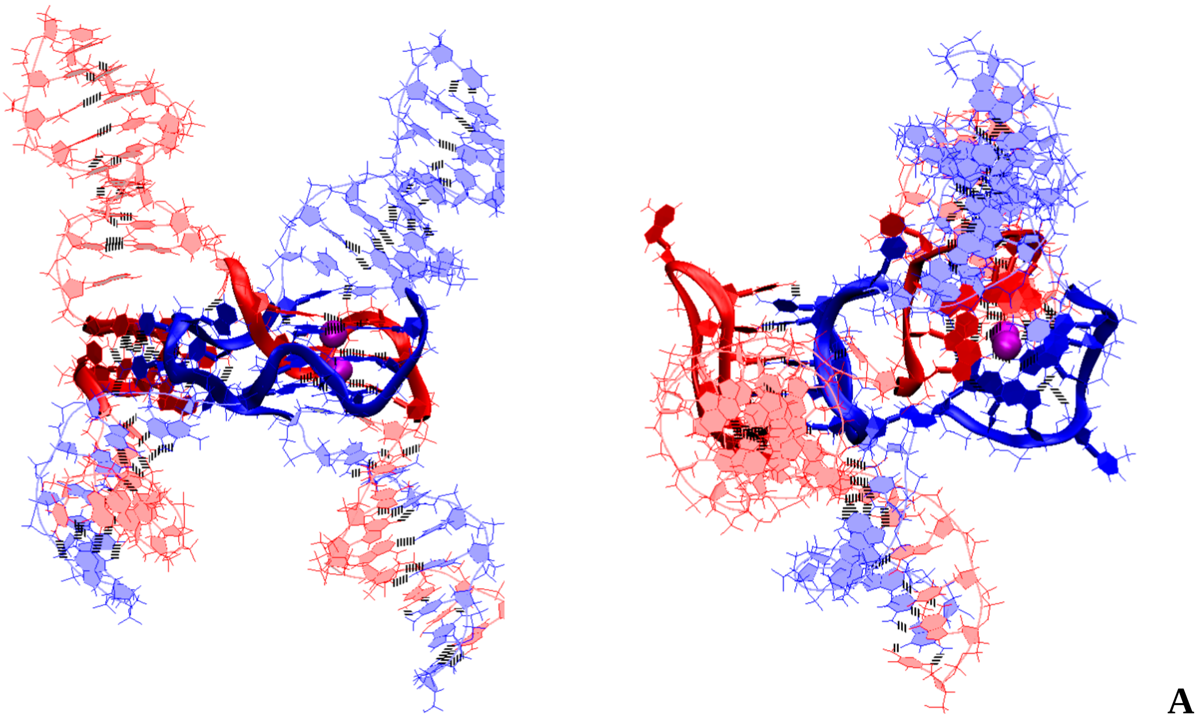


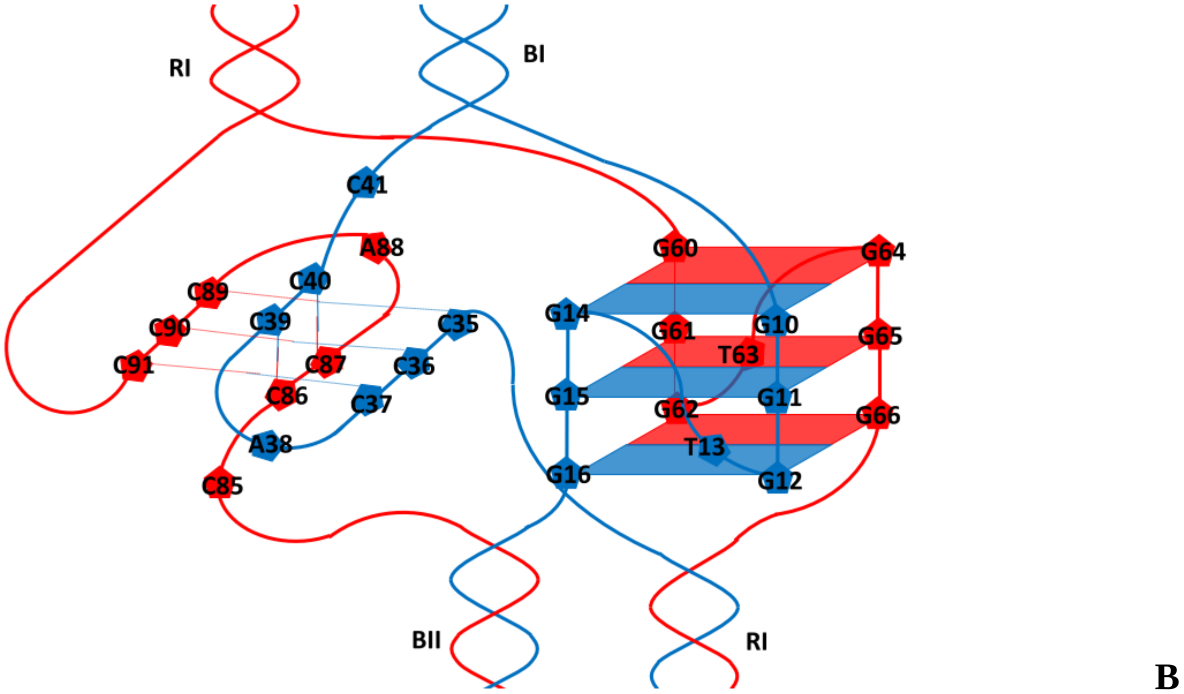


**Figure 1.C.S.1. Bimolecular complex with parallel G4-dimer and head-to-tail IM-dimer in case of the strands exchange: A** – conformation obtained at the last step of the MD trajectory (side and top view); **B** – the complex scheme.

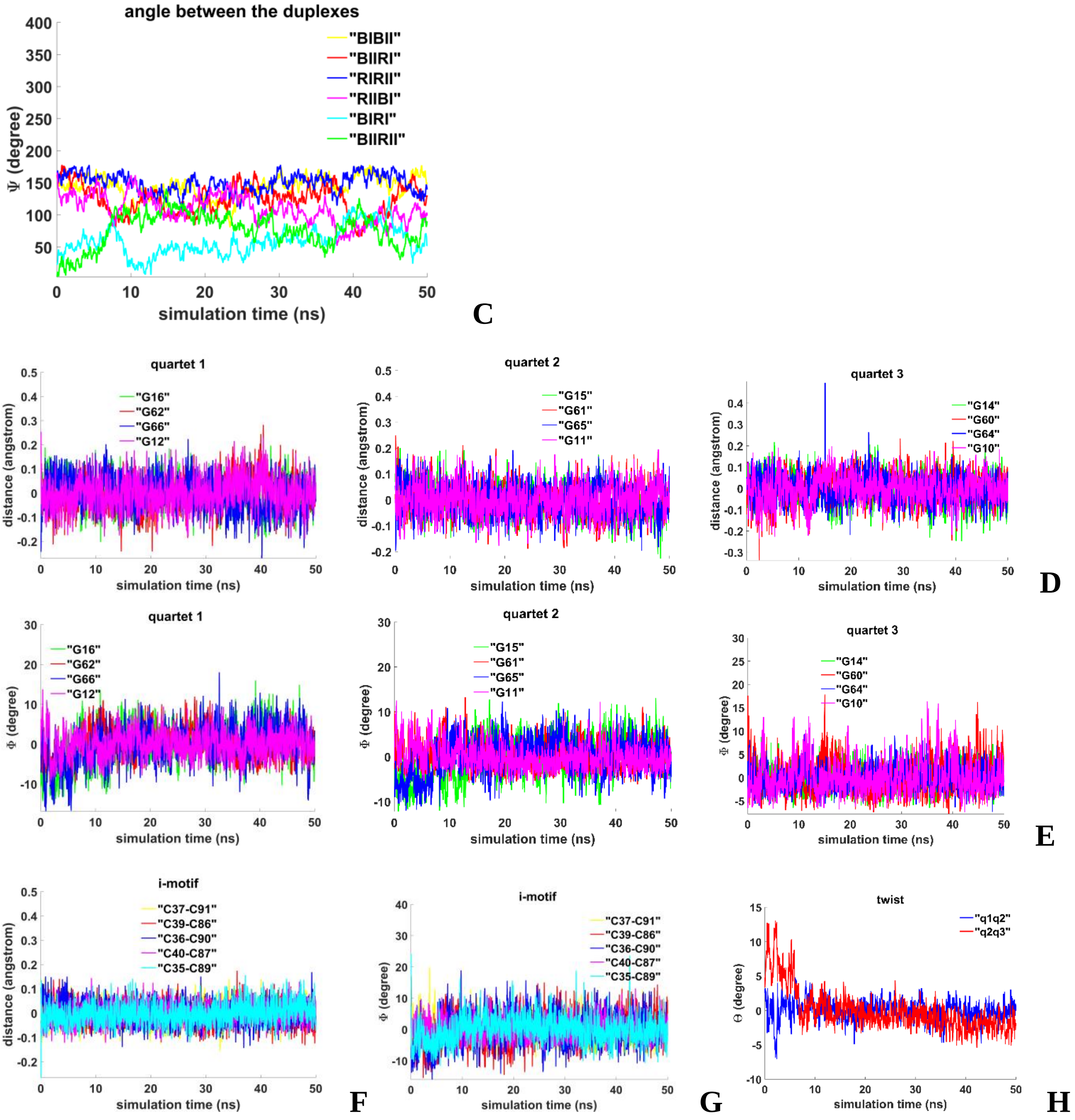


**Figure 1.C.S.2. Bimolecular complex with parallel G4-dimer and head-to-tail IM-dimer in case of the strands exchange: C** – angles between unmelted fragments of the duplexes; **D** - evolution of the values of distances from COMs of the guanines' bases to COMs of their containing tetrads, **E** - angles between normals to the guanines' bases and vectors connecting COMs of the boundary tetrads; **F** - distances between COMs of the cytosines' bases; **G** - angles between normals to the cytosines' bases; **H** – angles of rotation of the tetrads relative to each other.

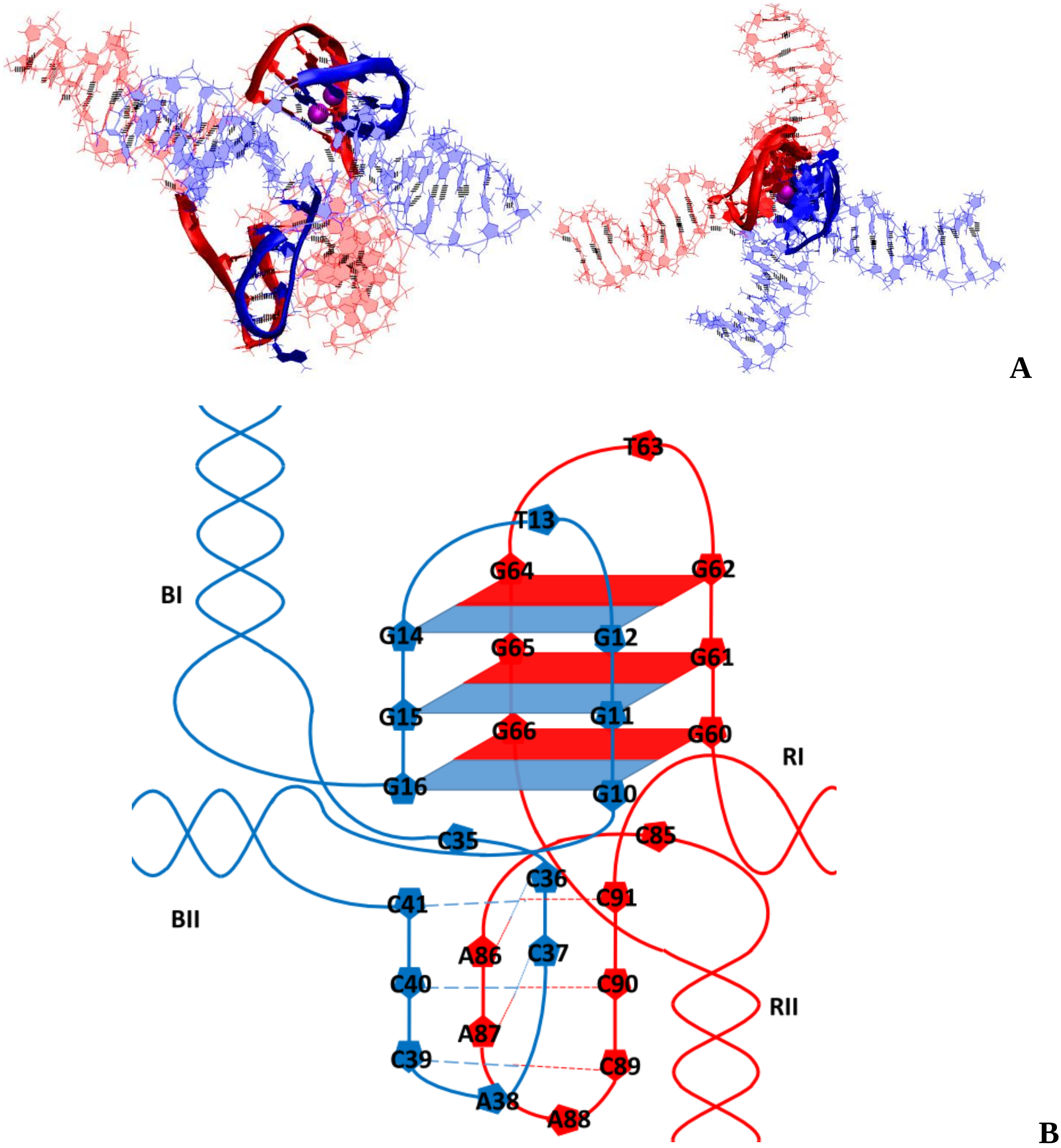


**Figure 1.D.S.1. Bimolecular complex with antiparallel G4-dimer and head-to-head IM-dimer: A** – conformation obtained at the last step of the MD trajectory (side and top view); **B** – the complex scheme.

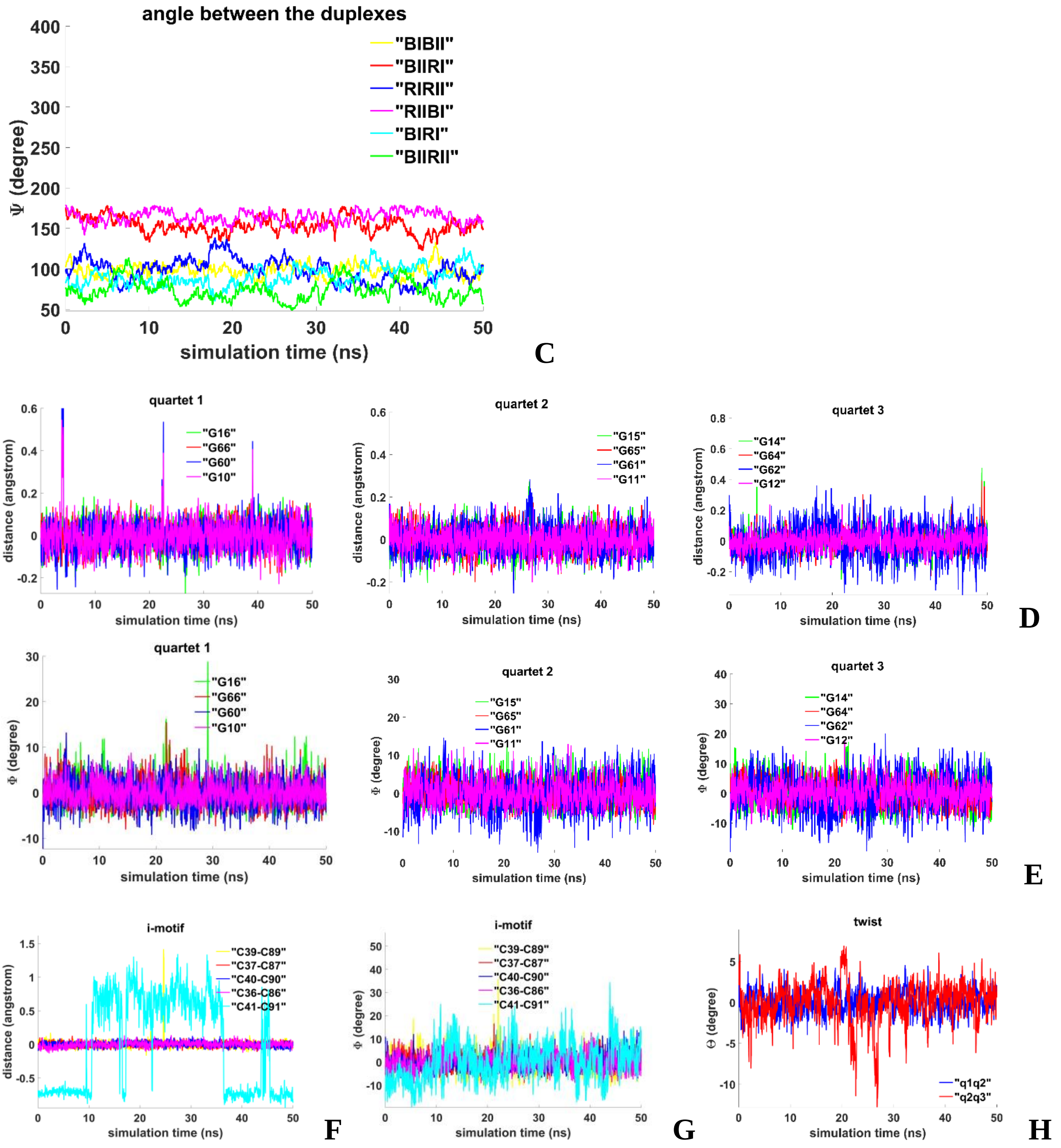


**Figure 1.D.S.2. Bimolecular complex with antiparallel G4-dimer and head-to-head IM-dimer: C** – angles between unmelted fragments of the duplexes; **D** - distances from COMs of the guanines' bases to COMs of their containing tetrads; **E** - angles between normals to the guanines' bases and vectors connecting COMs of the boundary tetrads; **F** - distances between COMs of the cytosines' bases; **G** - angles between normals to the cytosines' bases; **H** – angles of rotation of the tetrads relative to each other.

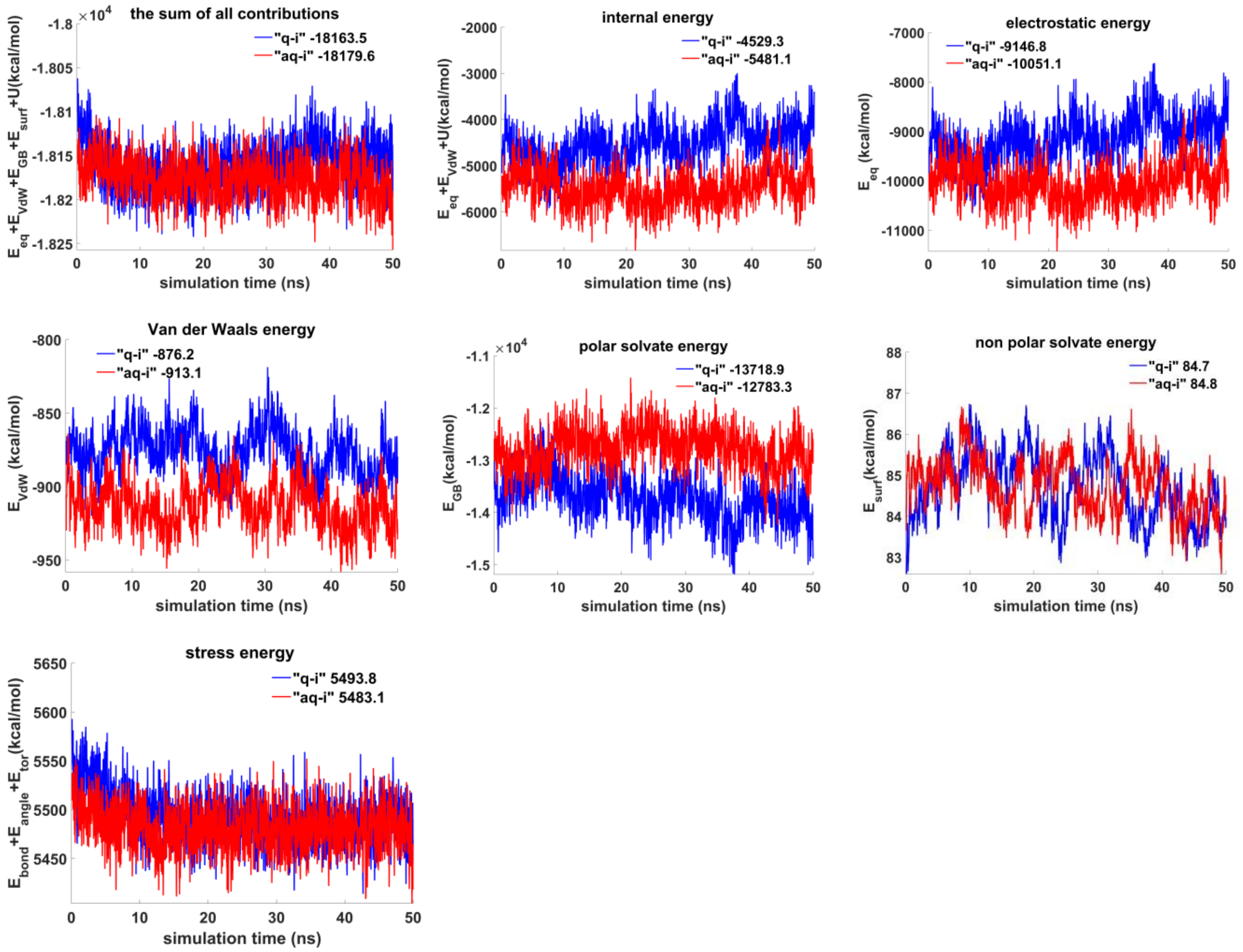


**Figure 1.S.E. The contributions to free energy for variants shown on Figure 2 and Figure 3 during MD calculations.** $E_{eq}$ – electrostatic, $E_{vdw}$ - Van der Waals, $E_{GB}$ - polar energy of solvation, $E_{surf}$ - non-polar energy of solvation due to the hydrophobic surface available to the solvent, $U = E_{bond} + E_{angle} + E_{tor}$ , e.g. $E_{bond}$, $E_{angle}$ and $E_{tor}$ – bond, angle and torsion stress energies. The energy plots were smoothed using moving average method (span = 5). Average energy values are indicated in the figure legends.

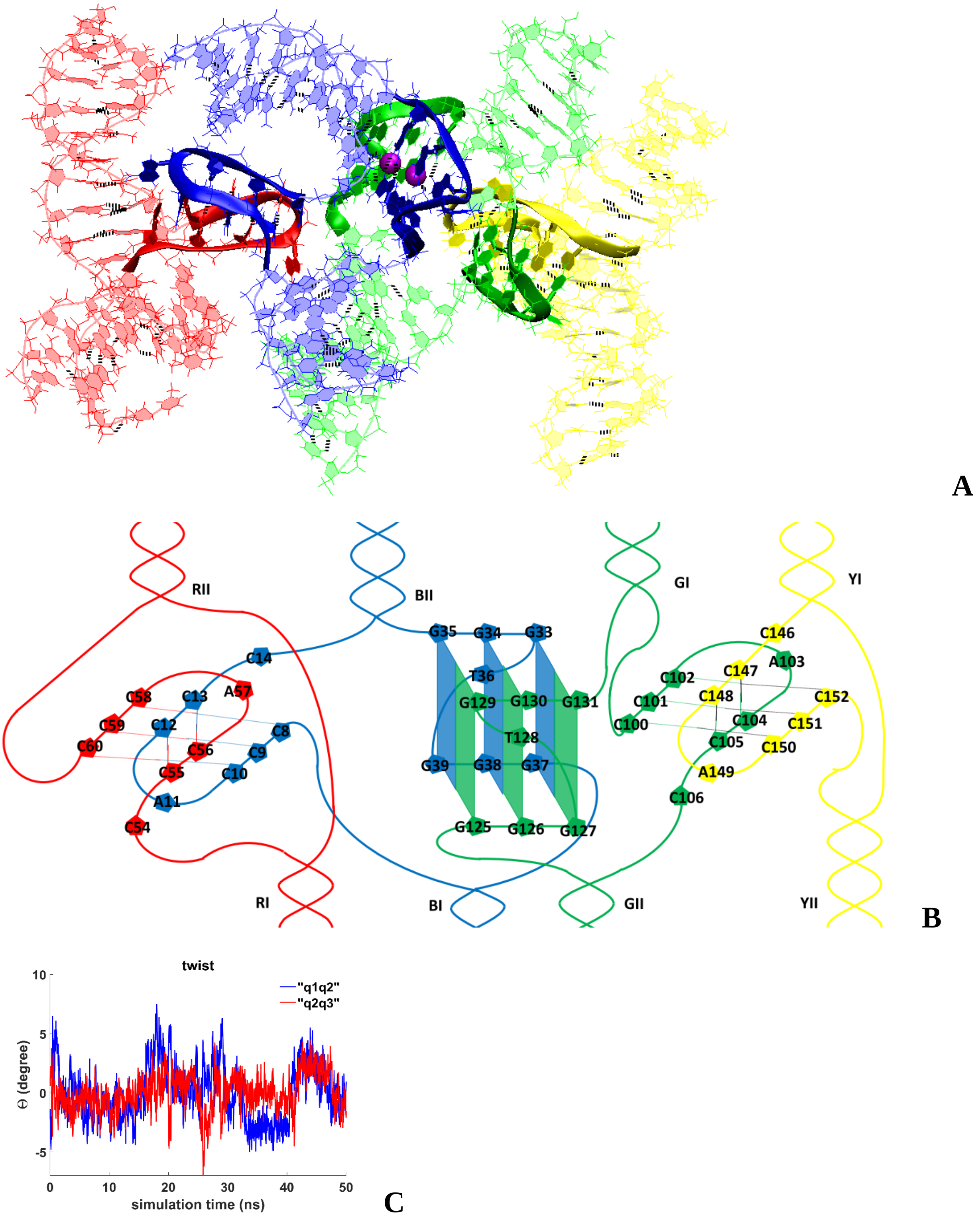


**Figure 2.A.S.1. Tetrameric complex with parallel G4-dimer and two head-to-tail IM-dimers: A** – conformation obtained at the last step of the MD trajectory (side and top view); **B** – the complex scheme; **C** – angles of rotation of the tetrads relative to each other.

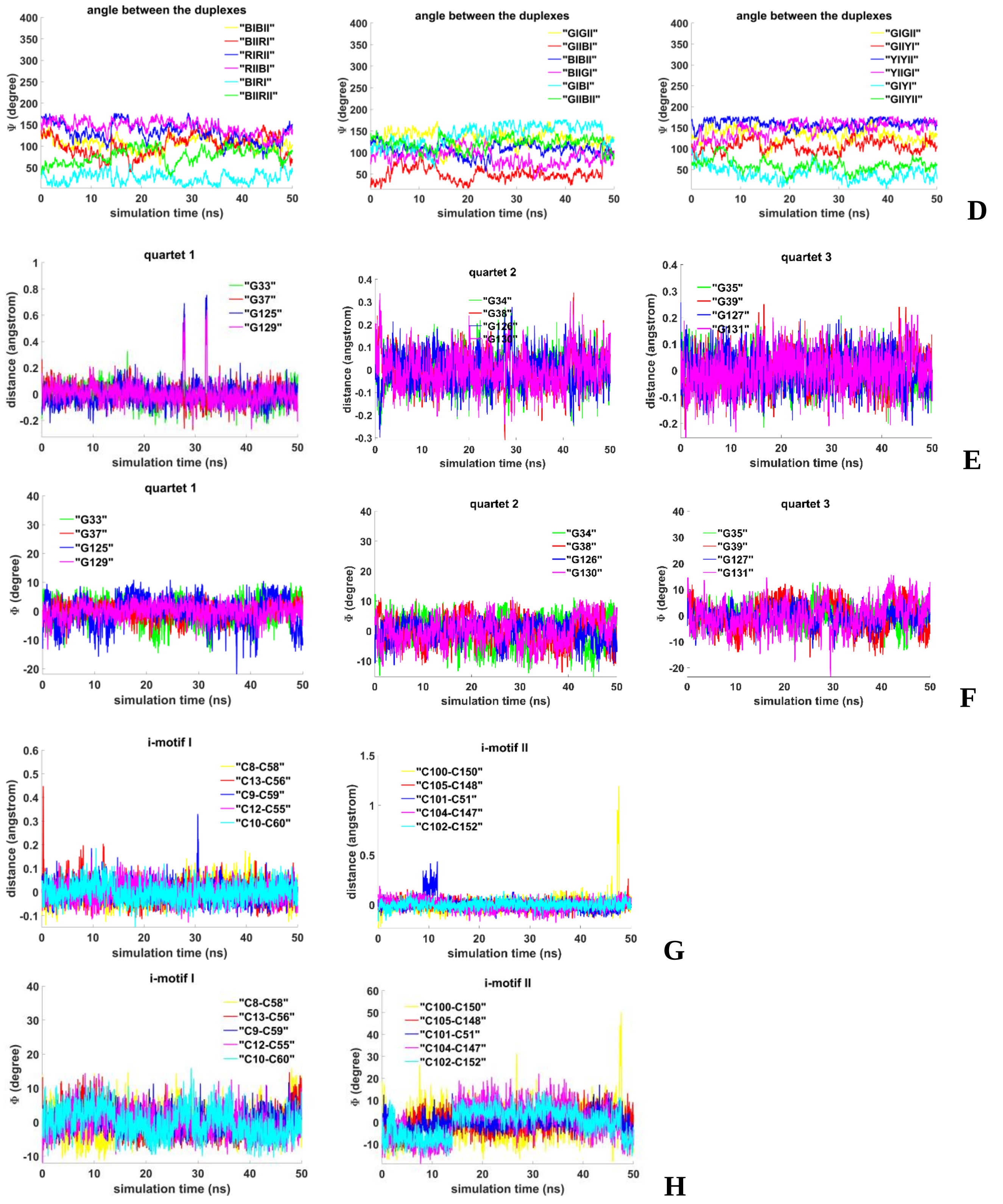


**Figure 2.A.S.2. Tetrameric complex with parallel G4-dimer and two head-to-tail IM-dimers: Tetrameric complex with parallel G4-dimer and two head-to-tail IM-dimers: D**– angles between unmelted fragments of the duplexes; **E** - distances from COMs of the guanines' bases to COMs of their containing tetrads; **F** - angles between normals to the guanines' bases and vectors connecting COMs of the boundary tetrads; **G** - distances between COMs of the cytosines' bases; **H** - angles between normals to the cytosines' bases.

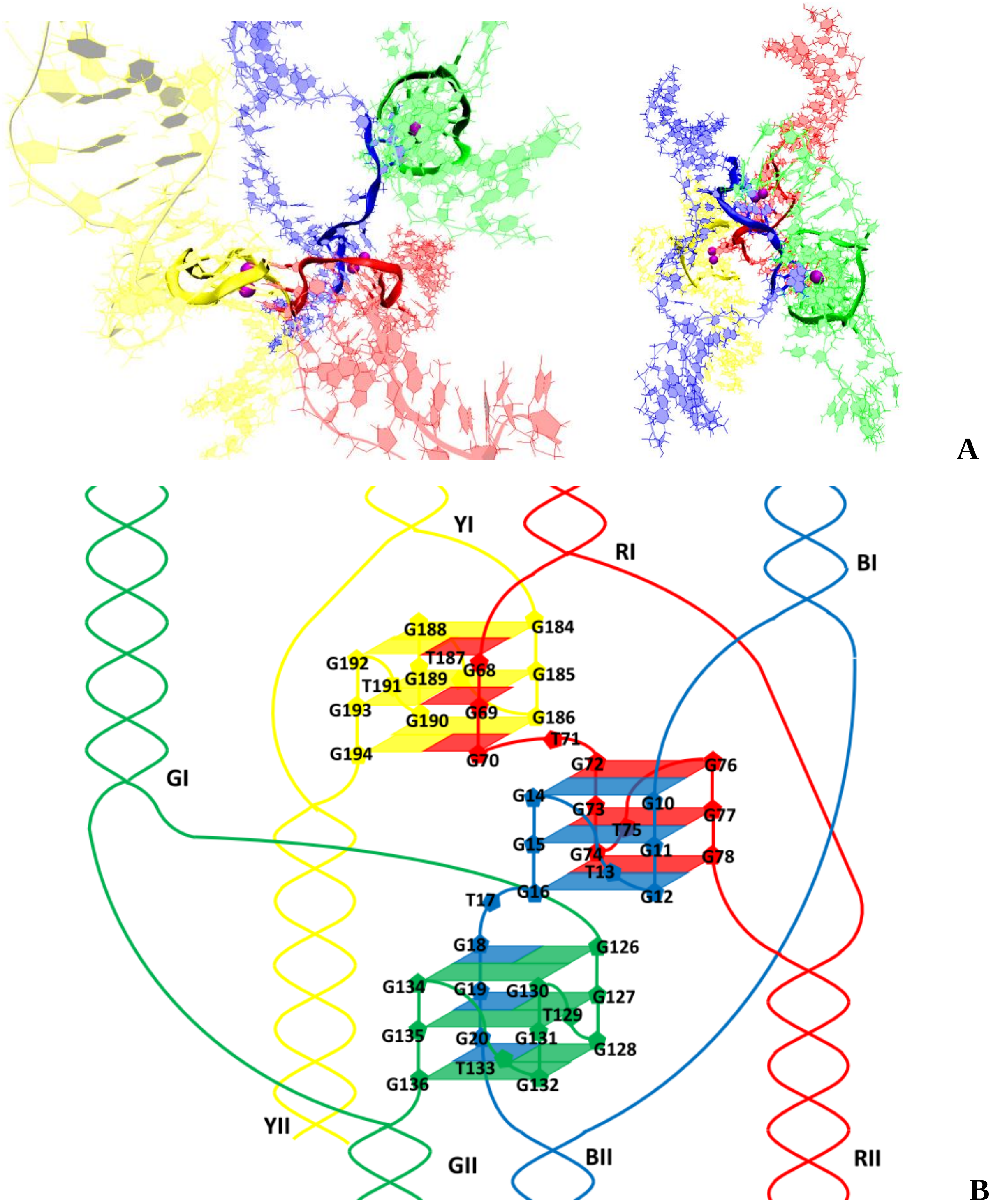


**Figure 2.B.S.1. Tetrameric complex with three parallel G4-dimers: A** – conformation obtained at the last step of the MD trajectory (side and top view); **B** – the complex scheme.

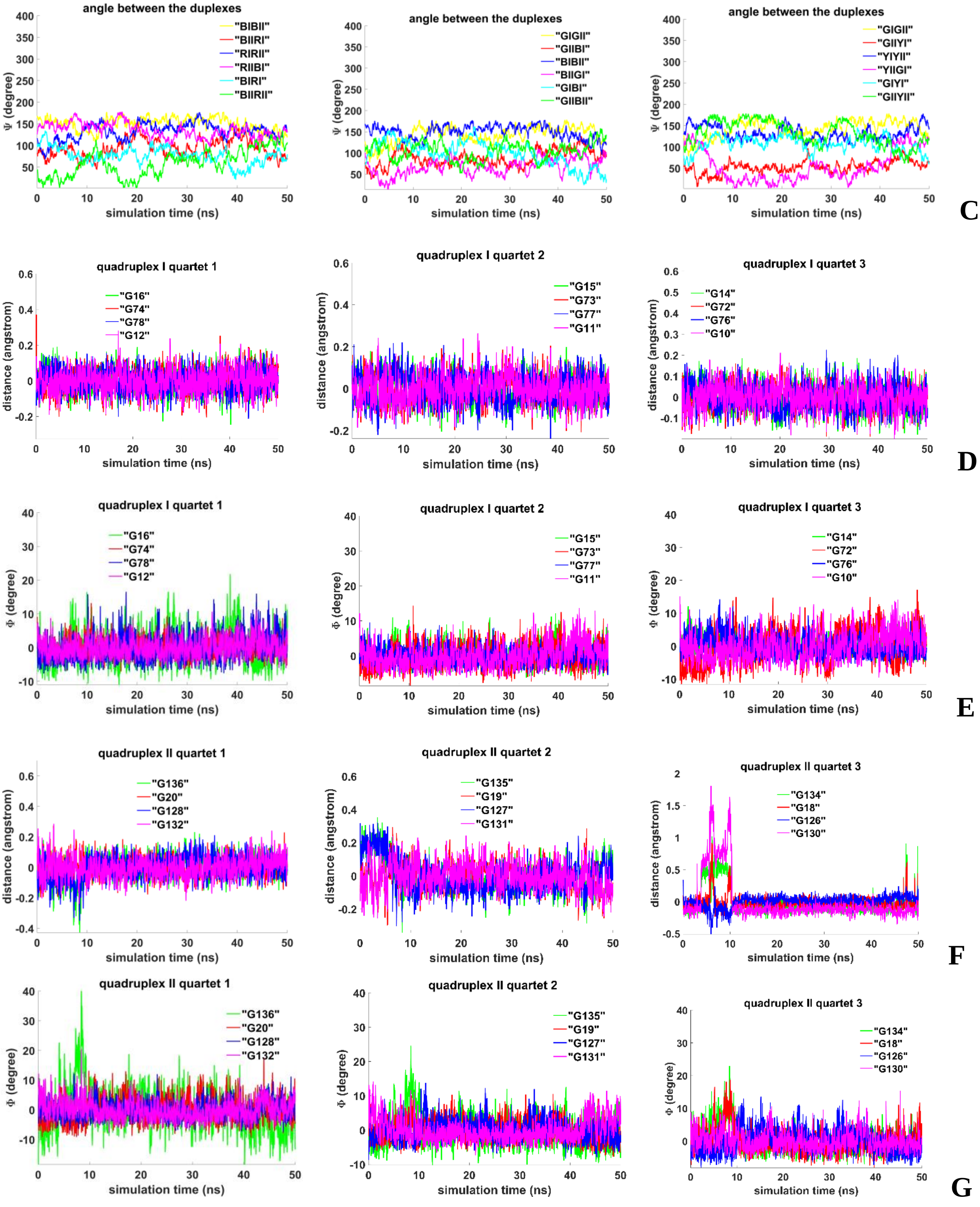


**Figure 2.B.S.2. Tetrameric complex with three parallel G4-dimers: C** – angles between unmelted fragments of the duplexes; **D, F** - distances from COMs of the guanines' bases to COMs of their containing tetrads; **E, G** - angles between normals to the guanines' bases and vectors connecting COMs of the boundary tetrads.

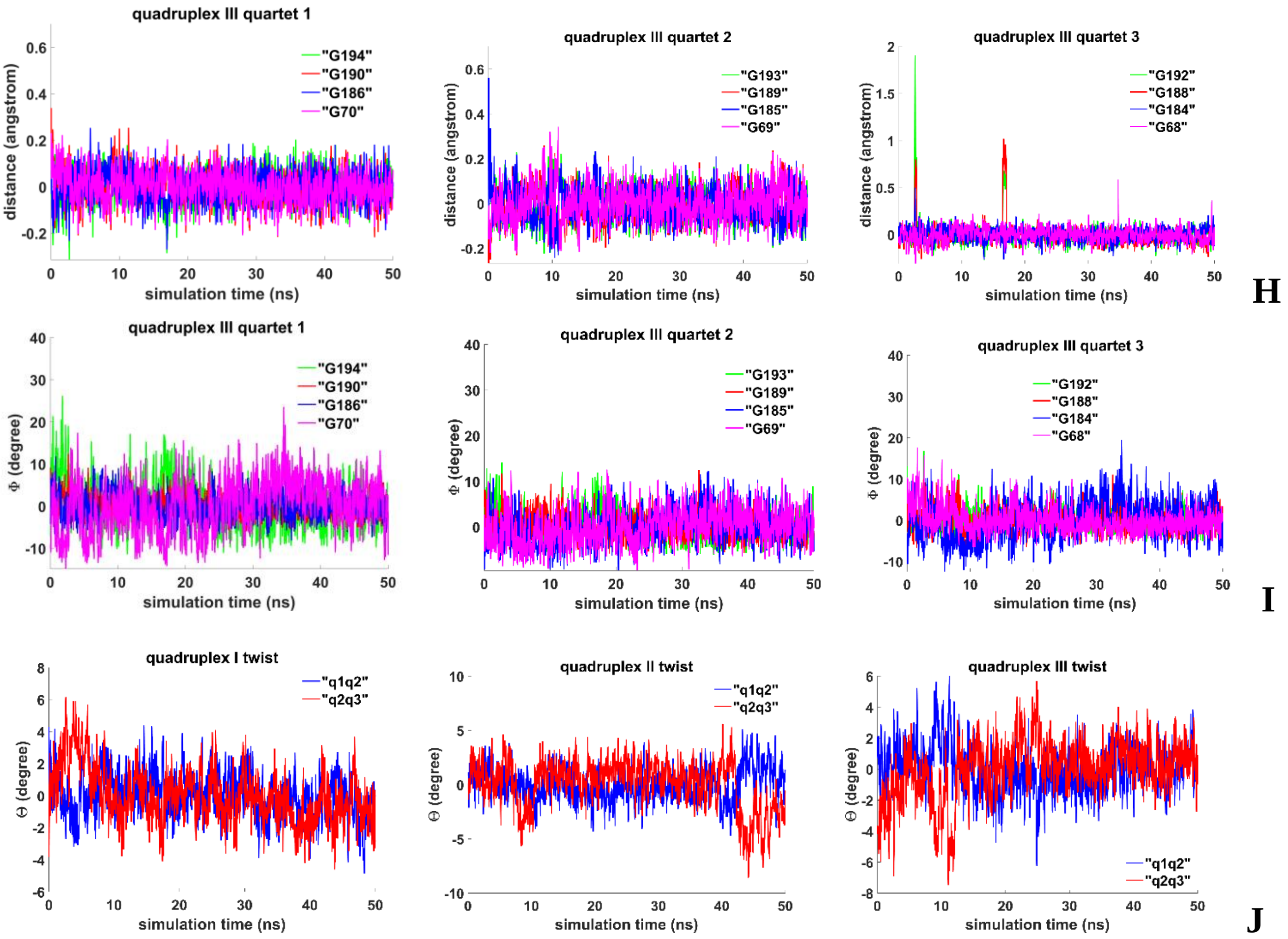


**Figure 2.B.S.3. Tetrameric complex with three parallel G4-dimers: H** - distances from COMs of the guanines' bases to COMs of their containing tetrads; **I** - angles between normals to the guanines' bases and vectors connecting COMs of the boundary tetrads; **J** - angles of rotation of the tetrads relative to each other.

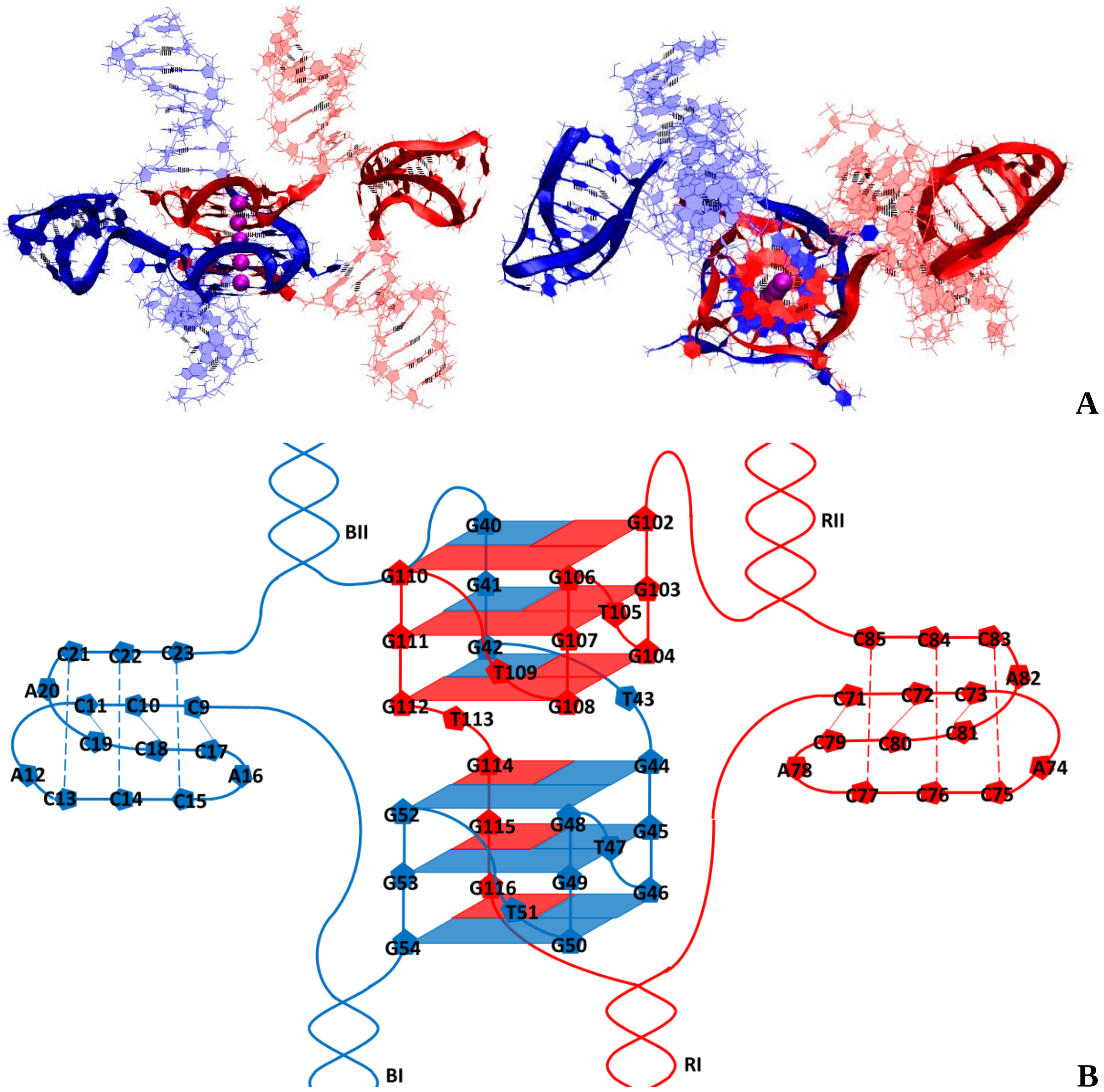


**Figure 3.1.A.S.1. Case 1 "1,3 hitch and two IM-monomers": A** – conformation obtained at the last step of the MD trajectory (side and top view); **B** – the complex scheme.

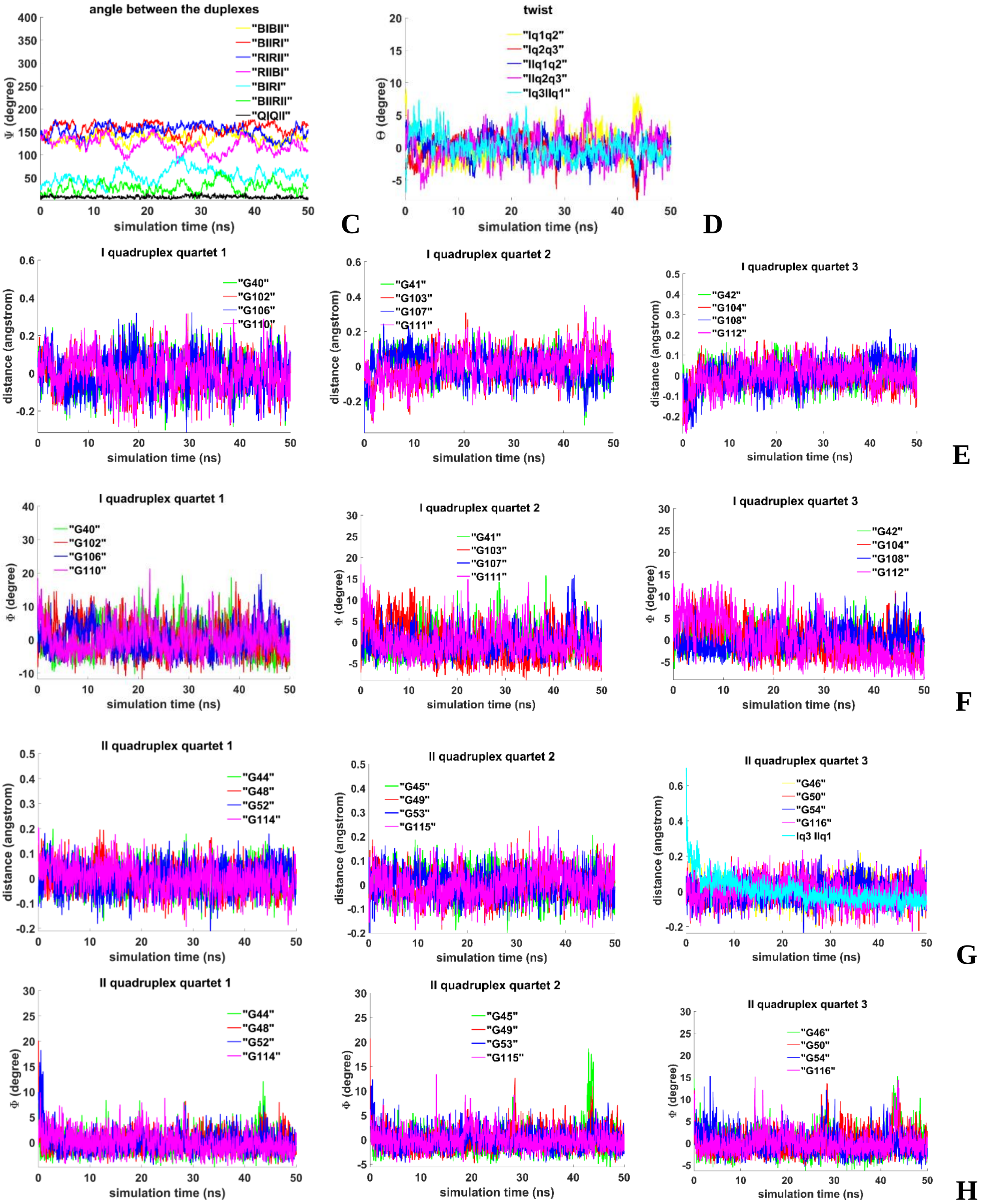


**Figure 3.1.A.S.2. Case 1 “1,3 hitch and two IM-monomers”: C** – angles between unmelted fragments of the duplexes, angle between axes passing through COMs of the boundary tetrads (**QIQII**); **D**– angles of rotation of the tetrads relative to each other; **E, G** - distances from COMs of the guanines’ bases to COMs of their containing tetrads, distance between COMs of the boundary tetrads (**Iq3 IIq1**); **F, H** - angles between normals to the guanines’ bases and vectors connecting COMs of the boundary tetrads.

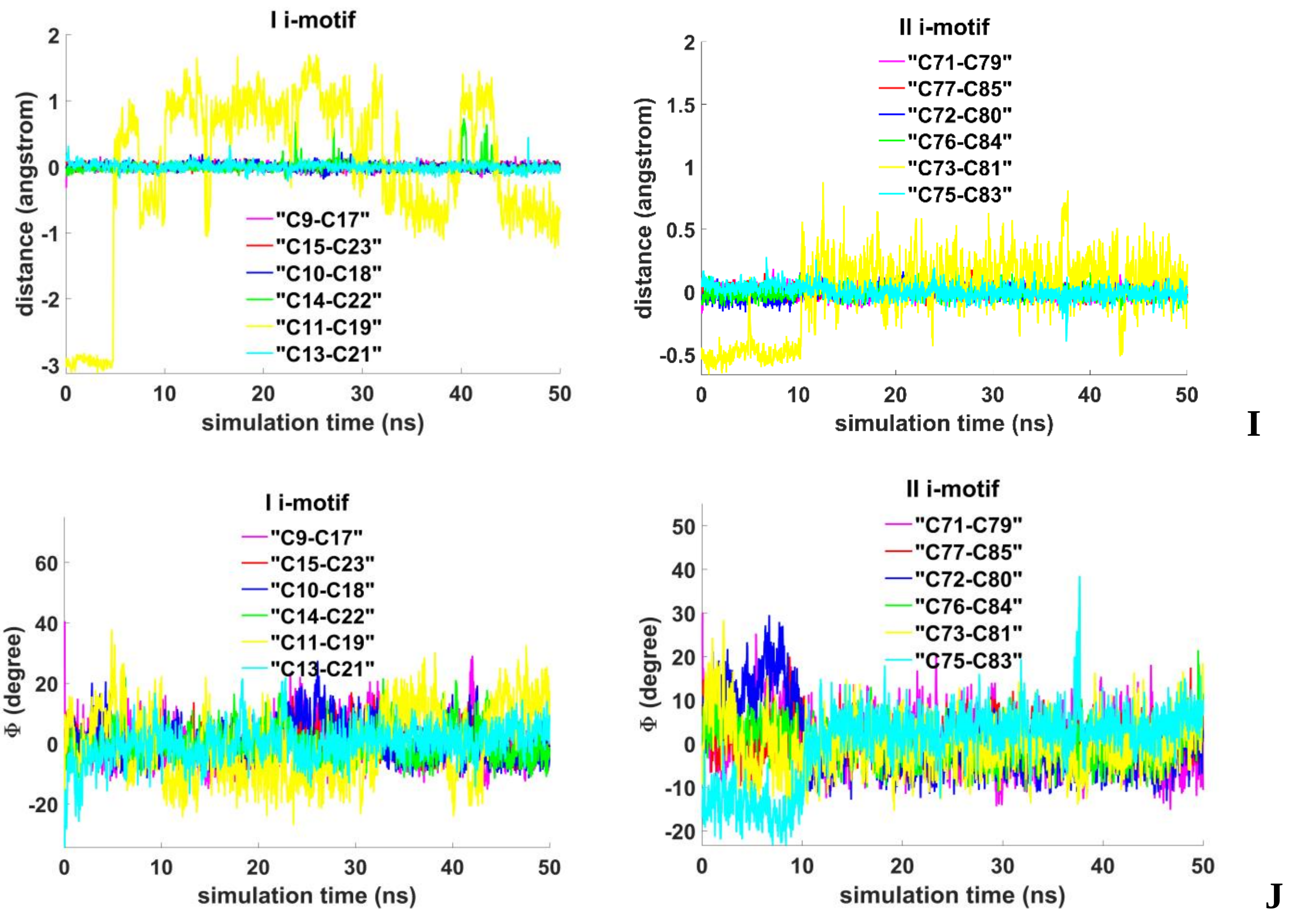


**Figure 3.1.A.S.3. Case 1 "1,3 hitch and two IM-monomers": I -** distances between COMs of the cytosines' bases; **J -** angles between normals to the cytosines' bases.

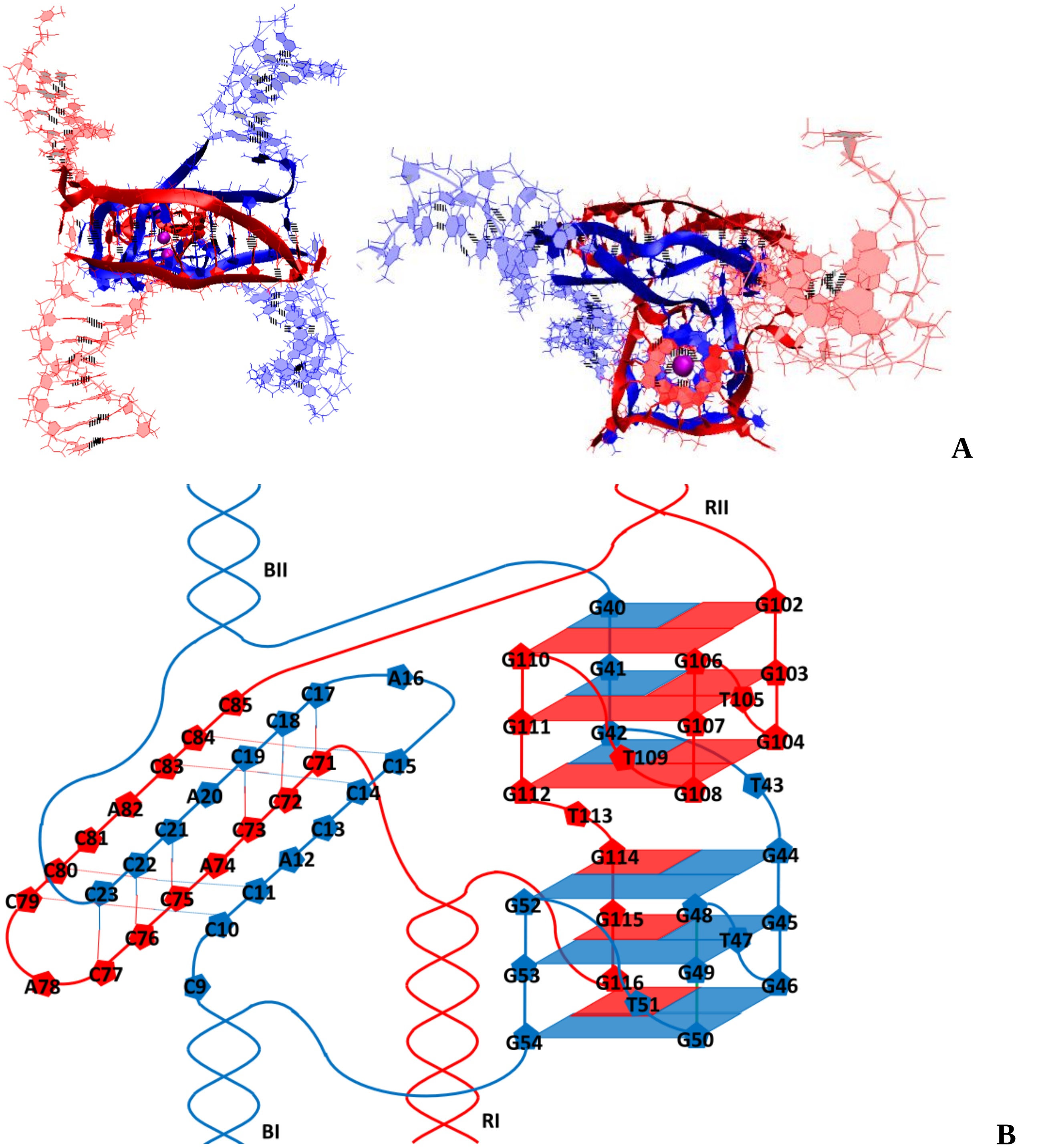


**Figure 3.1.B.S.1. Case 2: "1,3 hitch and head-to-tail IM-dimer"**: **A** – conformation obtained at the last step of the MD trajectory (side and top view); **B** – the complex scheme.

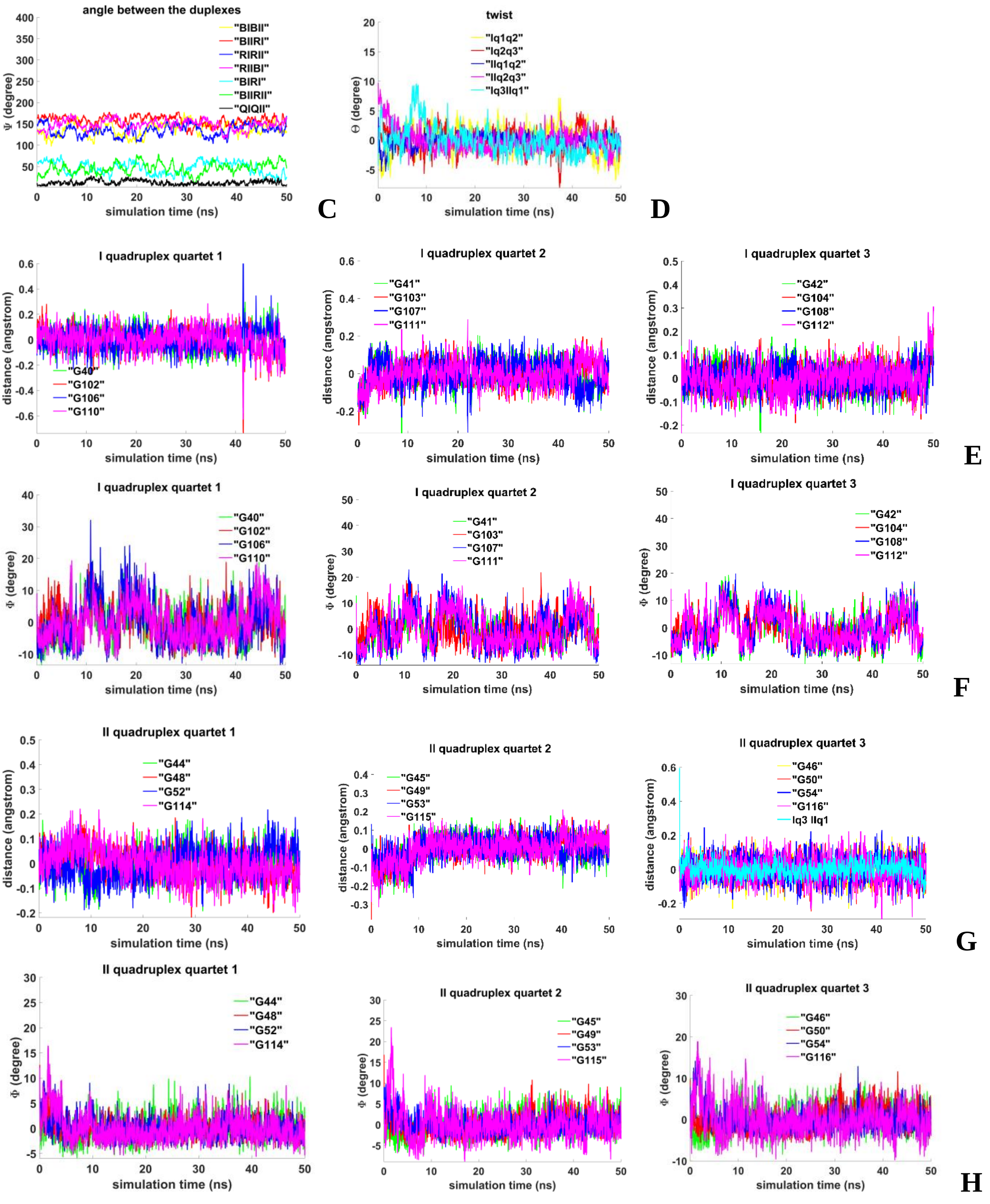


**Figure 3.1.B.S.2. Case 2: "1,3 hitch and head-to-tail IM-dimer"**: **C** – angles between unmelted fragments of the duplexes, angle between axes passing through COMs of the boundary tetrads (**QIQII**); **D**– angles of rotation of the tetrads relative to each other; **E, G** - distances from COMs of the guanines' bases to COMs of their containing tetrads, distance between COMs of the boundary tetrads (**Iq3 IIq1**); **F, H** - angles between normals to the guanines' bases and vectors connecting COMs of the boundary tetrads.

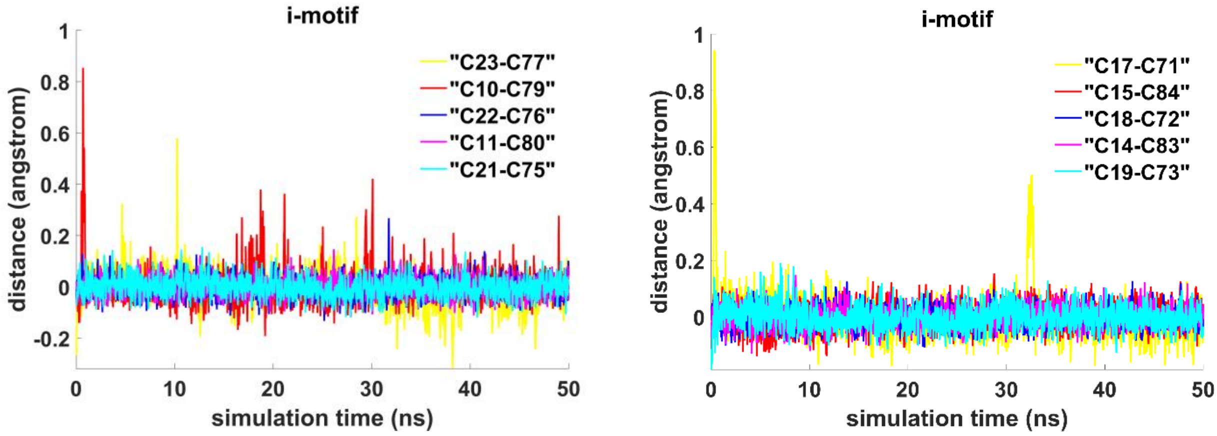


I

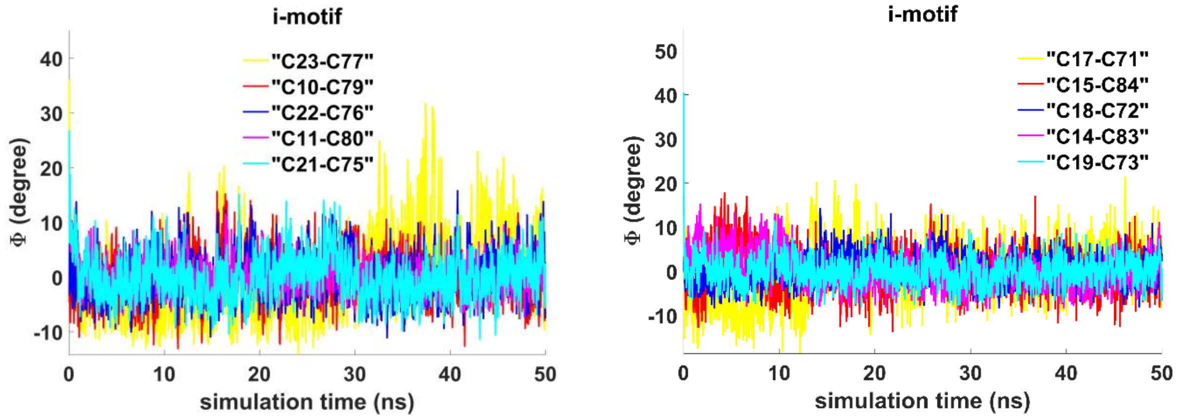


J

**Figure 3.1.B.S.3. Case 2: "1,3 hitch and head-to-tail IM-dimer": I -** distances between COMs of the cytosines' bases; **J -** angles between normals to the cytosines' bases.

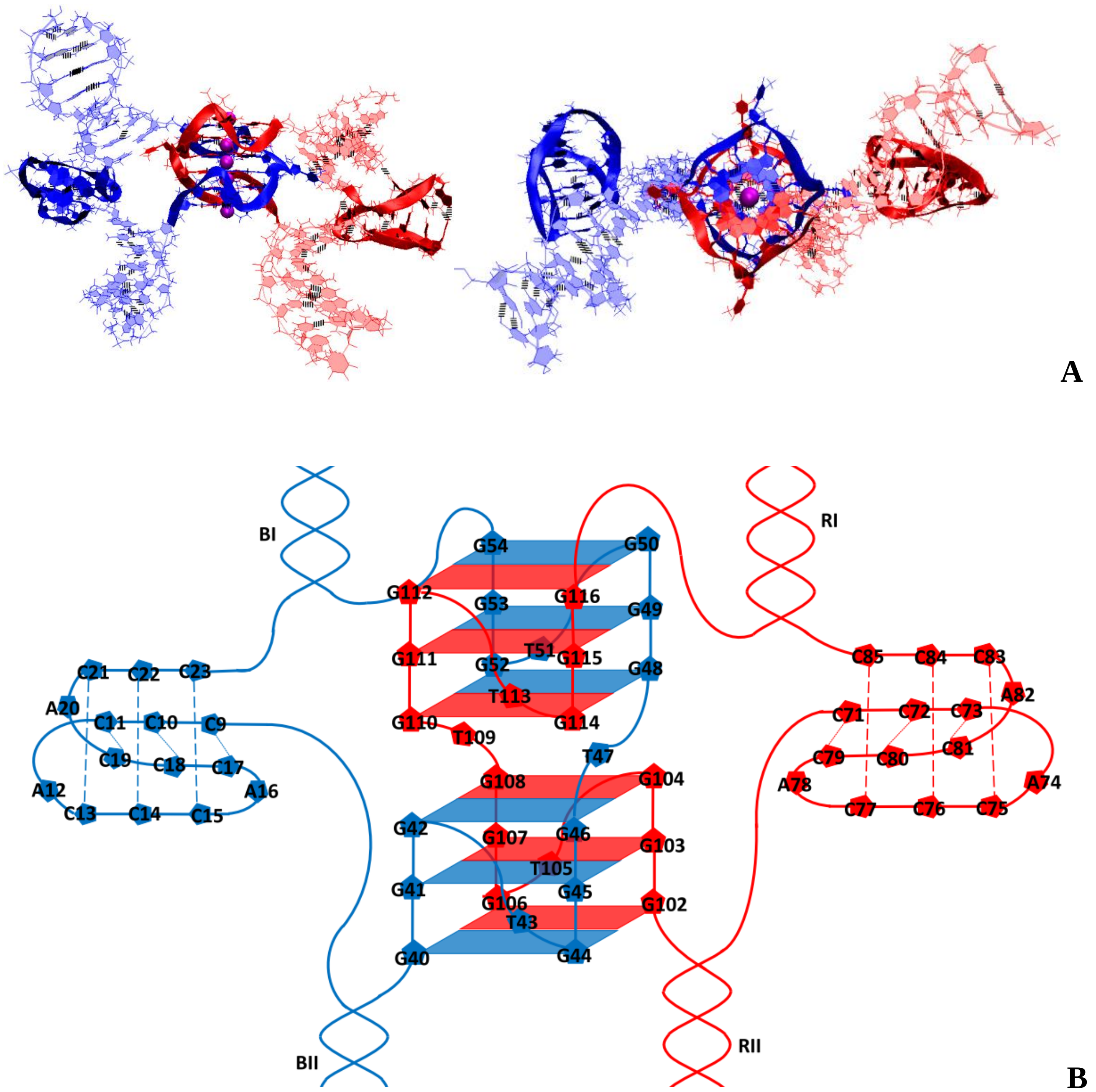


**Figure 3.1.C.S.1. Case 3: "1,2 hitch and two head-to-head IM-monomers with mutual girth of the strands"**: **A** – conformation obtained at the last step of the MD trajectory (side and top view); **B** – the complex scheme.

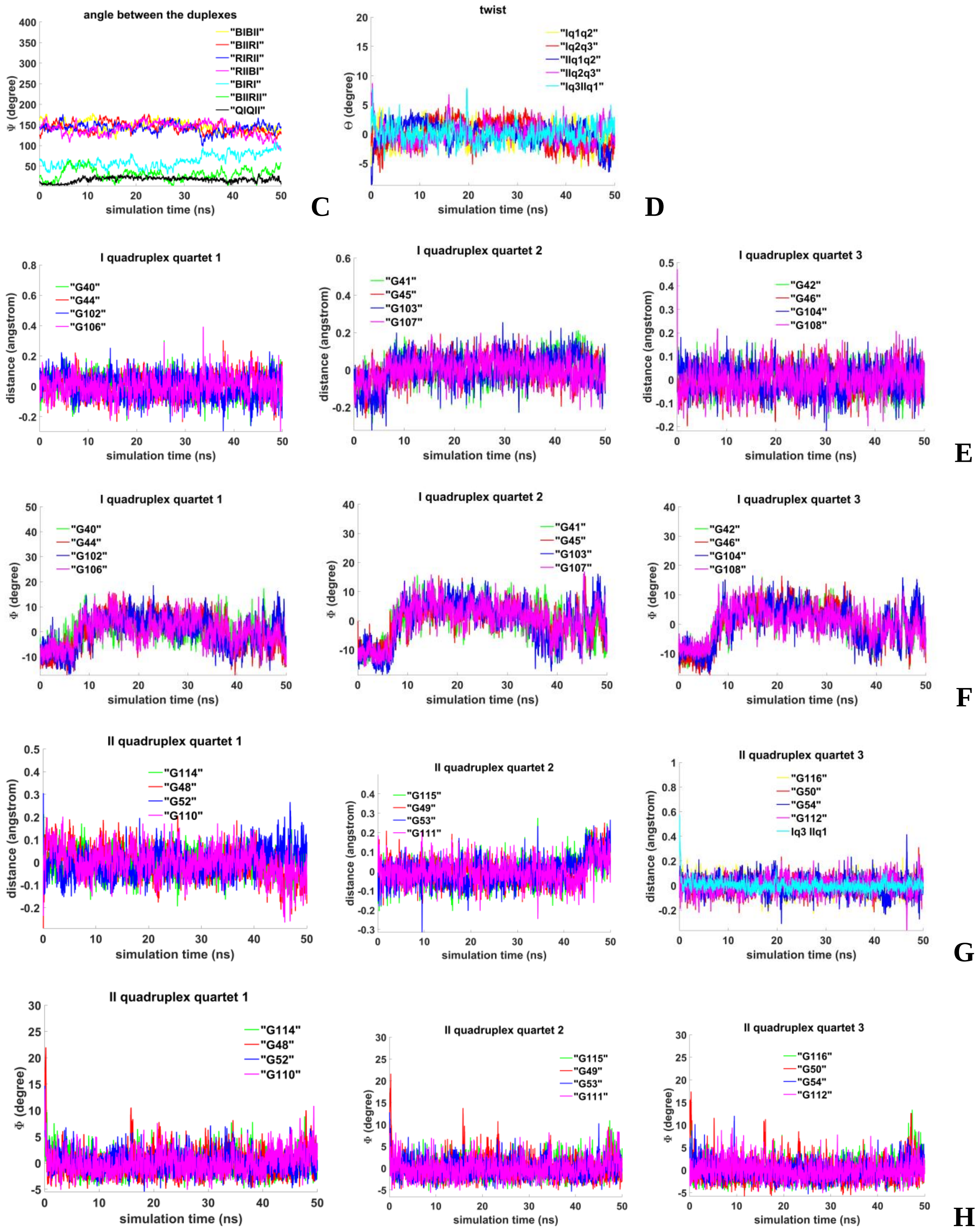


**Figure 3.1.C.S.2. Case 3: "1,2 hitch and two head-to-head IM-monomers with mutual girth of the strands"**: **C** – angles between unmelted fragments of the duplexes, angle between axes passing through COMs of the boundary tetrads (**QIQII**); **D**– angles of rotation of the tetrads relative to each other; **E, G** - distances from COMs of the guanines' bases to COMs of their containing tetrads, distance between COMs of the boundary tetrads (**Iq3 IIq1**); **F, H** - angles between normals to the guanines' bases and vectors connecting COMs of the boundary tetrads.

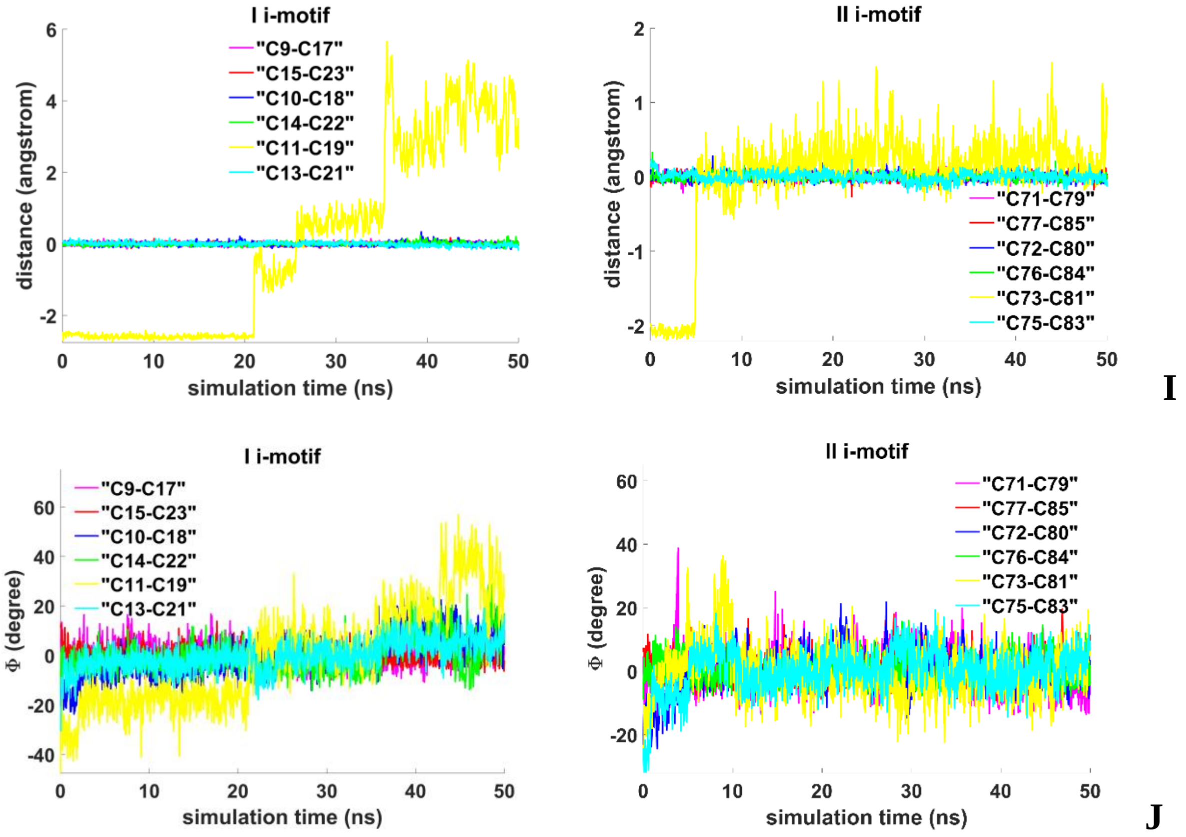


**Figure 3.1.C.S.3. Case 3: "1,2 hitch and two head-to-head IM-monomers with mutual girth of the strands"**: **I -** distances between COMs of the cytosines' bases; **J** - angles between normals to the cytosines' bases.

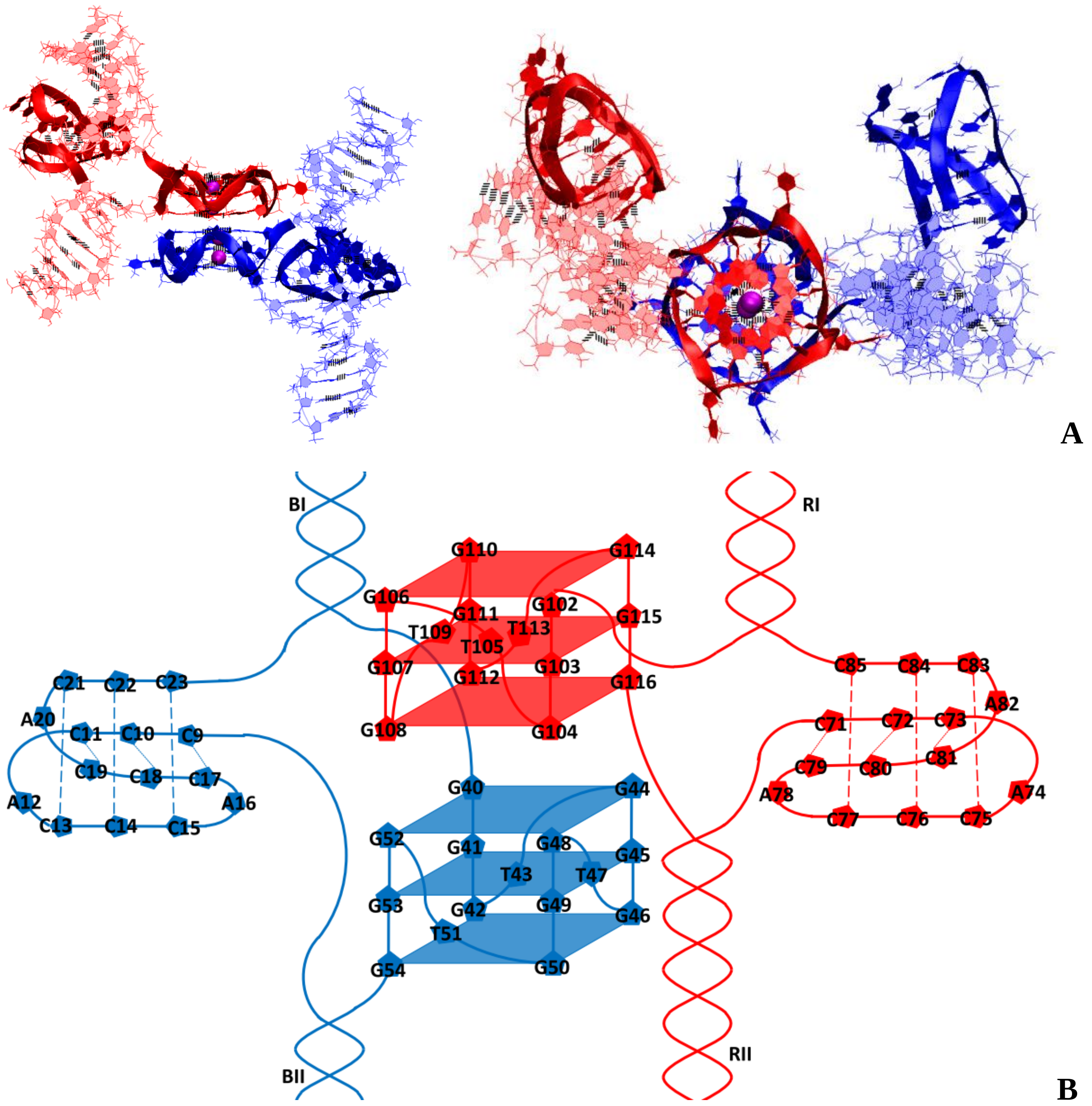


**Figure 3.2.A.S.1. Case 4: "Stacking with 2 IM-monomers"**: **A** – conformation obtained at the last step of the MD trajectory (side and top view); **B** – the complex scheme.

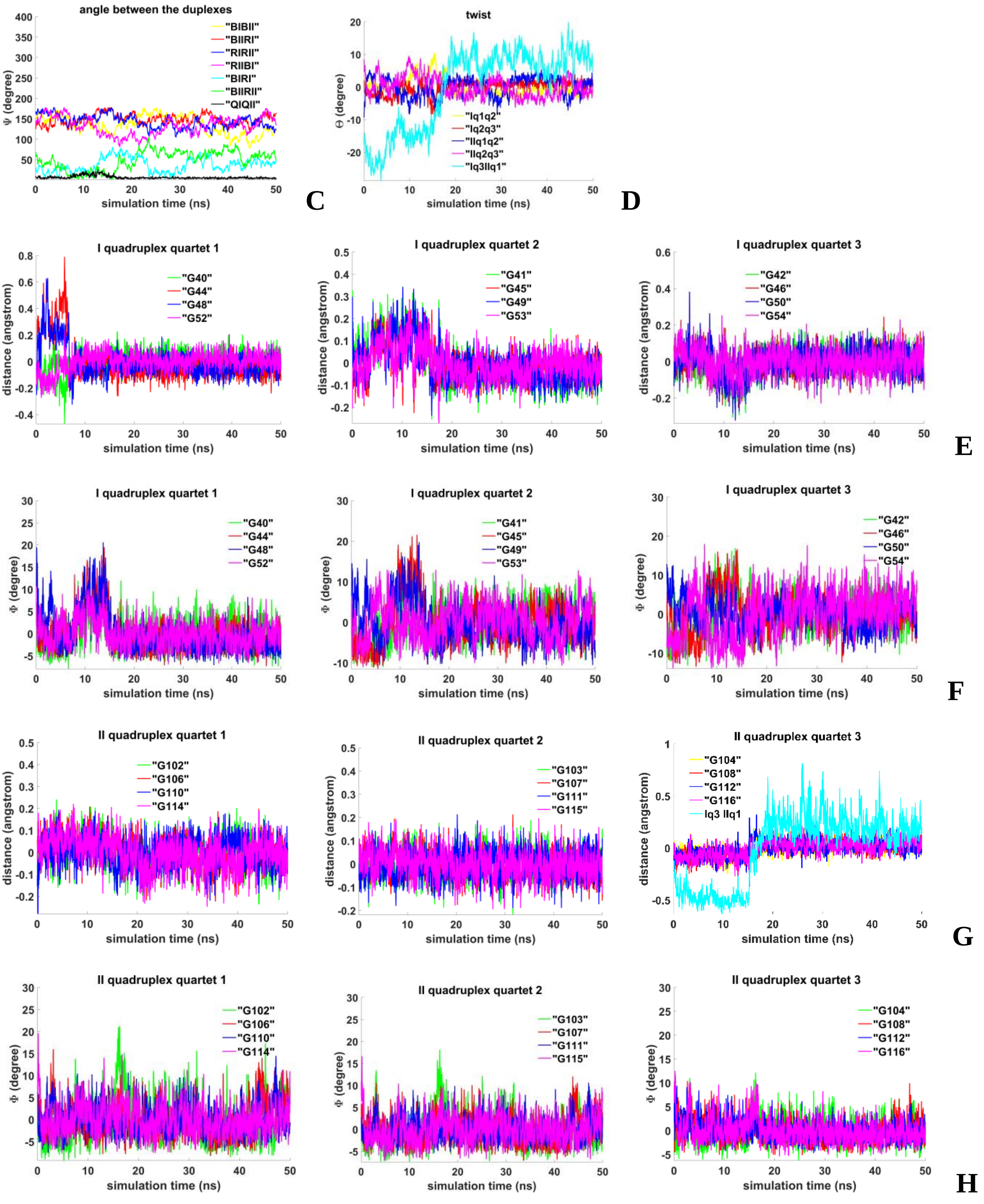


**Figure 3.2.A.S.2. Case 4: "Stacking with 2 IM-monomers"**: **C** – angles between unmelted fragments of the duplexes, angle between axes passing through COMs of the boundary tetrads (**QIQII**); **D**– angles of rotation of the tetrads relative to each other; **E, G** - distances from COMs of the guanines' bases to COMs of their containing tetrads, distance between COMs of the boundary tetrads (**Iq3 IIq1**); **F, H** - angles between normals to the guanines' bases and vectors connecting COMs of the boundary tetrads.

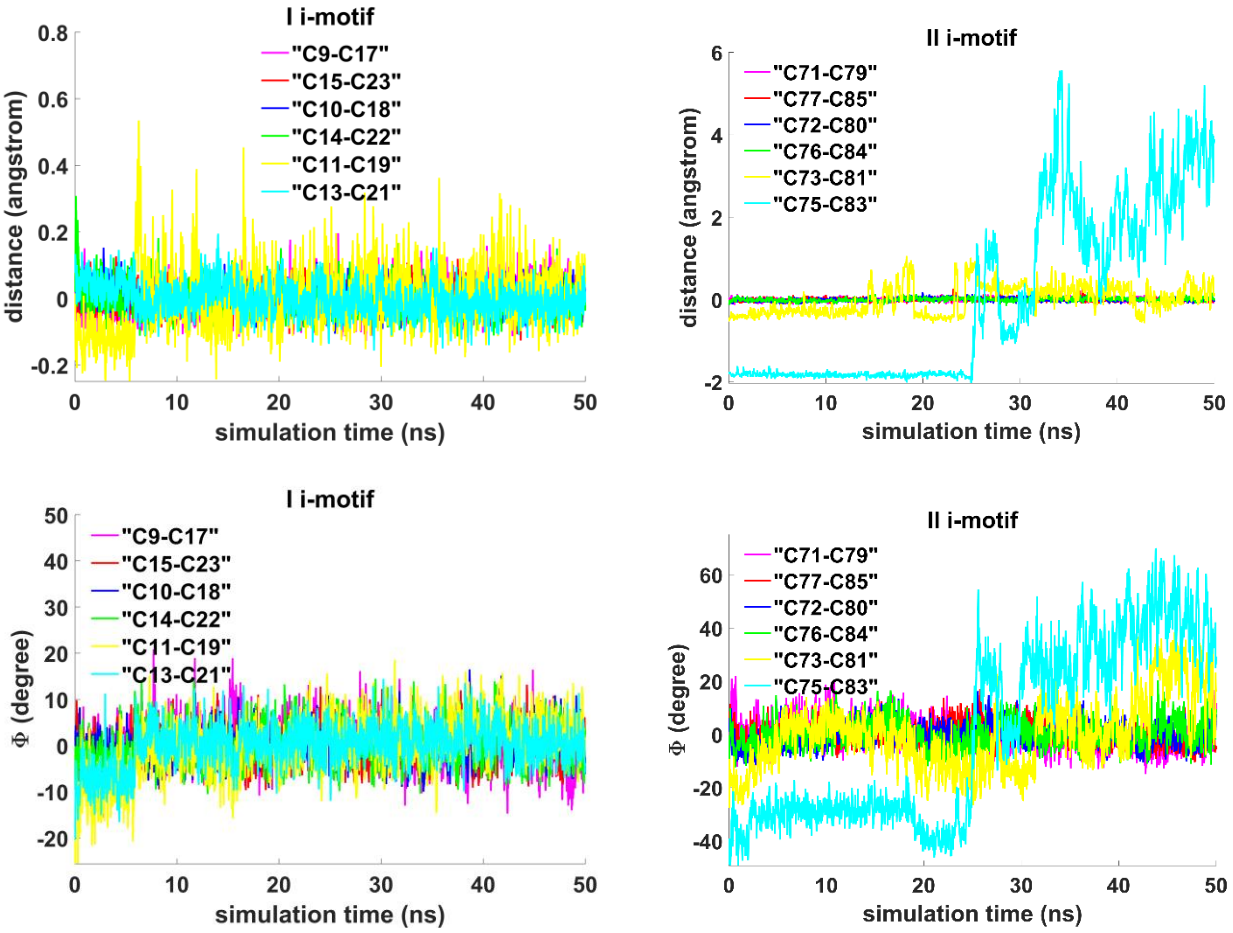


I

J

**Figure 3.2.A.S.3. Case 4: "Stacking with 2 IM-monomers"**: **I -** distances between COMs of the cytosines' bases; **J** - angles between normals to the cytosines' bases.

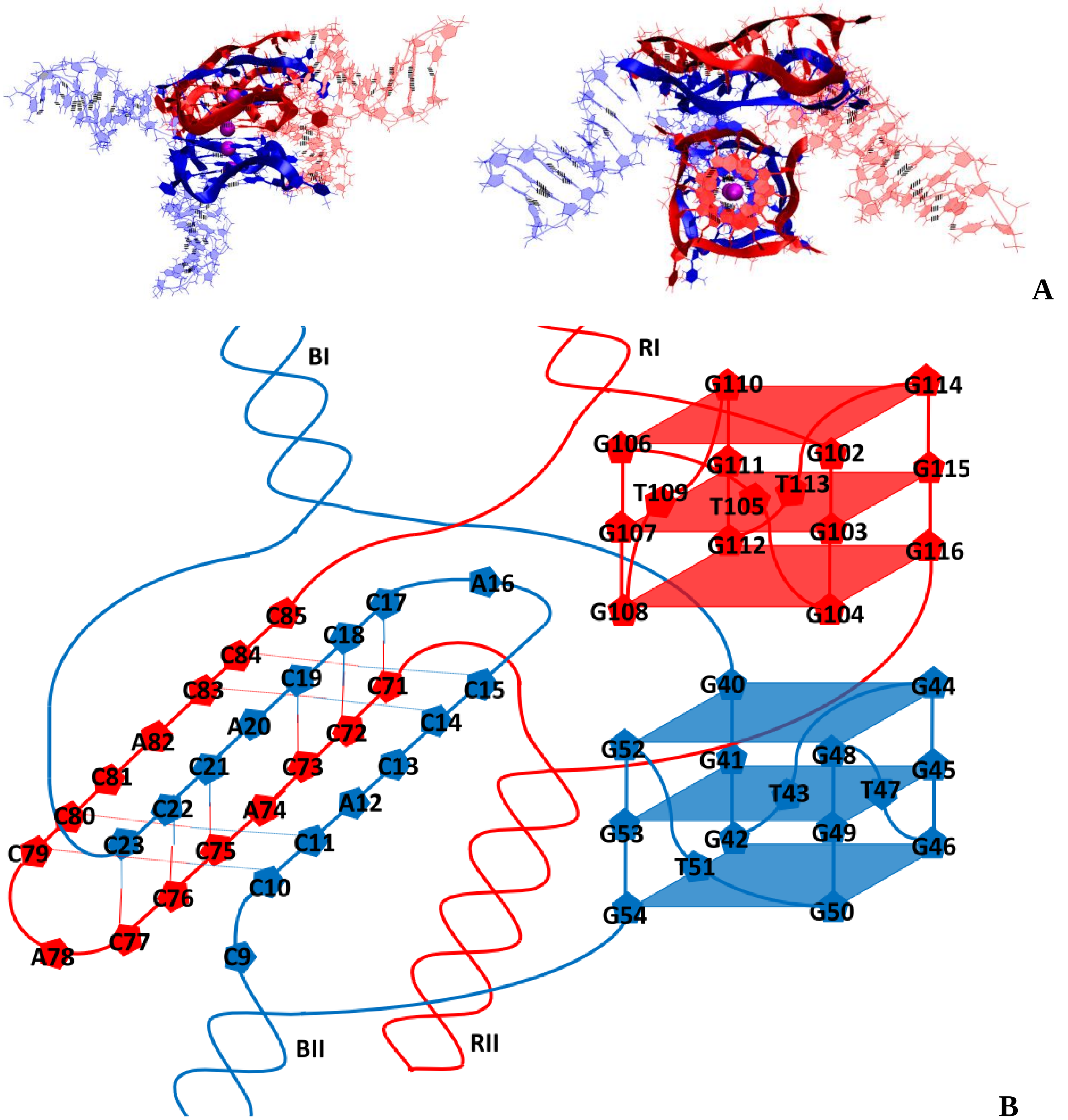


**Figure 3.2.B.S.1. Case 5: "Stacking and head-to-tail IM-dimer"**: **A** – conformation obtained at the last step of the MD trajectory (side and top view); **B** – the complex scheme.

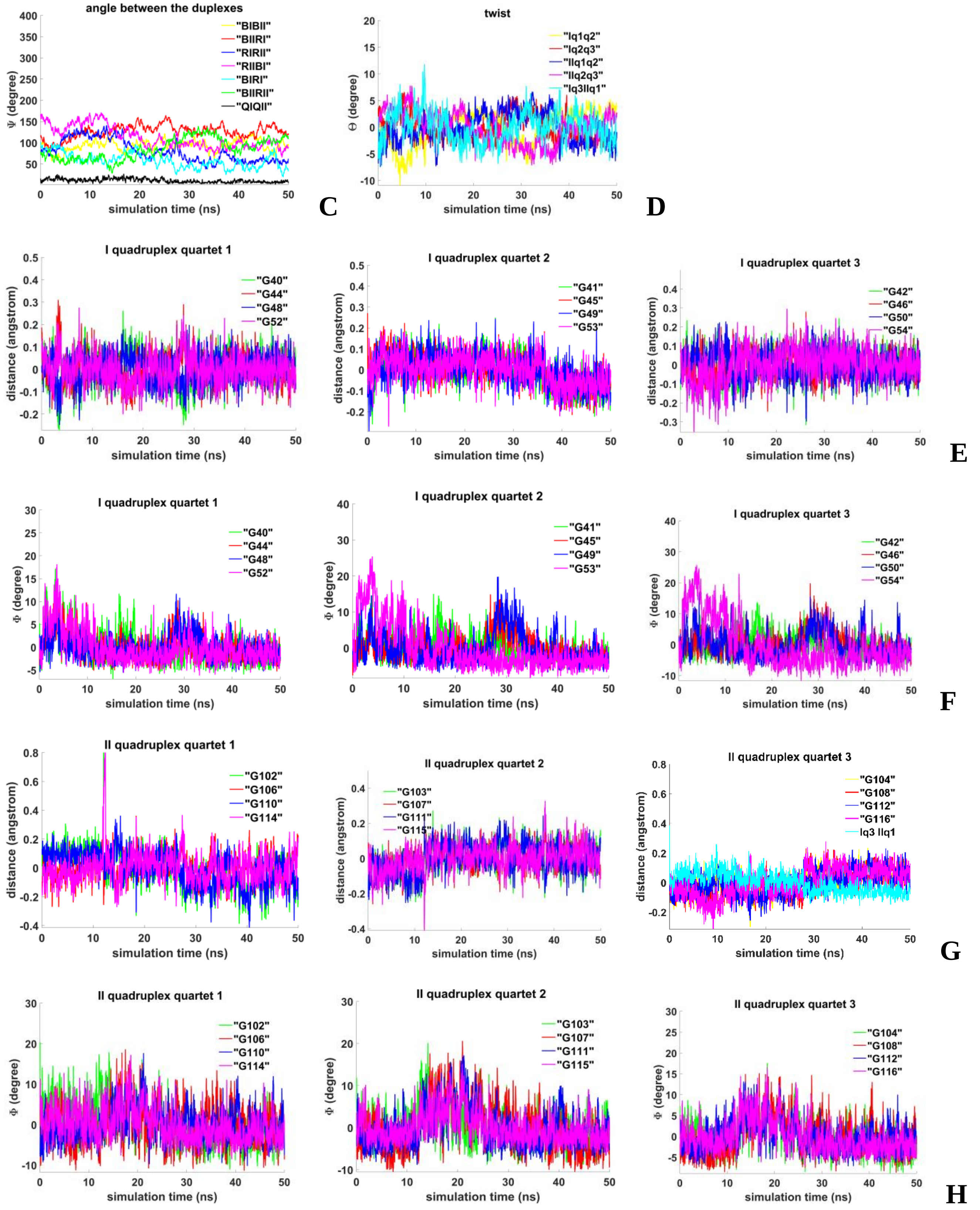


**Figure 3.2.B.S.2. Case 5: "Stacking and head-to-tail IM-dimer"**: **C** – angles between unmelted fragments of the duplexes, angle between axes passing through COMs of the boundary tetrads (**QIQII**); **D**– angles of rotation of the tetrads relative to each other; **E, G** - distances from COMs of the guanines' bases to COMs of their containing tetrads, distance between COMs of the boundary tetrads (**Iq3 IIq1**); **F, H** - angles between normals to the guanines' bases and vectors connecting COMs of the boundary tetrads.

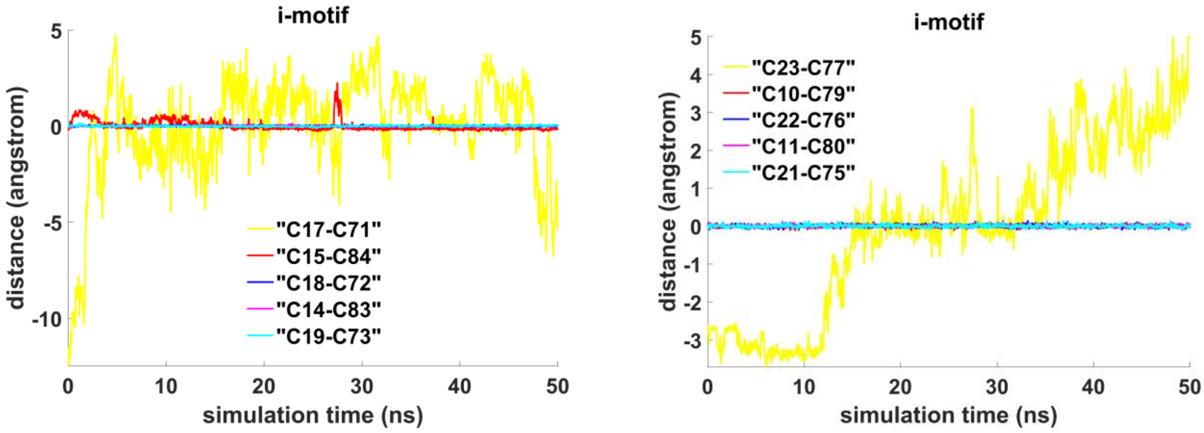


**I**

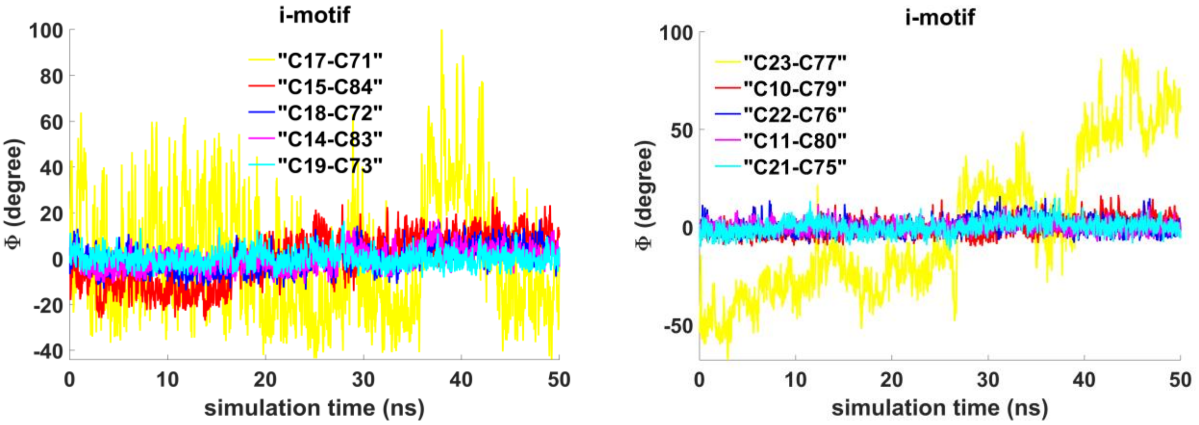


**J**

**Figure 3.2.B.S.3. Case 5: "Stacking and head-to-tail IM-dimer"**: **I -** distances between COMs of the cytosines' bases; **J -** angles between normals to the cytosines' bases.

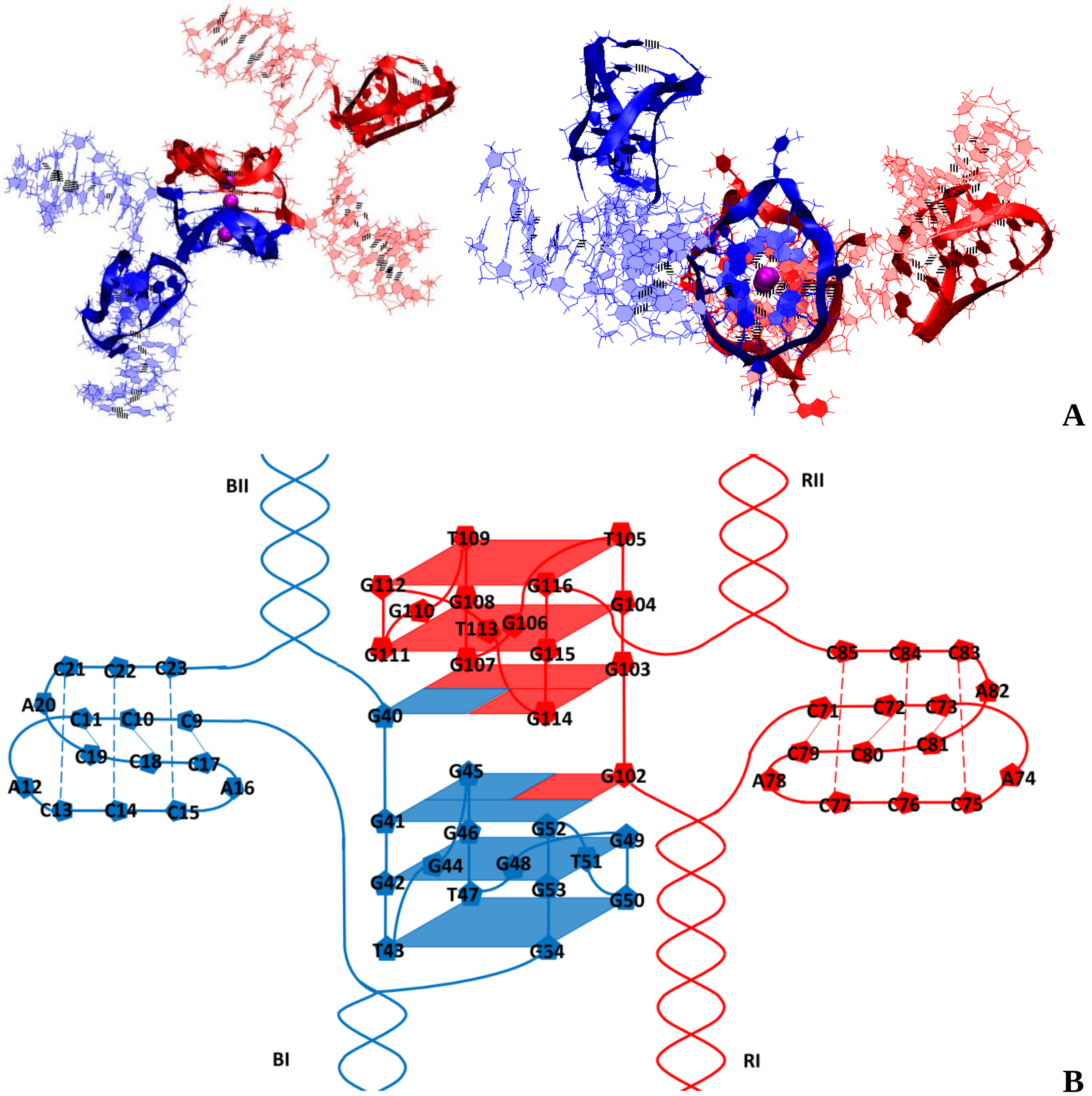


**Figure 3.2.C.S.1. Case 6: “Flip-flop interlock and two IM-monomers”**: **A** – conformation obtained at the last step of the MD trajectory (side and top view); **B** – the complex scheme.

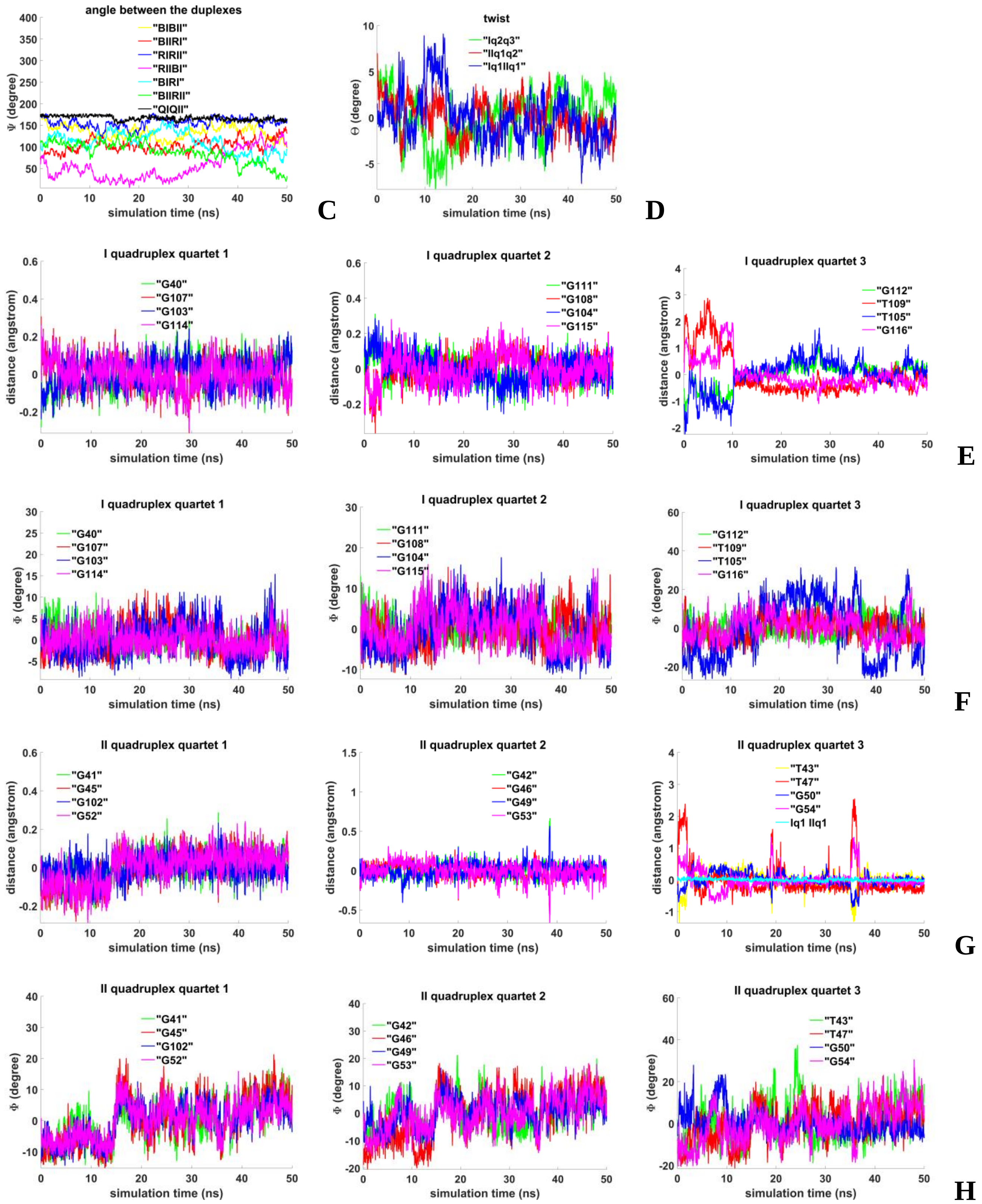


**Figure 3.2.C.S.2. Case 6: "Flip-flop interlock and two IM-monomers"**: **D -** angles between unmelted fragments of the duplexes, angle between axes passing through COMs of the boundary tetrads (**QIQII**); **E**– angles of rotation of the tetrads relative to each other; **F, H** - distances from COMs of the guanines' bases to COMs of their containing tetrads, distance between COMs of the boundary tetrads (**Iq1 IIq1**); **G, I** - angles between normals to the guanines' bases and vectors connecting COMs of the boundary tetrads.

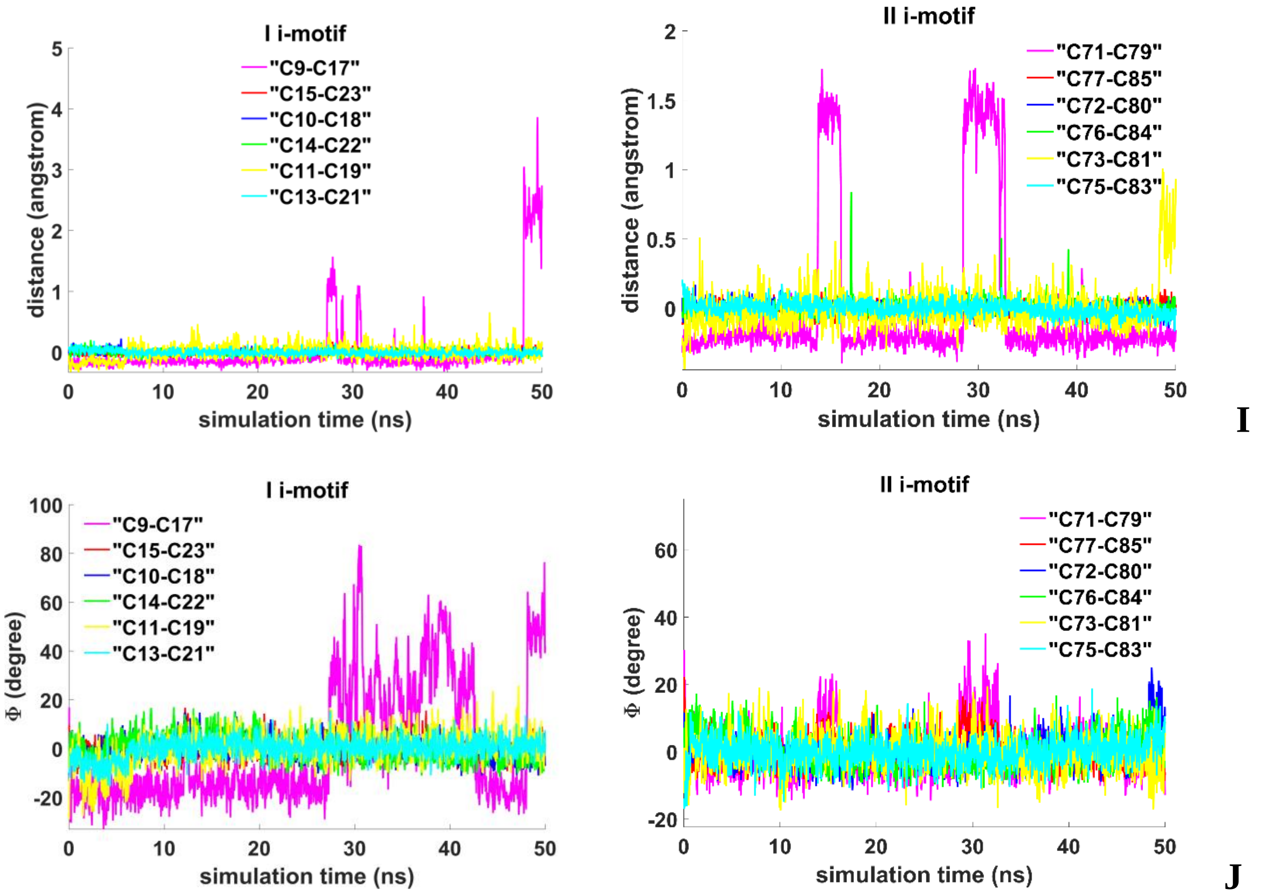


**Figure 3.2.C.S.3. Case 6: "Flip-flop interlock and two IM-monomers"**: **I -** distances between COMs of the cytosines' bases; **J** - angles between normals to the cytosines' bases.

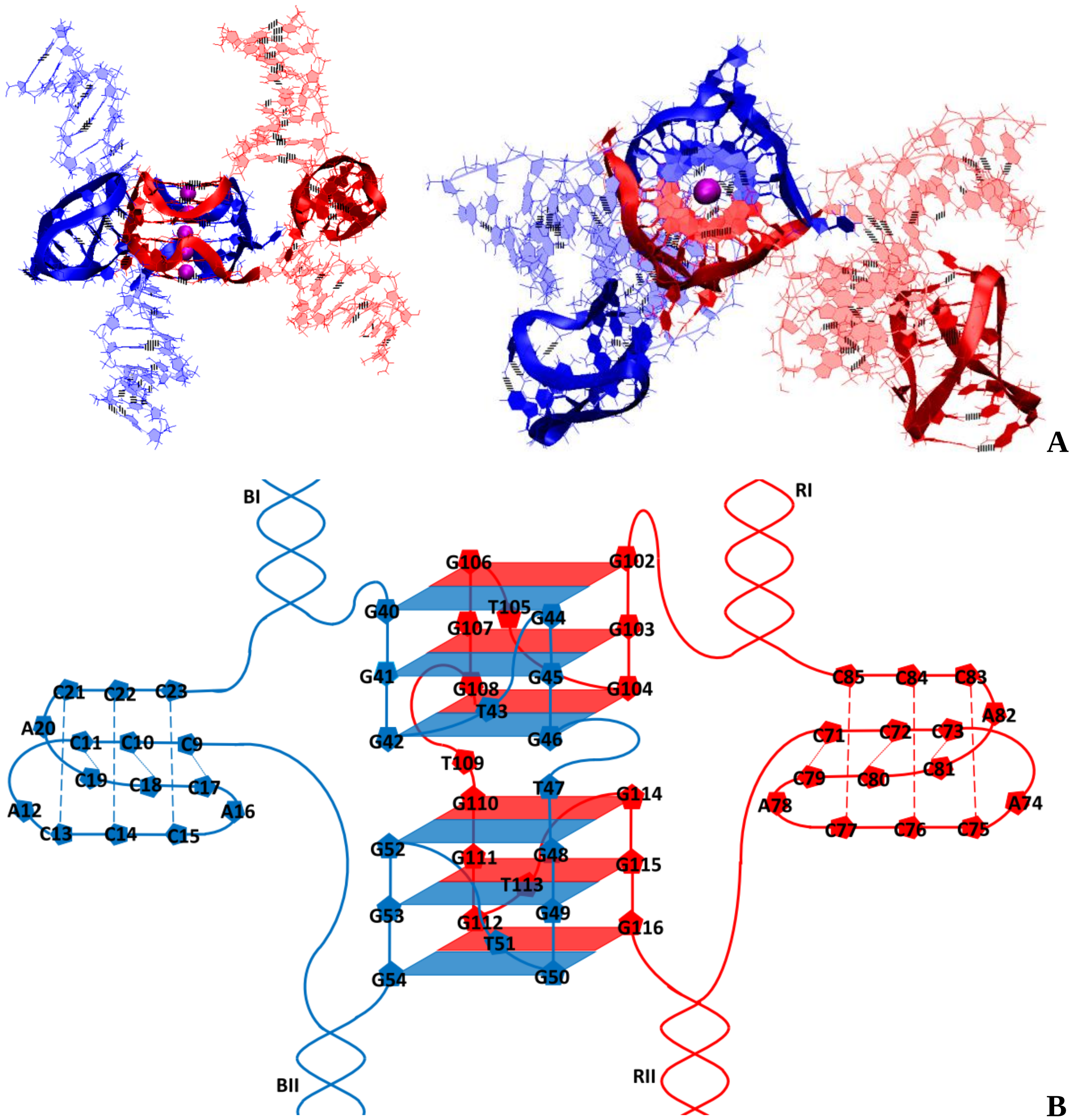


**Figure 3.3.A.S.1. Case 7: "Stacking of right and left handed parallel G4-dimers, and two IM-monomers"**: **A** – conformation obtained at the last step of the MD trajectory (side and top view); **B** – the complex scheme.

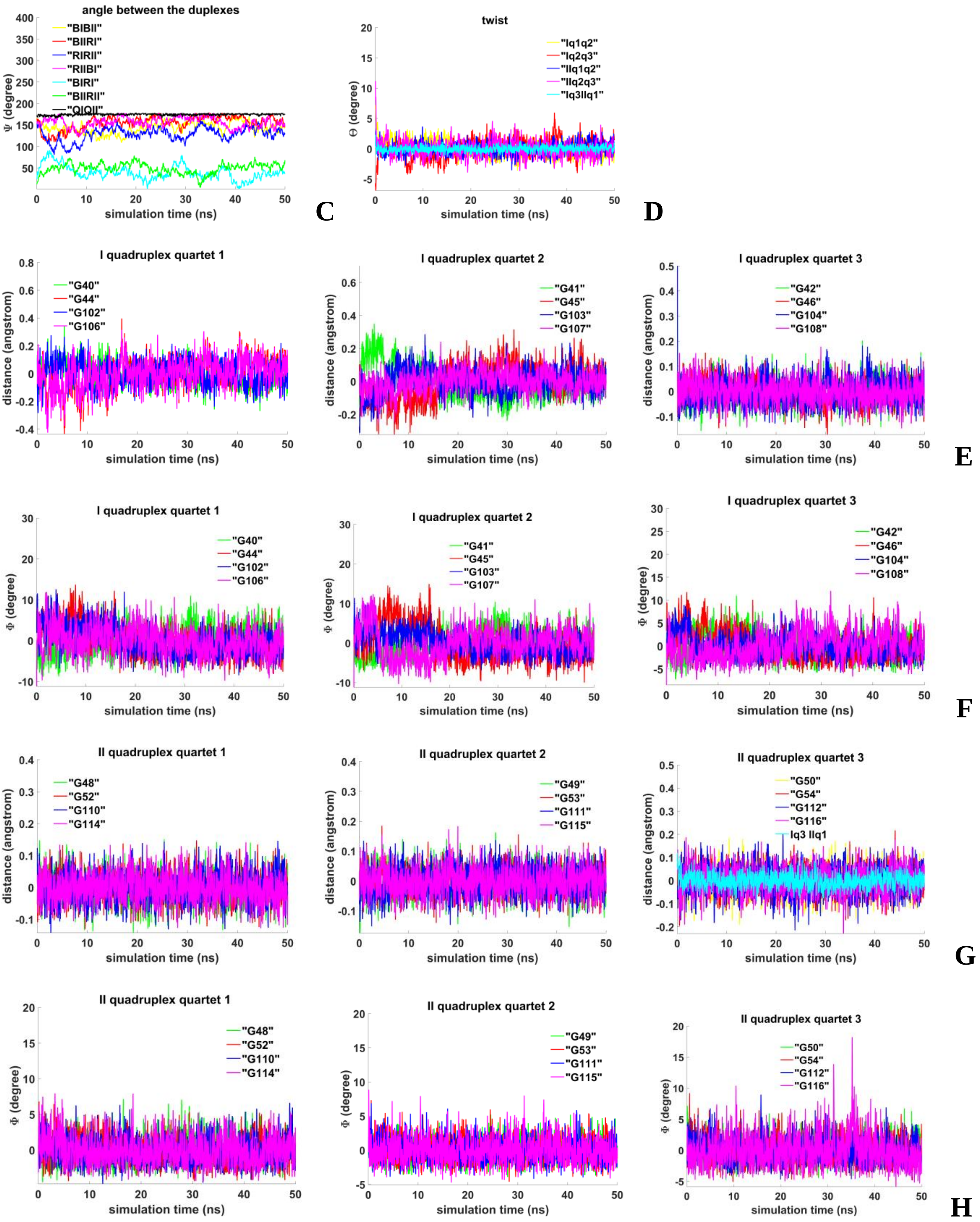


**Figure 3.3.A.S.2. Case 7: "Stacking of right and left handed parallel G4-dimers, and two IM-monomers"**: **C** – angles between unmelted fragments of the duplexes, angle between axes passing through COMs of the boundary tetrads (**QIQII**); **D**– angles of rotation of the tetrads relative to each other; **E, G** - distances from COMs of the guanines' bases to COMs of their containing tetrads, distance between COMs of the boundary tetrads (**Iq3 IIq1**); **F, H** - angles between normals to the guanines' bases and vectors connecting COMs of the boundary tetrads.

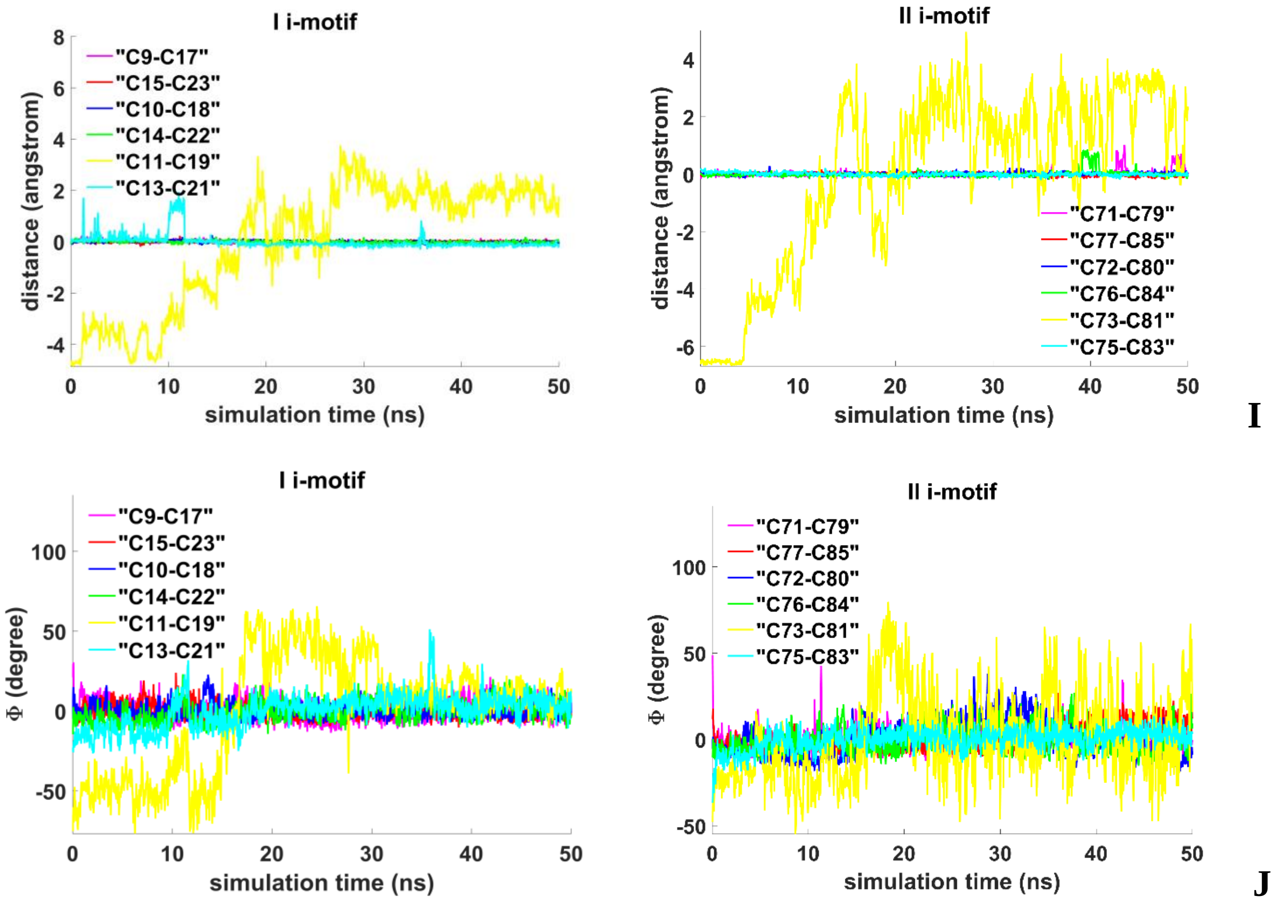


**Figure 3.3.A.S.3. Case 7: “Stacking of right and left handed parallel G4-dimers, and two IM-monomers”**: **I -** distances between COMs of the cytosines’ bases; **J** - angles between normals to the cytosines’ bases.

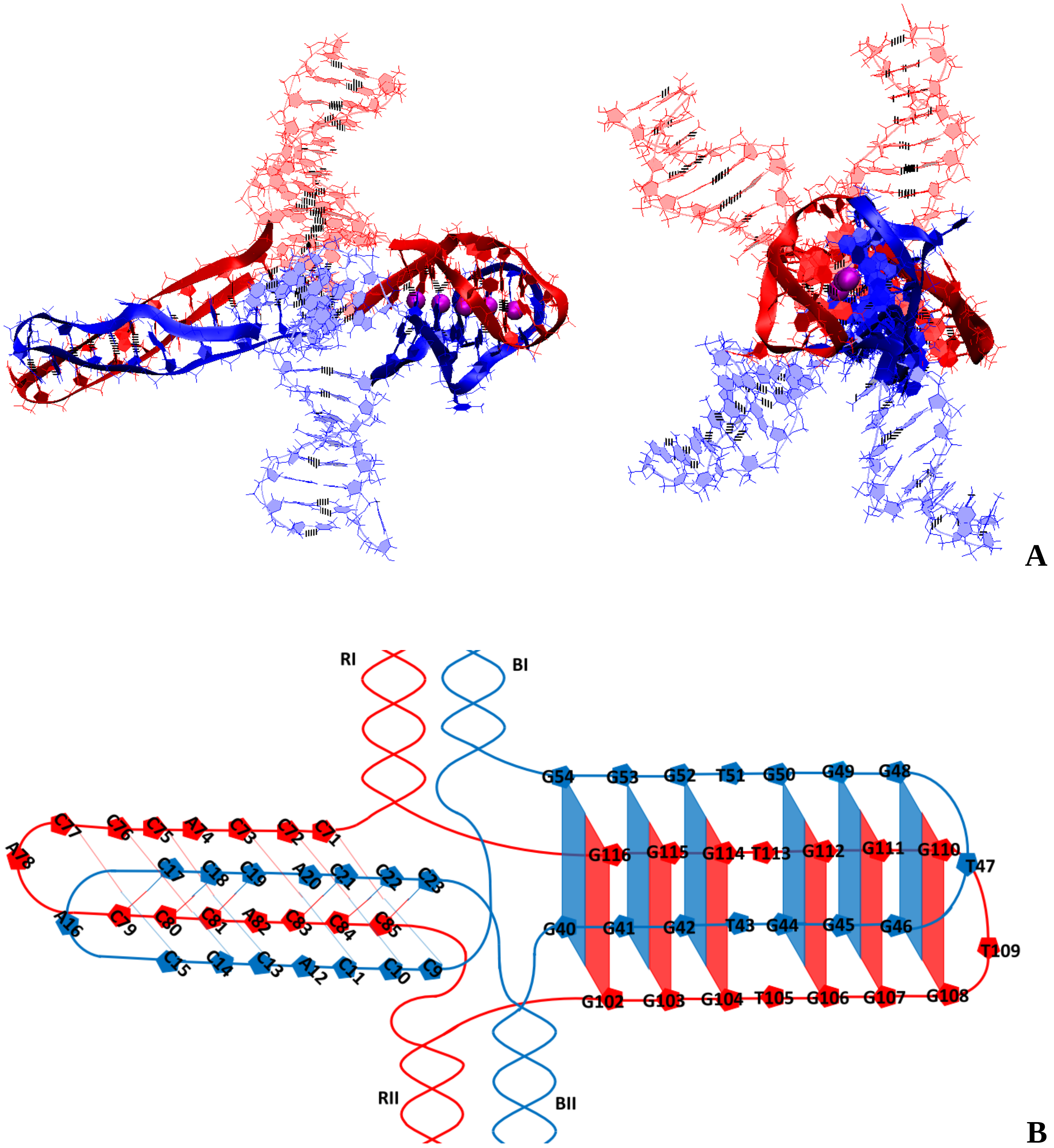


**Figure 3.3.B.S.1. Case 8: "Antiparallel G4-dimer and head-to-head IM-dimer"**: **A** – conformation obtained at the last step of the MD trajectory (side and top view); **B** – the complex scheme.

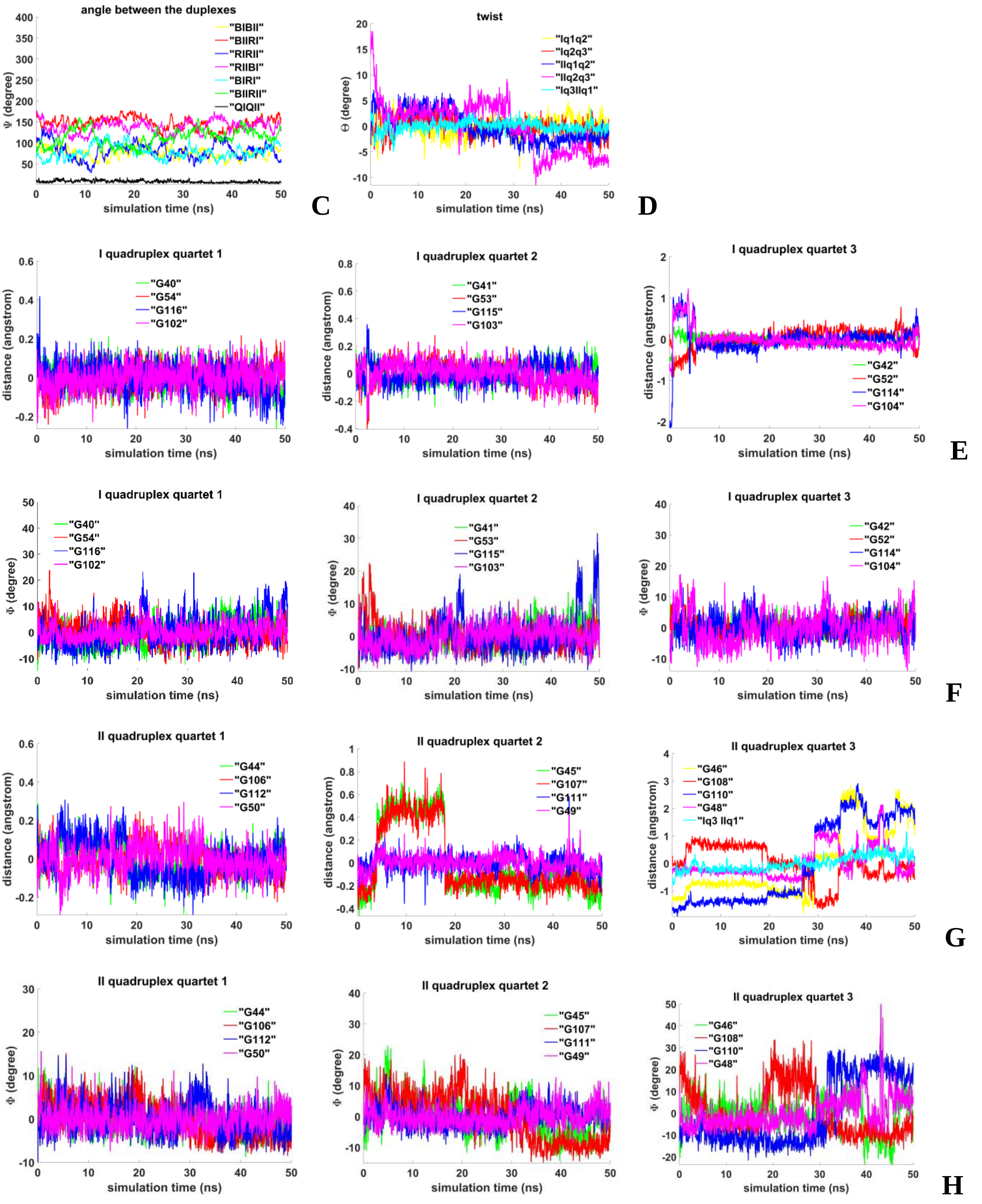


**Figure 3.3.B.S.2. Case 8: "Antiparallel G4-dimer and head-to-head IM-dimer: C** – angles between unmelted fragments of the duplexes, angle between axes passing through COMs of the boundary tetrads (**QIQII**); **D**– angles of rotation of the tetrads relative to each other; **E, G** - distances from COMs of the guanines' bases to COMs of their containing tetrads, distance between COMs of the boundary tetrads (**Iq3 IIq1**); **F, H** - angles between normals to the guanines' bases and vectors connecting COMs of the boundary tetrads.

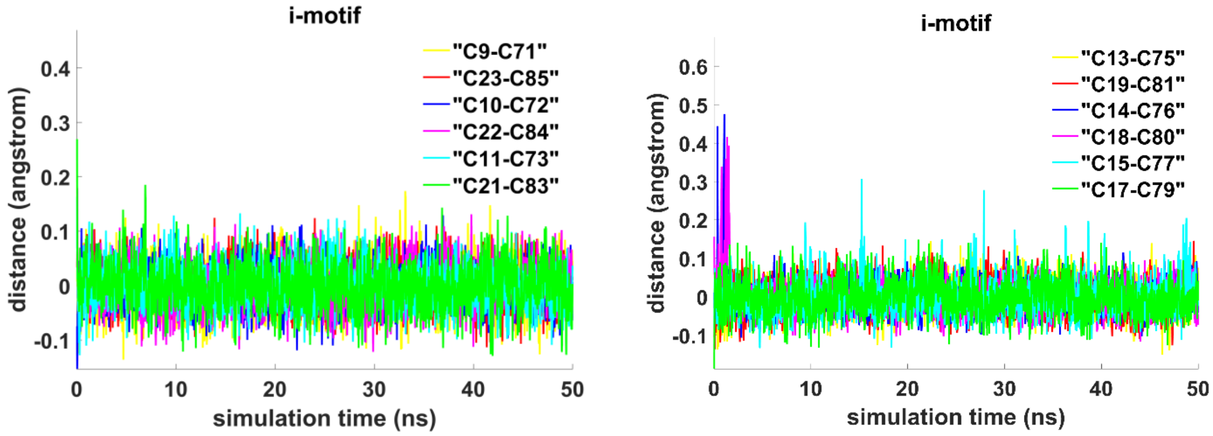


**I**

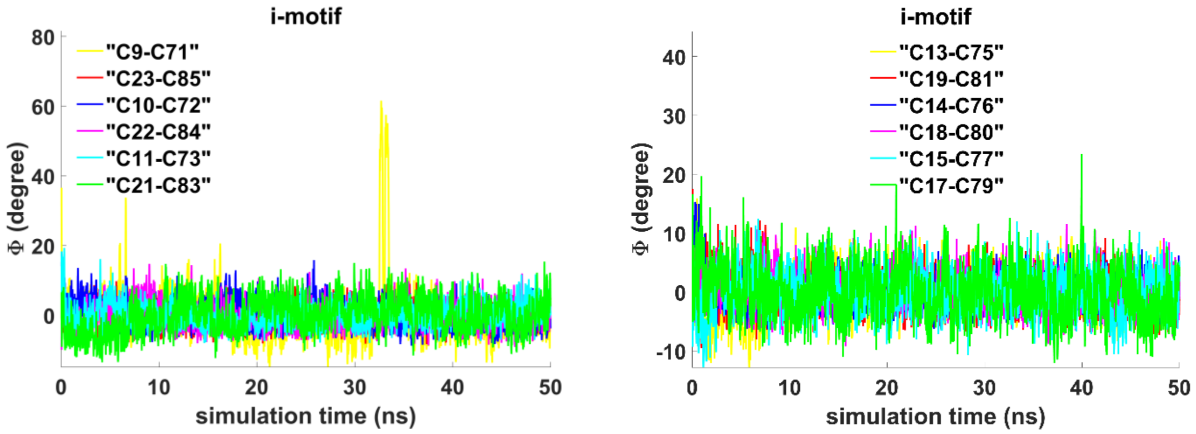


**J**

**Figure 3.3.B.S.3. Case 8: “Antiparallel G4-dimer and head-to-head IM-dimer”**: **I -** distances between COMs of the cytosines’ bases; **J** - angles between normals to the cytosines’ bases.

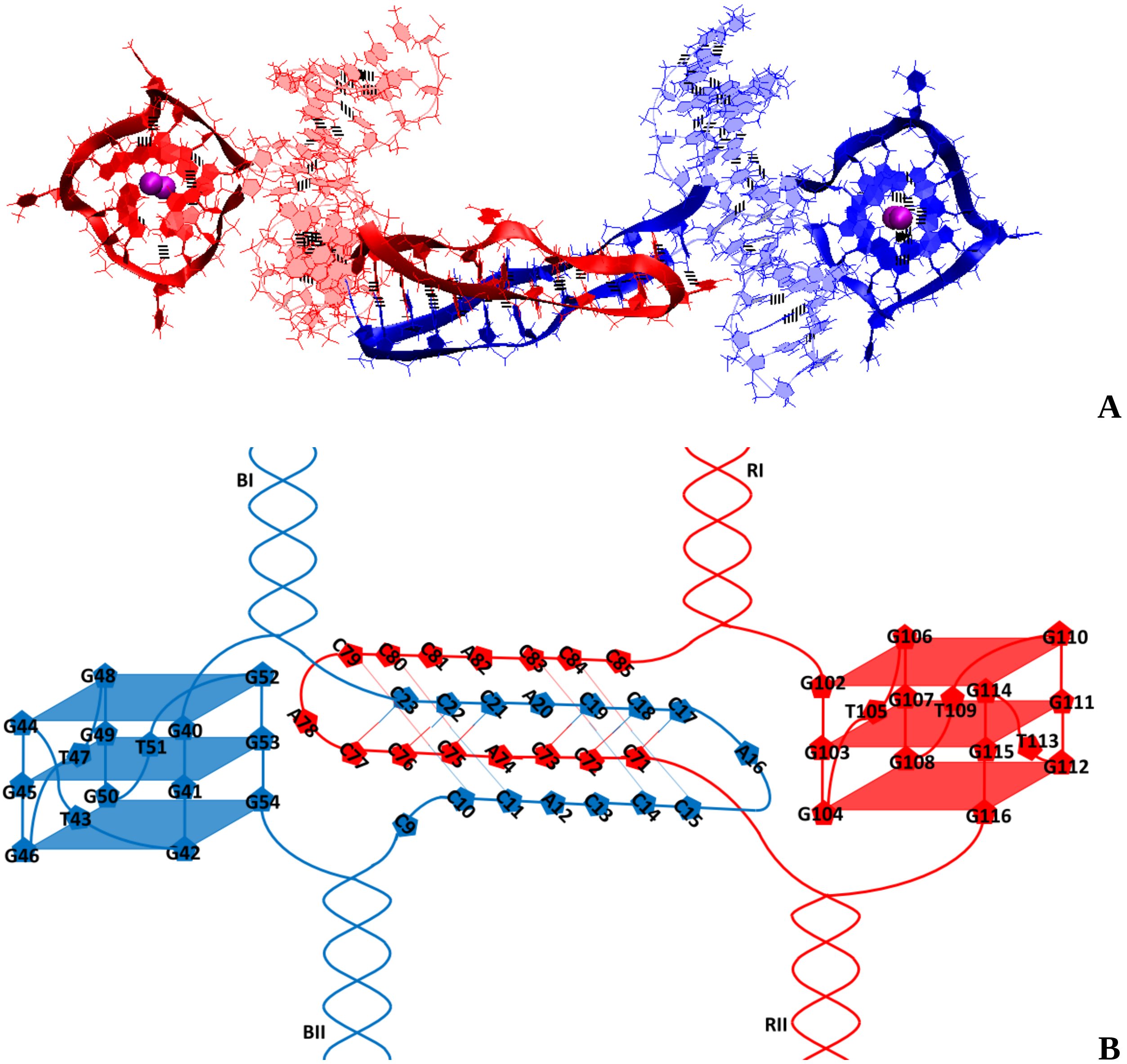


**Figure 3.3.C.S.1. Case 9: "Head-to-tail IM-dimer between two parallel G4-monomers"**: **A** – conformation obtained at the last step of the MD trajectory (side and top view); **B** – the complex scheme.

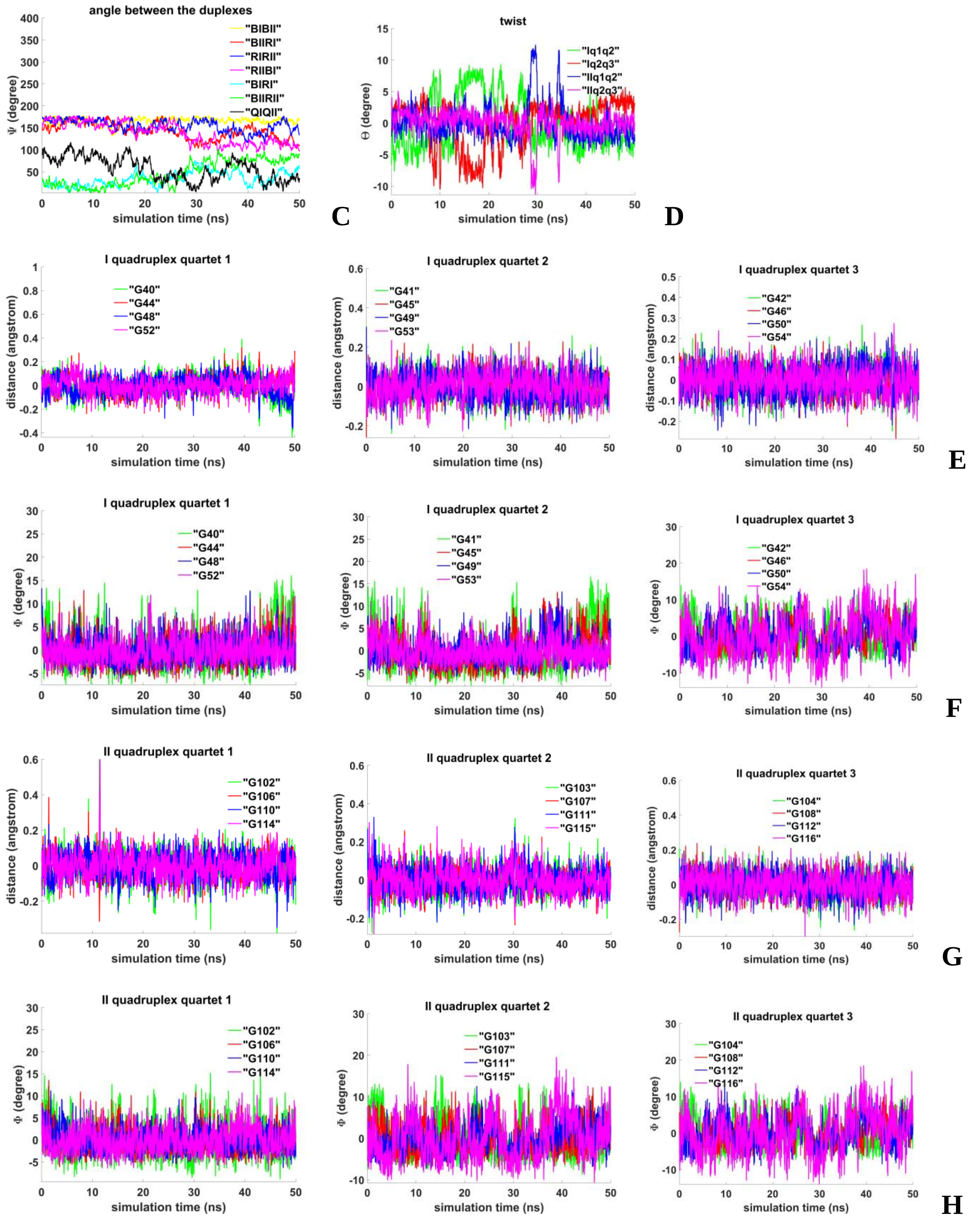


**Figure 3.3.C.S.2. Case 9: "Head-to-tail IM-dimer between two parallel G4-monomers"**: **C** – angles between unmelted fragments of the duplexes, angle between axes passing through COMs of the boundary tetrads (**QIQII**); **D** – angles of rotation of the tetrads relative to each other; **E, G** - distances from COMs of the guanines' bases to COMs of their containing tetrads; **F, H** - angles between normals to the guanines' bases and vectors connecting COMs of the boundary tetrads.

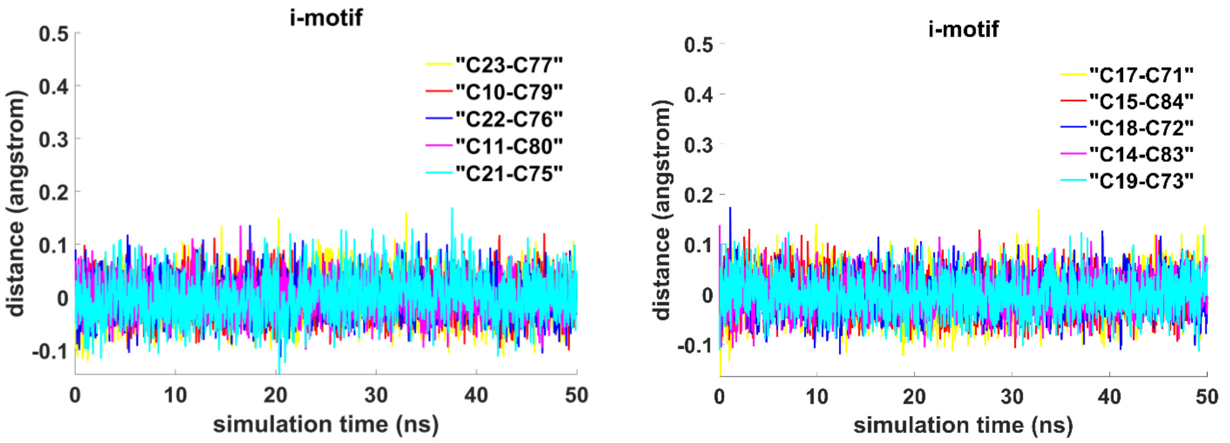


I

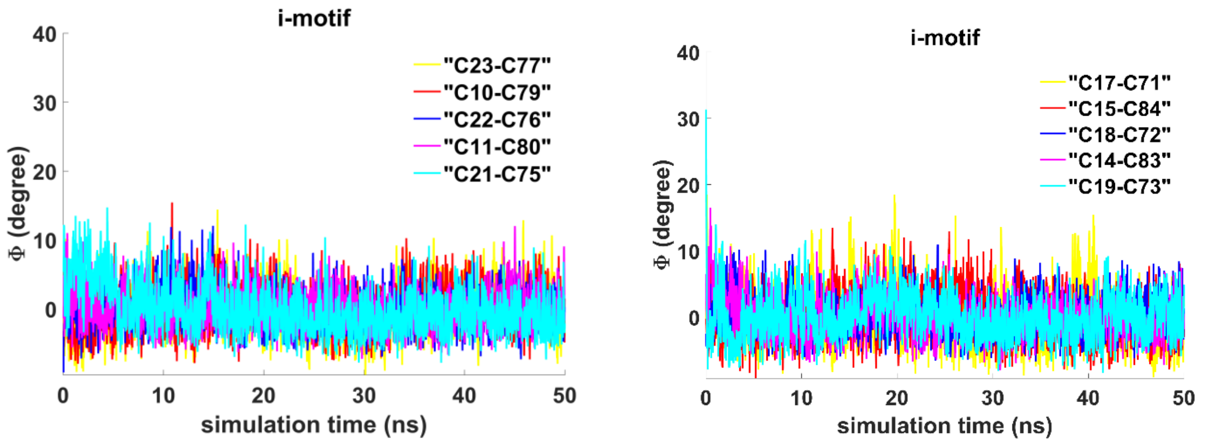


J

**Figure 3.3.C.S.3. Case 9: "Head-to-tail IM-dimer between two parallel G4-monomers"**: **I** - distances between COMs of the cytosines' bases; **J** - angles between normals to the cytosines' bases.

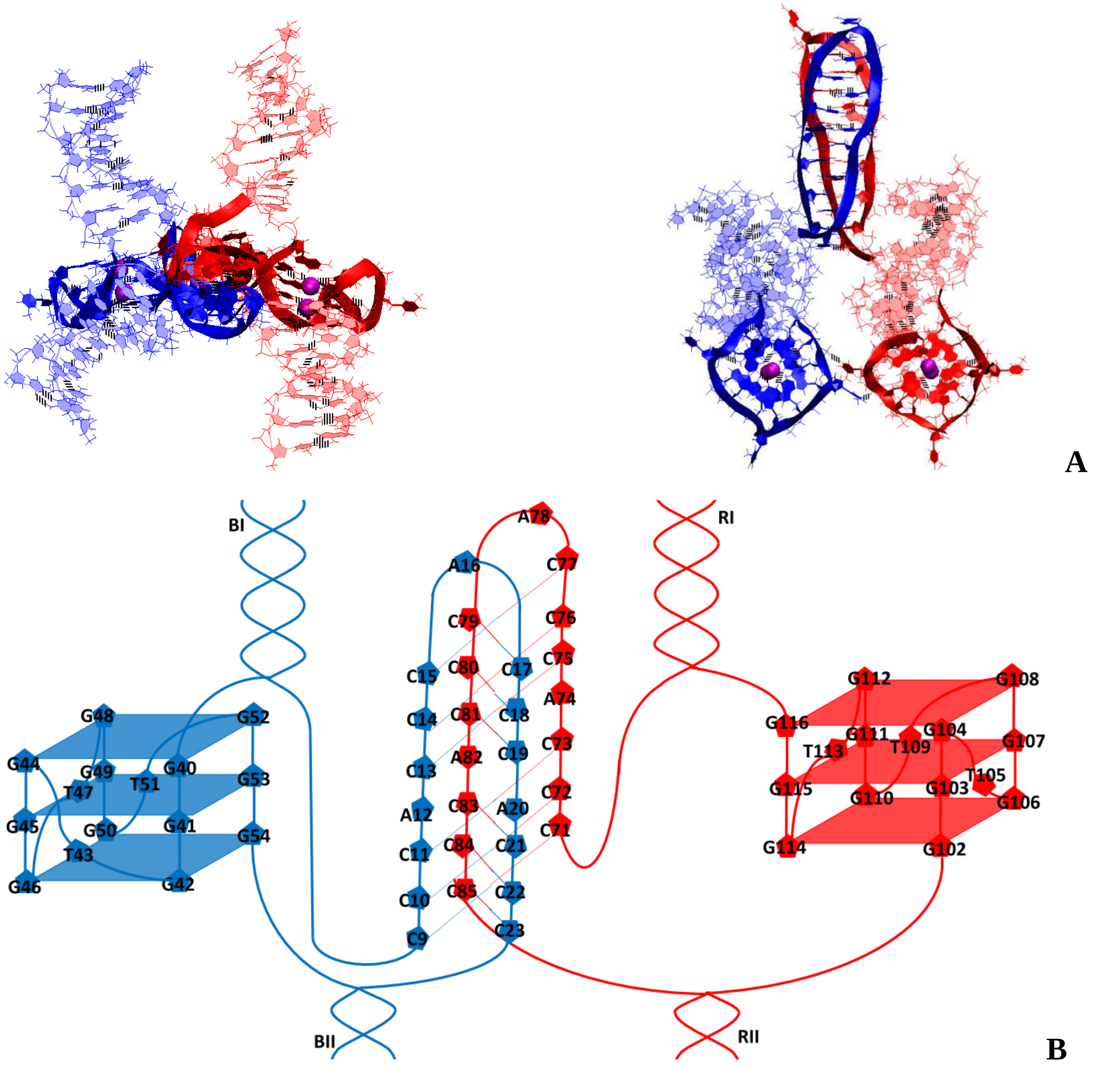


**Figure 3.4.A.S.1. Case 10: “Head-to-head IM-dimer between two parallel G4-monomers”**: **A** – conformation obtained at the last step of the MD trajectory (side and top view); **B** – the complex scheme.

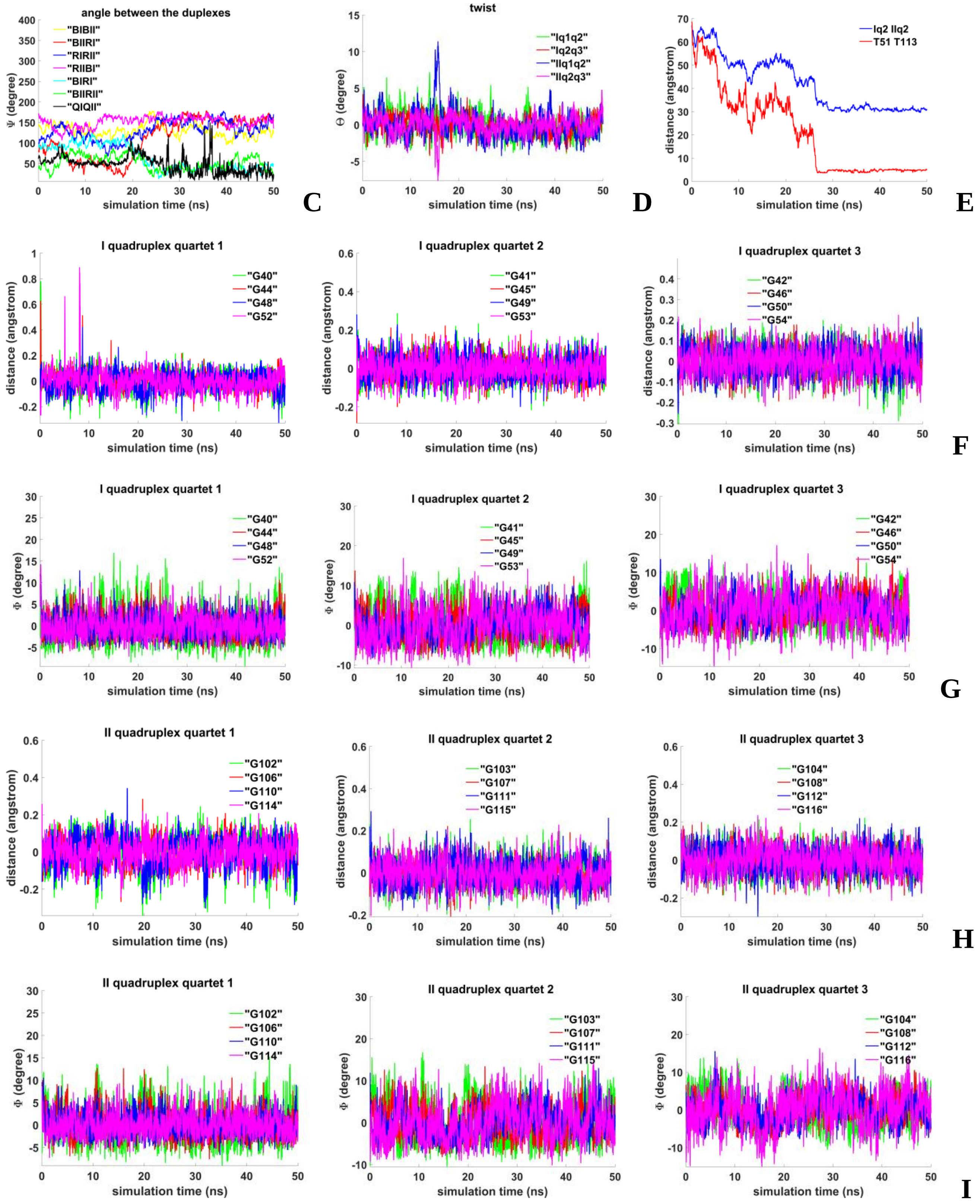


**Figure 3.4.A.S.2. Case 10: "Head-to-head IM-dimer between two parallel G4-monomers"**: **C** – angles between unmelted fragments of the duplexes, angle between axes passing through COMs of the boundary tetrads (**QIQII**); **D** – angles of rotation of the tetrads relative to each other; **E** – evolution of values of distances between COMs of the G4s and distances between COMs of bases of **T51** and **T113**; **F, H** - distances from COMs of the guanines' bases to COMs of their containing tetrad;, **G,I** - angles between normals to the guanines' bases and vectors connecting COMs of the boundary tetrads.

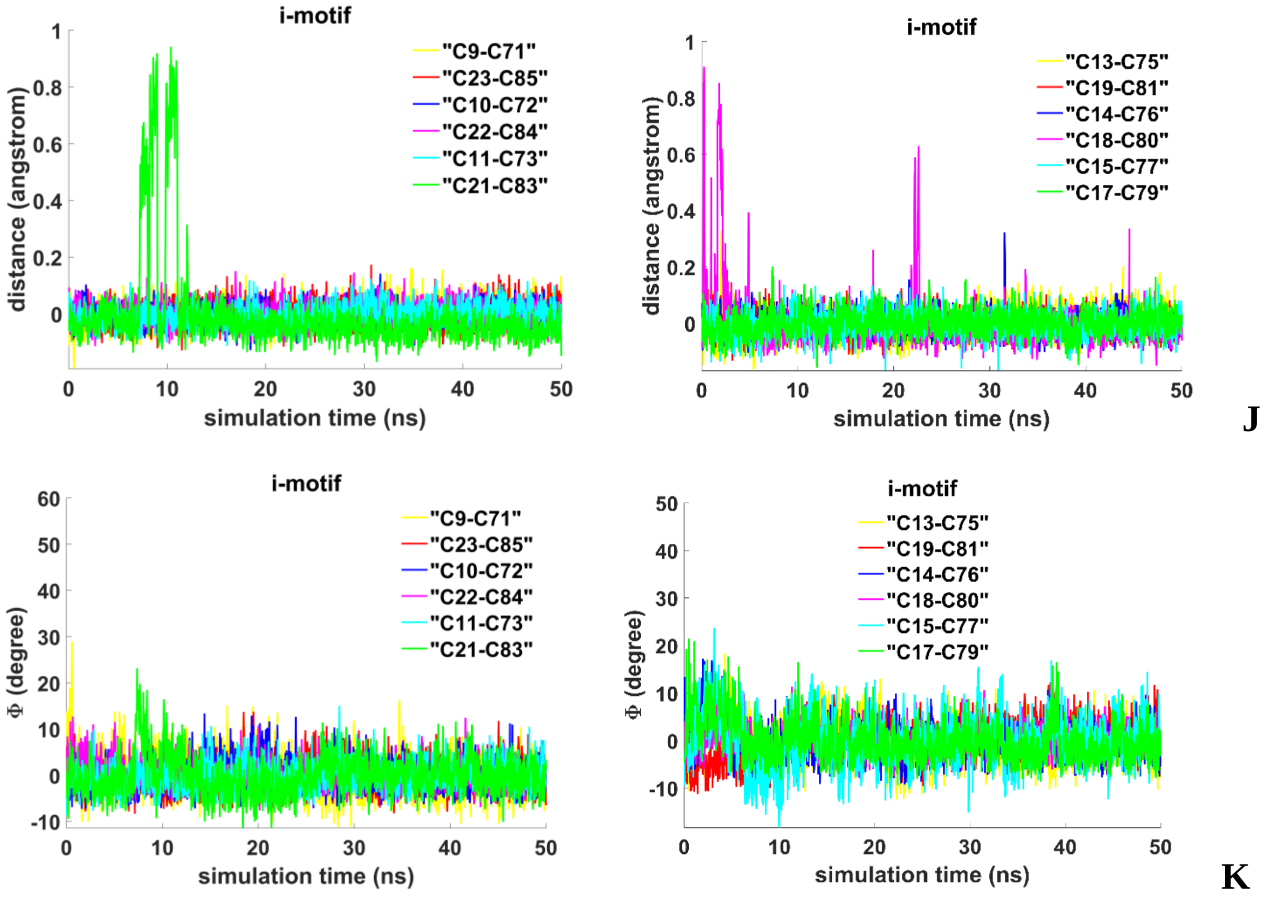


**Figure 3.4.A.S.3. Case 10: "Head-to-head IM-dimer between two parallel G4-monomers"**: **J** - distances between COMs of the cytosines' bases; **K** - angles between normals to the cytosines' bases.

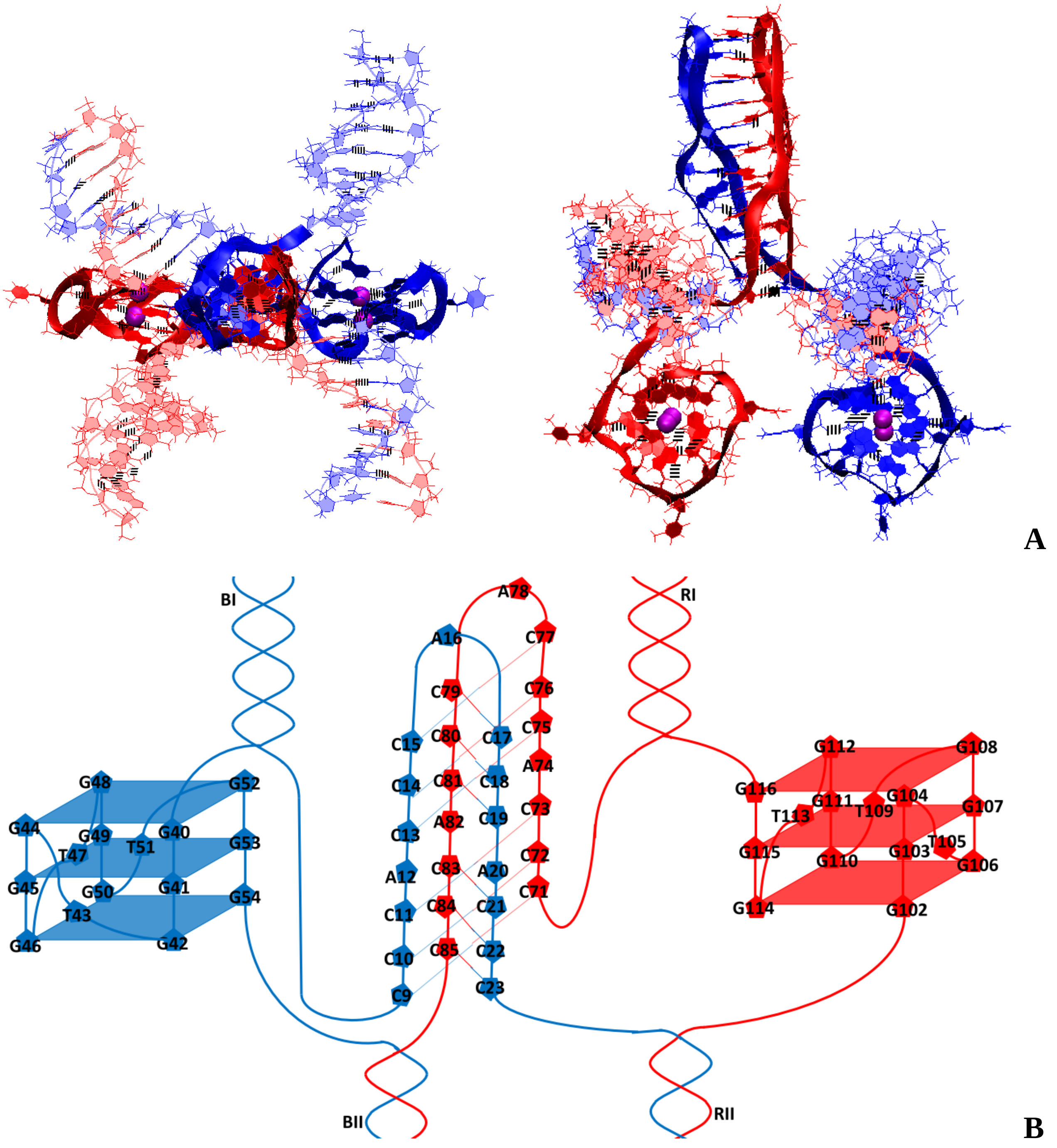


**Figure 3.4.B.S.1. Case 11: "Head-to-head IM-dimer between two parallel G4-monomers in case of the strands exchange"**: **A** – conformation obtained at the last step of the MD trajectory (side and top view); **B** – the complex scheme.

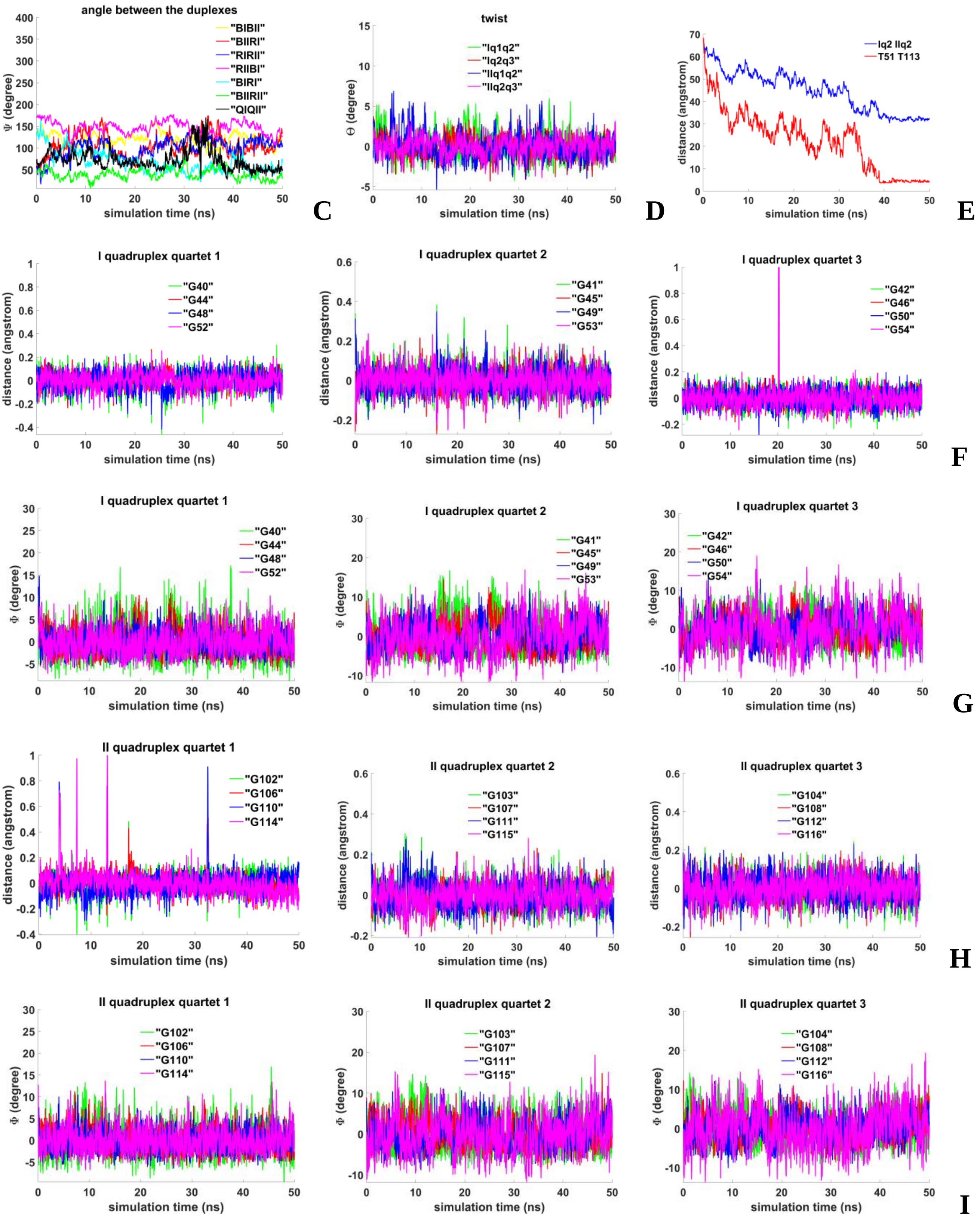


**Figure 3.4.B.S.2. Case 11: "Head-to-head IM-dimer between two parallel G4-monomers in case of the strands exchange"**: **C** – angles between unmelted fragments of the duplexes, angle between axes passing through COMs of the boundary tetrads (**QIQII**); **D** – angles of rotation of the tetrads relative to each other; **E** – evolution of values of distances between COMs of the G4s and distances between COMs of bases of **T51** and **T113**; **F, H** - distances from COMs of the guanines' bases to COMs of their containing tetrad; **G, I** - angles between normals to the guanines' bases and vectors connecting COMs of the boundary tetrads.

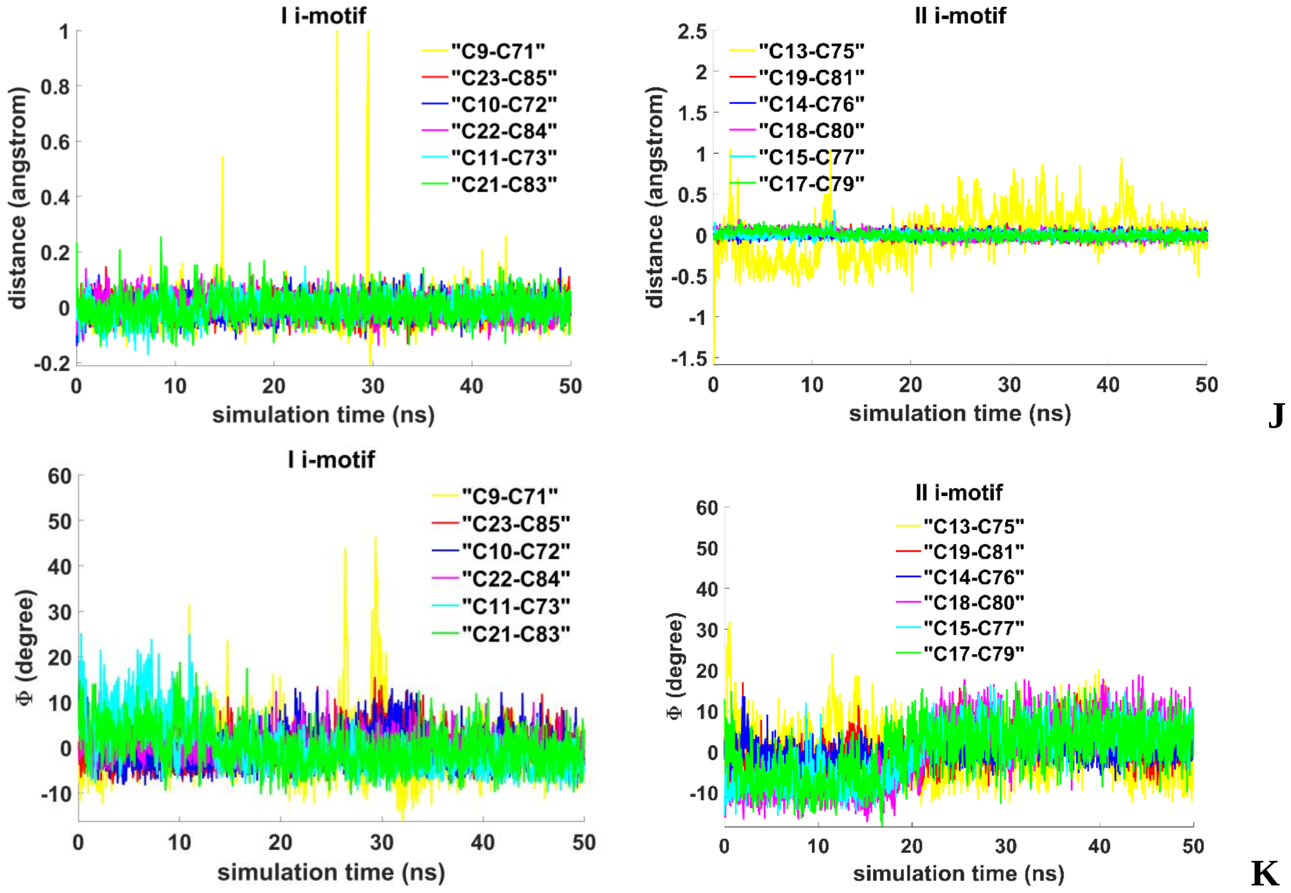


**Figure 3.4.B.S.3. Case 11: "Head-to-head IM-dimer between two parallel G4-monomers in case of the strands exchange"**: **J** - distances between COMs of the cytosines' bases; **K** - angles between normals to the cytosines' bases.

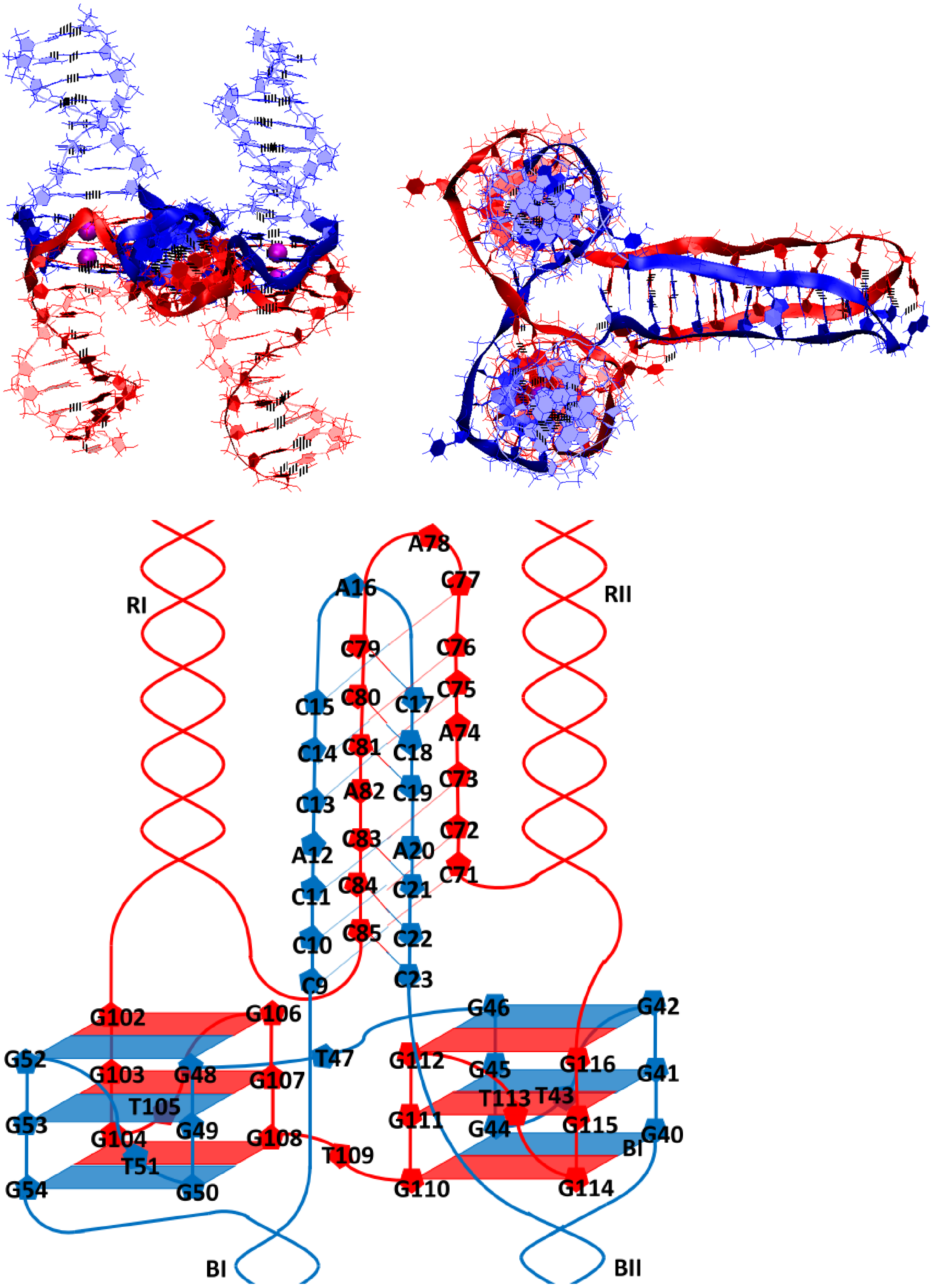


A

B

**Figure 3.4.C.S.1. Case 12: "Two parallel G4-dimer in the same plane and head-to-head IM-dimer between two unmelted fragments of duplexes with mutual girth of the strands"**: **A** – conformation obtained at the last step of the MD trajectory (side and top view); **B** – the complex scheme.

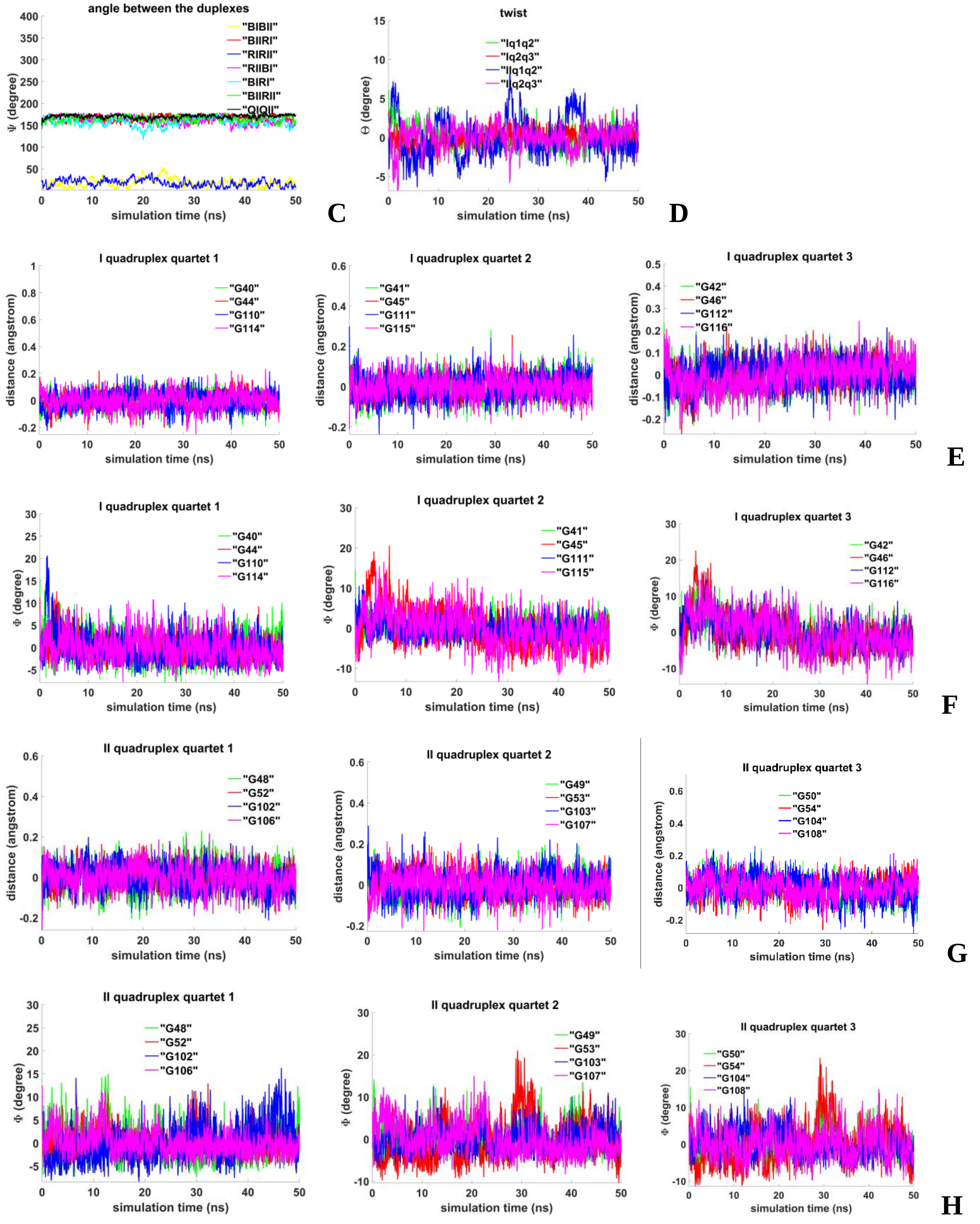


**Figure 3.4.C.S.2. Case 12: "Two parallel G4-dimer in the same plane and head-to-head IM-dimer between two unmelted fragments of duplexes with mutual girth of the strands"**: **C** – angles between unmelted fragments of the duplexes, angle between axes passing through COMs of the boundary tetrads (**QIQII**); **D** – angles of rotation of the tetrads relative to each other; **E, G** - distances from COMs of the guanines' bases to COMs of their containing tetrad; **F, H** - angles between normals to the guanines' bases and vectors connecting COMs of the boundary tetrads.

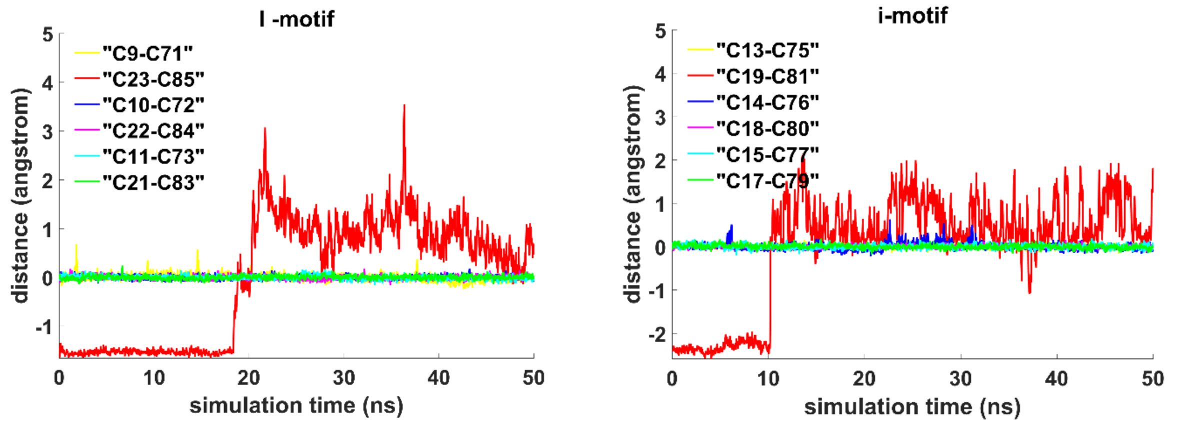


I

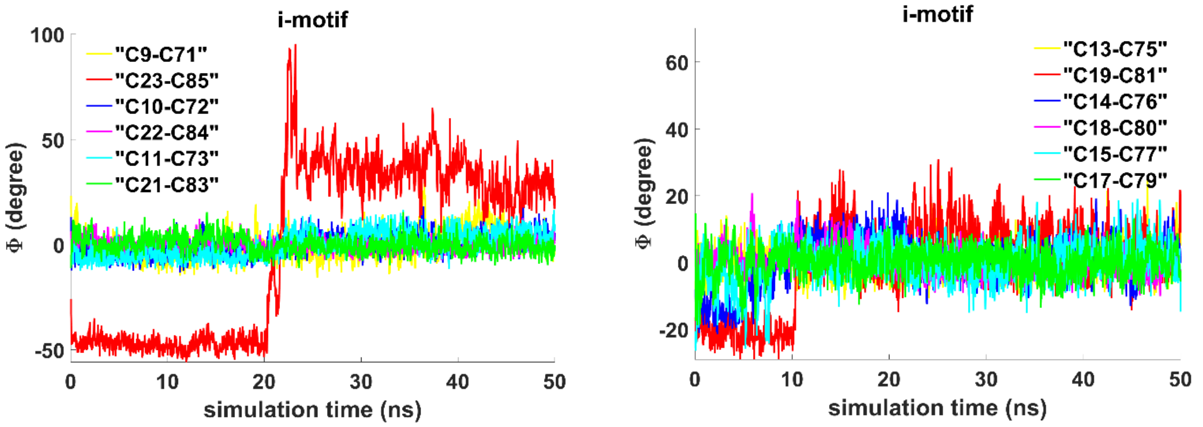


J

**Figure 3.4.C.S.3. Case 12: “Two parallel G4-dimer in the same plane and head-to-head IM-dimer between two unmelted fragments of duplexes with mutual girth of the strands”**: **I** - distances between COMs of the cytosines’ bases; **J** - angles between normals to the cytosines’ bases.

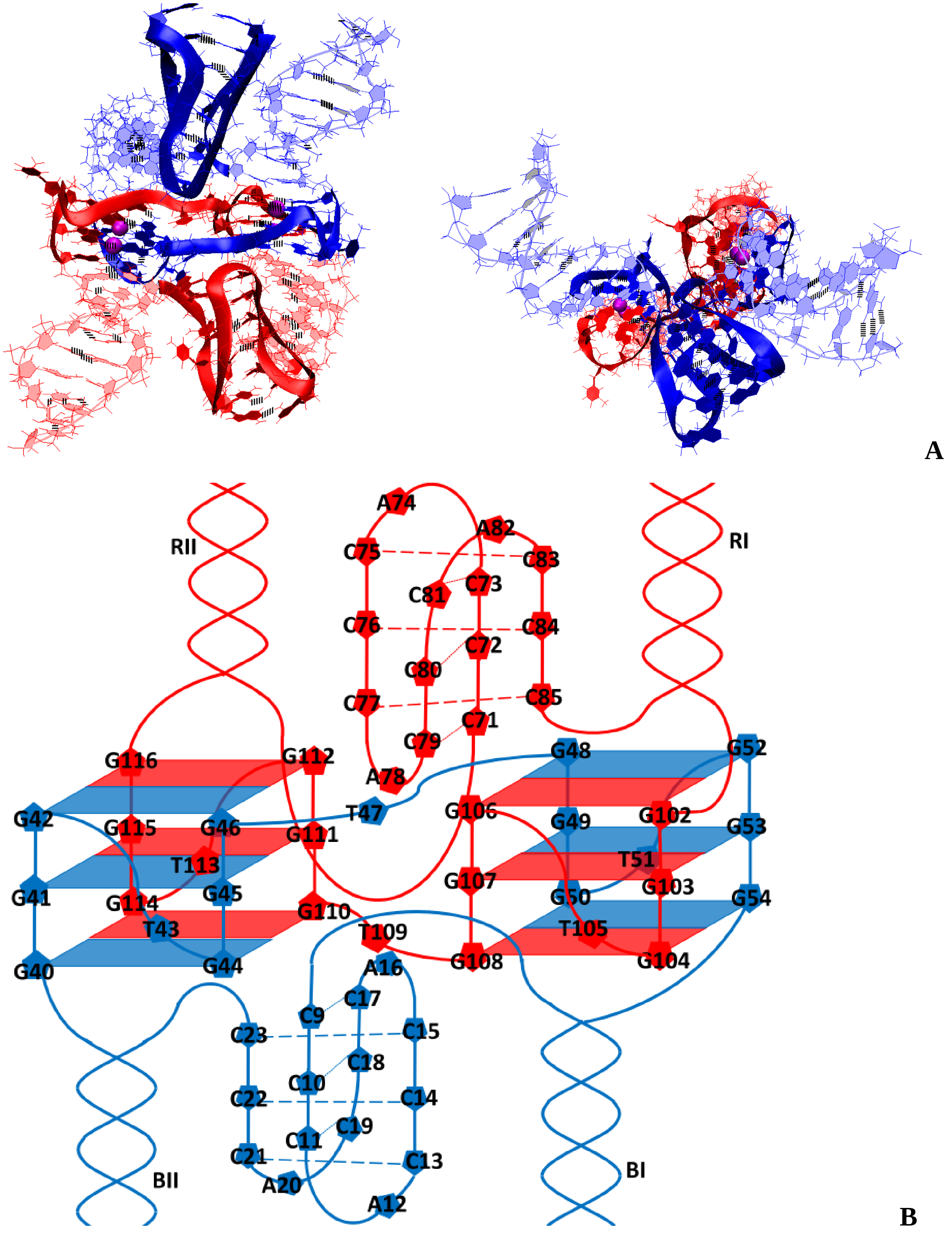


**Figure 3.5.A.S.1. Case 13: "Two parallel G4-dimers in the same plane and two IM-monomers clamped unmelted fragments of duplexes with mutual girth of the strands": A** – conformation obtained at the last step of the MD trajectory (side and top view); **B** – the complex scheme.

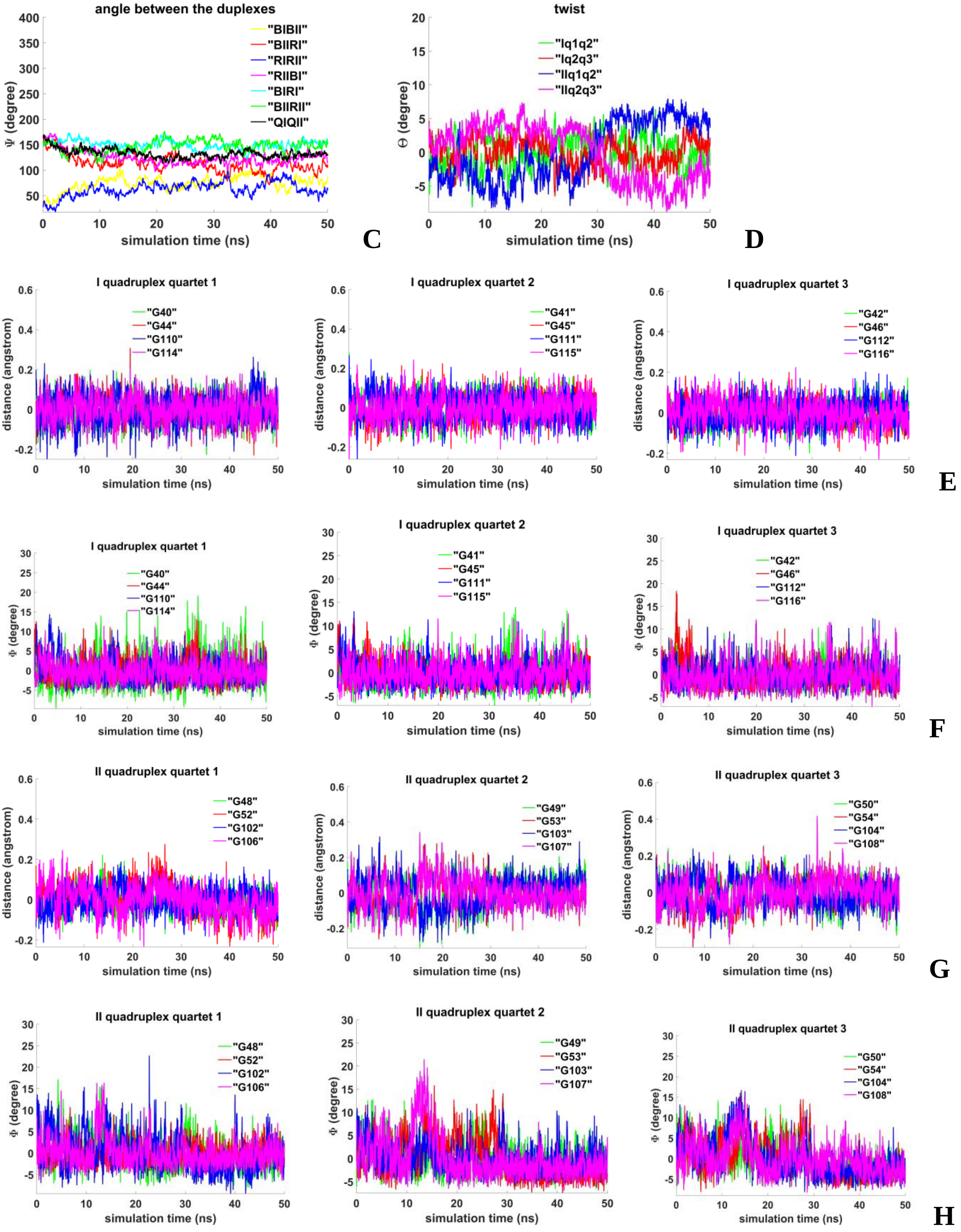


**Figure 3.5.A.S.2. Case 13: "Two parallel G4-dimers in the same plane and two IM-monomers clamped unmelted fragments of duplexes with mutual girth of the strands": C** – angles between unmelted fragments of the duplexes, angle between axes passing through COMs of the boundary tetrads (**QIQII**); **D**– angles of rotation of the tetrads relative to each other; **E, G** - distances from COMs of the guanines' bases to COMs of their containing tetrads; **F, H** - angles between normals to the guanines' bases and vectors connecting COMs of the boundary tetrads.

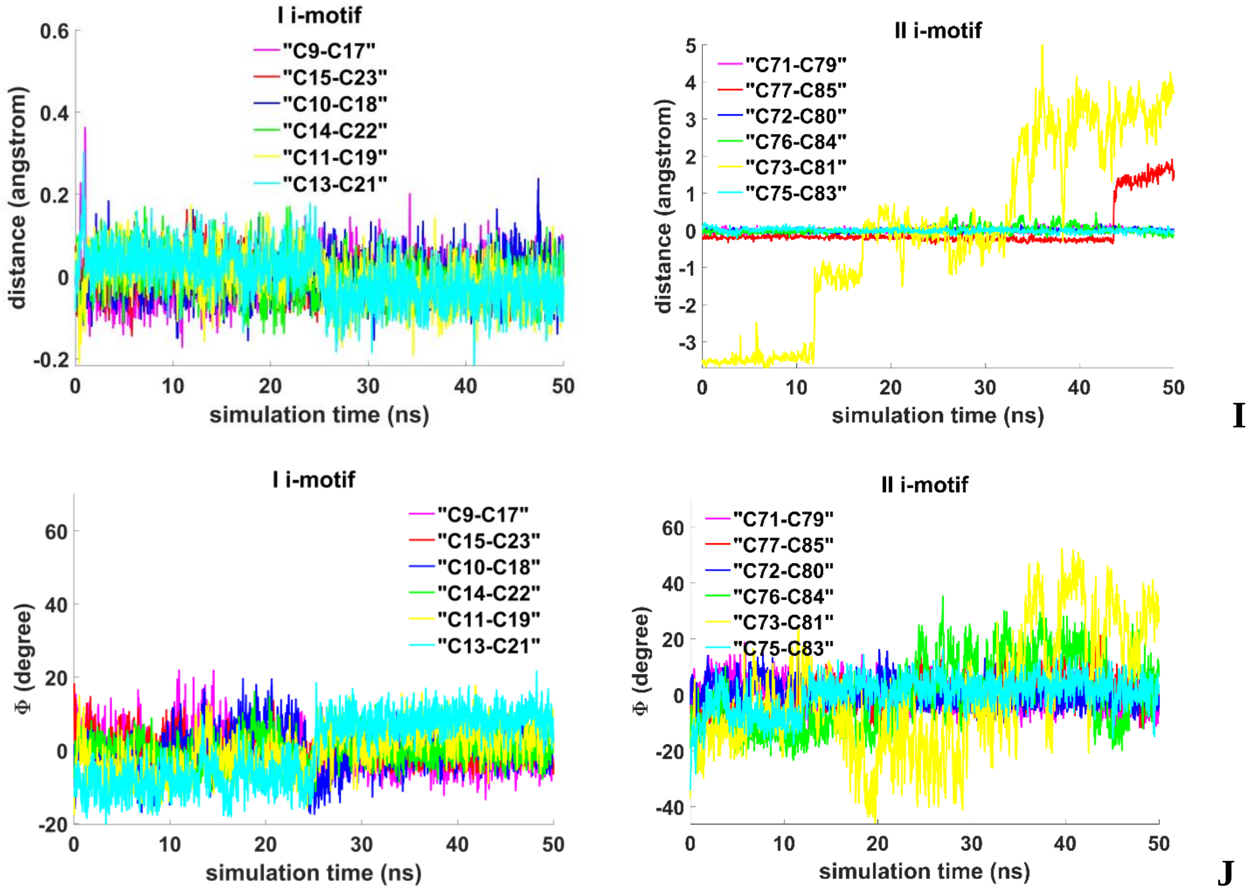


**Figure 3.5.A.S.3. Case 13: "Two parallel G4-dimers in the same plane and two IM-monomers clamped unmelted fragments of duplexes with mutual girth of the strands": I** - distances between COMs of the cytosines' bases; **J** - angles between normals to the cytosines' bases.

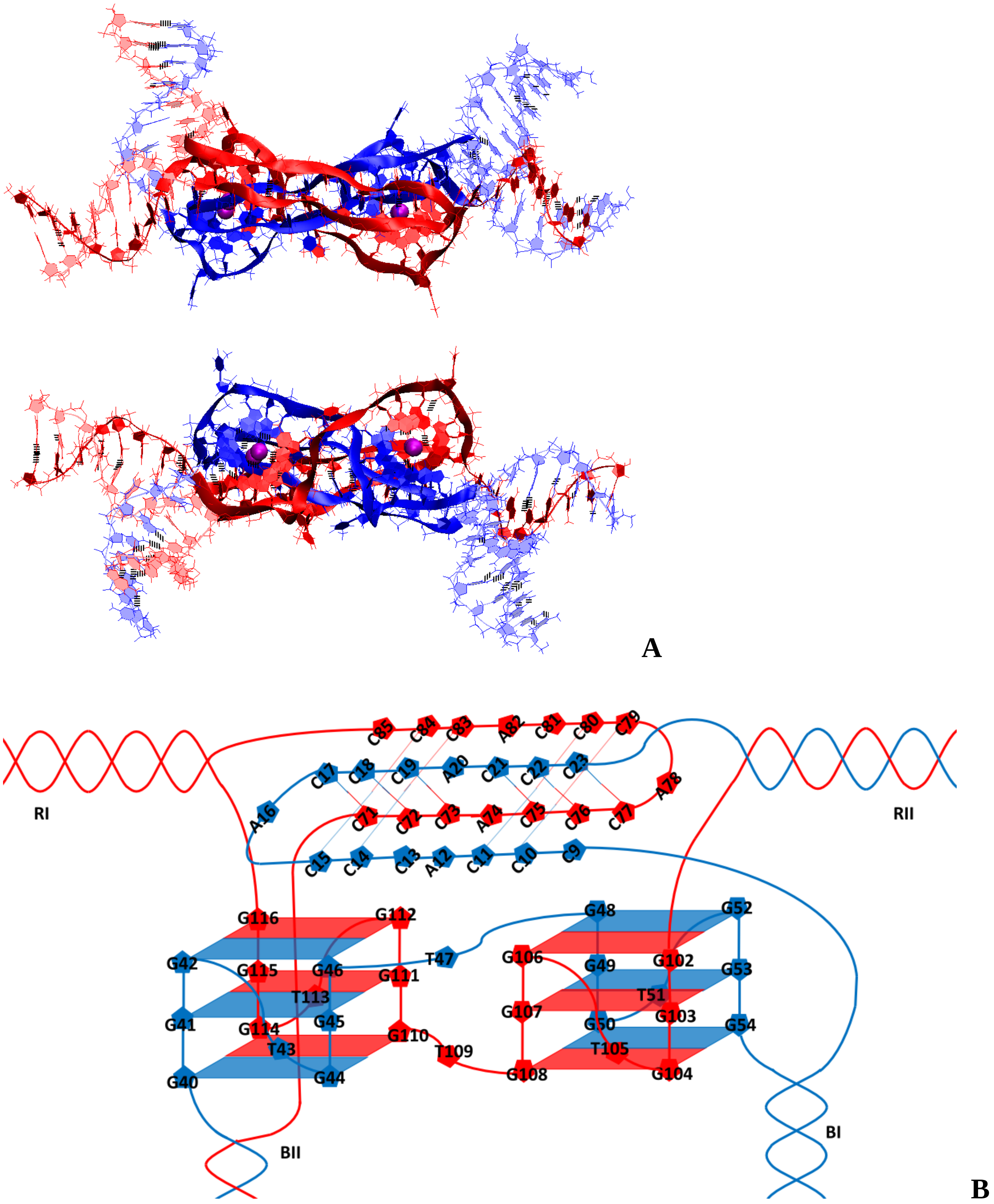


**Figure 3.5.B.S.1. Case 14: "Two parallel G4-dimers in the same plane and head-to-tail IM-dimer between two unmelted fragments of duplexes with exchange and mutual girth of the strands": A** – conformation obtained at the last step of the MD trajectory (side and top view); **B** – the complex scheme.

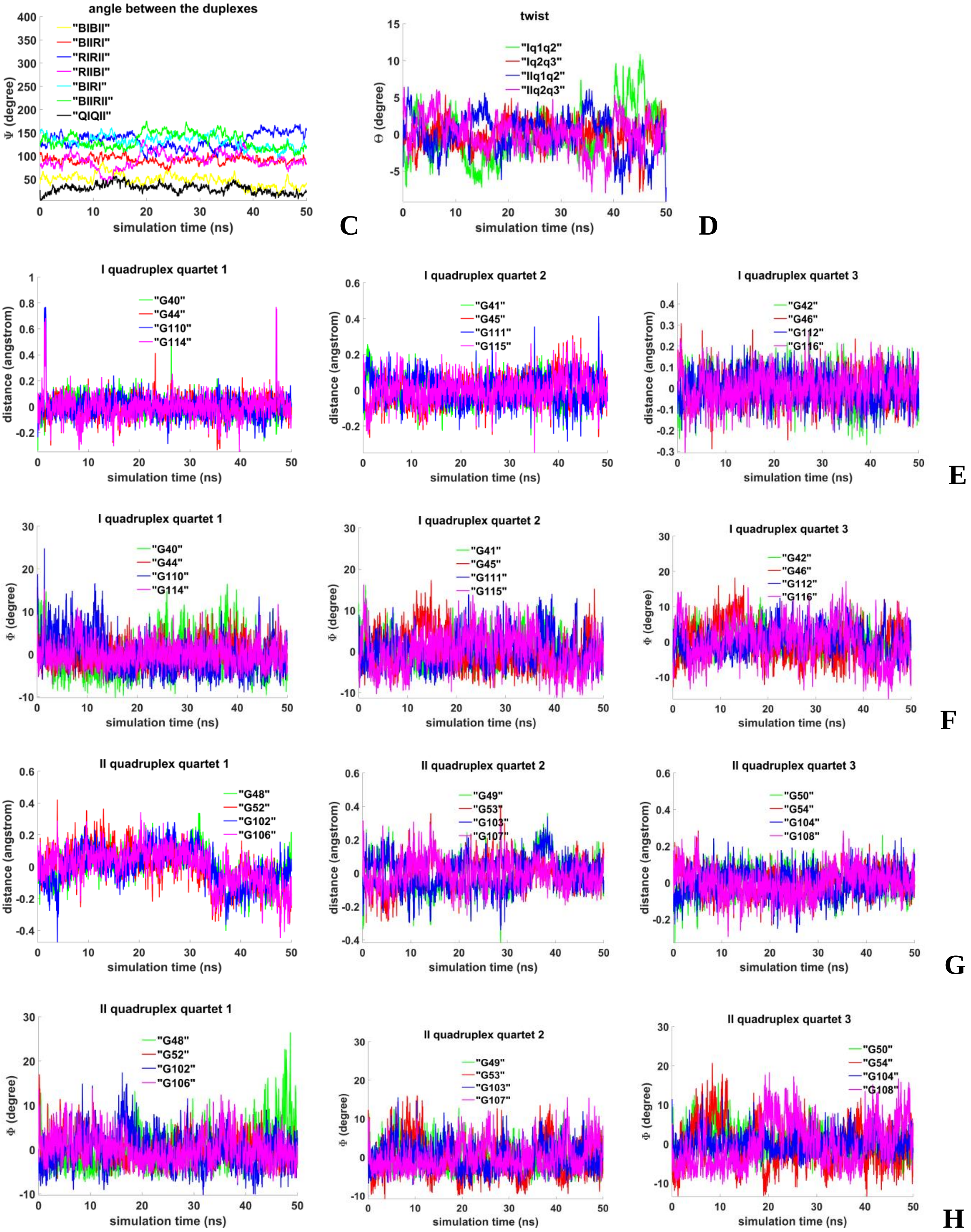


**Figure 3.5.B.S.2. Case 14: "Two parallel G4-dimers in the same plane and head-to-tail IM-dimer between two unmelted fragments of duplexes with exchange and mutual girth of the strands": C** - angles between unmelted fragments of the duplexes, angle between axes passing through COMs of the boundary tetrads (**QIQII**); **D** - angles of rotation of the tetrads relative to each other; **E, G** - distances from COMs of the guanines' bases to COMs of their containing tetrads; **F, H** - angles between normals to the guanines' bases and vectors connecting COMs of the boundary tetrads.

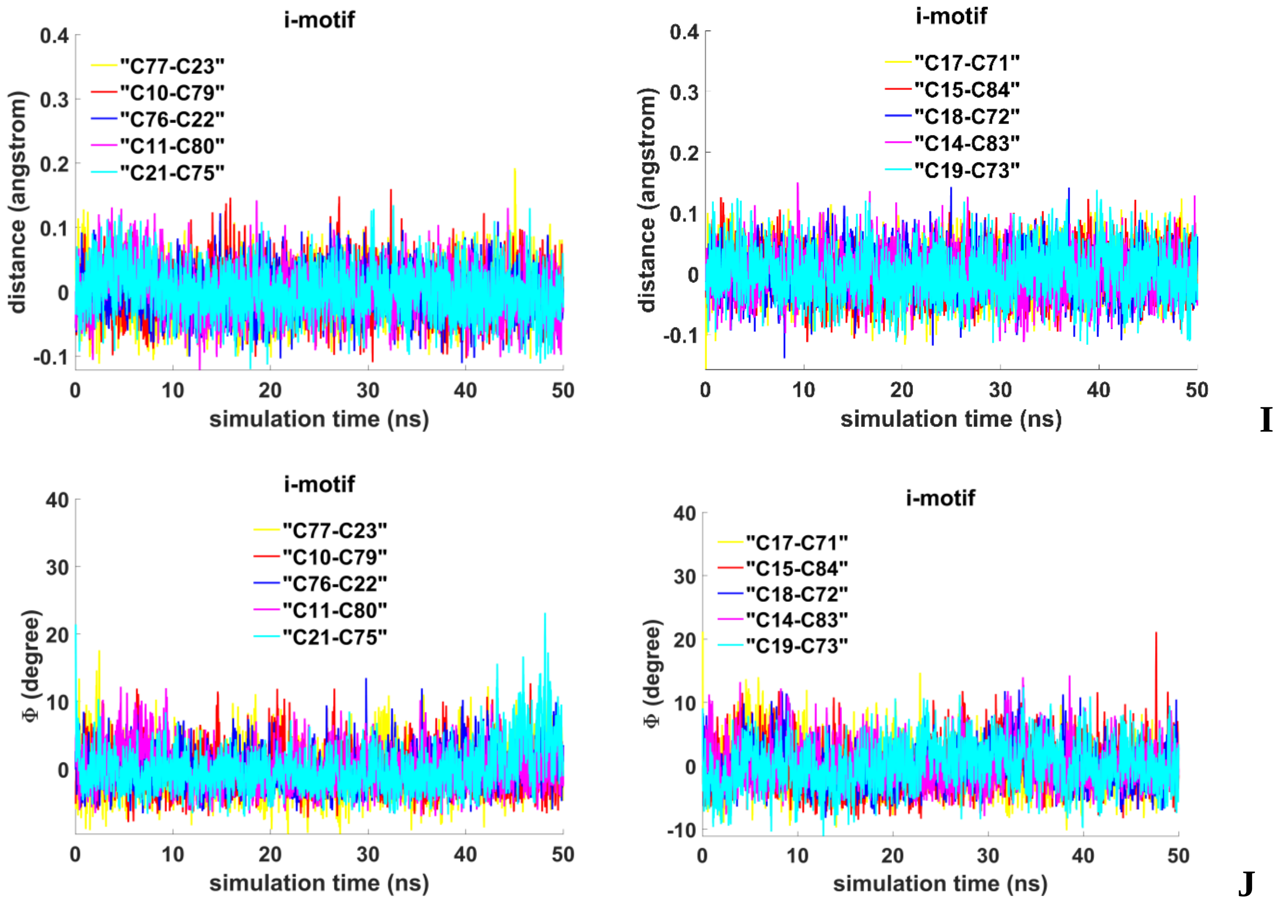


**Figure 3.5.B.S.3. Case 14: "Two parallel G4-dimers in the same plane and head-to-tail IM-dimer between two unmelted fragments of duplexes with exchange and mutual girth of the strands": I** - distances between COMs of the cytosines' bases; **J** - angles between normals to the cytosines' bases.

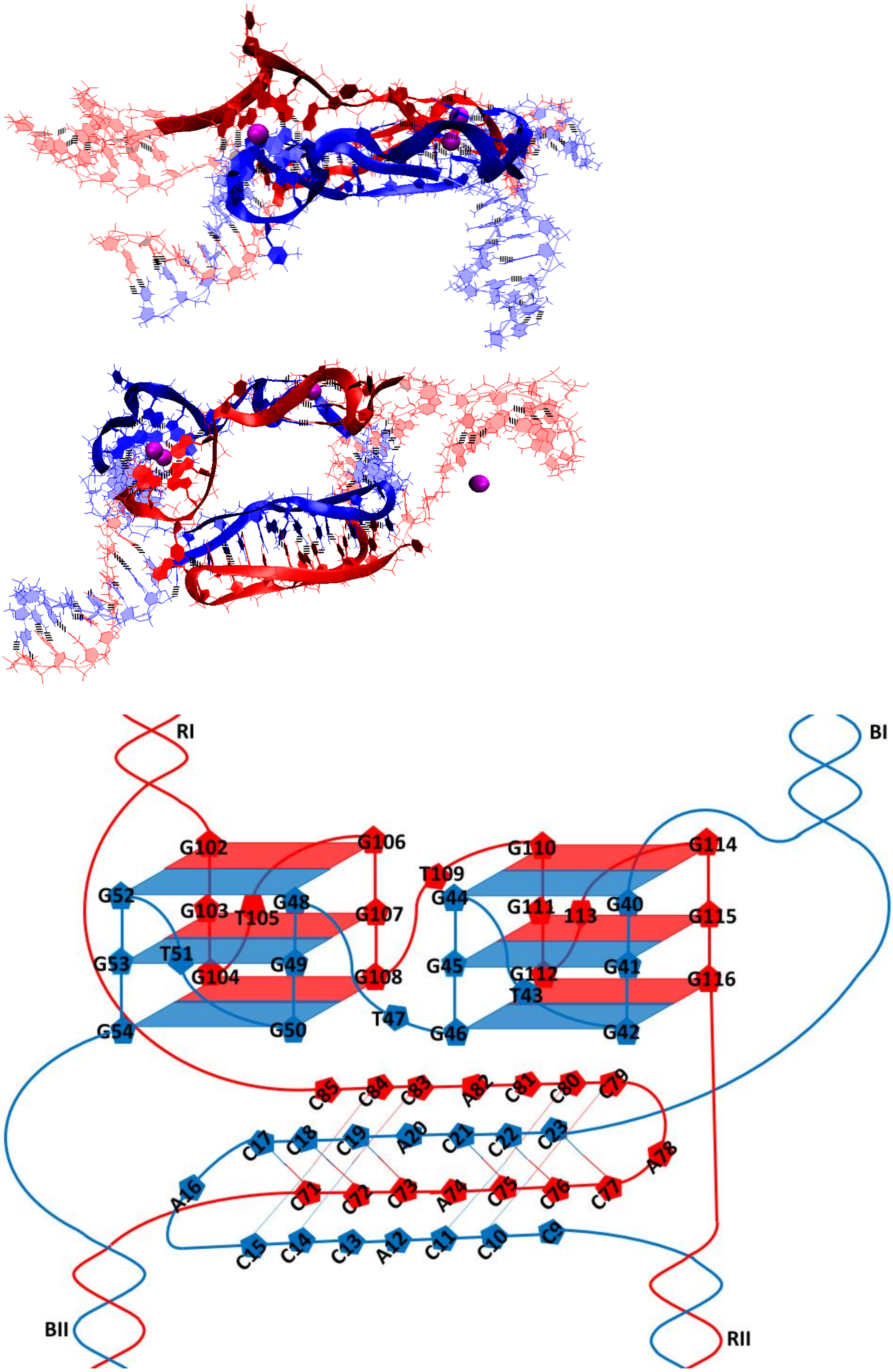


A

B

**Figure 3.5.C.S.1. Case 15: "Two parallel G4-dimers in the same plane and head-to-tail IM-dimer between two unmelted fragments of duplexes with the strands exchange": A** – conformation obtained at the last step of the MD trajectory (side and top view); **B** – the complex scheme.

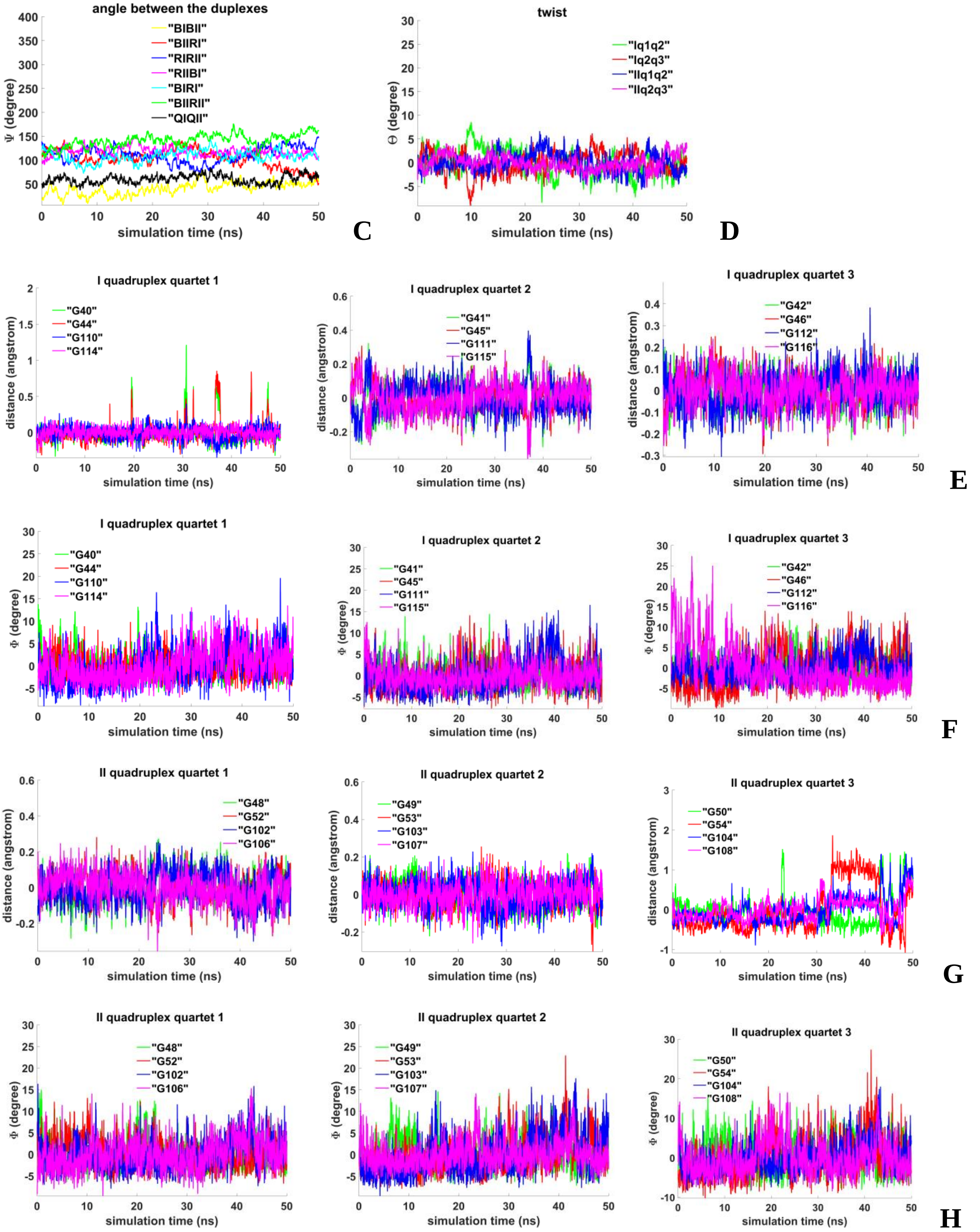


**Figure 3.5.C.S.2. Case 15: "Two parallel G4-dimers in the same plane and head-to-tail IM-dimer between two unmelted fragments of duplexes with the strands exchange" : C** – angles between unmelted fragments of the duplexes, angle between axes passing through COMs of the boundary tetrads (**QIQII**); **D**– angles of rotation of the tetrads relative to each other; **E, G -** distances from COMs of the guanines' bases to COMs of their containing tetrads; **F, H** - angles between normals to the guanines' bases and vectors connecting COMs of the boundary tetrads.

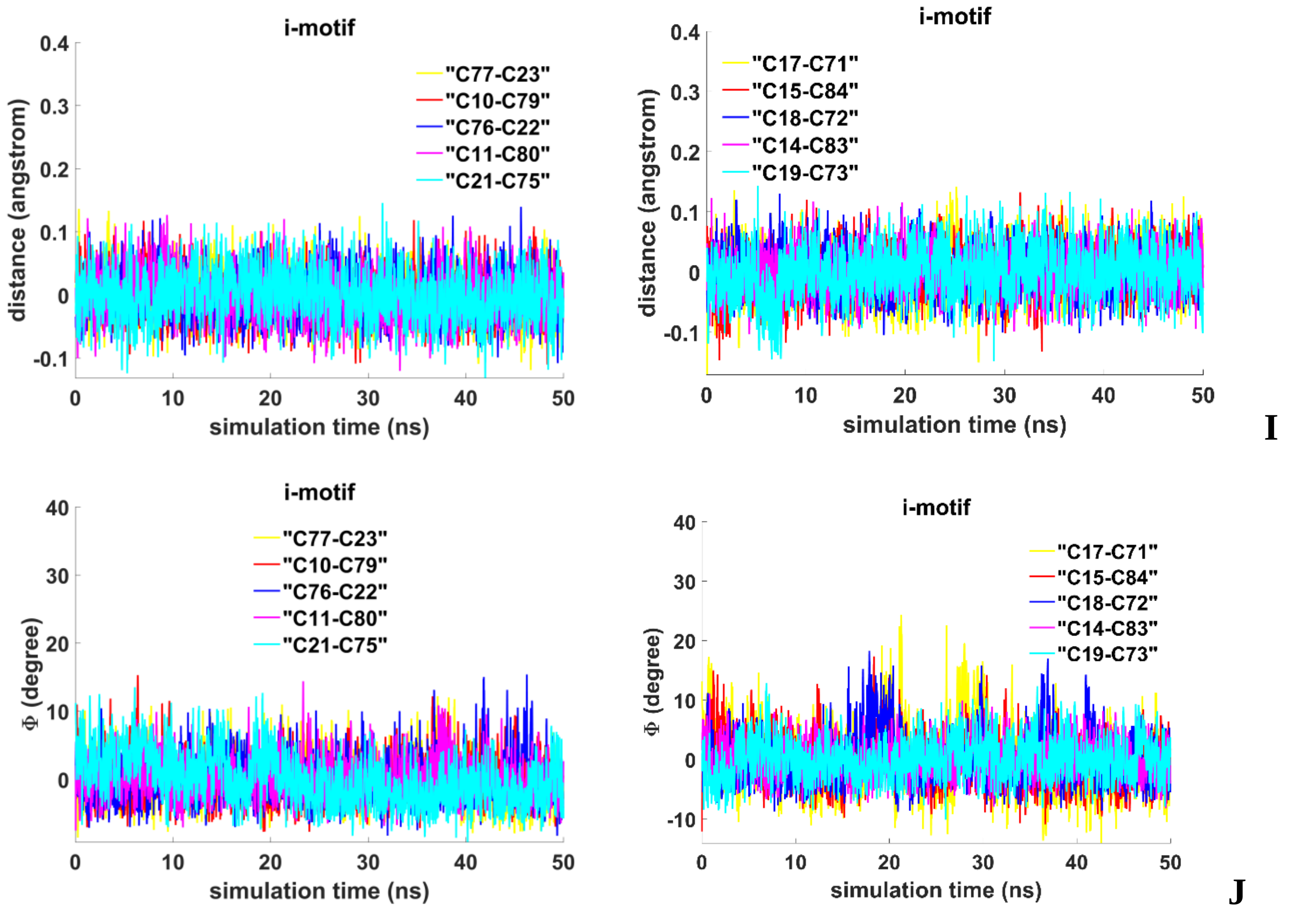


**Figure 3.5.C.S.3. Case 15: "Two parallel G4-dimers in the same plane and head-to-tail IM-dimer between two unmelted fragments of duplexes with the strands exchange": I** - distances between COMs of the cytosines' bases; **J** - angles between normals to the cytosines' bases.

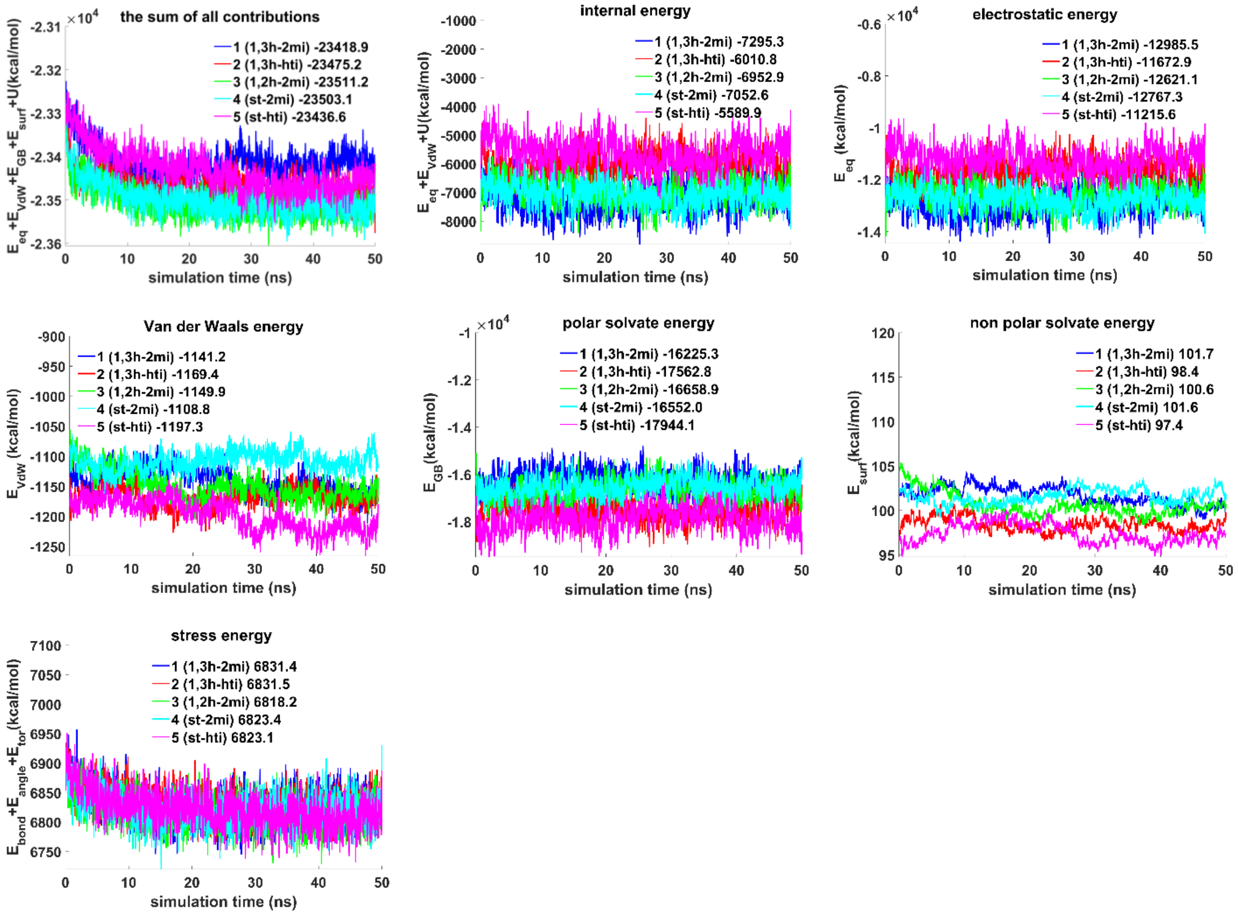


**Figure 3.S.E.1. The contributions to free energy during MD calculations for the variants of bimolecular complexes of unmelted fragments of duplexes containing $(G_3T)_3G_3$ and $(C_3A)_3C_3$ sequences with G4/IM in cases from 1 to 5**. $E_{eq}$ – electrostatic, $E_{vdw}$ - Van der Waals, $E_{GB}$ - polar energy of solvation, $E_{surf}$ - non-polar energy of solvation due to the hydrophobic surface available to the solvent, $U = E_{bond} + E_{angle} + E_{tor}$ , e.g. $E_{bond}$, $E_{angle}$ and $E_{tor}$ –bond, angle and torsion stress energies. The energy plots were smoothed using moving average method (span = 5). Average energy values are indicated in the figure legends.

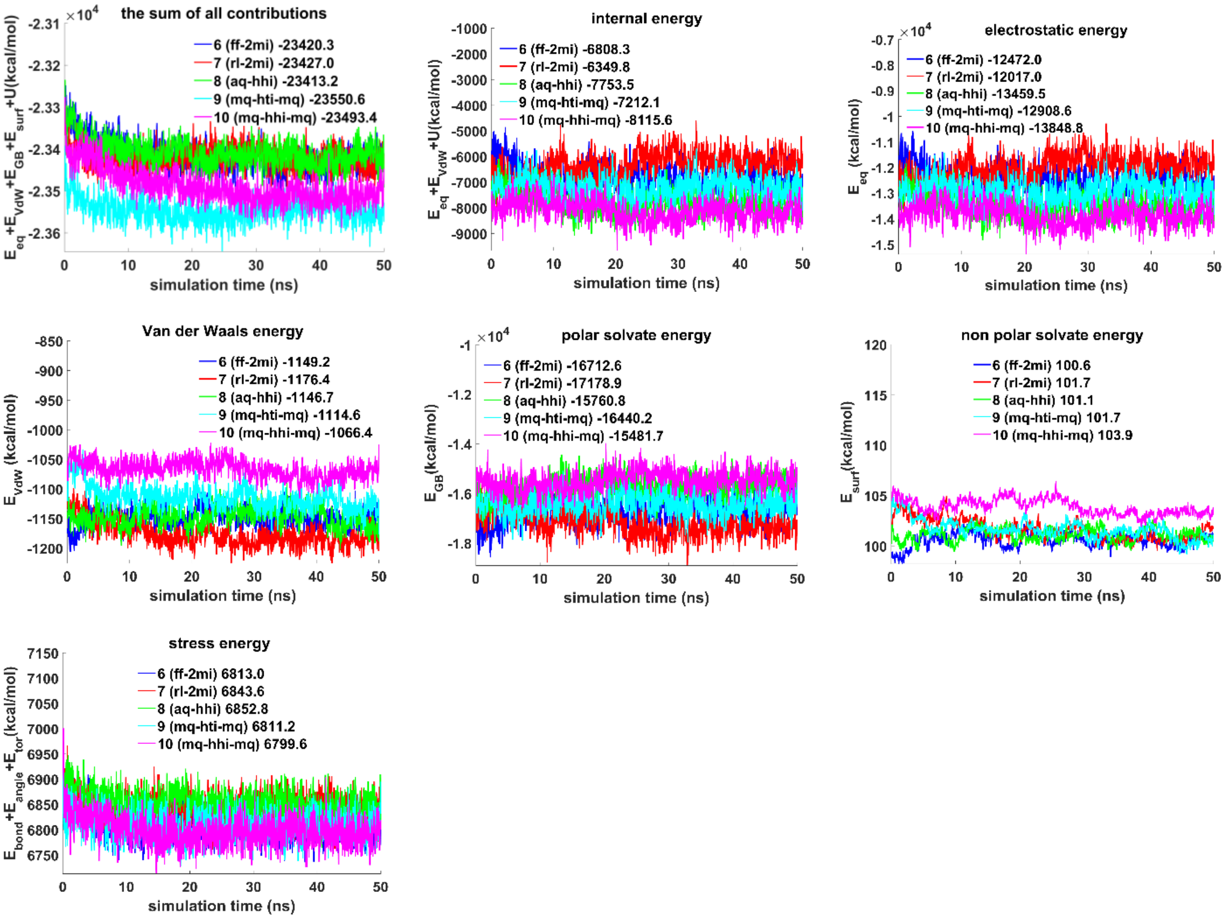


**Figure 3.S.E.2. The contributions to free energy during MD calculations for the variants of bimolecular complexes of unmelted fragments of duplexes containing $(G_3T)_3G_3$ and $(C_3A)_3C_3$ sequences with G4/IM in cases from 6 to 10**. $E_{eq}$ – electrostatic, $E_{vdw}$ - Van der Waals, $E_{GB}$ - polar energy of solvation, $E_{surf}$ - non-polar energy of solvation due to the hydrophobic surface available to the solvent, $U = E_{bond} + E_{angle} + E_{tor}$ , e.g. $E_{bond}$, $E_{angle}$ and $E_{tor}$ –bond, angle and torsion stress energies. The energy plots were smoothed using moving average method (span = 5). Average energy values are indicated in the figure legends.

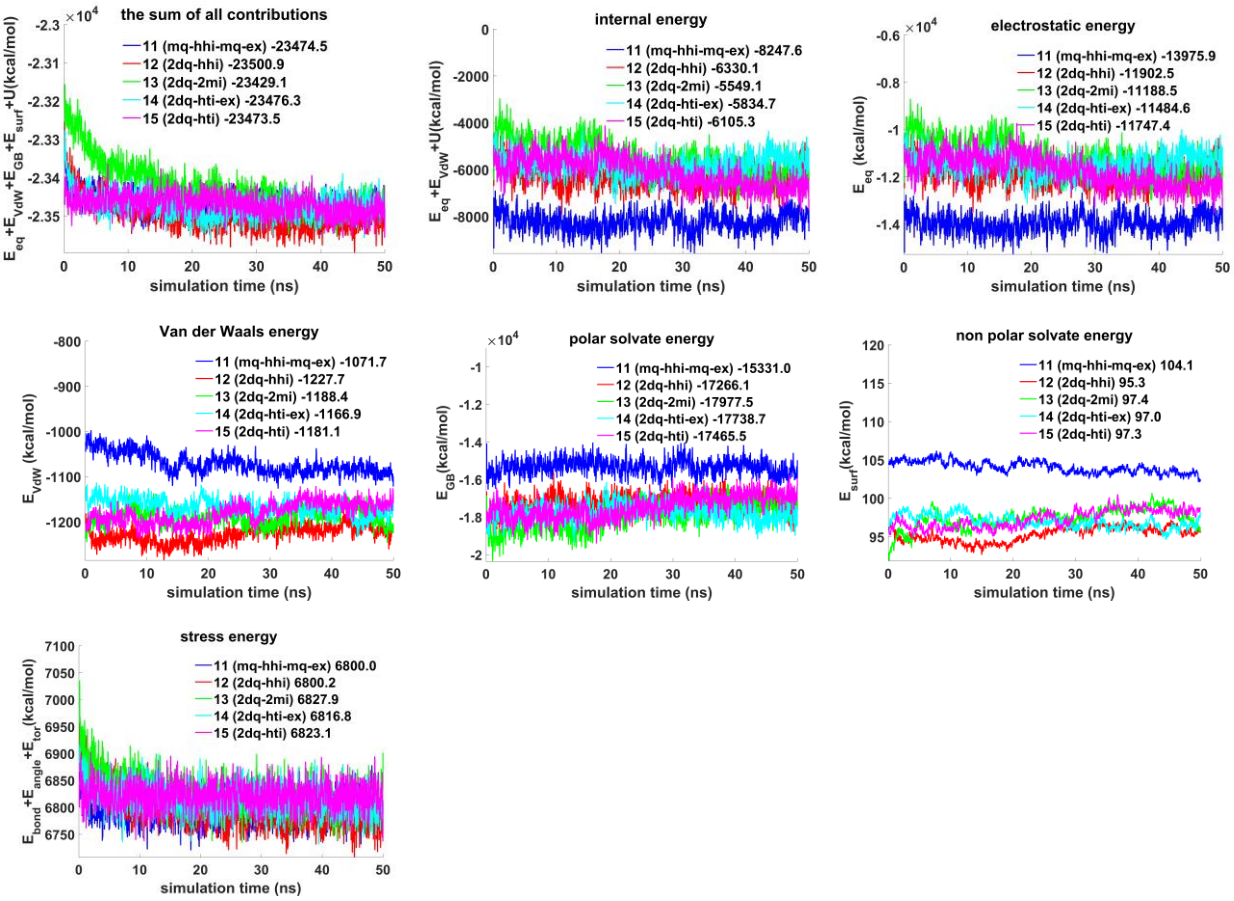


**Figure 3.S.E.3. The contributions to free energy during MD calculations for the variants of bimolecular complexes of unmelted fragments of duplexes containing $(G_3T)_3G_3$ and $(C_3A)_3C_3$ sequences with G4/IM in cases from 11 to 15**. $E_{eq}$ – electrostatic, $E_{vdw}$ - Van der Waals, $E_{GB}$ - polar energy of solvation, $E_{surf}$ - non-polar energy of solvation due to the hydrophobic surface available to the solvent, $U = E_{bond} + E_{angle} + E_{tor}$ , e.g. $E_{bond}$, $E_{angle}$ and $E_{tor}$ –bond, angle and torsion stress energies. The energy plots were smoothed using moving average method (span = 5). Average energy values are indicated in the figure legends.

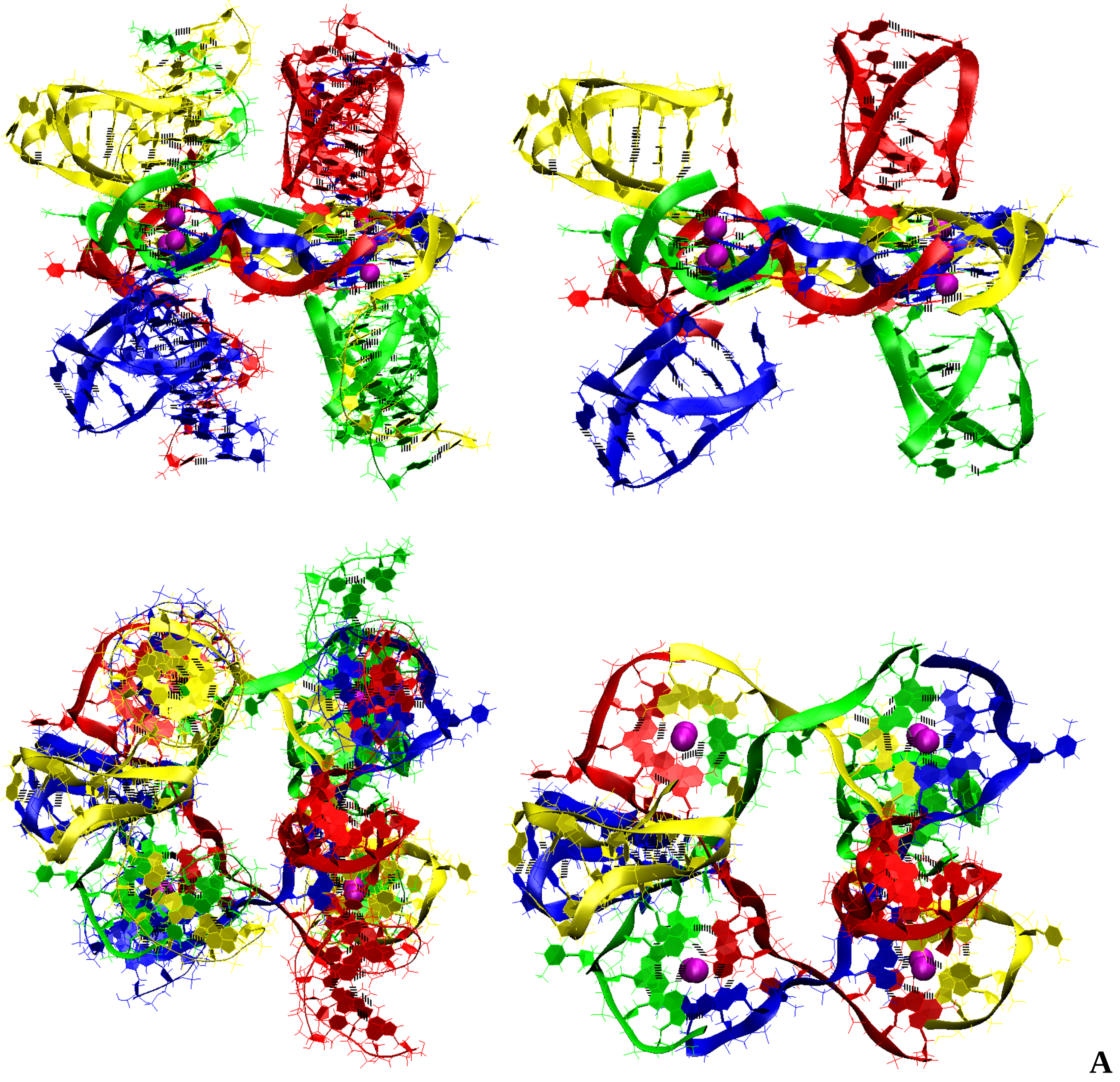


**Figure 4.1.S.1. Case 1: “Four parallel G4-dimers in the same plane and four IM-monomers between the unmelted fragments of duplexes with exchange and mutual girth of the strands”: A -** conformation obtained at the last step of the MD trajectory. Top row is the side view, bottom row is the top view. On the left in both rows are views with unmelted fragments of the duplexes, on the right there are only G4s and IMs.

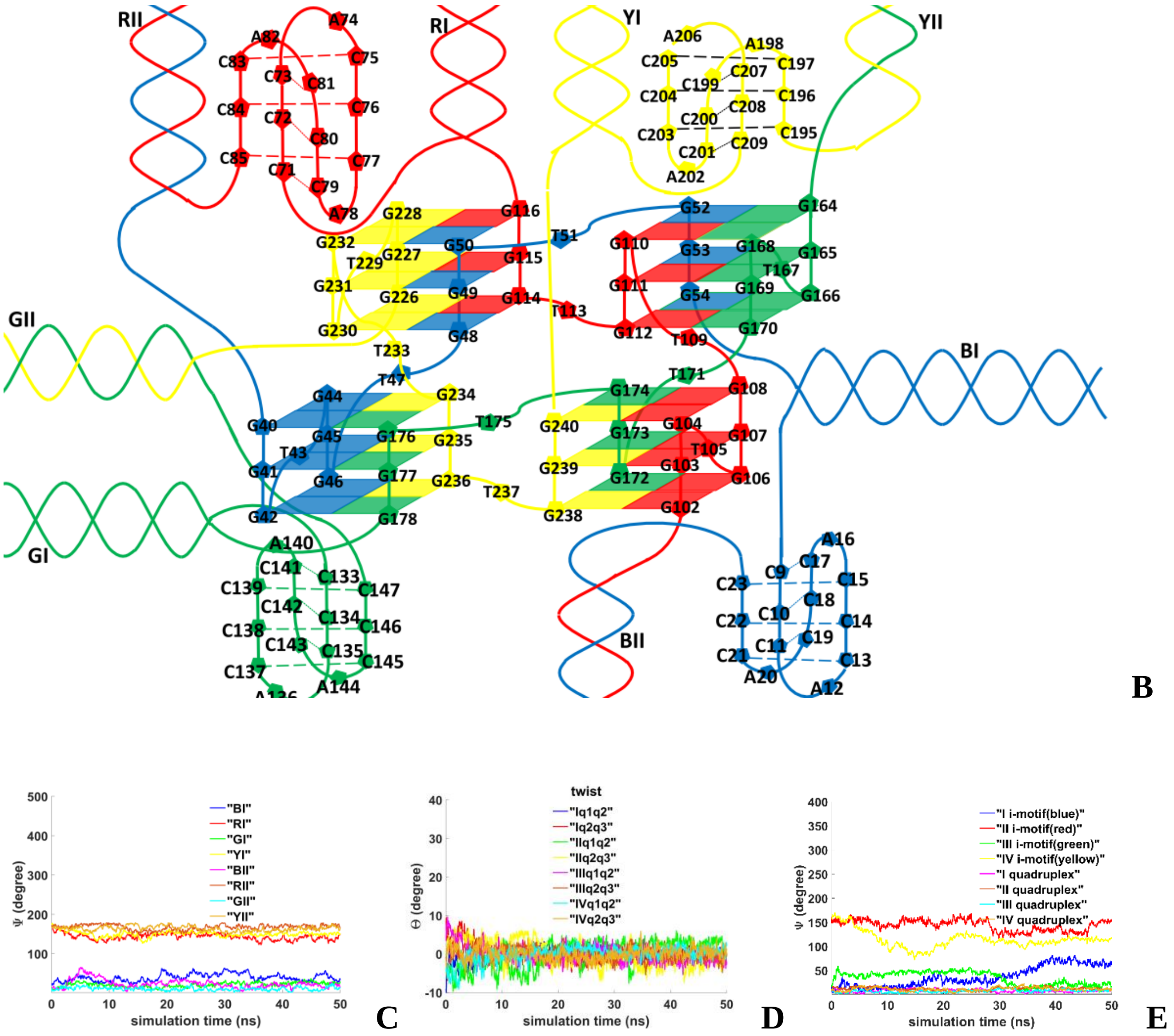


**Figure 4.1.S.2. Case 1: "Four parallel G4-dimers in the same plane and four IM-monomers between the unmelted fragments of duplexes with exchange and mutual girth of the strands": B** – the complex scheme; **C** – angles between straight lines, passing through the COMs of the first and the last complementary pairs of unmelted fragments of duplexes, and planes, containing COMs of the G4s tetrads; **D**– angles of rotation of the tetrads relative to each other; **E** - angles between the straight line, passing through the COM of all upper tetrads and the COM of all lower tetrads, and the straight lines, passing through the COMs of the upper and lower tetrads, in the G4s' case and straight lines, passing through the COMs of the boundary cytosine pairs, in the IMs' case.

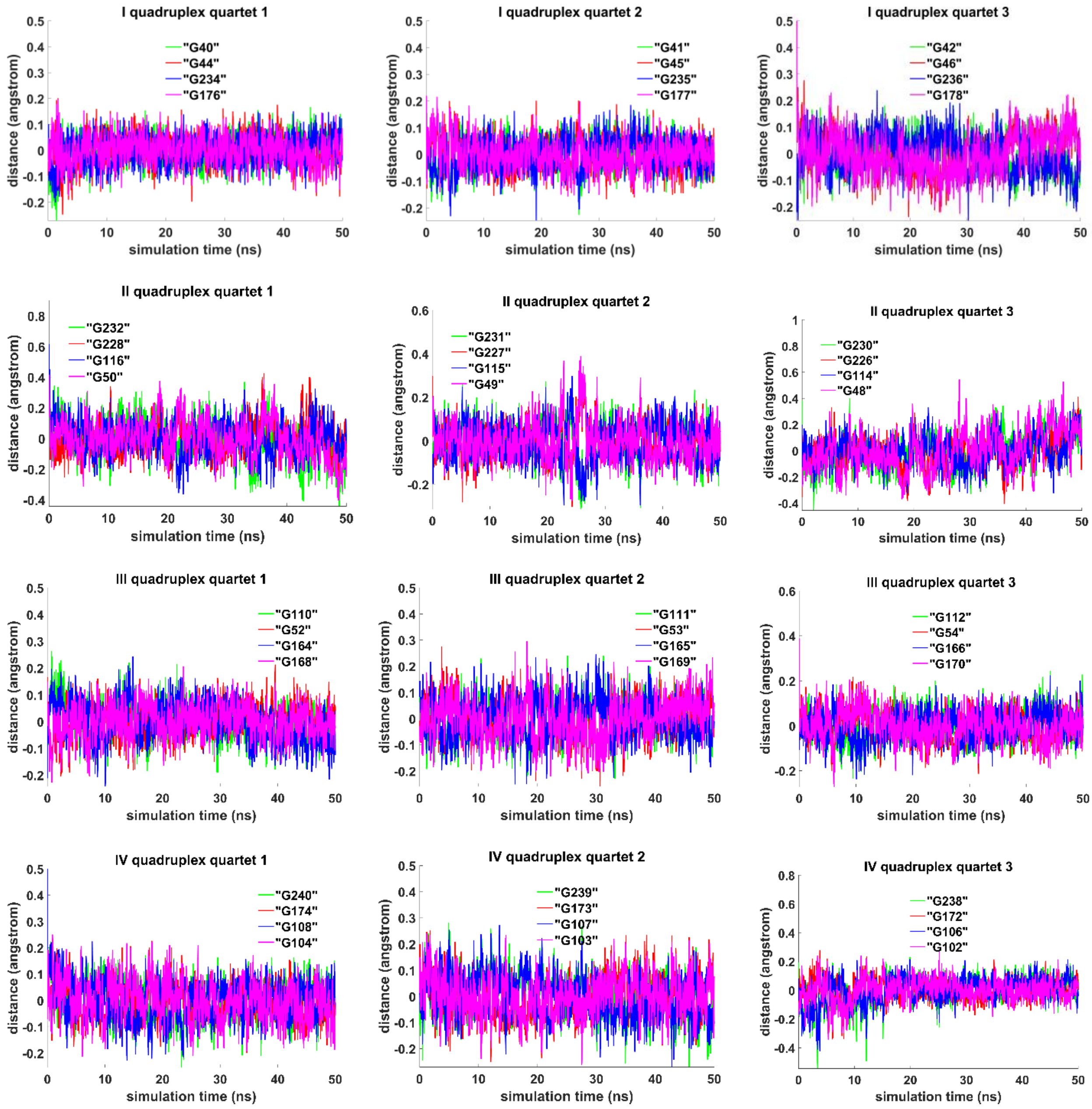


**Figure 4.1.S.3. Case 1: "Four parallel G4-dimers in the same plane and four IM-monomers between the unmelted fragments of duplexes with exchange and mutual girth of the strands":** distances from COMs of the guanines' bases to COMs of their containing tetrad.

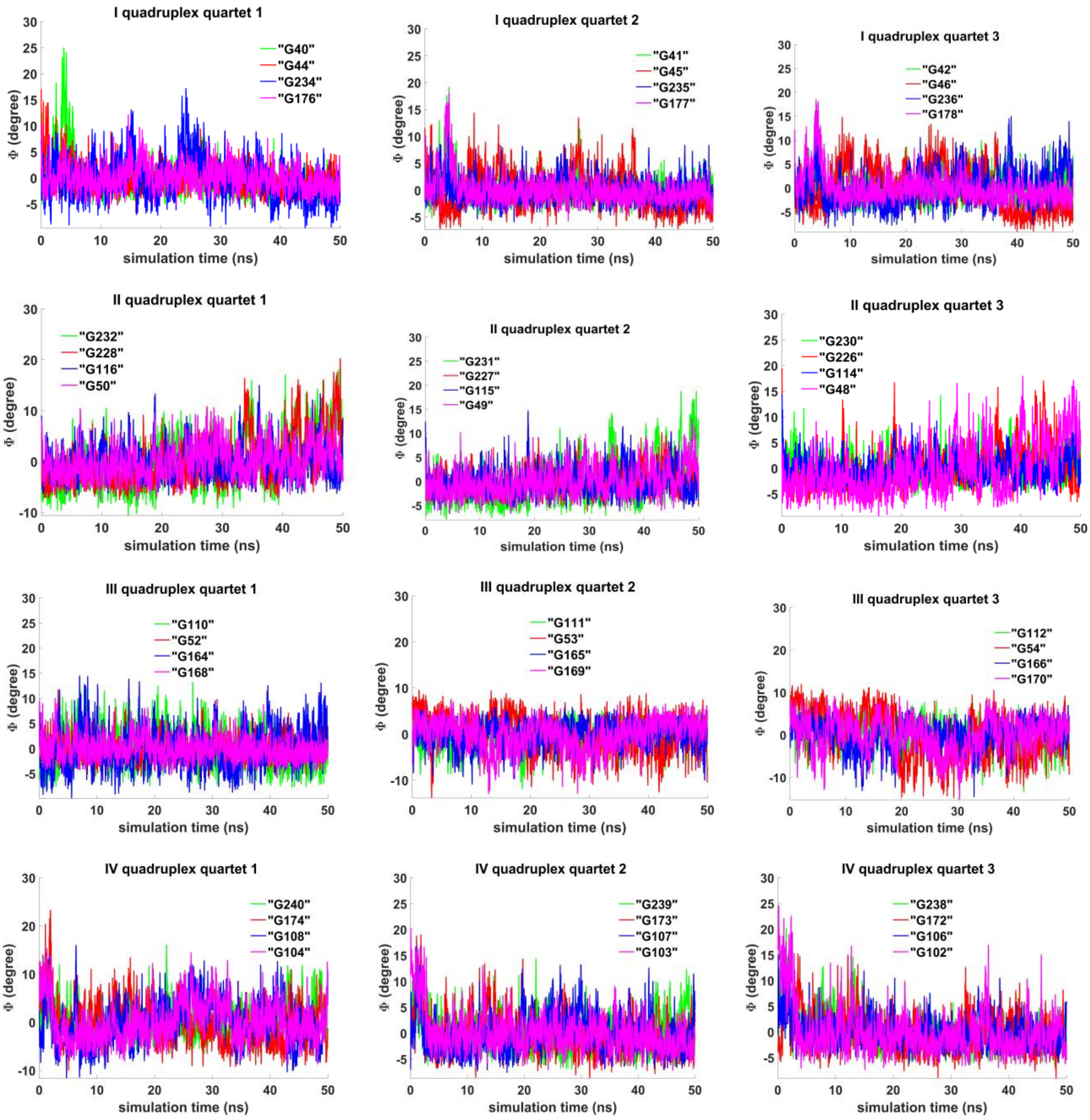


**Figure 4.1.S.4. Case 1: "Four parallel G4-dimers in the same plane and four IM-monomers between the unmelted fragments of duplexes with exchange and mutual girth of the strands":** angles between normals to the guanine' bases and vectors connecting COMs of the boundary tetrads.

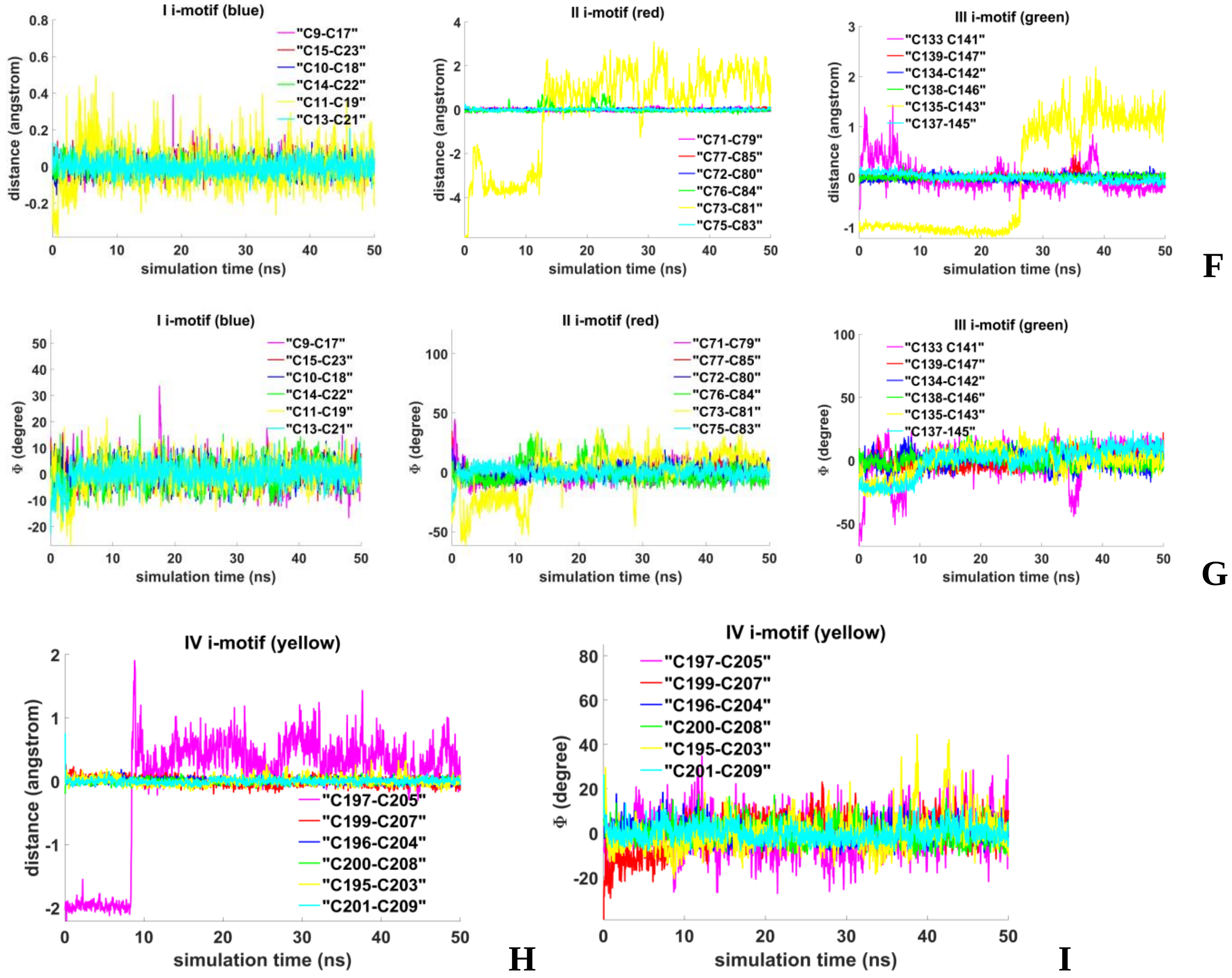


**Figure 4.1.S.5. Case 1: "Four parallel G4-dimers in the same plane and four IM-monomers between the unmelted fragments of duplexes with exchange and mutual girth of the strands": F, H** - distances between COMs of the cytosines' bases; **G, I** - angles between normals to the cytosines' bases.

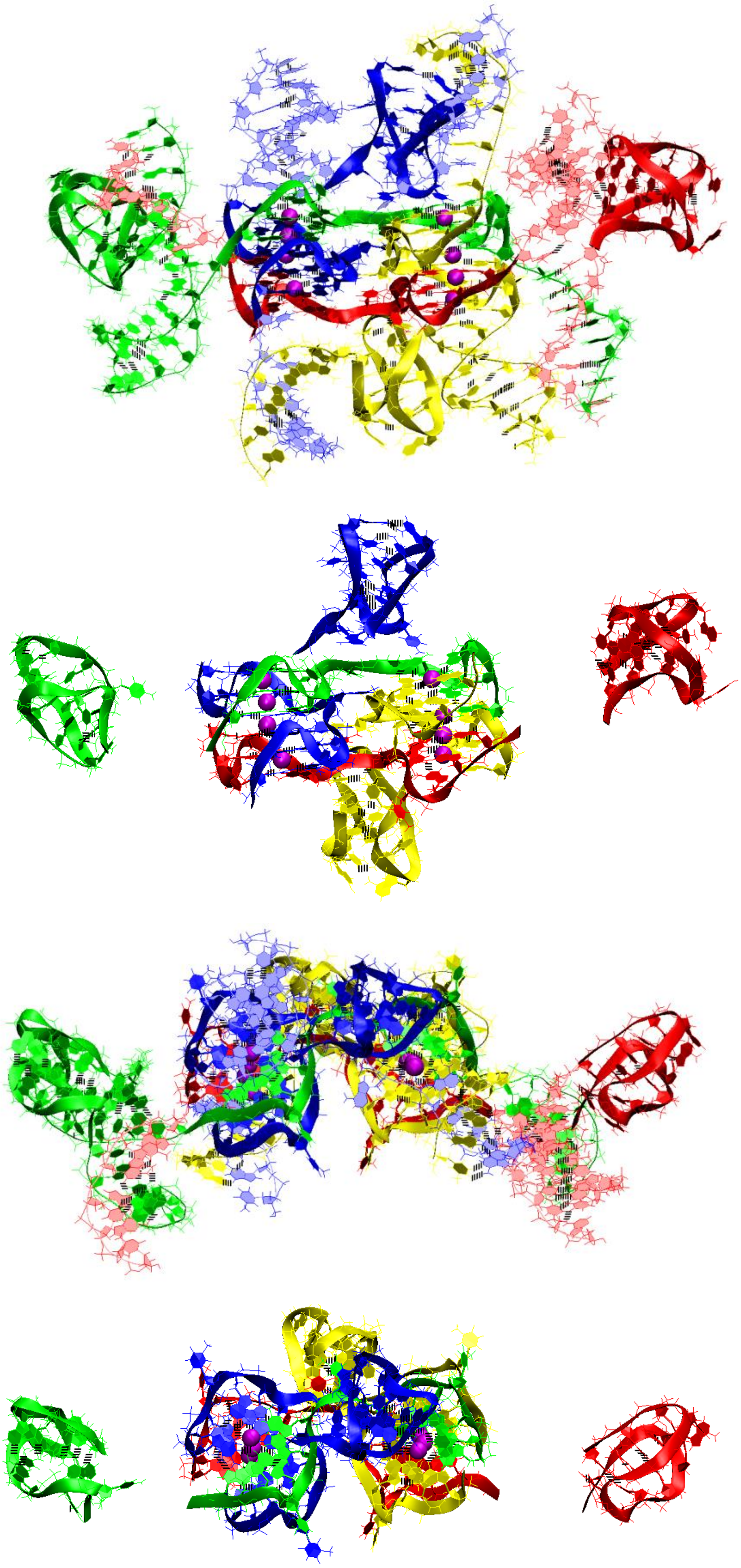

A

**Figure 4.2.S.1. Case 2: "Four parallel G4-dimers in two stacks and four IM-monomers with exchange and mutual girth of the strands": A-** conformation obtained at the last step of the MD trajectory. The first row is the side view with unmelted fragments of the duplexes, the second row is the side view with only G4s and IMs, the third row is top view with unmelted fragments of the duplexes, and the fourth row is top view with only G4s and IMs.

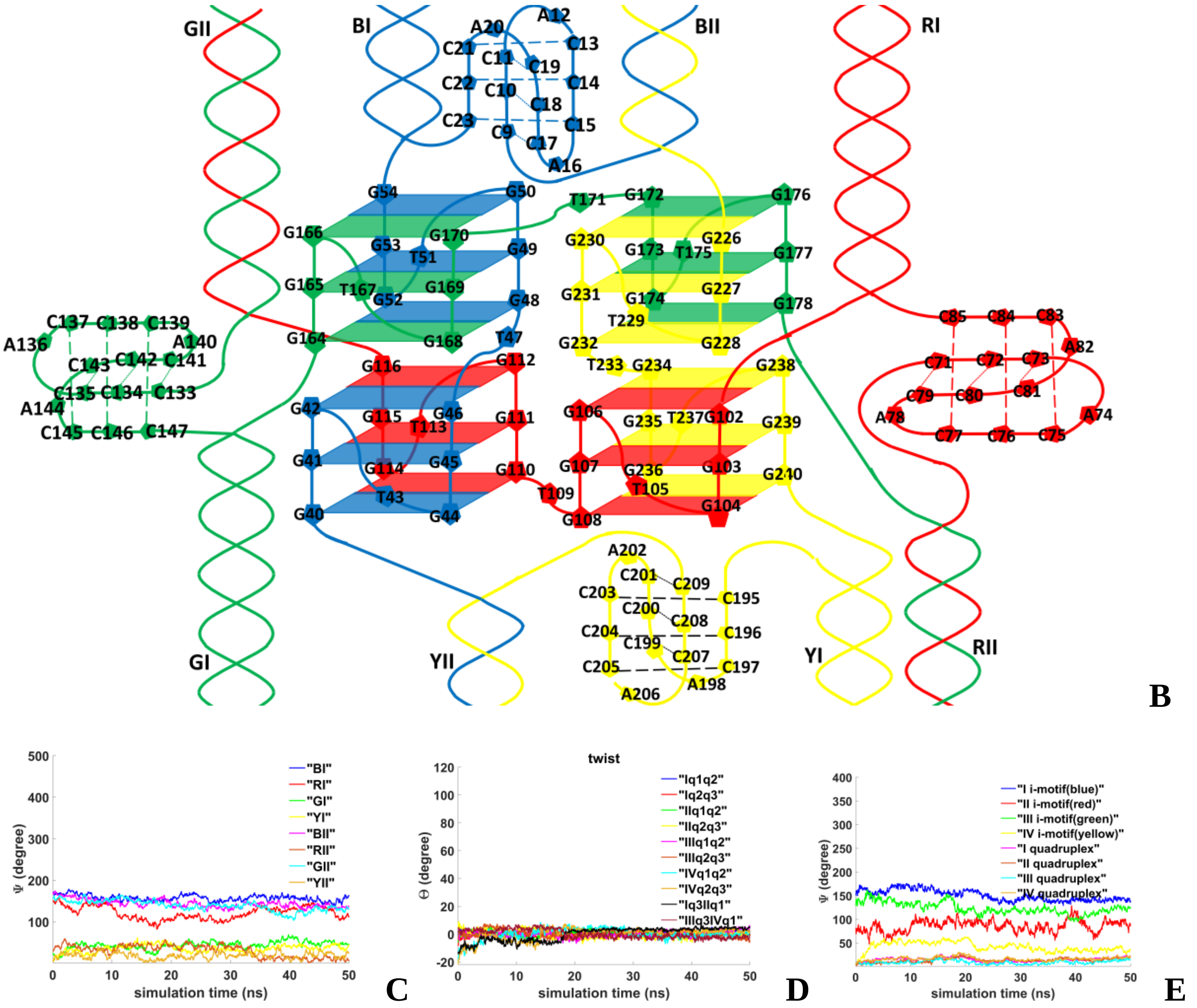


**Figure 4.2.S.2. Case 2: "Four parallel G4-dimers in two stack and four IM-monomers with exchange and mutual girth of the strands": B** – the complex scheme; **C** – angles between straight lines, passing through COMs of the first and the last complementary pairs of unmelted fragments of the duplexes, and straight line, passing through COMs of boundary tetrads of the G4s; **D**– angles of rotation of the tetrads relative to each other; **E** - angles between the straight line, passing through the COM of all upper tetrads and the COM of all lower tetrads, and the straight lines, passing through the COMs of the upper and lower tetrads, in the G4s' case and straight lines, passing through the COMs of the boundary cytosine pairs, in the IMs' case.

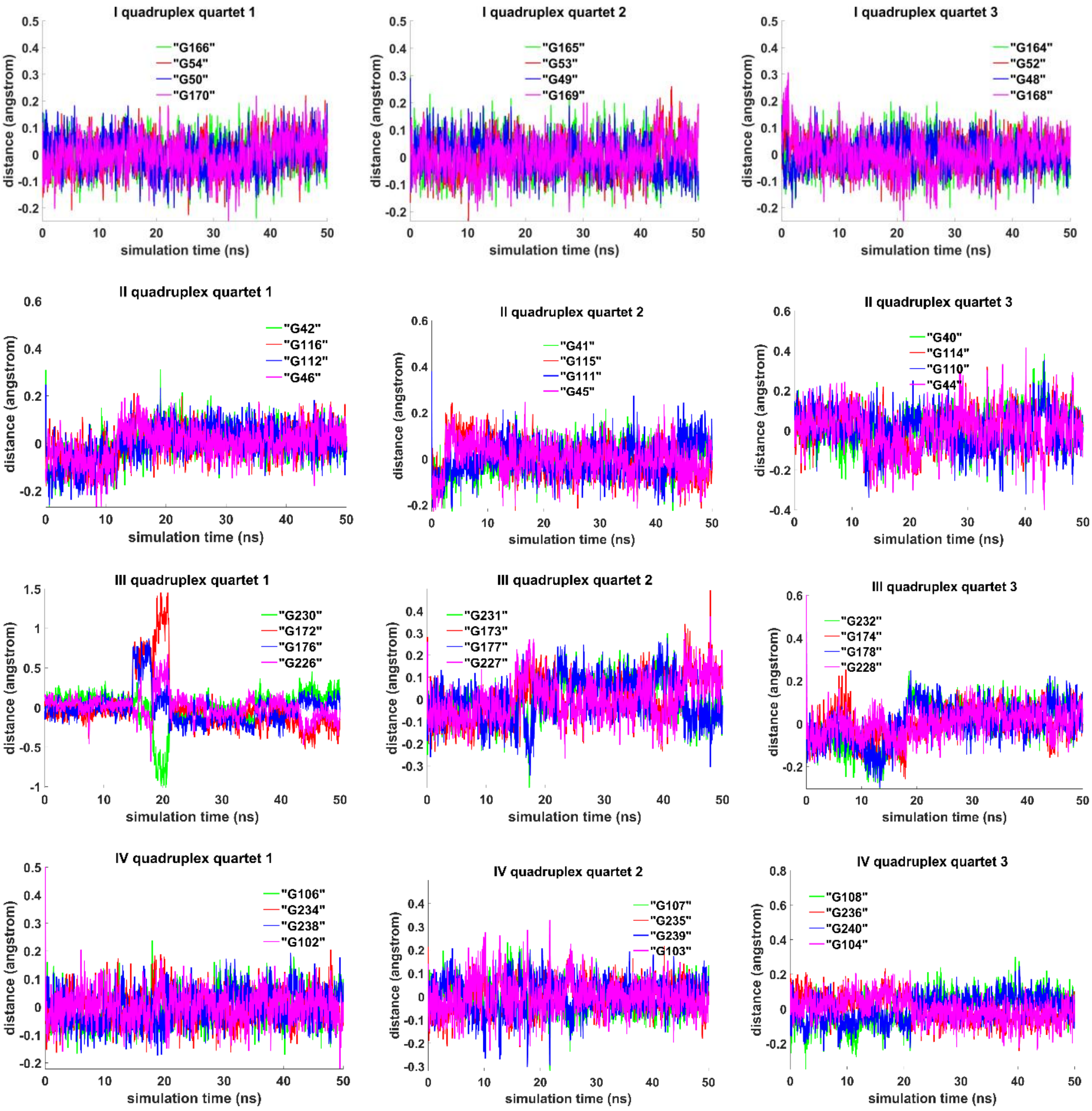


**Figure 4.2.S.3. Case 2: "Four parallel G4-dimers in two stack and four IM-monomers with exchange and mutual girth of the strands":** distances from COMs of the guanines' bases to COMs of their containing tetrad.

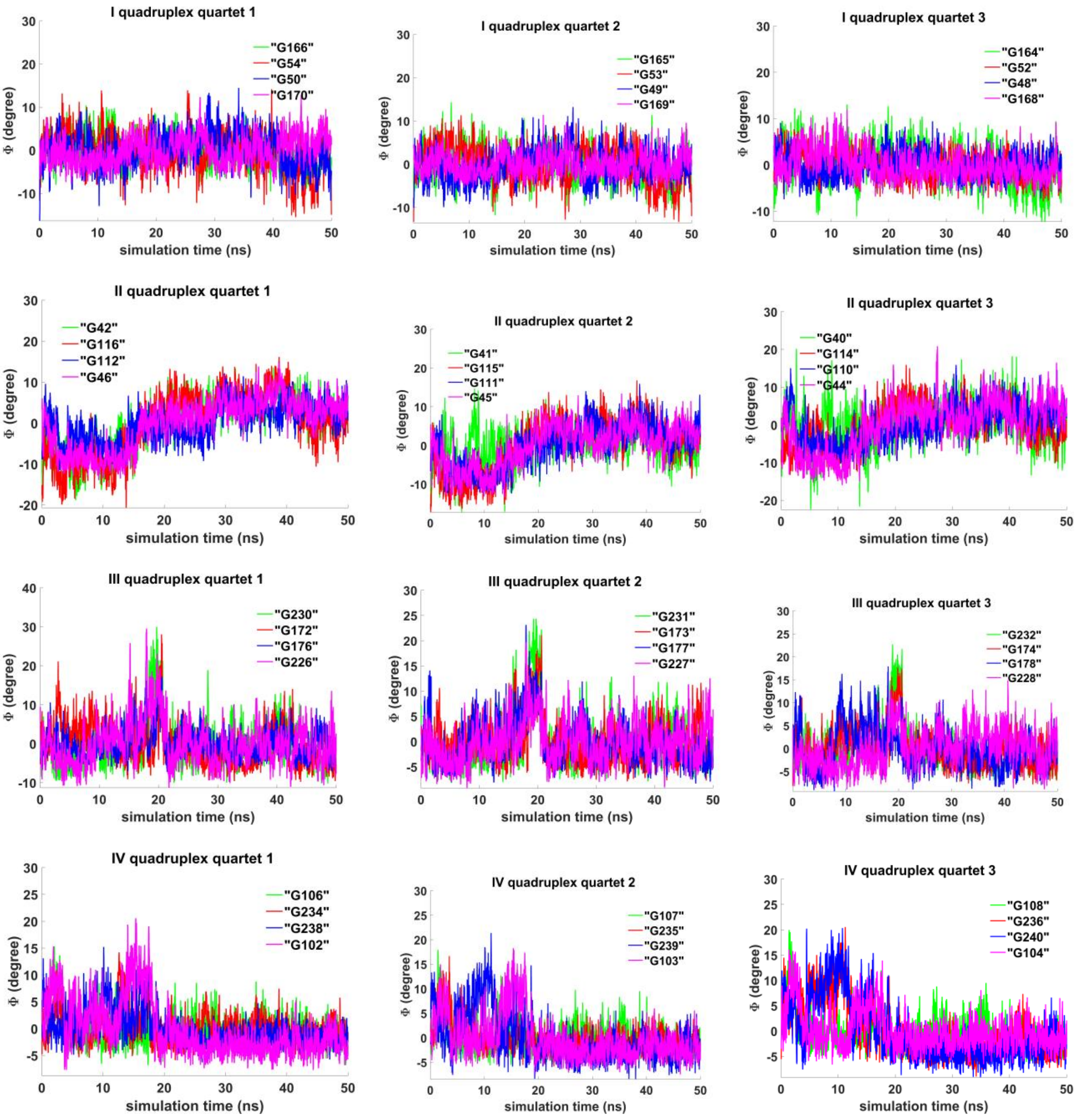


**Figure 4.2.S.4. Case 2: "Four parallel G4-dimers in two stack and four IM-monomers with exchange and mutual girth of the strands":** angles between normals to the guanine' bases and vectors connecting COMs of the boundary tetrads.

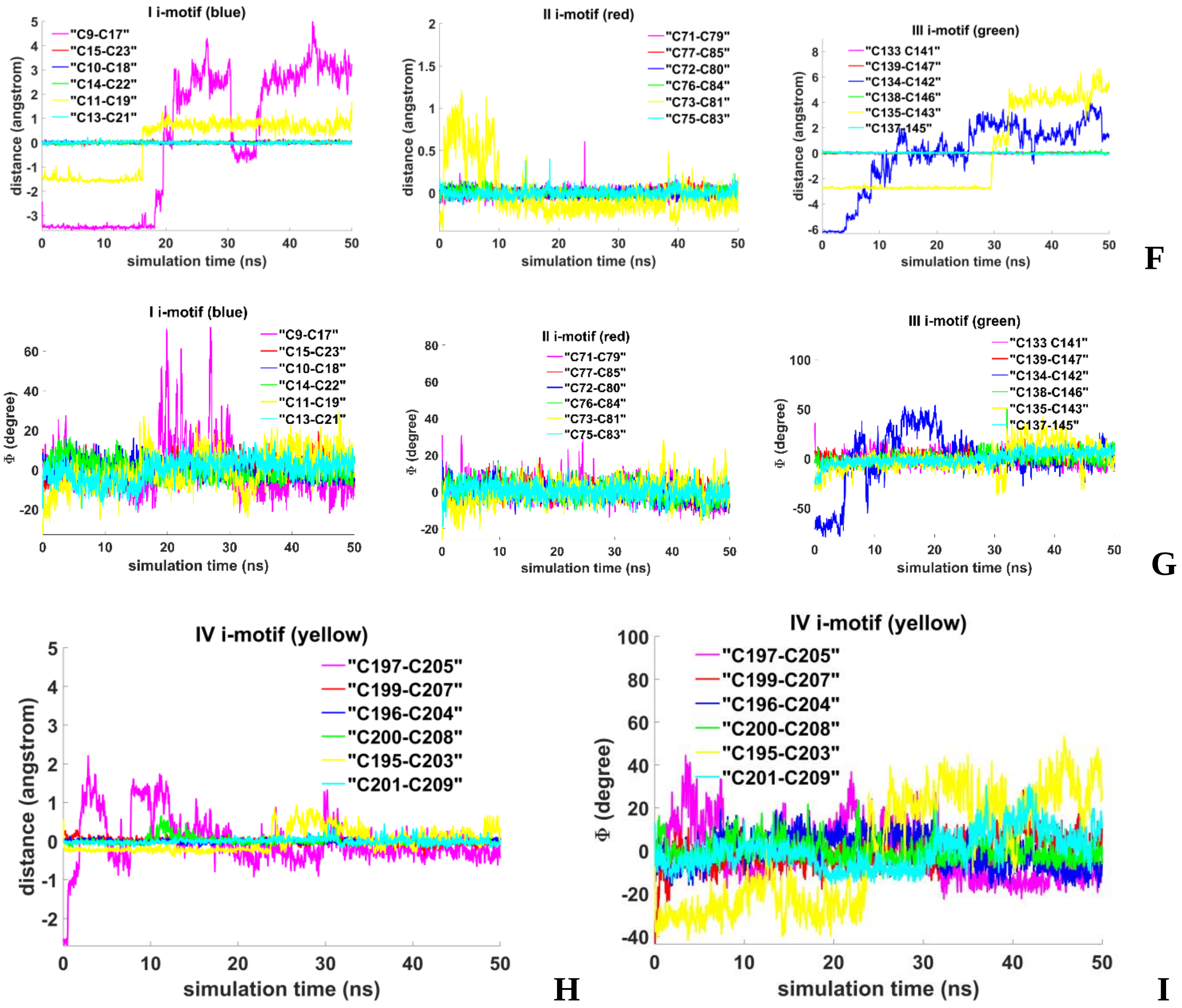


**Figure 4.2.S.5. Case 2: "Four parallel G4-dimers in two stack and four IM-monomers with exchange and mutual girth of the strands": F, H** - distances between COMs of the cytosines' bases; **G, I** - angles between normals to the cytosines' bases.

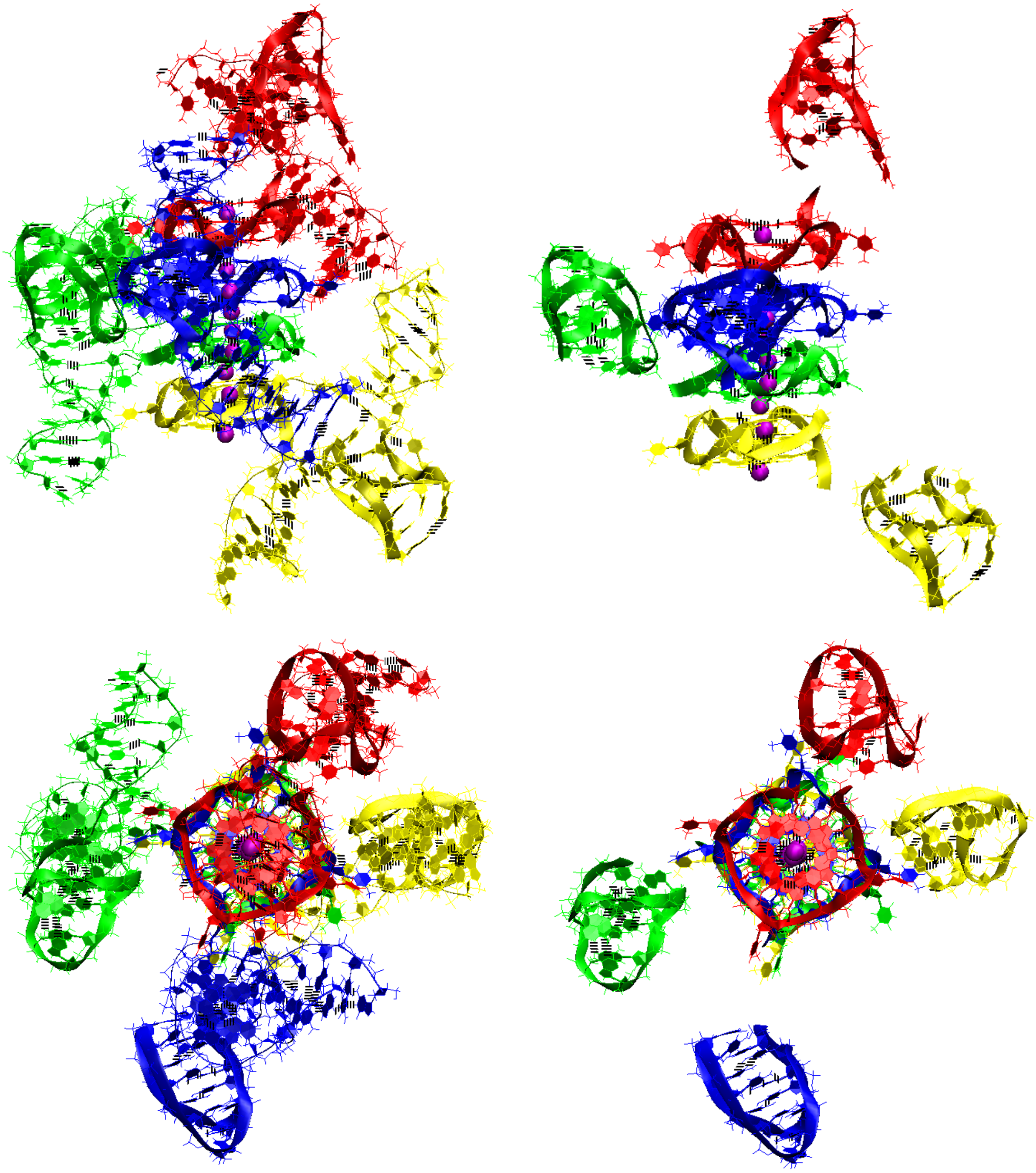

**A**

**Figure 4.3.S.1. Case 3: “Stacking of four parallel G4-dimers and four IM-monomers”: A-** conformation obtained at the last step of the MD trajectory. Top row is the side view, bottom row is the top view. On the left in both rows are views with unmelted fragments of the duplexes, on the right there are only G4s and IMs.

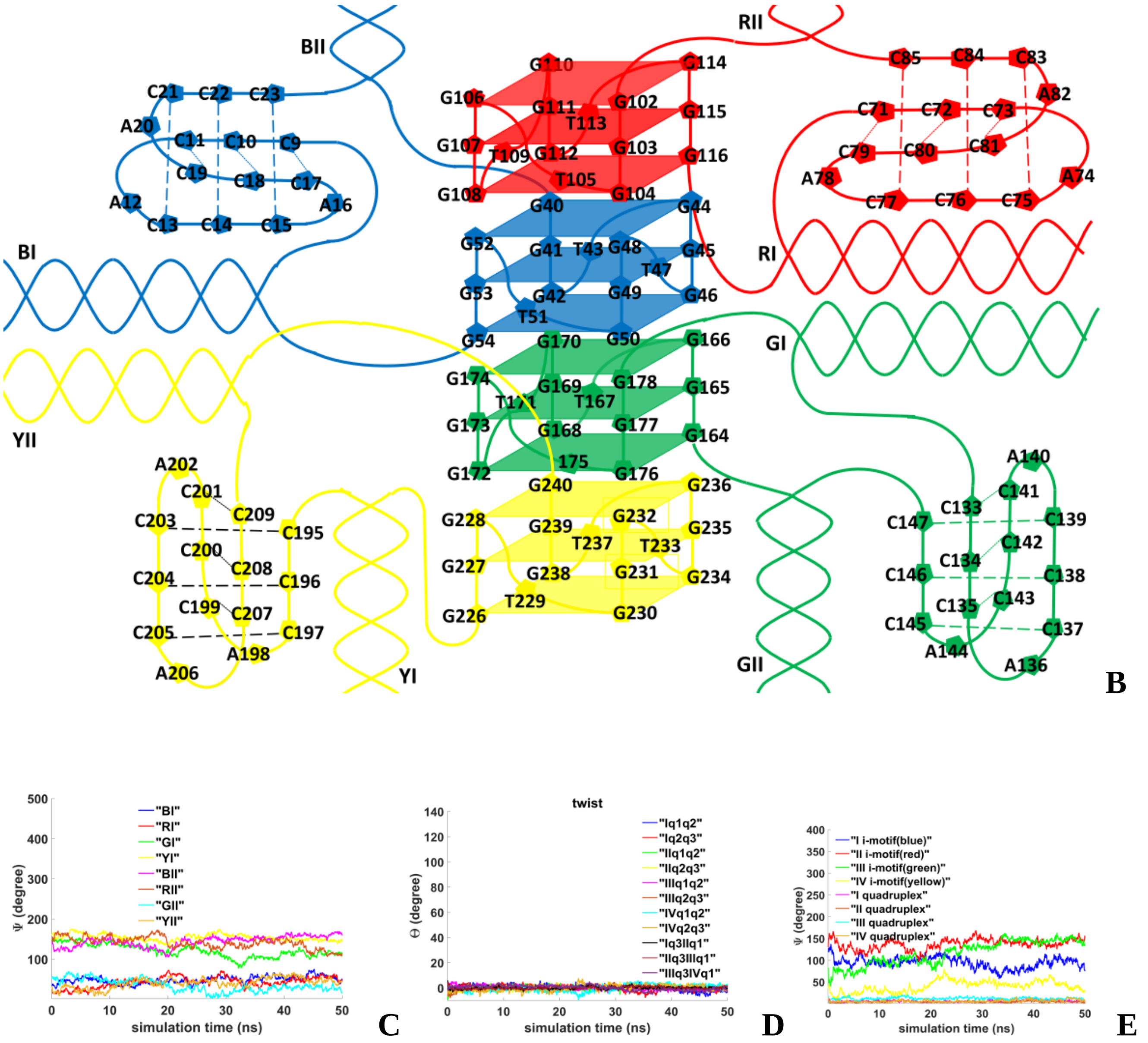


**Figure 4.3.S.2. Case 3: "Stacking of four parallel G4-dimers and four IM-monomers": B** – the complex scheme; **C** – angles between straight lines, passing through COMs of the first and the last complementary pairs of unmelted fragments of the duplexes, and straight line, passing through COMs of boundary tetrads of the G4s; **D**– angles of rotation of the tetrads relative to each other; **E** - angles between the straight line, passing through the COM of all upper tetrads and the COM of all lower tetrads, and the straight lines, passing through the COMs of the upper and lower tetrads, in the G4s' case and straight lines, passing through the COMs of the boundary cytosine pairs, in the IMs' case.

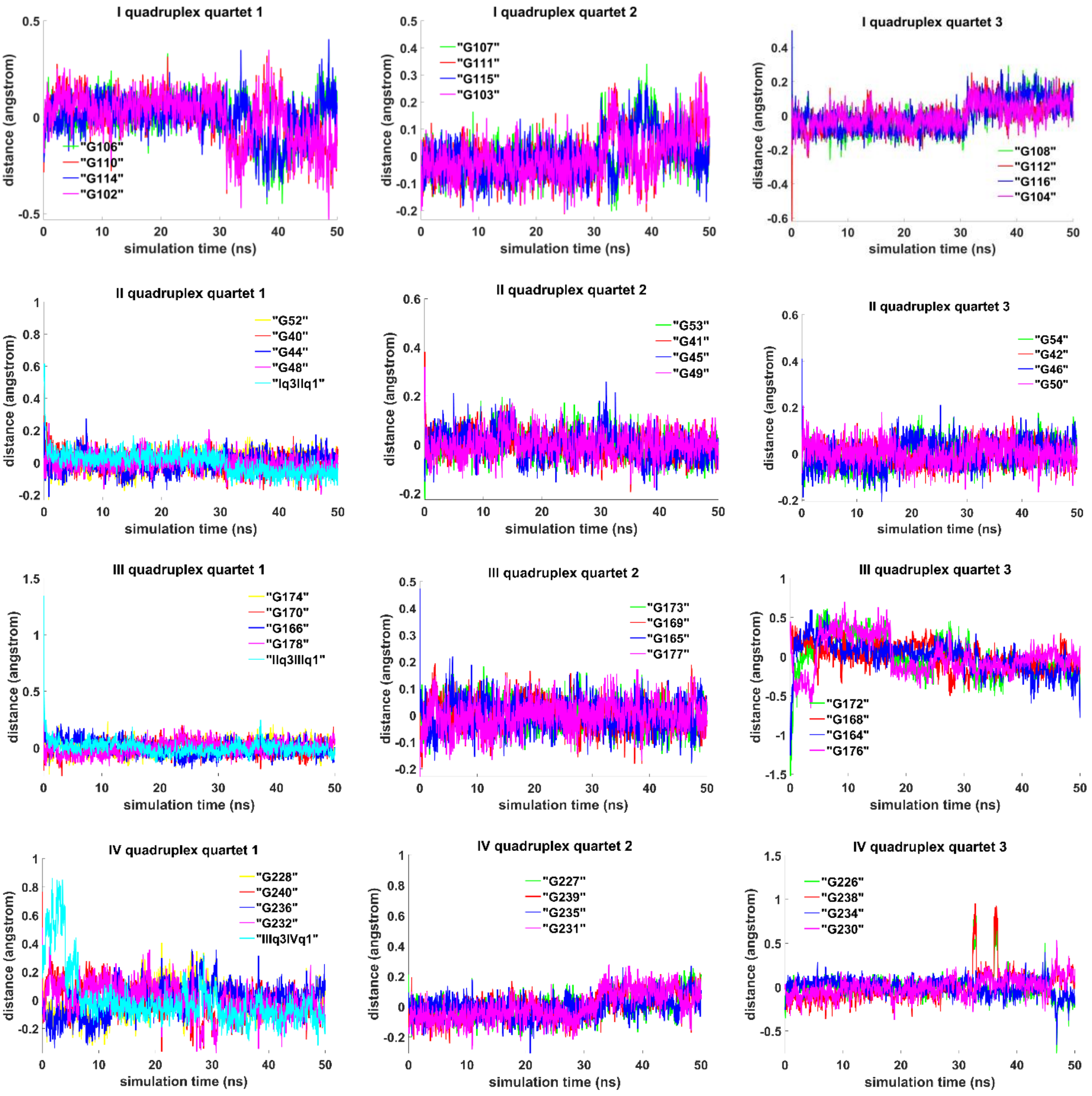


**Figure 4.3.S.3. Case 3: "Stacking of four parallel G4-dimers and four IM-monomers":** distances from COMs of the guanines' bases to COMs of their containing tetrad, distance between COMs of the boundary tetrads (**Iq3 IIq1, IIq3 IIIq1, IIIq3 IVq1**).

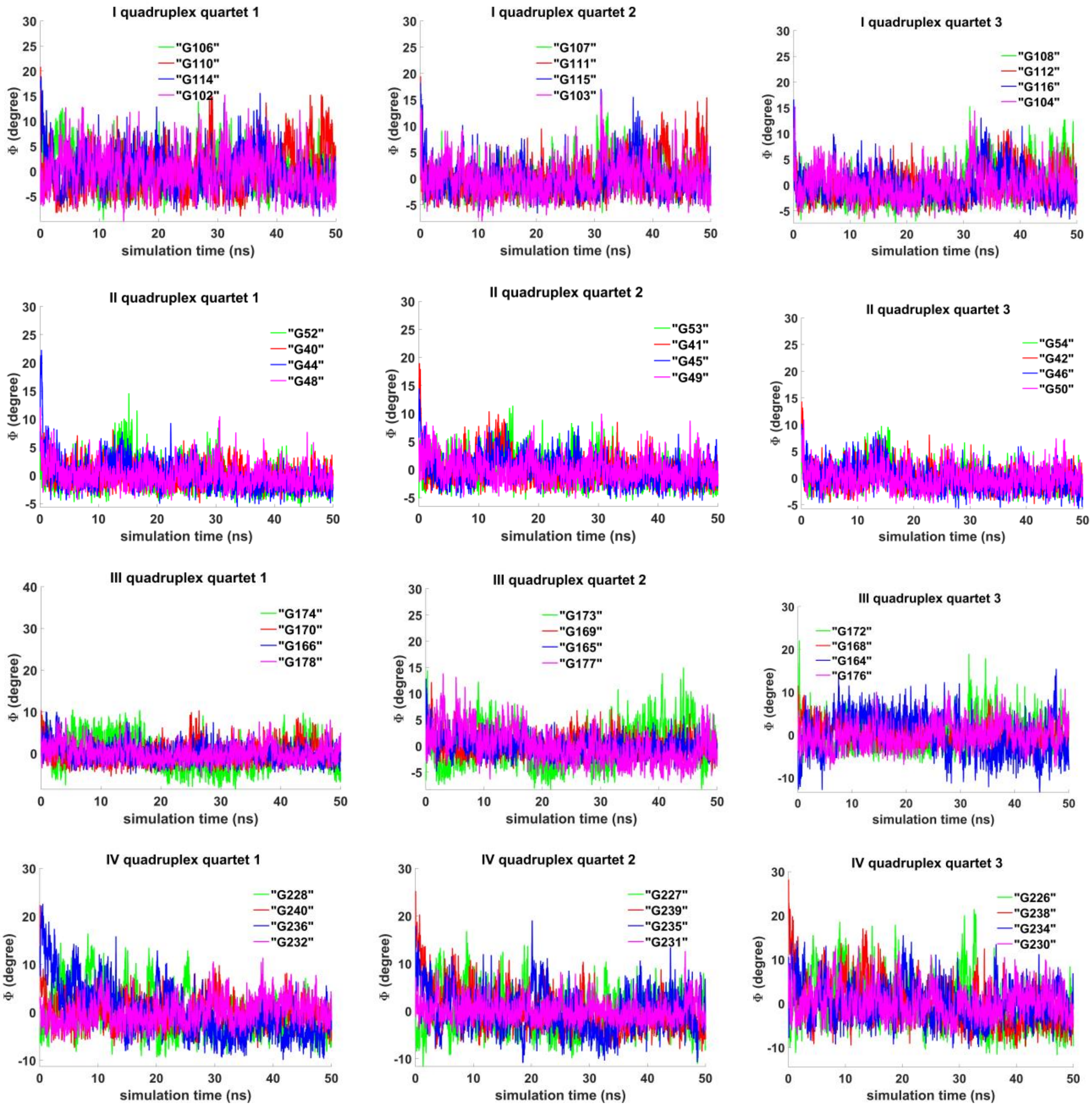


**Figure 4.3.S.4. Case 3: "Stacking of four parallel G4-dimers and four IM-monomers":** angles between normals to the guanine' bases and vectors connecting COMs of the boundary tetrads.

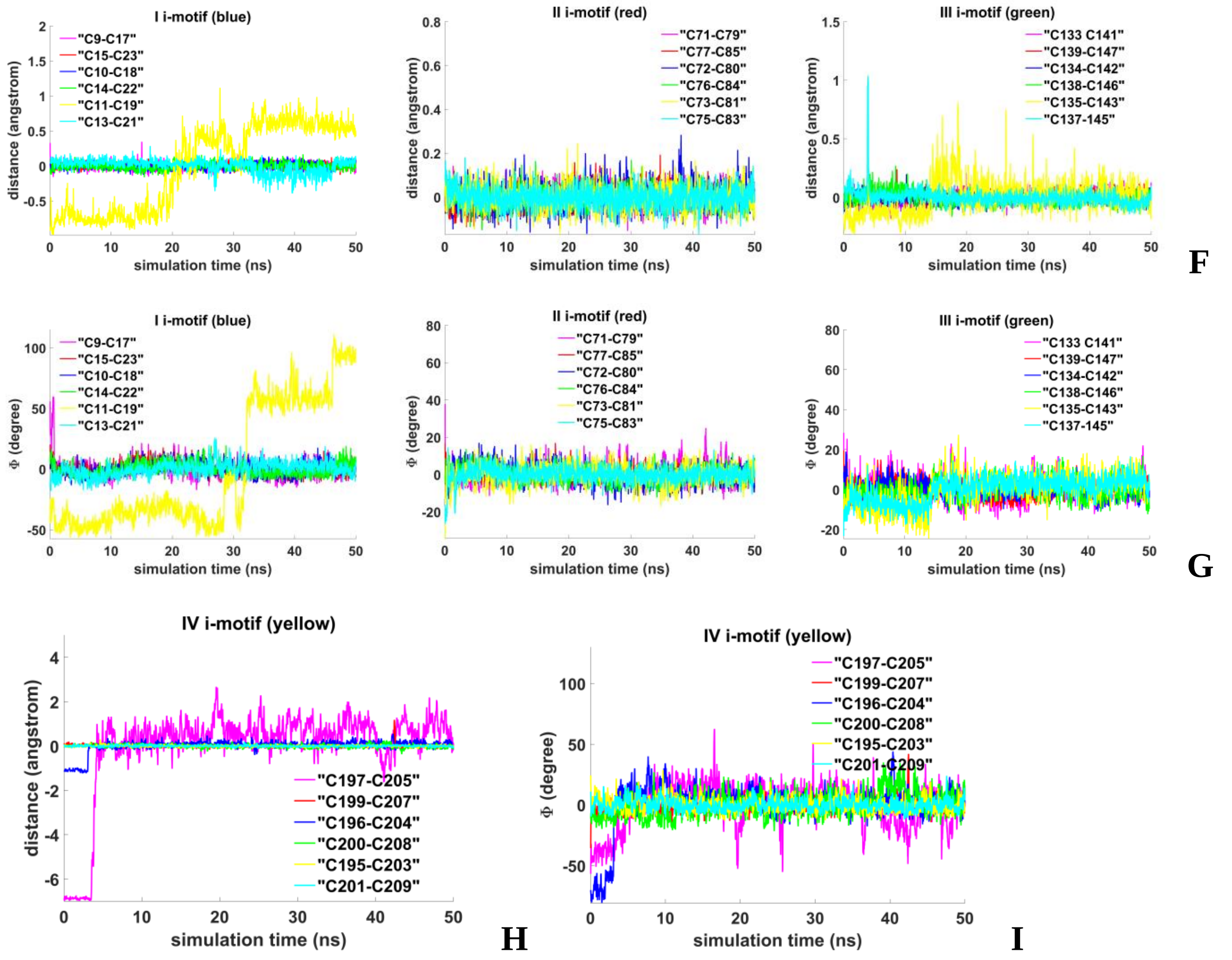


**Figure 4.3.S.5. Case 3: "Stacking of four parallel G4-dimers and four IM-monomers": F, H** - distances between COMs of the cytosines' bases; **G, I** - angles between normals to the cytosines' bases.

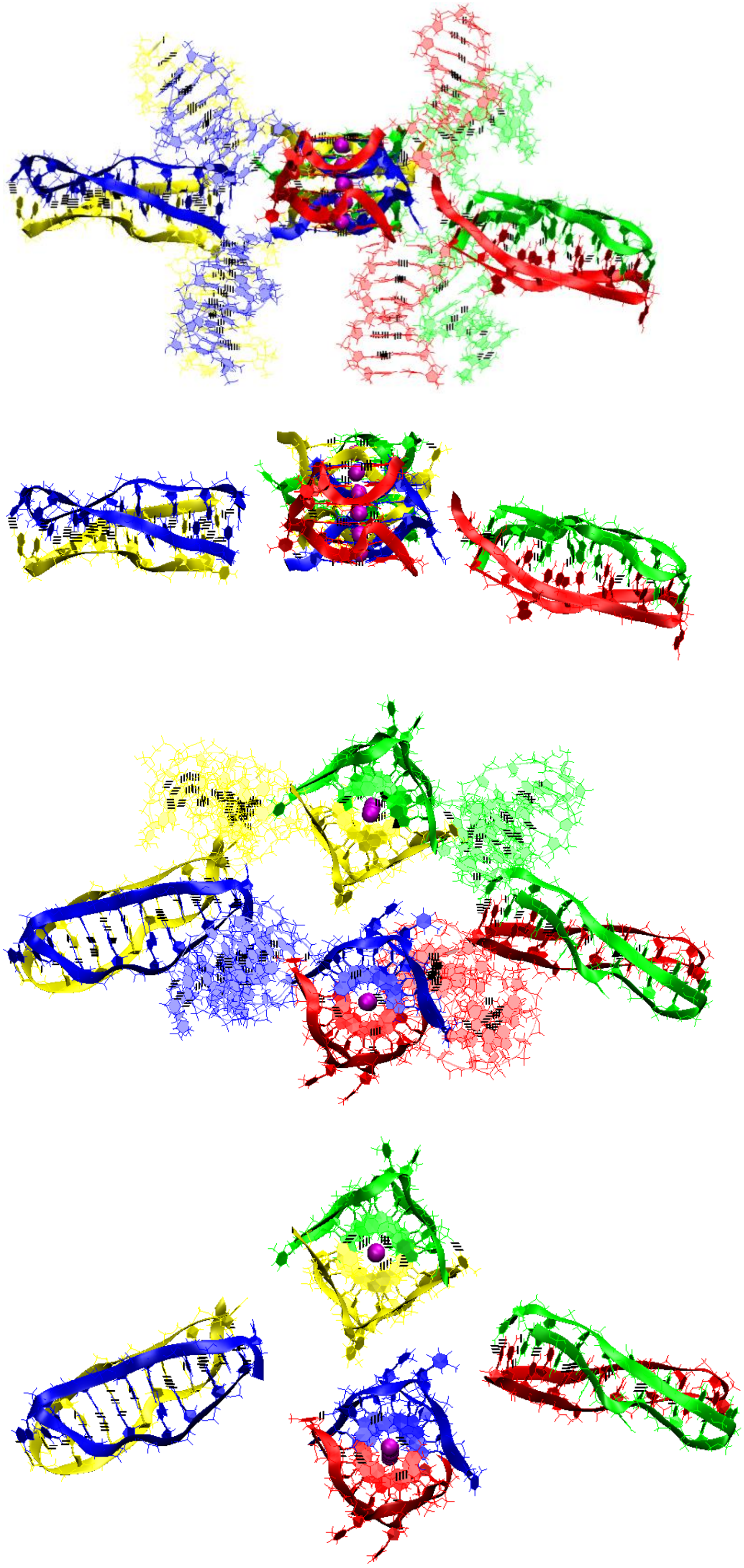

**A**

**Figure 4.4.S.1. Case 4: "Two parallel stack with right and left handed G4-dimers, and two head-to-head IM-dimers":** **A -** conformation obtained at the last step of the MD trajectory. The first row is the side view with unmelted fragments of the duplexes, the second row is the side view with only G4s and IMs, the third row is top view with unmelted fragments of the duplexes, and the fourth row is top view with only G4s and IMs.

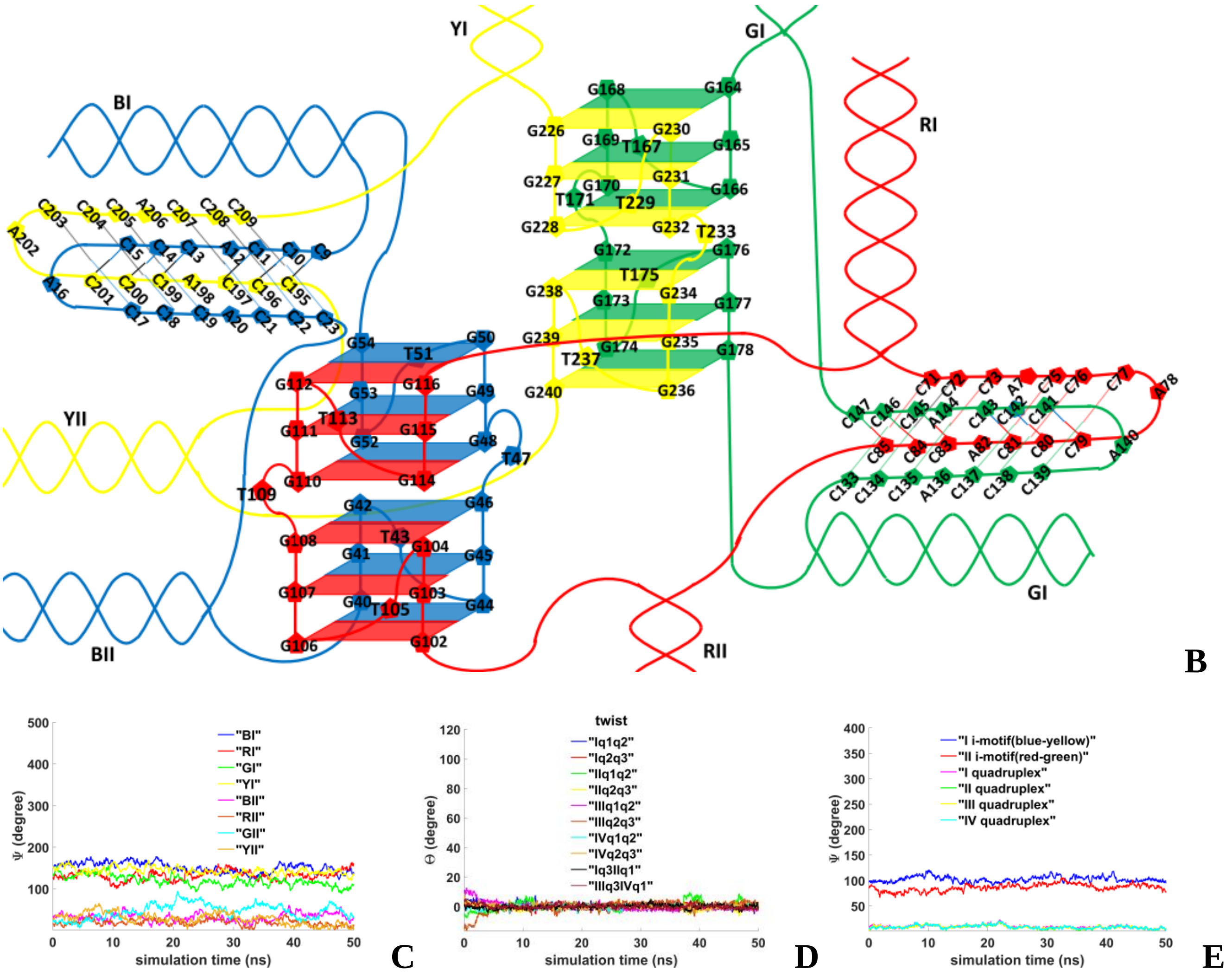


**Figure 4.4.S.2. Case 4: "Two parallel stack with right and left handed G4-dimers, and two head-to-head IM-dimers": B -** the complex scheme; **C -** angles between the straight lines, passing through the COMs of the first and the last complementary pairs unmelted fragments of duplexes, and the straight line passing through the COMs of boundary tetrads of the G4s; **D -** angles of rotation of the tetrads relative to each other; **E** - angles between the straight line, passing through the COM of all upper tetrads and the COM of all lower tetrads, and the straight lines, passing through the COMs of the upper and lower tetrads, in the G4s' case and straight lines, passing through the COMs of the boundary cytosine pairs, in the IMs' case.

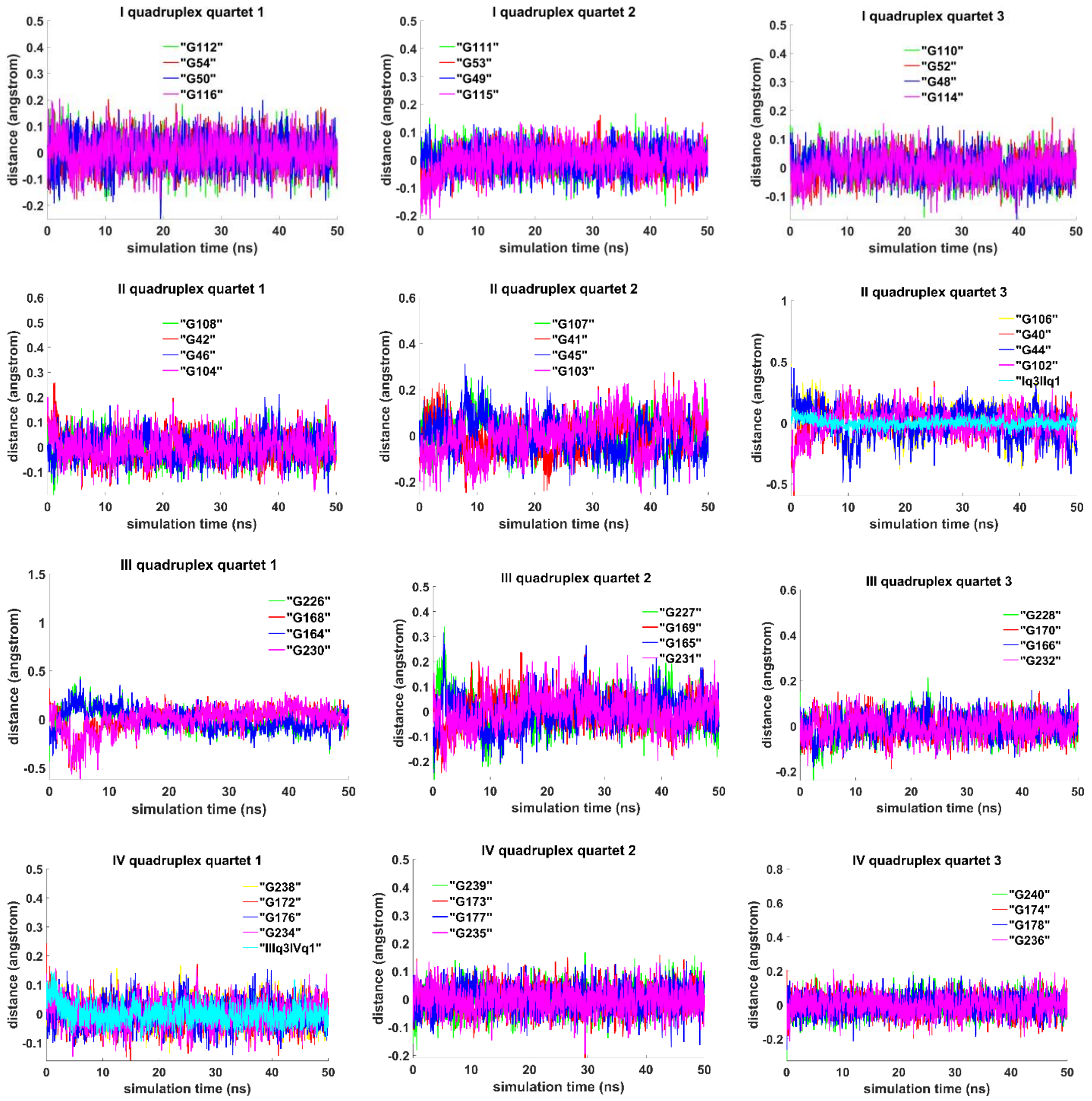


**Figure 4.4.S.3. Case 4: "Two parallel stack with right and left handed G4-dimers, and two head-to-head IM-dimers":** distances from COMs of the guanines' bases to COMs of their containing tetrad, distance between COMs of the boundary tetrads (**Iq3 IIq1, IIIq3 IVq1**).

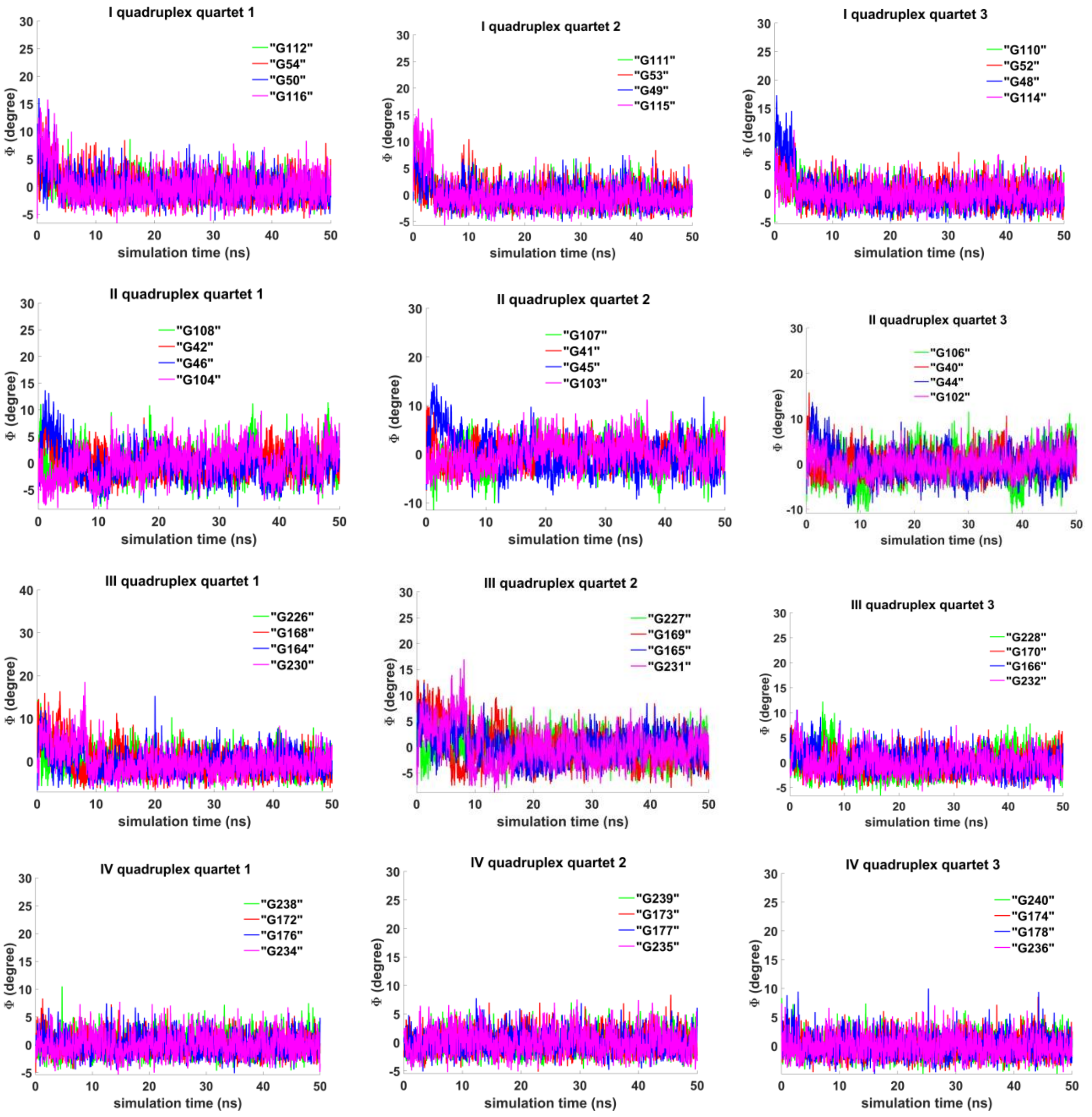


**Figure 4.4.S.4. Case 4: "Two parallel stack with right and left handed G4-dimers, and two head-to-head IM-dimers":** angles between normals to the guanine' bases and vectors connecting COMs of the boundary tetrads.

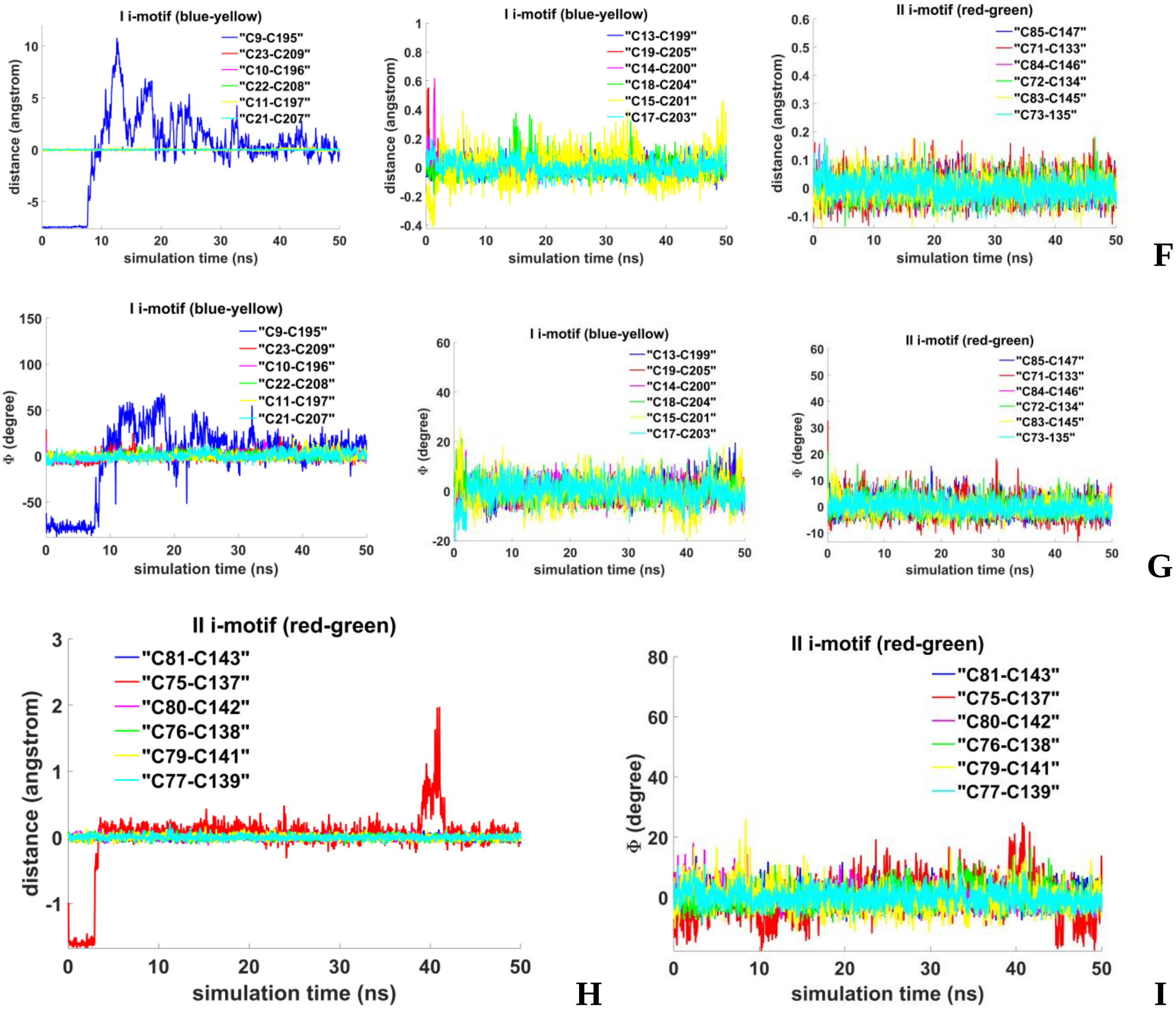


**Figure 4.4.S.5. Case 4: "Two parallel stack with right and left handed G4-dimers, and two head-to-head IM-dimers":** **F, H** - distances between COMs of the cytosines' bases; **G, I** - angles between normals to the cytosines' bases.

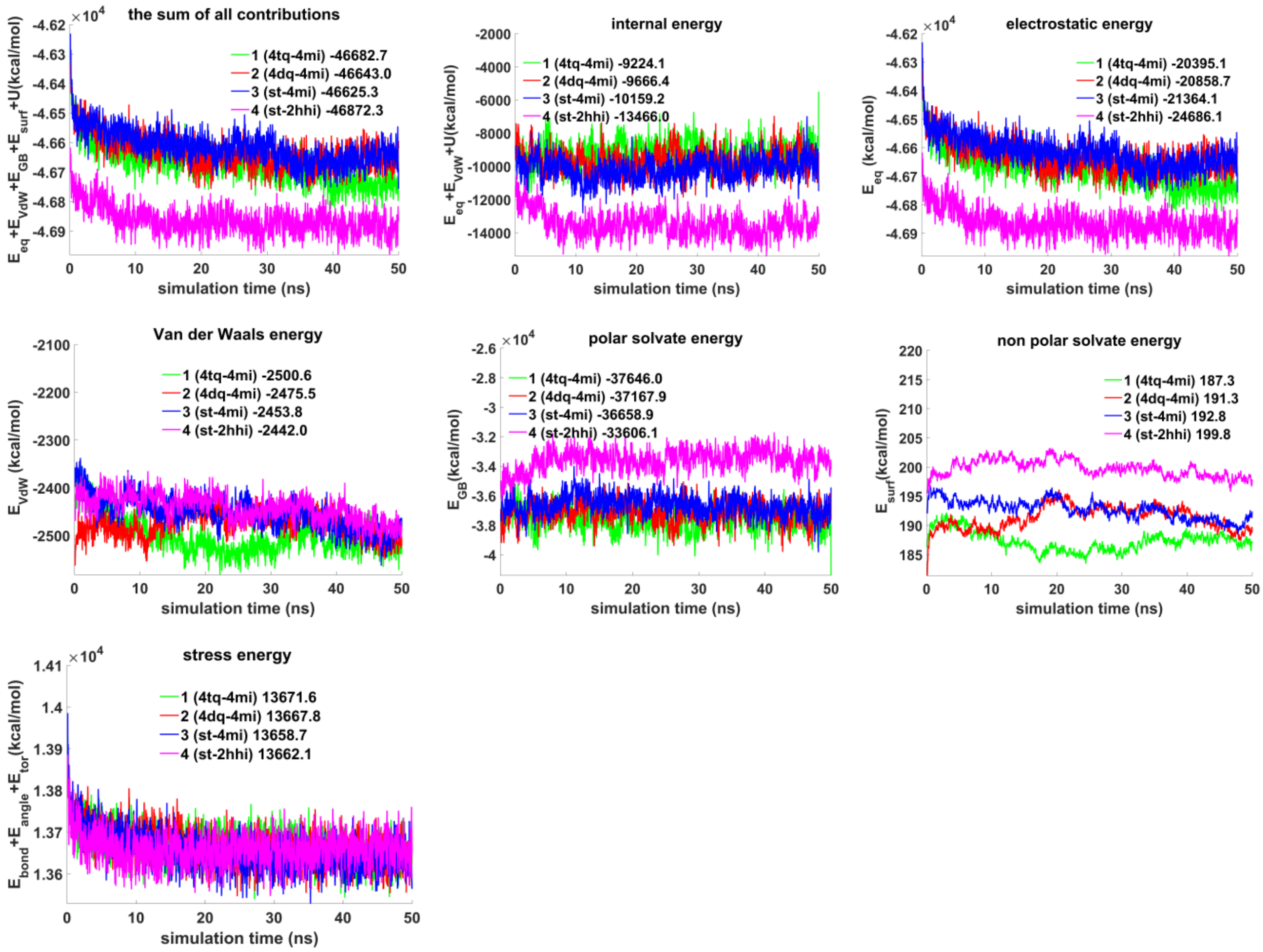


**Figure 4.S.E. The contributions to free energy during MD calculations for the variants of tetrameric complex of unmelted fragments of duplexes containing $(G_3T)_3G_3$ and $(C_3A)_3C_3$ fragments.** $E_{eq}$ – electrostatic, $E_{vdw}$ - Van der Waals, $E_{GB}$ - polar energy of solvation, $E_{surf}$ - non-polar energy of solvation due to the hydrophobic surface available to the solvent, $U = E_{bond} + E_{angle} + E_{tor}$ , e.g. $E_{bond}$, $E_{angle}$ and $E_{tor}$ –bond, angle and torsion stress energies. The energy plots were smoothed using moving average method (span = 5). Average energy values are indicated in the figure legends.

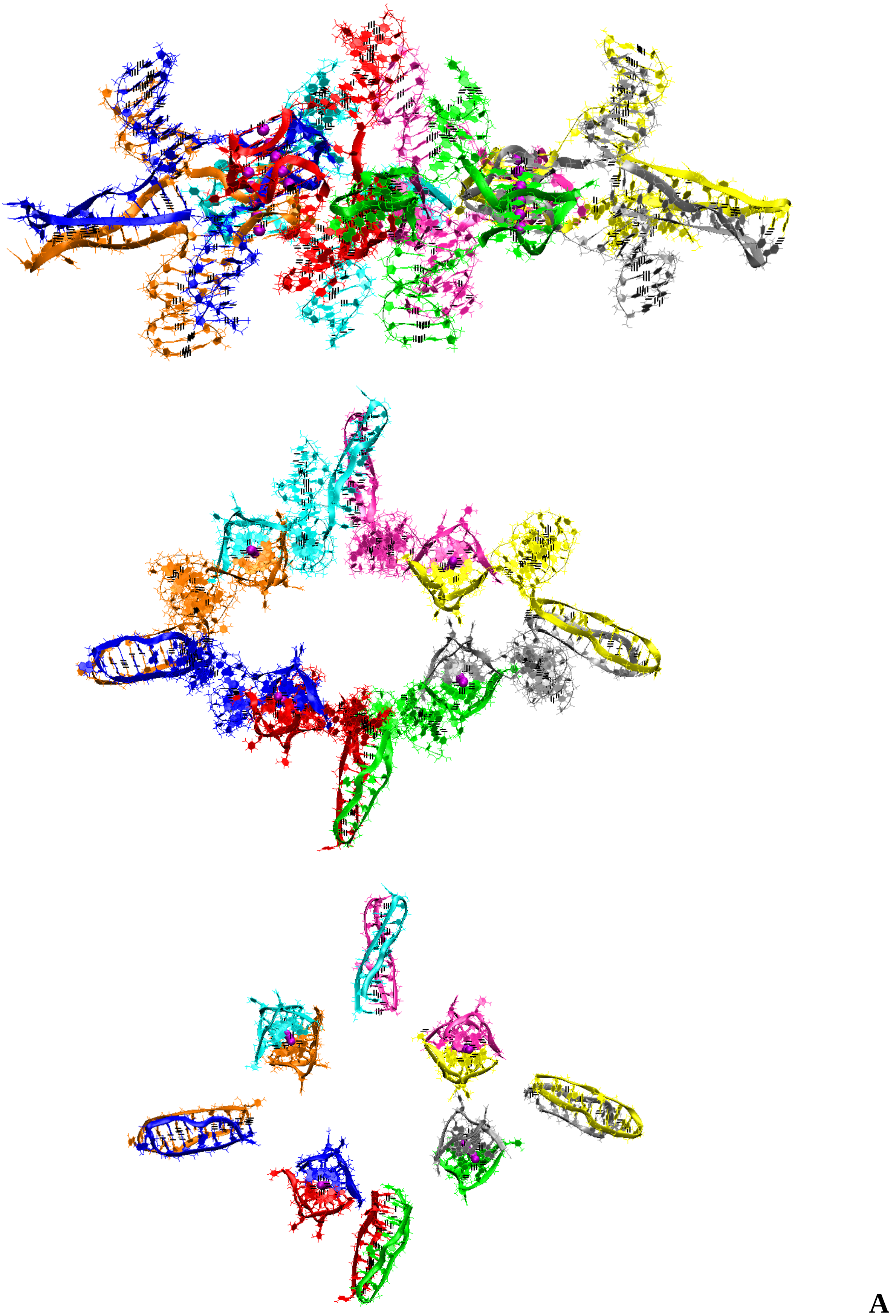


**Figure 5.S.1. Octameric complex with four parallel stack with right and left handed G4-dimers and four head-to-head IM-dimers: A** – conformation obtained at the last step of the MD trajectory (side view, top view, view with only G4s and IMs).

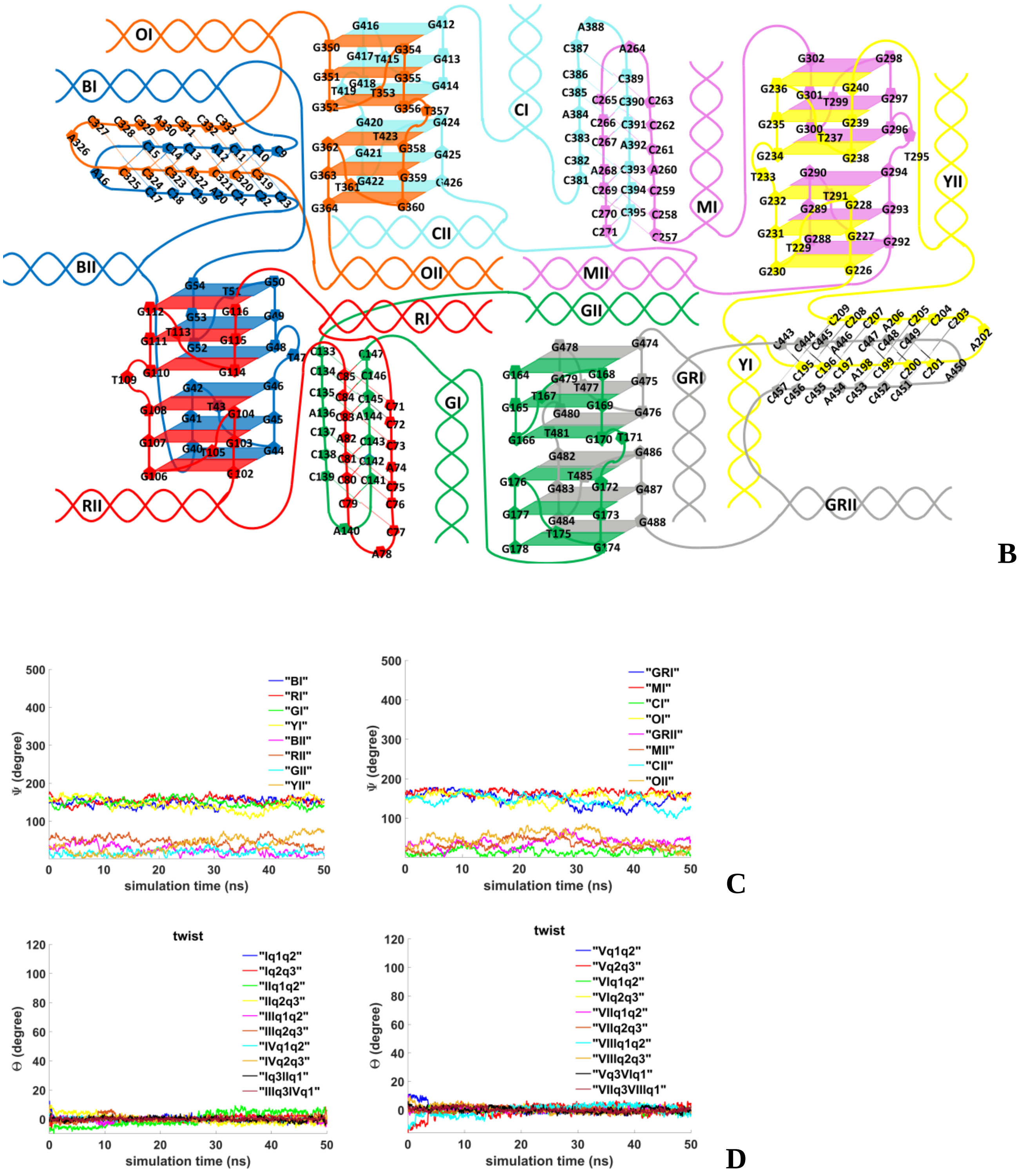


**Figure 5.S.2. Octameric complex with four parallel stack with right and left handed G4-dimers and four head-to-head IM-dimers: B** – the complex scheme; **C** – angles between the straight lines, passing through the COMs of the first and the last complementary pairs unmelted fragments of duplexes, and the straight line passing through the COMs of boundary tetrads of the G4s; **D**– angles of rotation of the tetrads relative to each other.

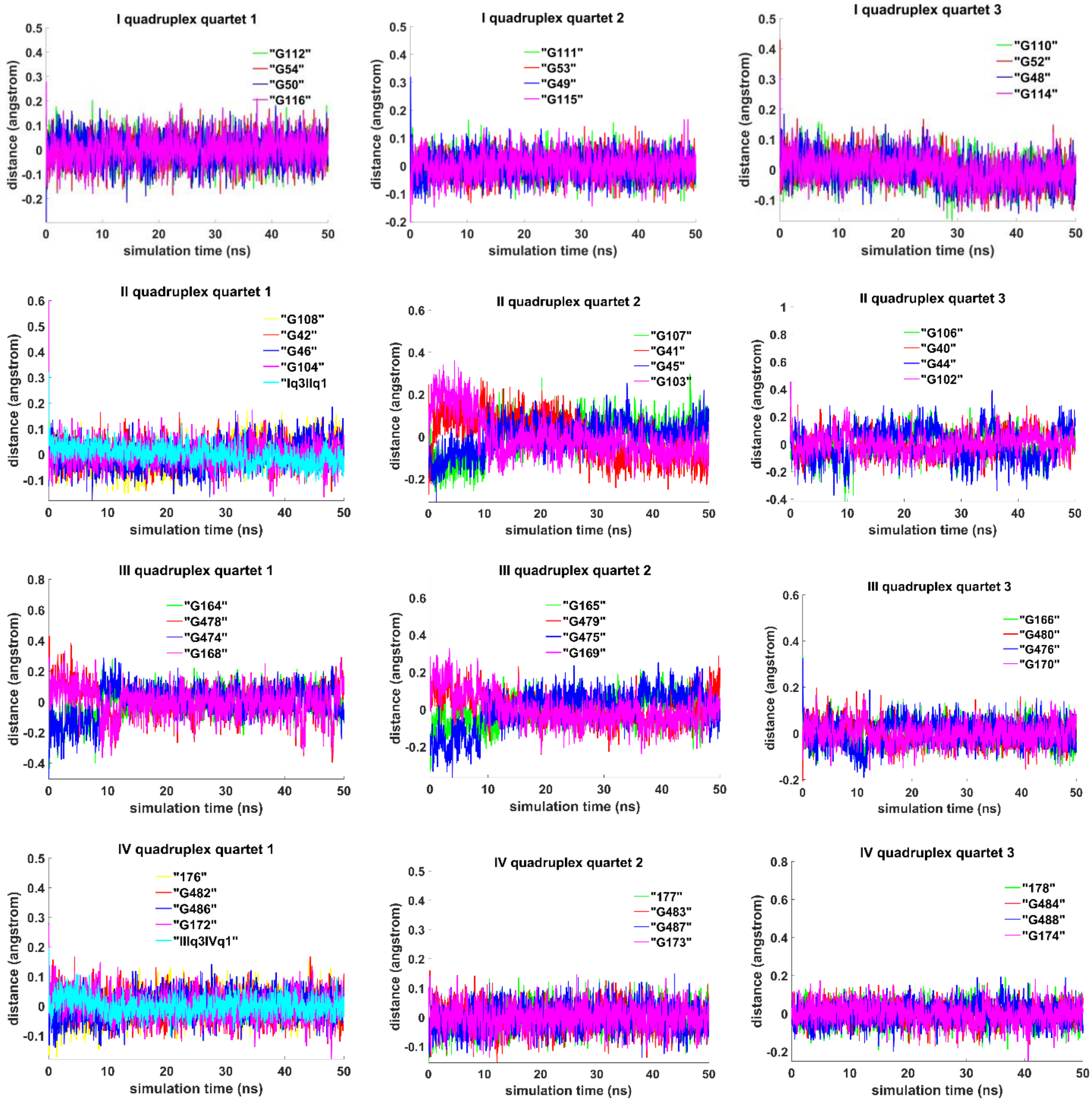


**Figure 5.S.3. Octameric complex with four parallel stack with right and left handed G4-dimers and four head-to-head IM-dimers: :** distances from COMs of the guanines' bases to COMs of their containing tetrad, distance between COMs of the boundary tetrads (**Iq3 IIq1, IIIq3 IVq1**).

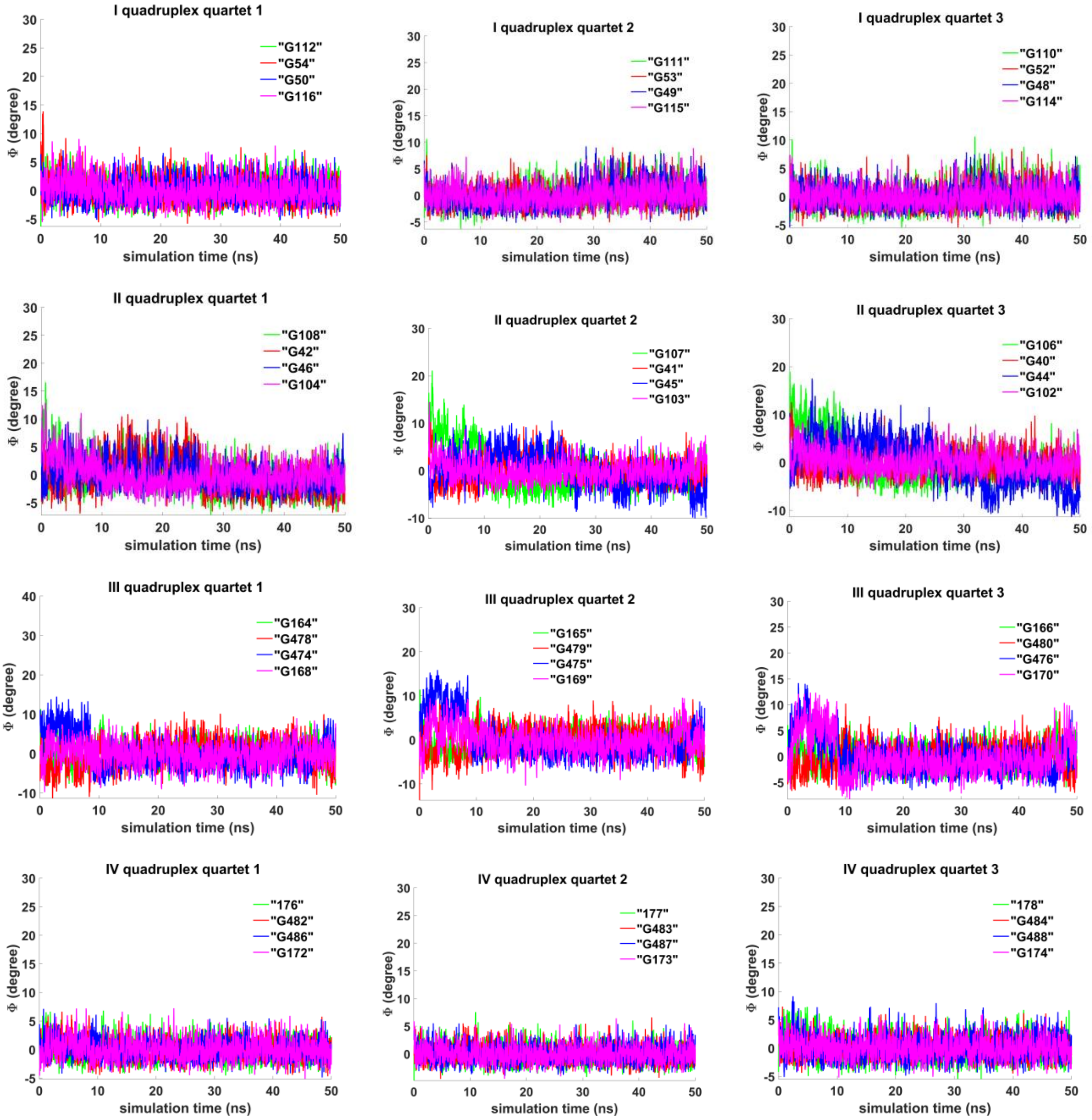


**Figure 5.S.4. Octameric complex with four parallel stack with right and left handed G4-dimers and four head-to-head IM-dimers: :** angles between normals to the guanines' bases and vectors connecting COMs of the boundary tetrads.

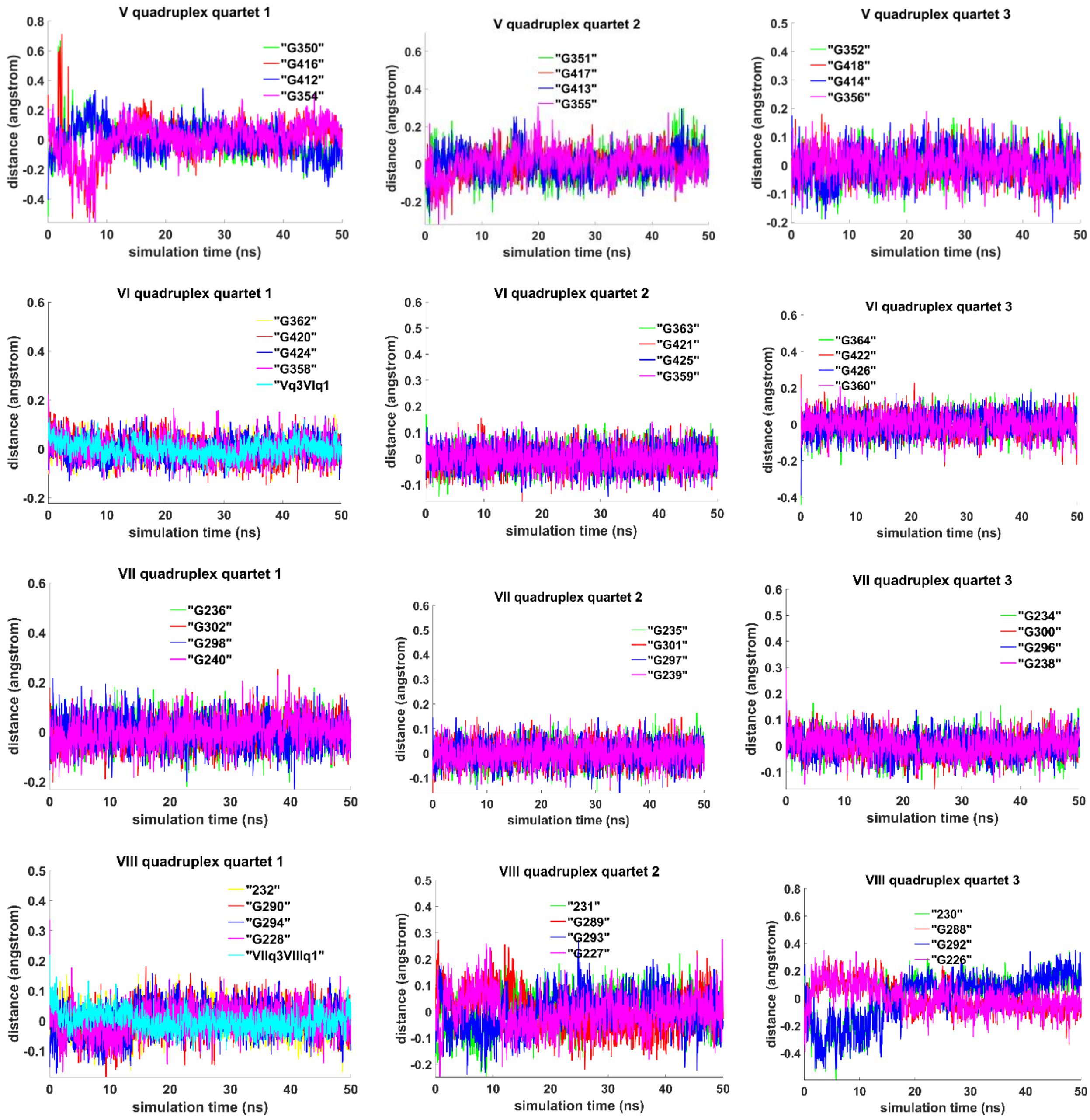


**Figure 5.S.5. Octameric complex with four parallel stack with right and left handed G4-dimers and four head-to-head IM-dimers:** distances from COMs of the guanines' bases to COMs of their containing tetrad, distance between COMs of the boundary tetrads (**Vq3 VIq1, VIIq3 VIIIq1**).

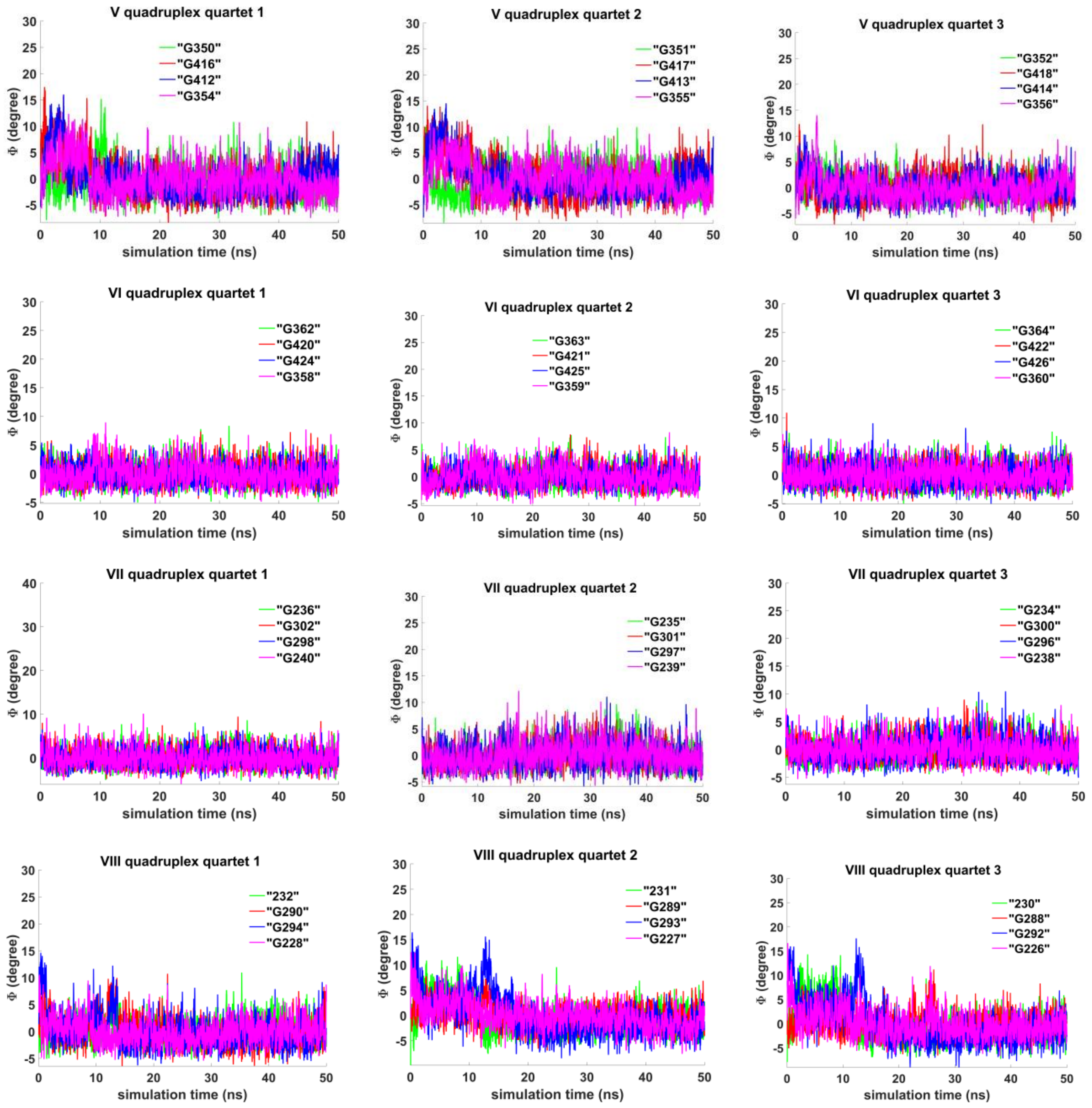


**Figure 5.S.6. Octameric complex with four parallel stack with right and left handed G4-dimers and four head-to-head IM-dimers: :** angles between normals to the guanines' bases and vectors connecting COMs of the boundary tetrads.

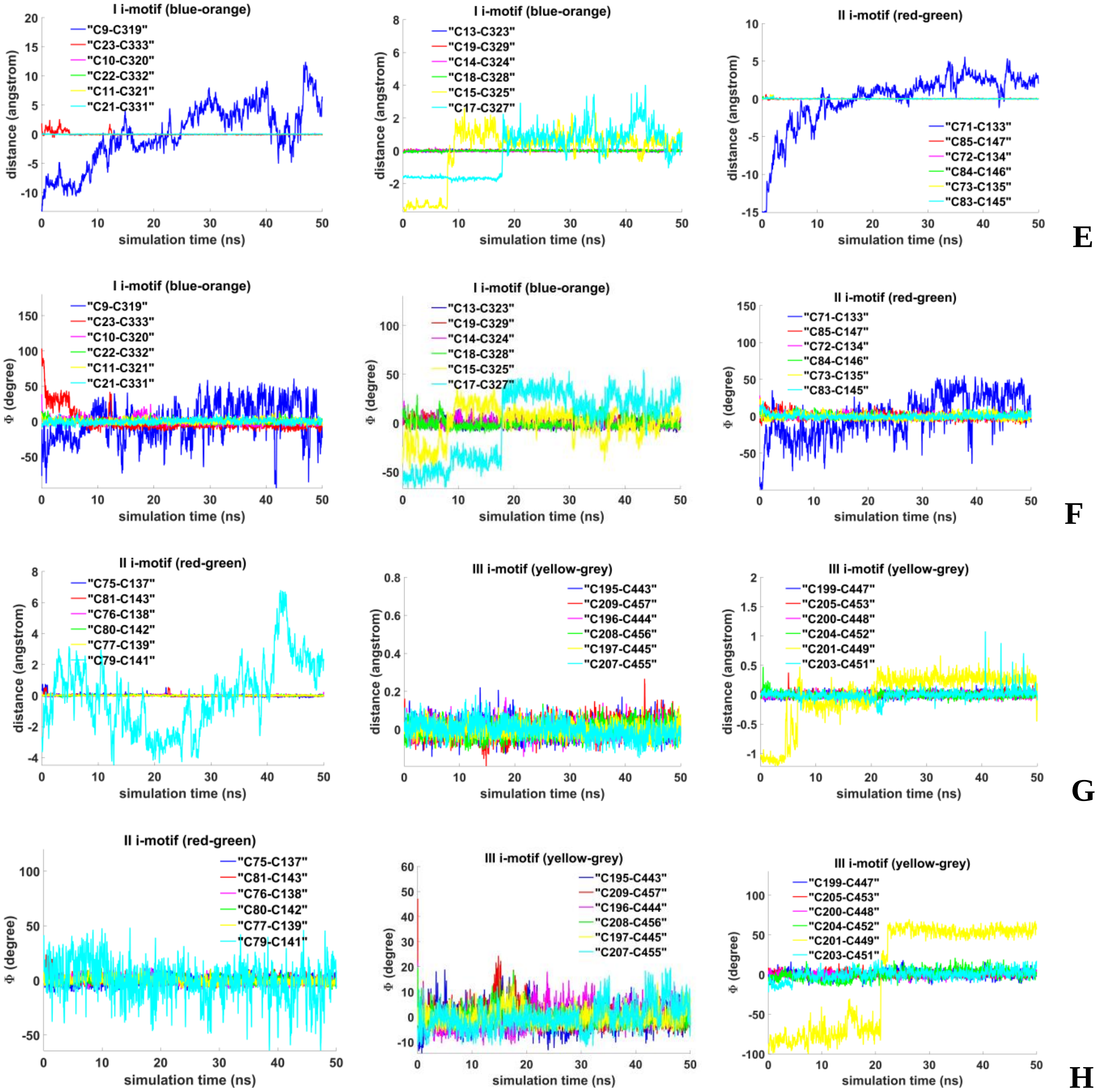


**Figure 5.S.7. Octameric complex with four parallel stack with right and left handed G4-dimers and four head-to-head IM-dimers: : E, G** - distances between COMs of the cytosines' bases; **F, H** - angles between normals to the cytosines' bases.

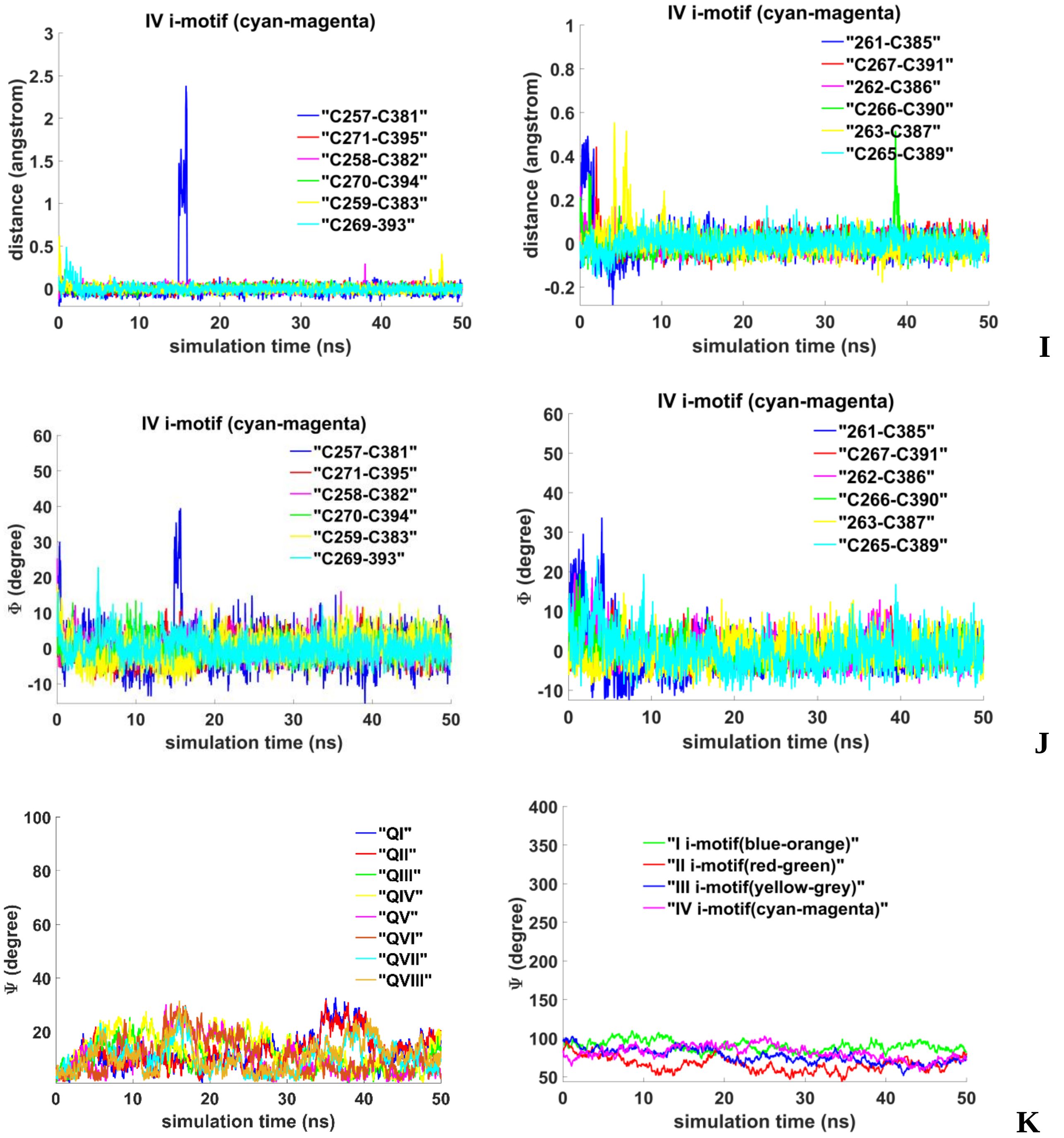


**Figure 5.S.8. Octameric complex with four parallel stack with right and left handed G4-dimers and four head-to-head IM-dimers: I** - distances between COMs of the cytosines' bases; **J** - angles between normals to the cytosines' bases; **K** - evolution of angle values between straight line, passing through COM of all upper tetrads and COM of all lower tetrads, and straight lines, passing through the COMs of the upper and lower quarters, in the case of G4s, and straight lines, passing through the COMs of the middle pairs of cytosines, in the case of the IMs.

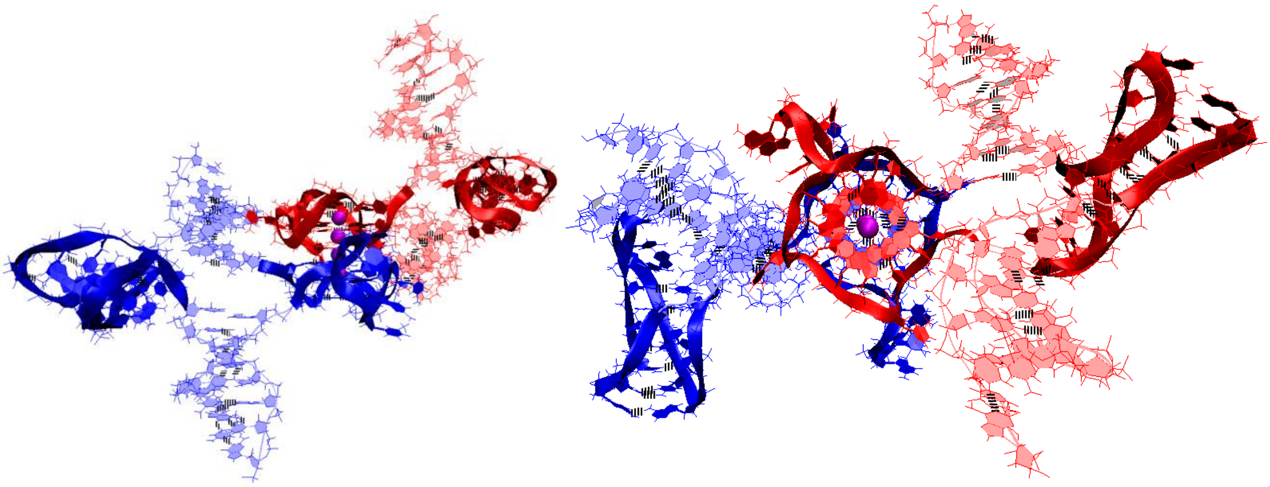

**A**

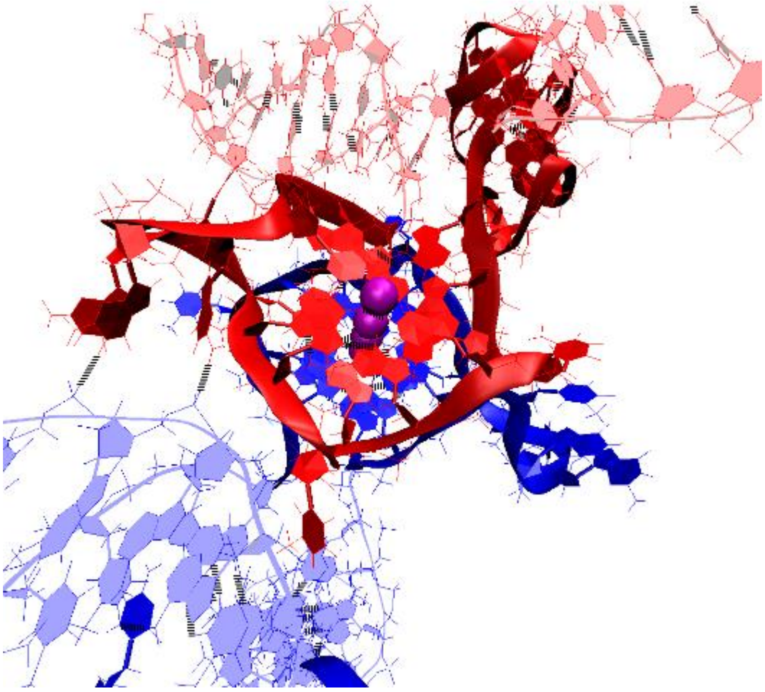

**B**

**Figure 6.S.1. Stacking and two IM-monomers**: **A, B** – conformation obtained at the last step of the MD trajectory (**A** - side and top view, **B** - enlarged view of quadruplexes).

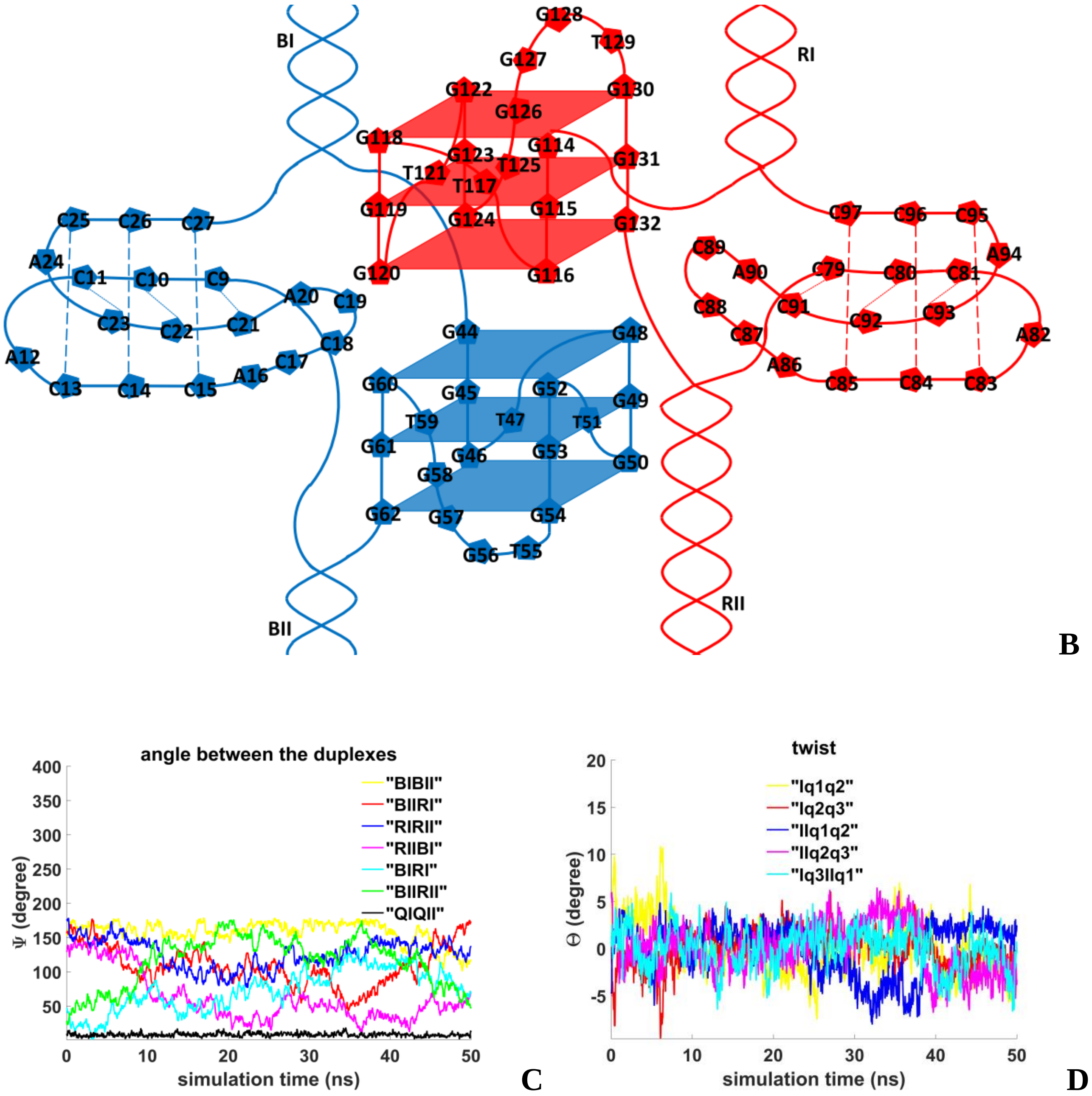


**Figure 6.S.2. Stacking and two IM-monomers**: **B** – the complex scheme; **C** – angles between unmelted fragments of the duplexes and axes passing through COMs of the tetrads, angle between axes passing through COMs of the boundary tetrads (**QIQII**); **D** – angles of rotation of the tetrads relative to each other.

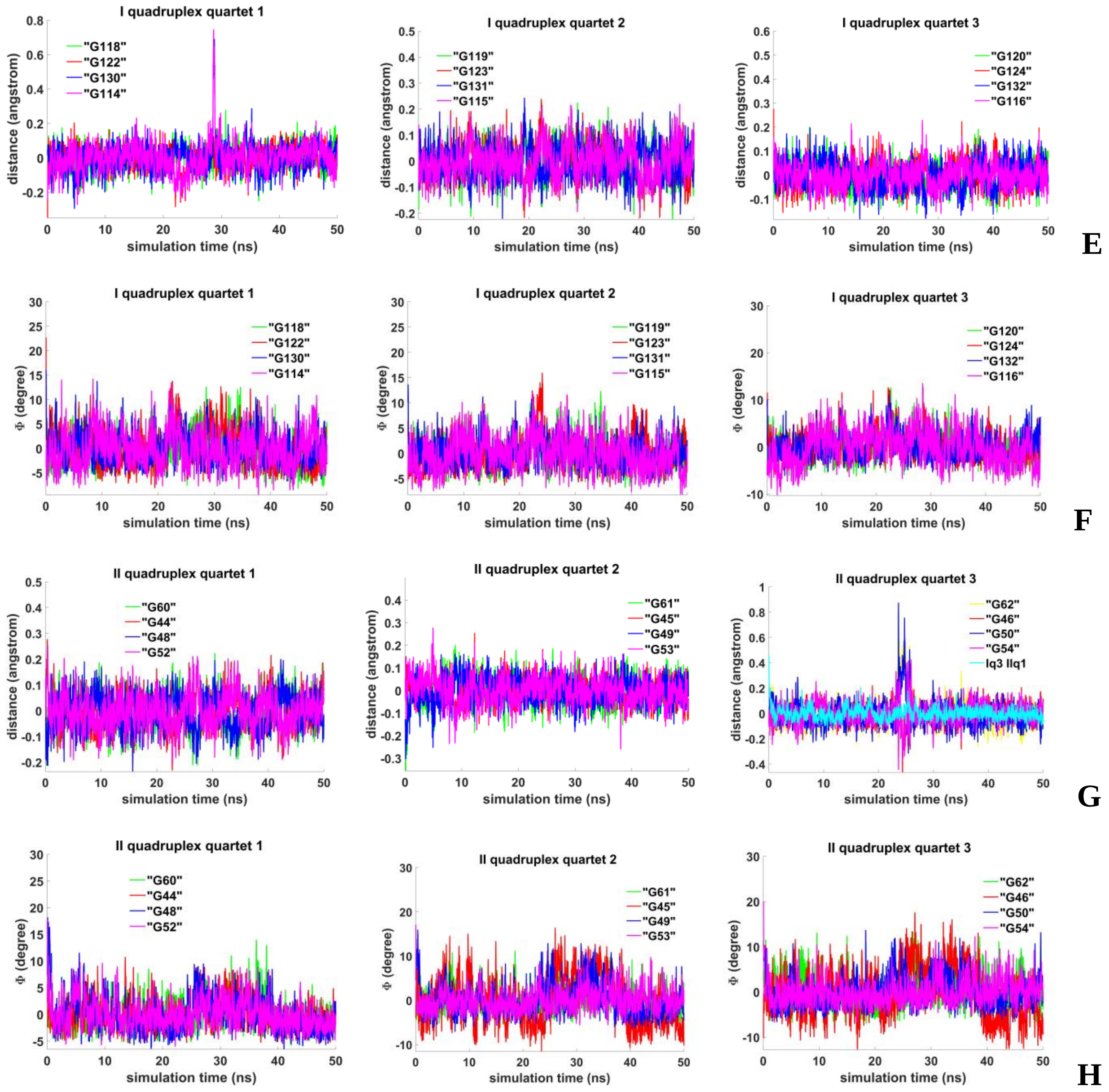


**Figure 6.S.3. Stacking and two IM-monomers**: **E, G** - distances from COMs of the guanines' bases to COMs of their containing tetrads, distance between COMs of the boundary tetrads (**Iq3 IIq1**); **F, H** - angles between normals to the guanines' bases and vectors connecting COMs of the boundary tetrads.

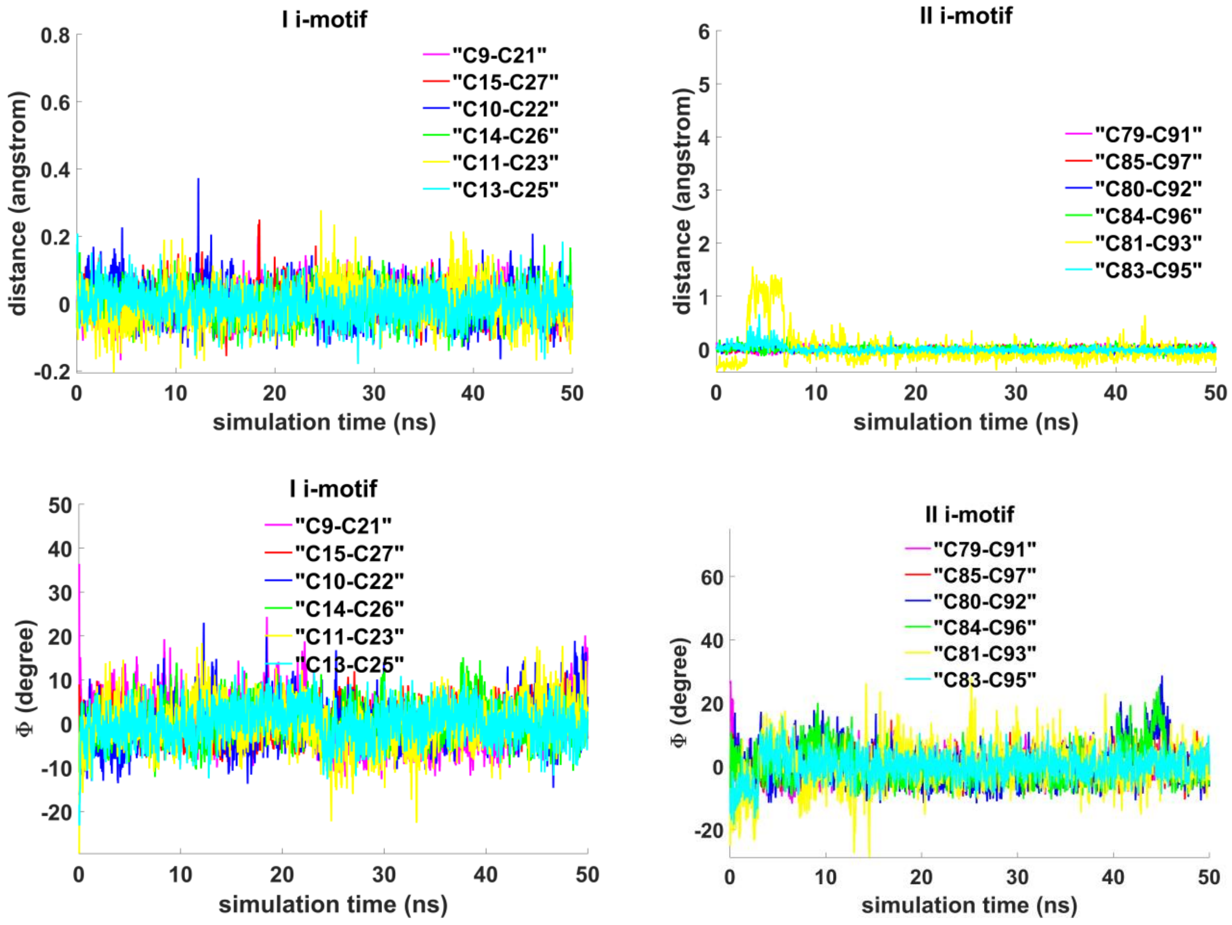


J

K

**Figure 6.S.4. Stacking and two IM-monomers**: **J -** distances between COMs of the cytosines' bases; **K** - angles between normals to the cytosines' bases.

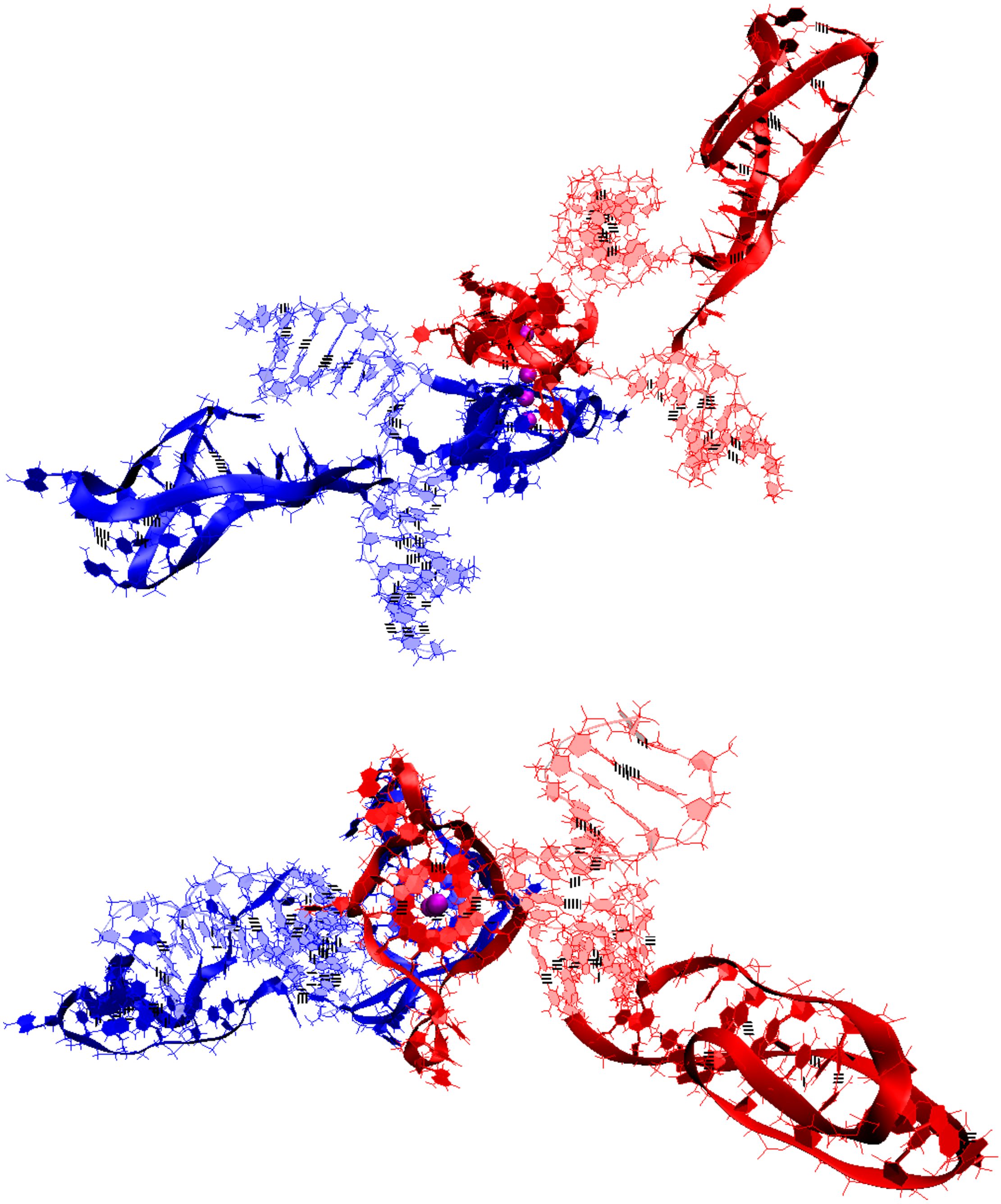

**A**

**Figure 7.1.A.S.1. Case 1: “Stacking and two IM-monomers”**: **A** – conformation obtained at the last step of the MD trajectory (side and top view);

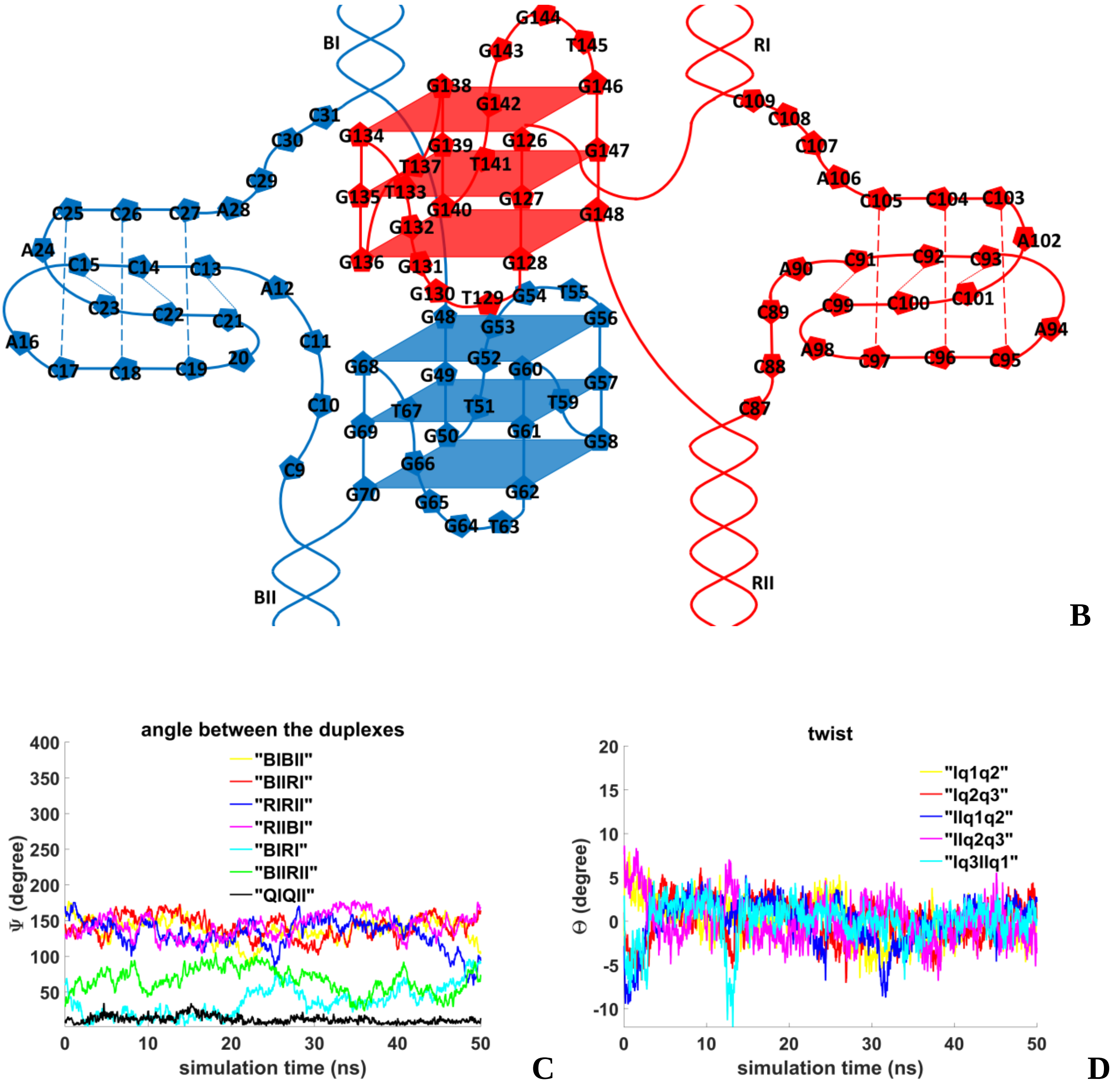


**Figure 7.1.A.S.2. Case 1: "Stacking and two IM-monomers"**: **B** – the complex scheme; **C** – angles between unmelted fragments of the duplexes and axes passing through COMs of the tetrads, angle between axes passing through COMs of the boundary tetrads (**QIQII**); **D**– angles of rotation of the tetrads relative to each other.

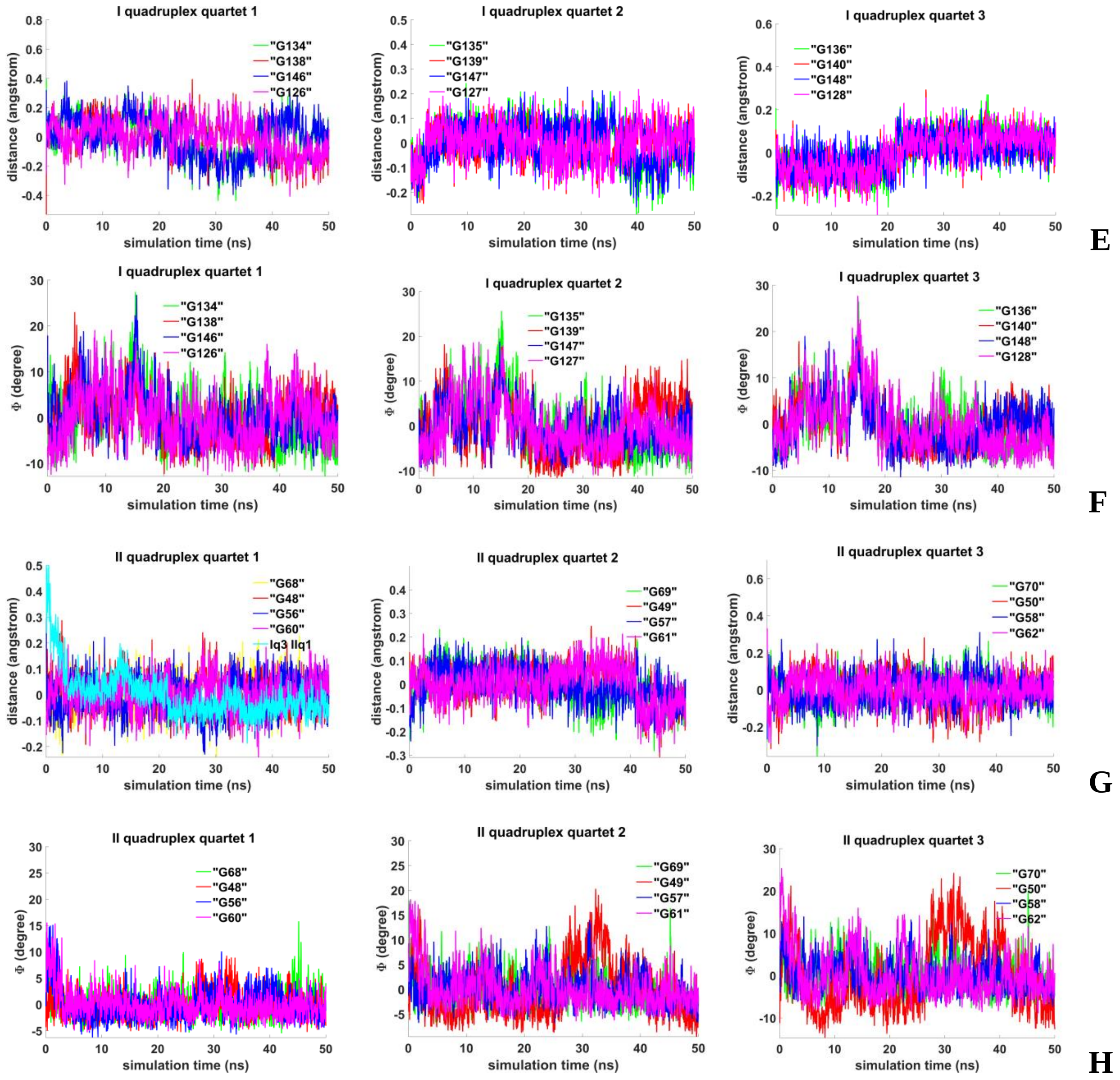


**Figure 7.1.A.S.3. Case 1: "Stacking and two IM-monomers: E, G -** distances from COMs of the guanines' bases to COMs of their containing tetrads, distance between COMs of the boundary tetrads **(Iq3 IIq1)**; **F, H** - angles between normals to the guanines' bases and vectors connecting COMs of the boundary tetrads.

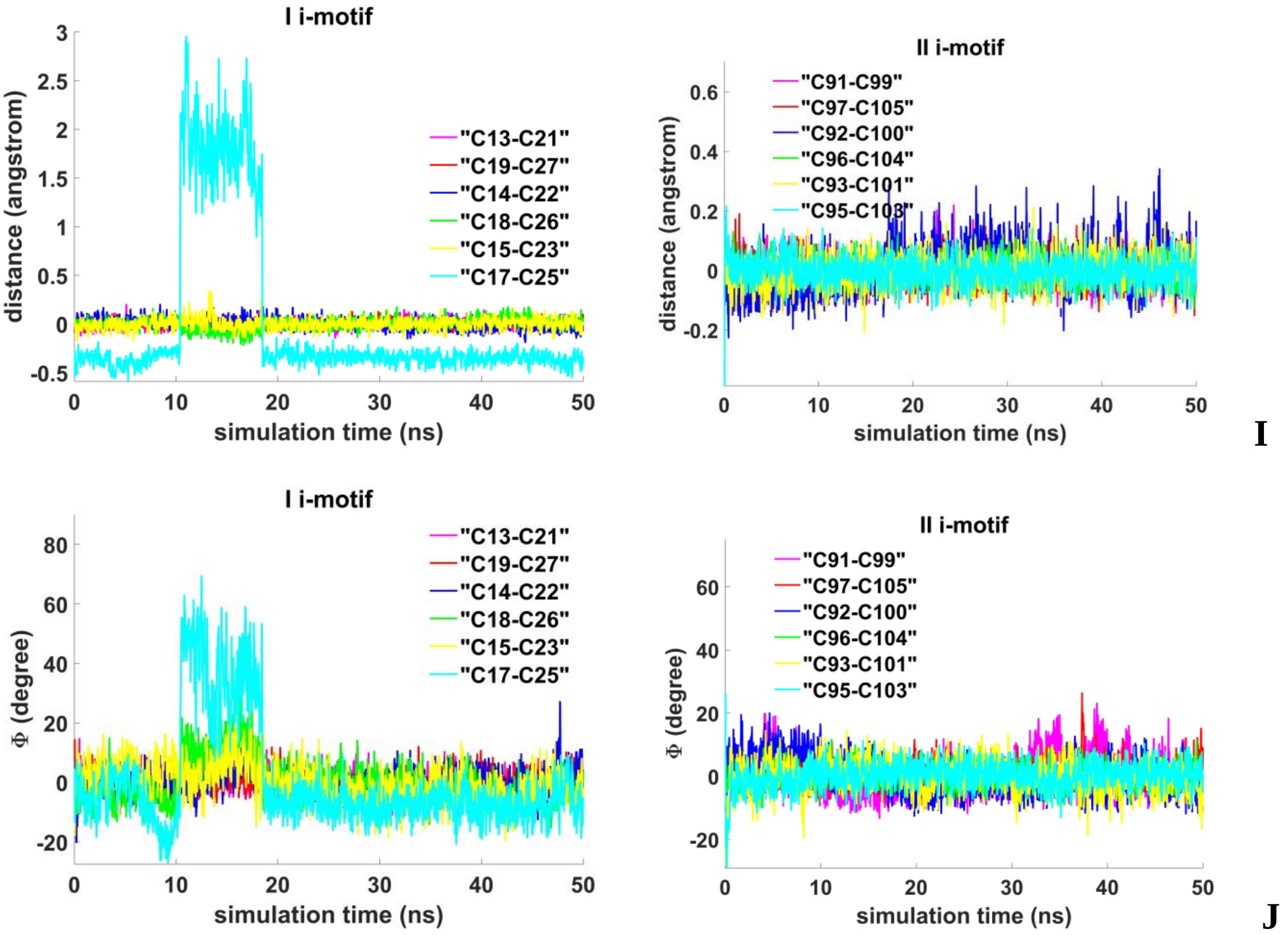


**Figure 7.1.A.S.4. Case 1: "Stacking and two IM-monomers"**: **I** - distances between COMs of the cytosines' bases; **J** - angles between normals to the cytosines' bases.

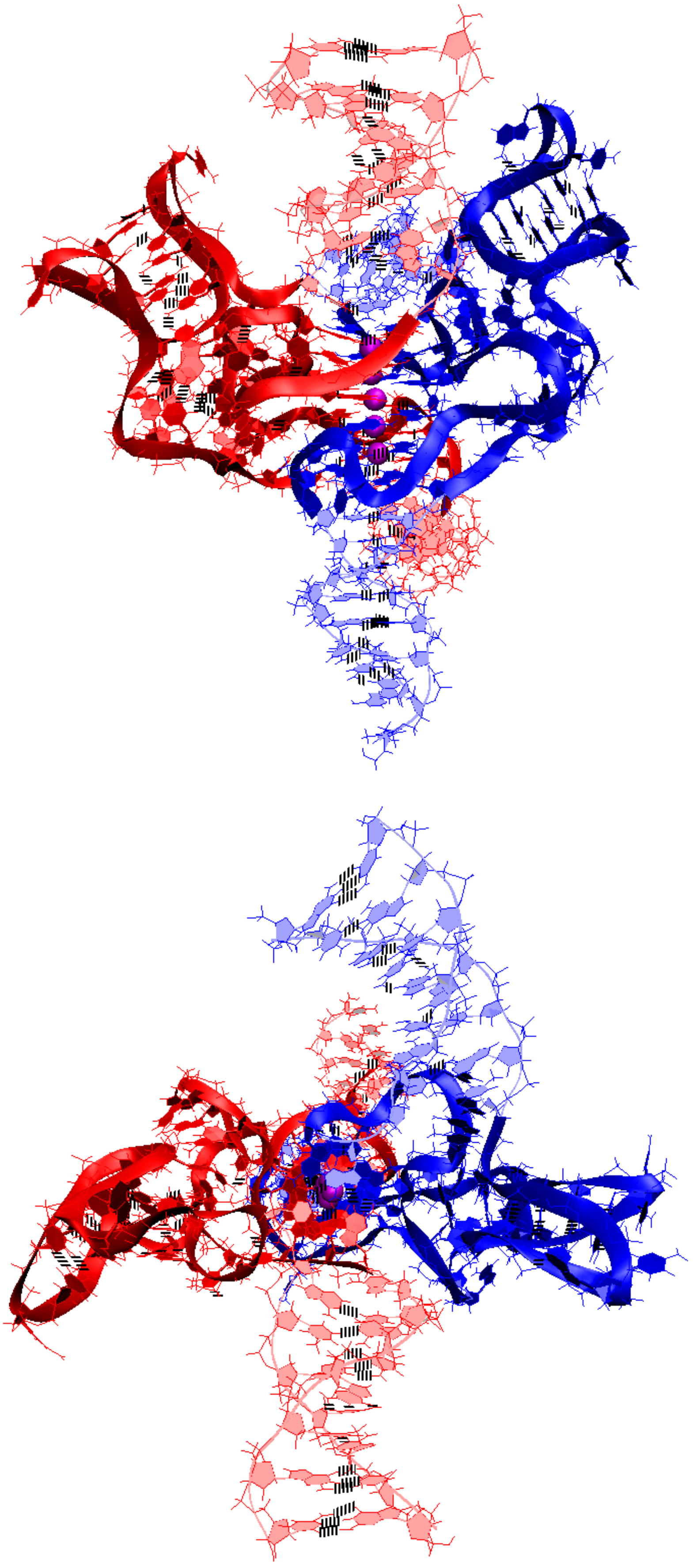

**A**

**Figure 7.1.B.S.1. Case 2: "1,2 hitch and two IM-monomers with two mini-duplex"**: **A** – conformations obtained at the last step of the MD trajectory (side and top view).

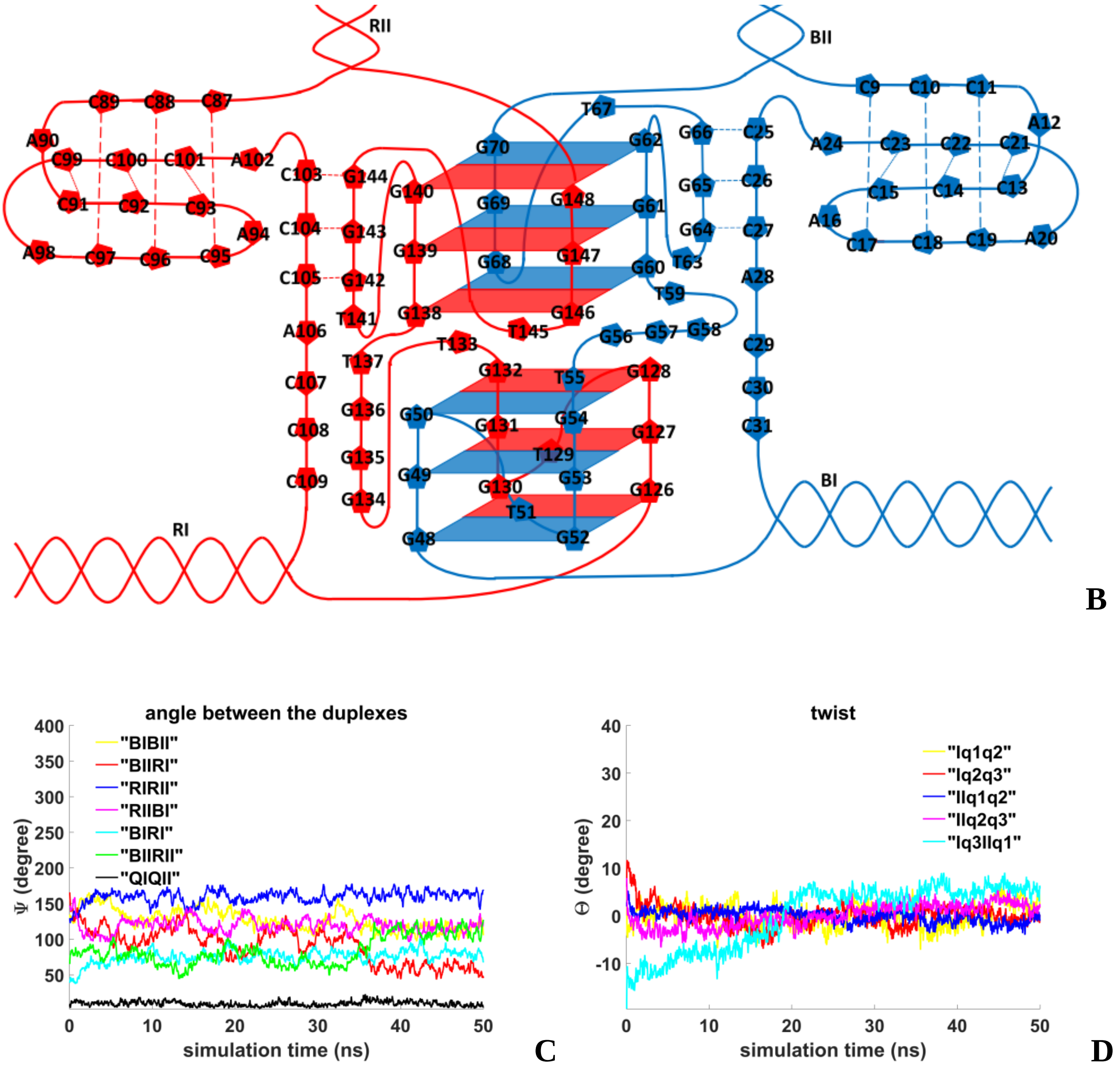


**Figure 7.1.B.S.2. Case 2: “1,2 hitch and two IM-monomers with two mini-duplex”**: **B** – the complex scheme; **C** – angles between unmelted fragments of the duplexes and axes passing through COMs of the tetrads, angle between axes passing through COMs of the boundary tetrads (**QIQII**); **D** – angles of rotation of the tetrads relative to each other.

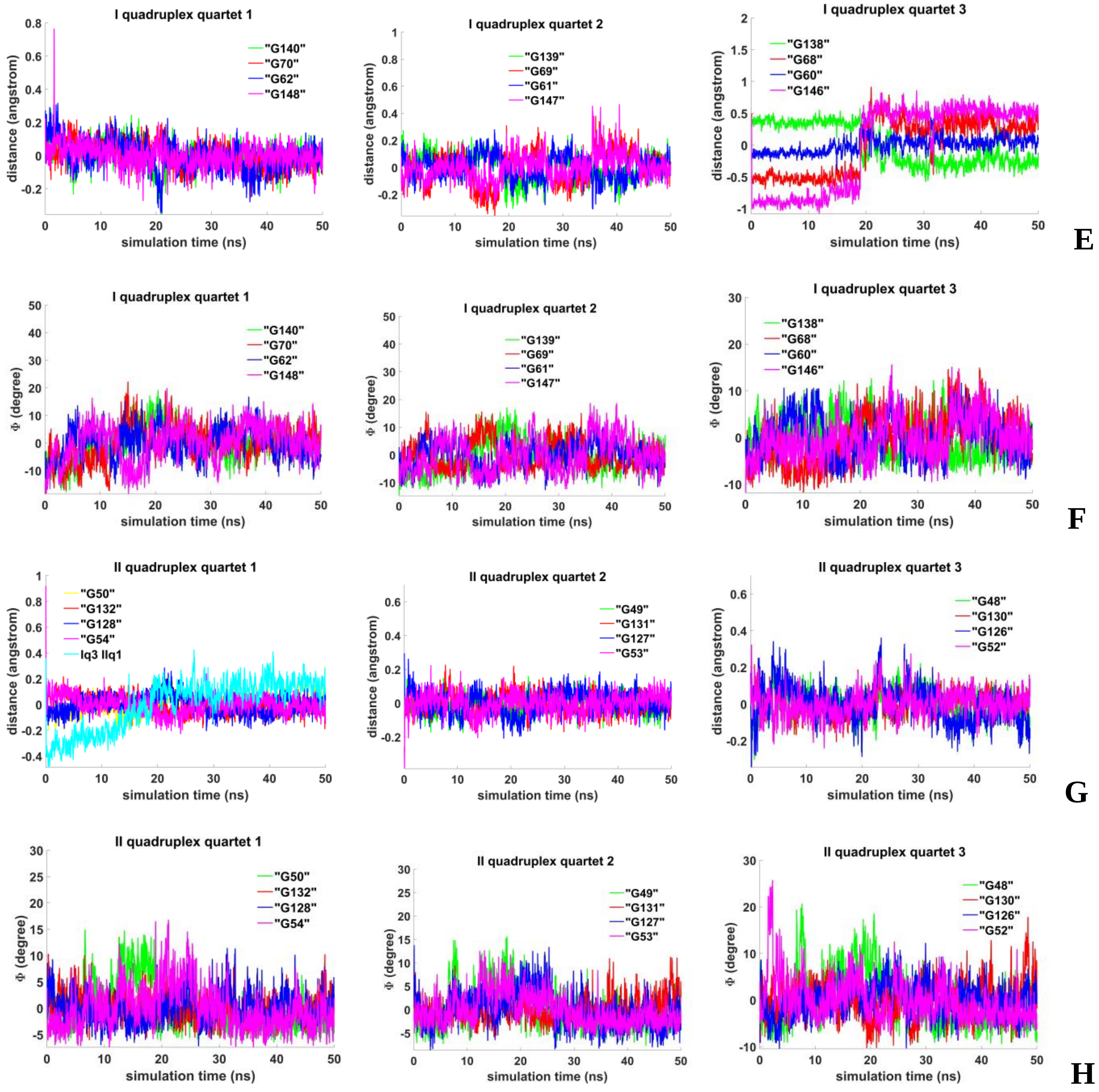


**Figure 7.1.B.S.3. Case 2: "1,2 hitch and two IM-monomers with two mini-duplex"**: **E, G -** distances from COMs of the guanines' bases to COMs of their containing tetrads, distance between COMs of the boundary tetrads (**Iq3 IIq1**); **F, H** - angles between normals to the guanines' bases and vectors connecting COMs of the boundary tetrads.

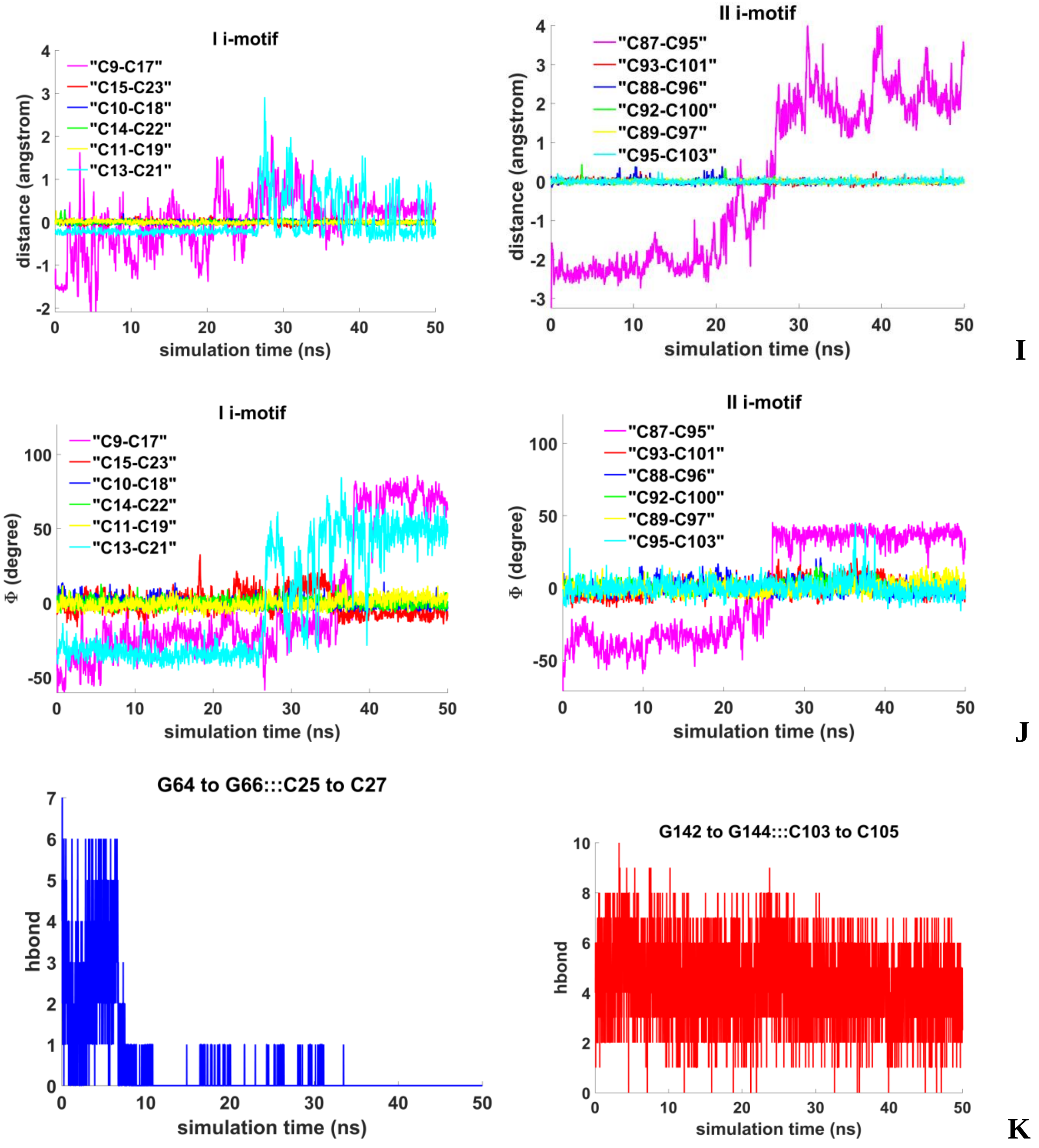


**Figure 7.1.B.S.4. Case 2: "1,2 hitch and two IM-monomers with two mini-duplexes"**: **I** - distances between COMs of the cytosines' bases; **J** - angles between normals to the cytosines' bases ; **K** - number of hydrogen bonds in mini-duplexes.

| donor | acceptor | occupancy | donor | acceptor | occupancy |
|---|---|---|---|---|---|
| DC103-Side-N4 | DG144-Side-O6 | 23,75% | DG64-Side-N1 | DC27-Side-N3 | 0,05% |
| DC104-Side-N4 | DG143-Side-O6 | 54,58% | DG64-Side-N2 | DC27-Side-O2 | 0,06% |
| DC105-Side-N4 | DG142-Side-O6 | 37,31% | DG65-Side-N1 | DC26-Side-N3 | 4,71% |
| DG144-Side-N2 | DC103-Side-O2 | 18,63% | DG66-Side-N1 | DC25-Side-N3 | 4,42% |
| DG144-Side-N1 | DC103-Side-N3 | 23,18% | DG65-Side-N2 | DC26-Side-O2 | 3,84% |
| DG143-Side-N1 | DC104-Side-N3 | 48,17% | DG66-Side-N2 | DC25-Side-O2 | 3,41% |
| DG143-Side-N2 | DC104-Side-O2 | 62,06% | DC27-Side-N4 | DG64-Side-O6 | 0,26% |
| DG142-Side-N1 | DC105-Side-N3 | 61,81% | DC26-Side-N4 | DG65-Side-O6 | 2,56% |
| DG142-Side-N2 | DC105-Side-O2 | 62,23% | DC25-Side-N4 | DG66-Side-O6 | 4,09% |
| DG144-Side-N2 | DC105-Side-O4' | 33,66% | DG66-Side-N2 | DC26-Side-O2 | 0,04% |
| DG142-Side-N1 | DC105-Side-O2 | 0,12% | DC26-Side-N4 | DG64-Side-O6 | 0,30% |
| DG144-Side-N2 | DC104-Side-O2 | 0,47% | DG66-Side-N1 | DC25-Side-O2 | 0,09% |
| DG142-Side-N2 | DC105-Side-N3 | 0,10% | DG65-Side-N1 | DC26-Side-O2 | 0,09% |
| DG143-Side-N2 | DC105-Side-O2 | 0,06% | DC25-Side-N4 | DG65-Side-O6 | 1,09% |
| DG144-Side-N1 | DC103-Side-O2 | 0,01% | DG65-Side-N2 | DC26-Side-N3 | 0,02% |
| DG144-Side-N2 | DC104-Side-O4' | 0,10% | DC26-Side-N4 | DG64-Side-N7 | 0,07% |
| | | | DG66-Side-N2 | DC25-Side-N3 | 0,07% |
| | | | DG66-Side-N2 | DC26-Side-N3 | 0,02% |
| | | | DC26-Side-N4 | DG66-Side-O6 | 0,29% |

**Table .1(Appendix to Figure 7.1.B.S.4. Case 2)** Percentage of snapshots with hydrogen bonds generated by pairs in the mini-duplexes.

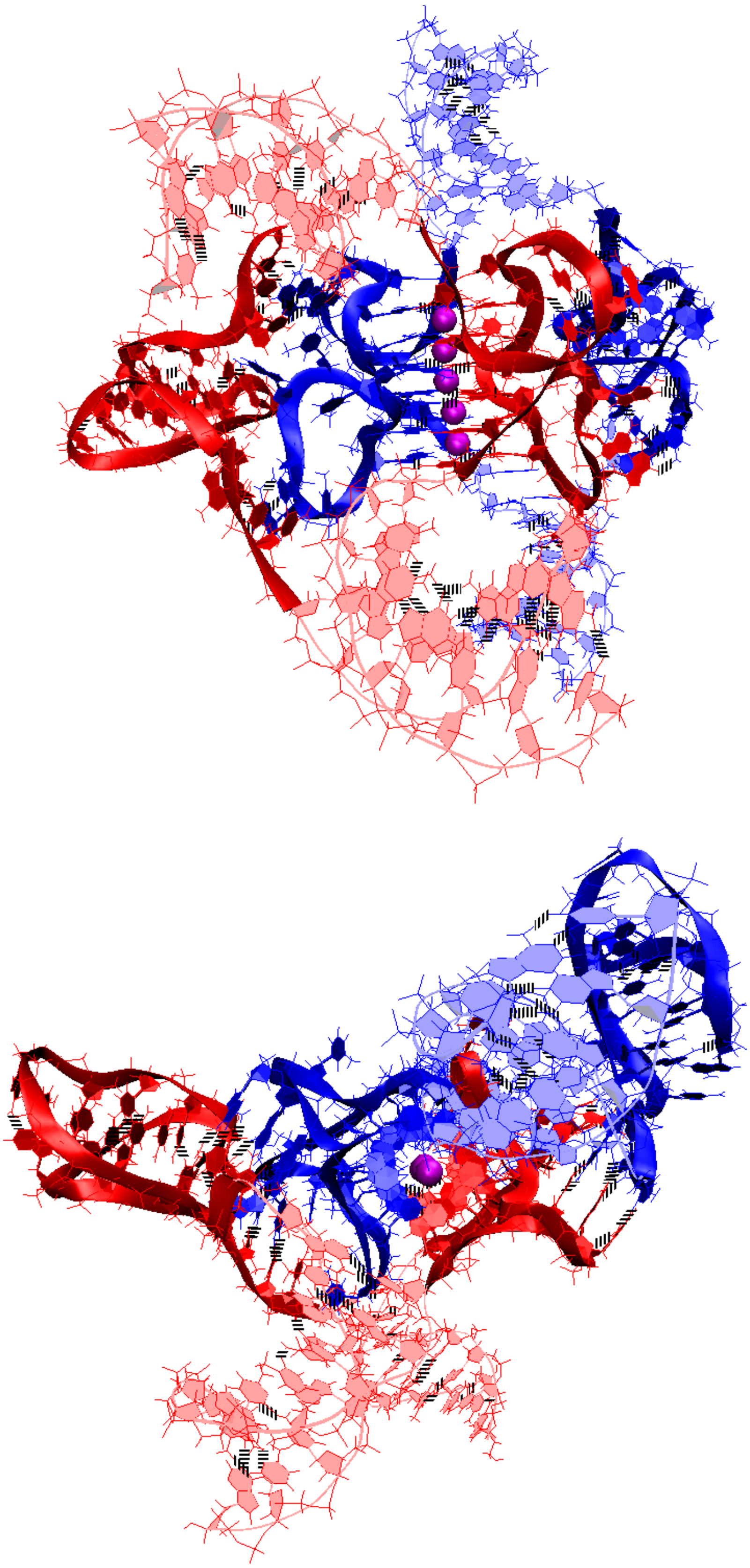

**A**

**Figure 7.2.A.S.1. Case 3: “Stacking of right and left handed parallel G4-dimers, and two IM-monomers with 4 mini-unmelted fragments of duplexes”**: **A** – conformation obtained at the last step of the MD trajectory (side and top view).

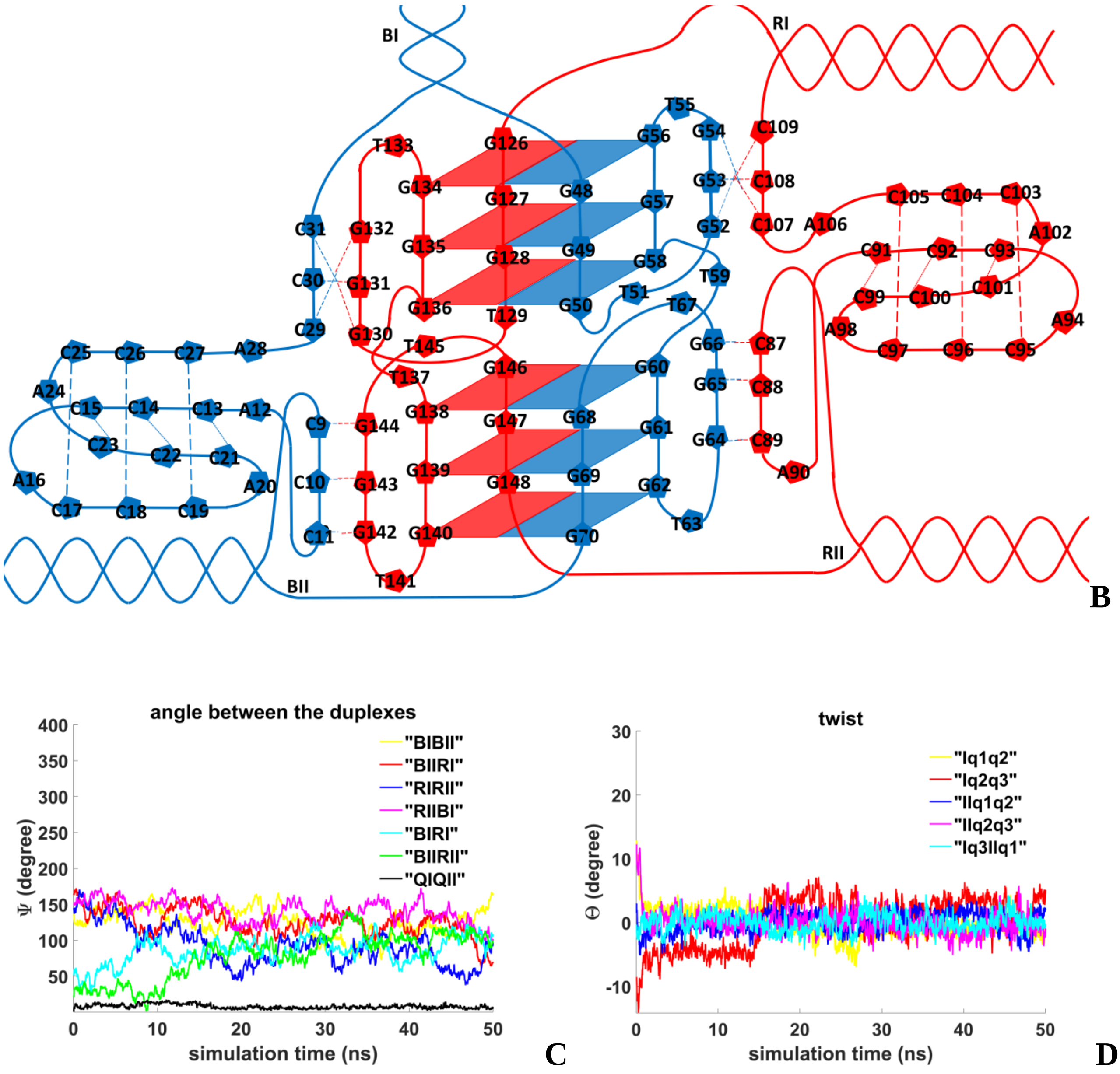


**Figure 7.2.A.S.2. Case 3: "Stacking of right and left handed parallel G4-dimers, and two IM-monomers with 4 mini-unmelted fragments of duplexes"**: **B** – the complex scheme; **C** – angles between unmelted fragments of the duplexes and axes passing through COMs of the tetrads, angle between axes passing through COMs of the boundary tetrads (**QIQII**); **D**– angles of rotation of the tetrads relative to each other.

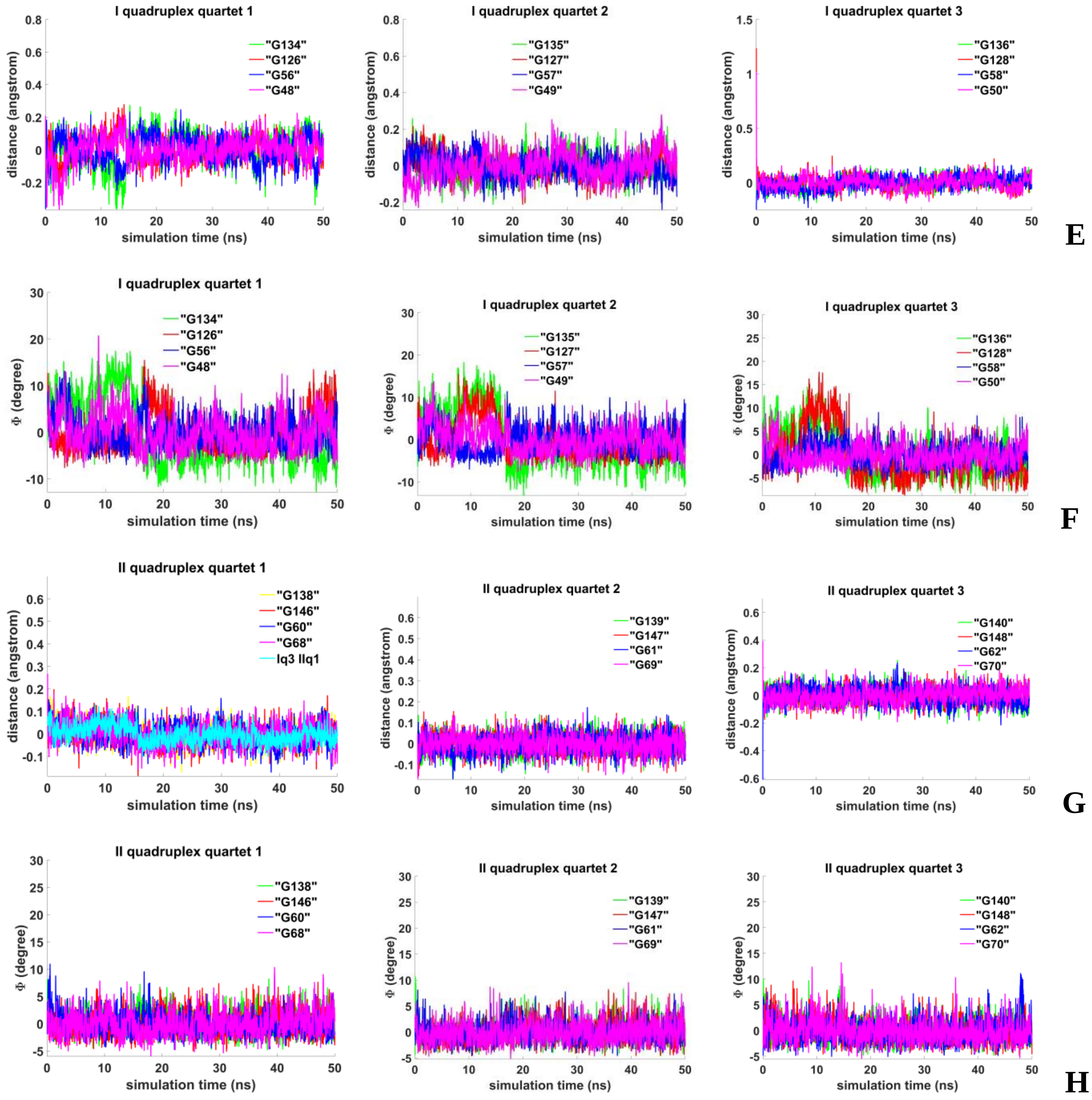


**Figure 7.2.A.S.3. Case 3: "Stacking of right and left handed parallel G4-dimers, and two IM-monomers with 4 mini-unmelted fragments of duplexes"**: **E, G -** distances from COMs of the guanines' bases to COMs of their containing tetrads, distance between COMs of the boundary tetrads (**Iq3 IIq1**); **F, H** - angles between normals to the guanines' bases and vectors connecting COMs of the boundary tetrads.

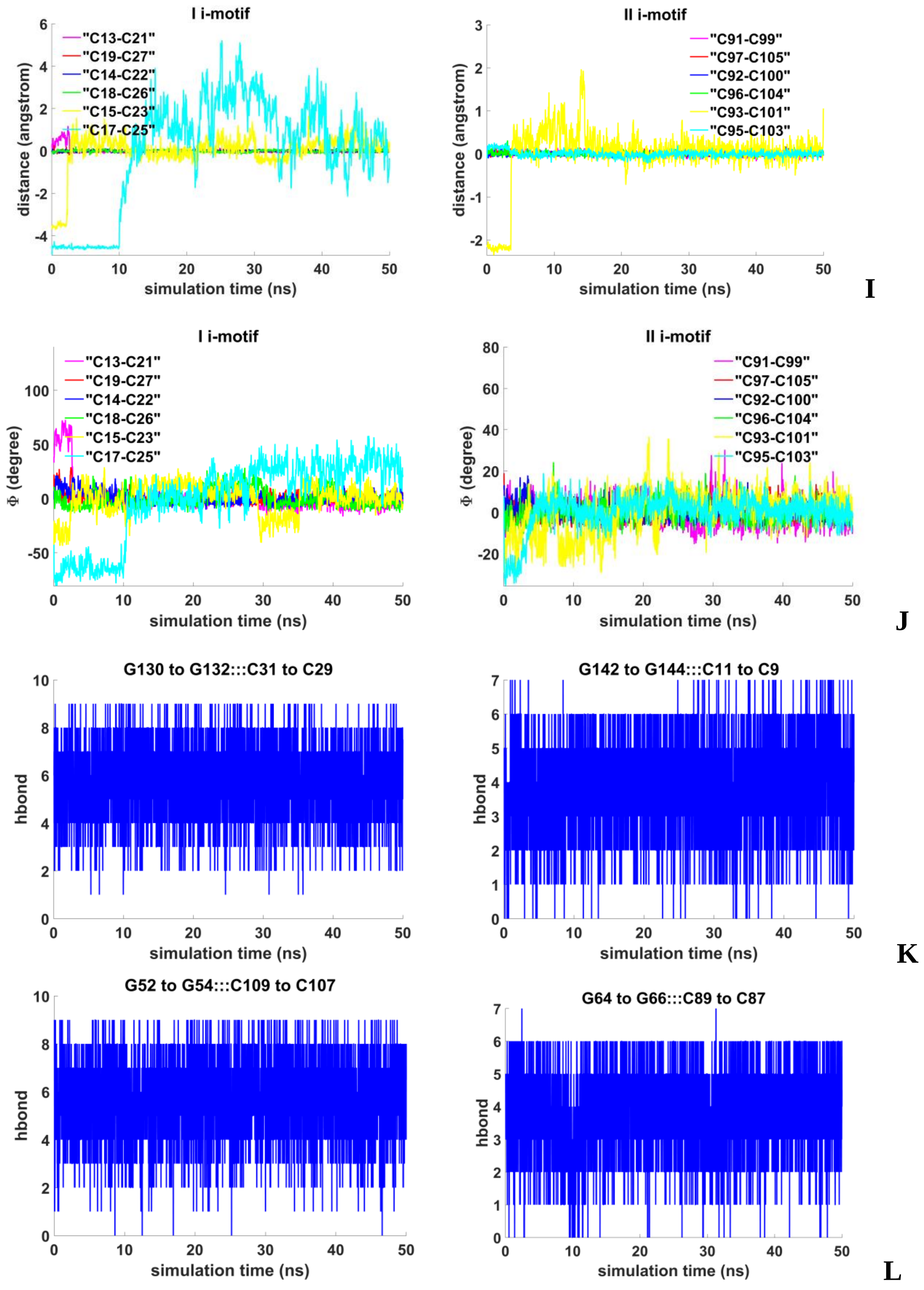


**Figure 7.2.A.S.4. Case 3: "Stacking of right and left handed parallel G4-dimers, and two IM-monomers with 4 mini-unmelted fragments of duplexes"**: **I** - distances between COMs of the cytosines' bases; **J** - angles between normals to the cytosines' bases; **K**, **L** - number of hydrogen bonds in mini-duplexes.

| donor | acceptor | occupancy | donor | acceptor | occupancy |
|---|---|---|---|---|---|
| DG64-Side-N2 | DC89-Side-O2 | 65,13% | DG52-Side-N2 | DC109-Side-O2 | 64,74% |
| DG65-Side-N1 | DC88-Side-N3 | 67,29% | DG53-Side-N2 | DC108-Side-O2 | 69,62% |
| DG65-Side-N2 | DC88-Side-O2 | 58,94% | DG52-Side-N1 | DC109-Side-N3 | 66,71% |
| DG64-Side-N1 | DC89-Side-N3 | 71,45% | DG53-Side-N1 | DC108-Side-N3 | 66,90% |
| DC89-Side-N4 | DG64-Side-O6 | 60,55% | DC109-Side-N4 | DG52-Side-O6 | 50,19% |
| DC88-Side-N4 | DG65-Side-O6 | 54,05% | DC108-Side-N4 | DG53-Side-O6 | 62,89% |
| DG66-Side-N2 | DC87-Side-O2 | 0,07% | DG54-Side-N1 | DC107-Side-N3 | 64,71% |
| DG66-Side-N1 | DC87-Side-O2 | 0,02% | DG54-Side-N2 | DC107-Side-O2 | 58,39% |
| DG66-Side-N1 | DC87-Side-N3 | 0,01% | DC107-Side-N4 | DG54-Side-O6 | 56,24% |
| DC87-Side-N4 | DG66-Side-OP2 | 0,06% | DG52-Side-N2 | DC109-Side-N3 | 0,12% |
| DG64-Side-N2 | DC89-Side-N3 | 0,01% | DG52-Side-N1 | DC109-Side-N4 | 0,02% |
| DC88-Side-N4 | DG64-Side-O6 | 0,46% | DG54-Side-N2 | DC107-Side-N3 | 0,02% |
| DG65-Side-N2 | DC88-Side-N3 | 0,10% | DG52-Side-N1 | DC109-Side-O2 | 0,05% |
| DG64-Side-N1 | DC89-Side-O2 | 0,12% | DC107-Side-N4 | DG53-Side-O6 | 0,01% |
| DG65-Side-N2 | DC89-Side-O2 | 0,02% | DG54-Side-N1 | DC107-Side-O2 | 0,04% |
| DC87-Side-N4 | DG65-Side-O6 | 0,86% | | | |
| DC87-Side-N4 | DG65-Side-N7 | 0,02% | | | |

| donor | acceptor | occupancy | donor | acceptor | occupancy |
|---|---|---|---|---|---|
| DC11-Side-N4 | DG142-Side-O6 | 58,37% | DC31-Side-N4 | DG130-Side-O6 | 54.13% |
| DC10-Side-N4 | DG143-Side-O6 | 55,63% | DC30-Side-N4 | DG131-Side-O6 | 63.98% |
| DG143-Side-N2 | DC10-Side-O2 | 50,86% | DG131-Side-N2 | DC30-Side-O2 | 69.14% |
| DG144-Side-N1 | DC9-Side-N3 | 0,09% | DG132-Side-N2 | DC29-Side-O2 | 54.57% |
| DG142-Side-N1 | DC11-Side-N3 | 65,86% | DG132-Side-N1 | DC29-Side-N3 | 73.71% |
| DG143-Side-N1 | DC10-Side-N3 | 69,97% | DG130-Side-N1 | DC31-Side-N3 | 67.47% |
| DC9-Side-N4 | DG144-Side-O6 | 0,41% | DG130-Side-N2 | DC31-Side-O2 | 66.30% |
| DG144-Side-N2 | DC9-Side-O2 | 0,24% | DG131-Side-N1 | DC30-Side-N3 | 74.33% |
| DG142-Side-N2 | DC11-Side-O2 | 64,56% | DC29-Side-N4 | DG132-Side-O6 | 58.39% |
| DG144-Side-N2 | DC9-Side-N3 | 0,21% | DC29-Side-N4 | DG131-Side-O6 | 0.20% |
| DG144-Side-N1 | DC9-Side-O2 | 0,07% | DG130-Side-N2 | DC31-Side-N3 | 0.05% |
| DG144-Side-N2 | DC10-Side-O2 | 0,02% | DG130-Side-N1 | DC31-Side-O2 | 0.02% |
| DG144-Side-N2 | DC11-Side-O4' | 0,02% | DG132-Side-N2 | DC30-Side-N3 | 0.01% |
| DG144-Side-N2 | DC11-Side-O2 | 0,01% | DC30-Side-N4 | DG130-Side-O6 | 0.01% |
| DG144-Side-N2 | DC11-Side-O3' | 0,27% | DG132-Side-N2 | DC29-Side-N3 | 0.01% |
| DG142-Side-N2 | DC11-Side-N3 | 0,04% | | | |
| DG144-Side-N1 | DC11-Side-O3' | 22,68% | | | |
| DG144-Side-N1 | DC11-Side-O4' | 0,09% | | | |
| DC10-Side-N4 | DG142-Side-O6 | 0,01% | | | |
| DG143-Side-N1 | DC10-Side-O2 | 0,01% | | | |
| DG143-Side-N2 | DC11-Side-O2 | 0,04% | | | |
| DC9-Side-N4 | DG143-Side-O6 | 0,05% | | | |

**Table .2. (Appendix to Figure 7.2.A.S.4. Case 3) Percentage of snapshots with hydrogen bonds generated by pairs in the mini-duplexes.**

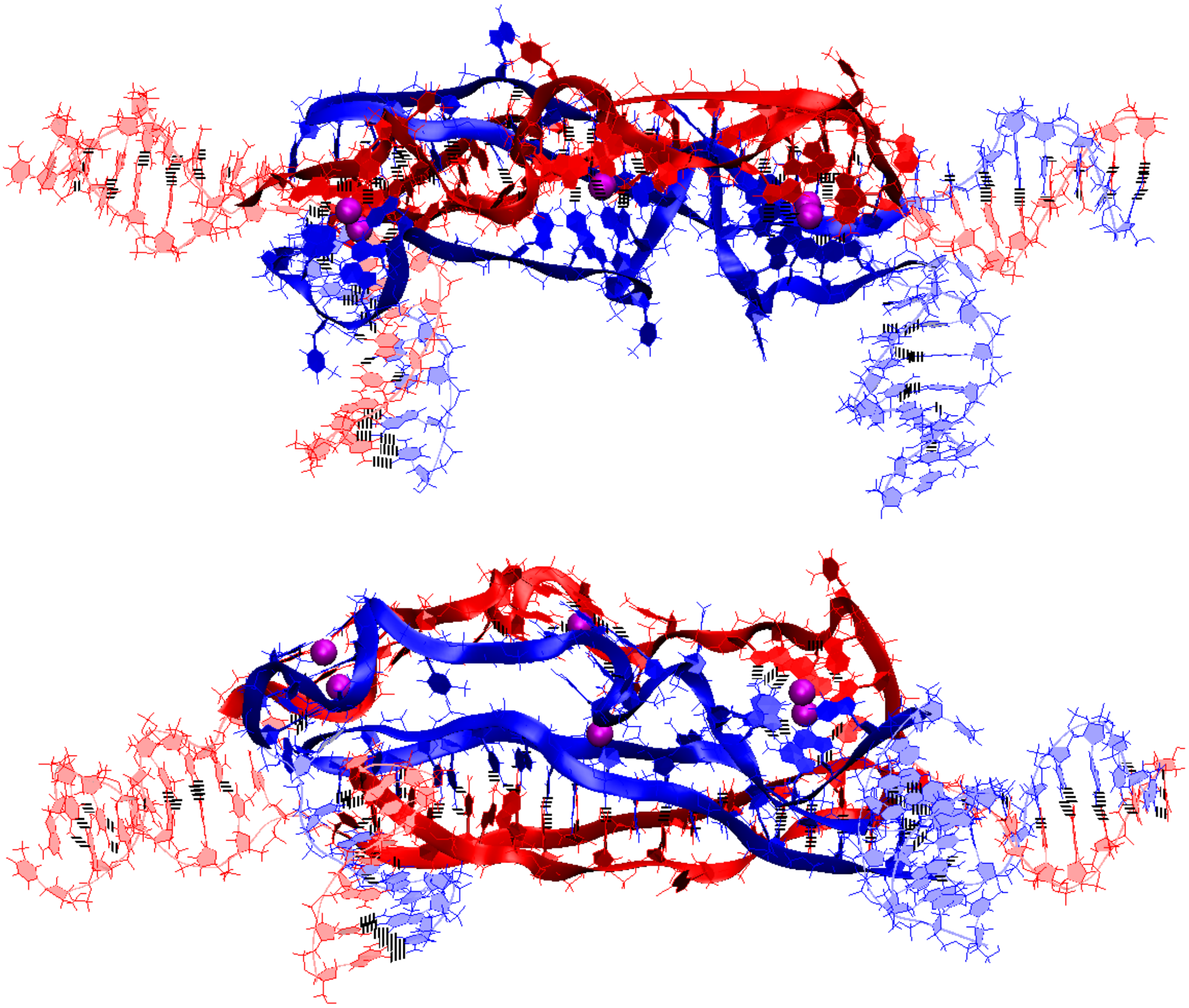

**A**

**Figure 7.2.B.S.1. Case 4: "Three parallel G4-dimers in the same plane and head-to-tail IM-dimer with the strands exchange"**: **A** – conformation obtained at the last step of the MD trajectory (side and top view).

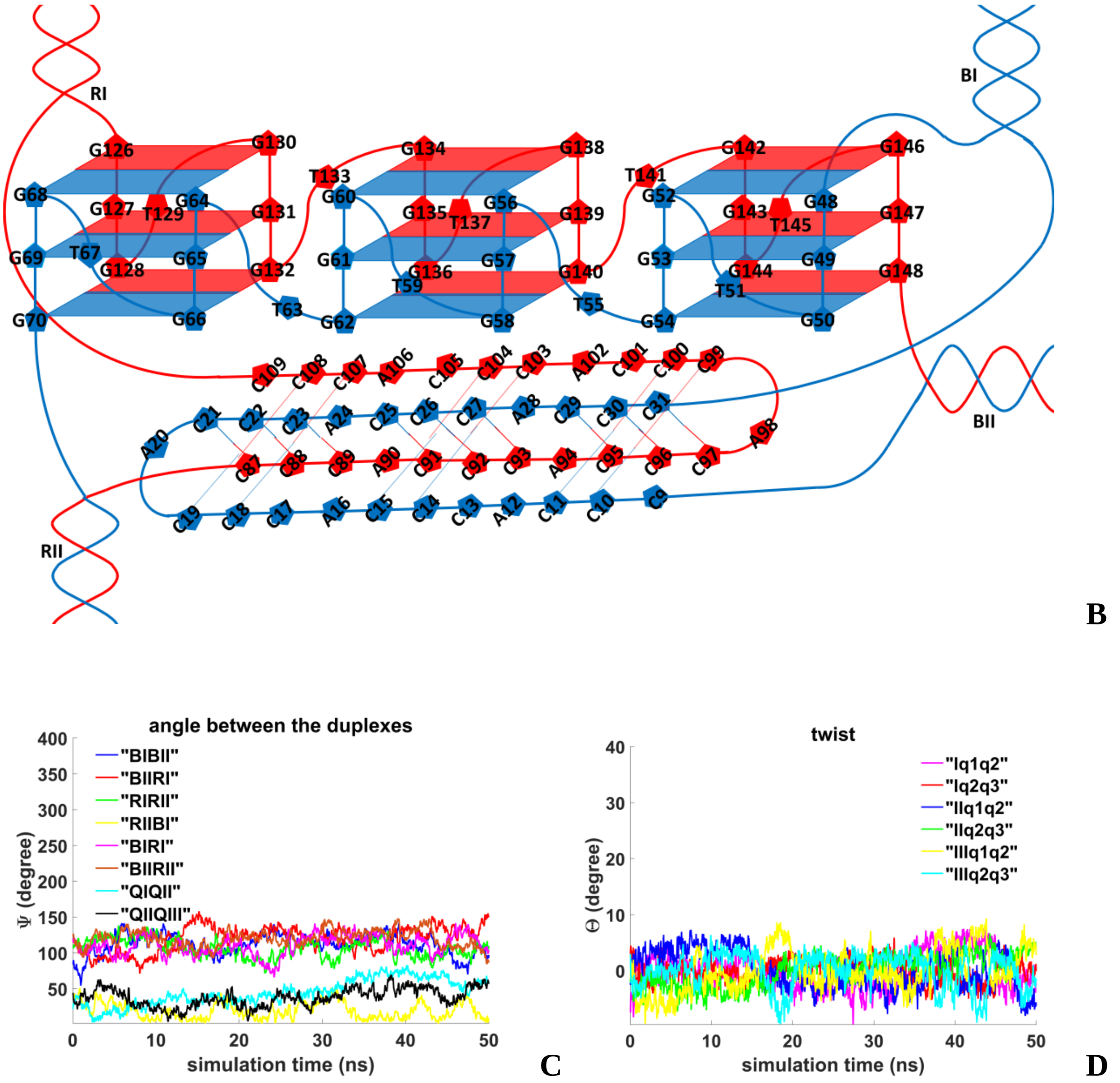


**Figure 7.2.B.S.2. Case 4: "Three parallel G4-dimers in the same plane and head-to-tail IM-dimer with the strands exchange"**: **B** – the complex scheme; **C** – angles between unmelted fragments of the duplexes and axes passing through COMs of the tetrads, angle between axes passing through COMs of the boundary tetrads (**QIQII, QIIQIII**); **D** – angles of rotation of the tetrads relative to each other.

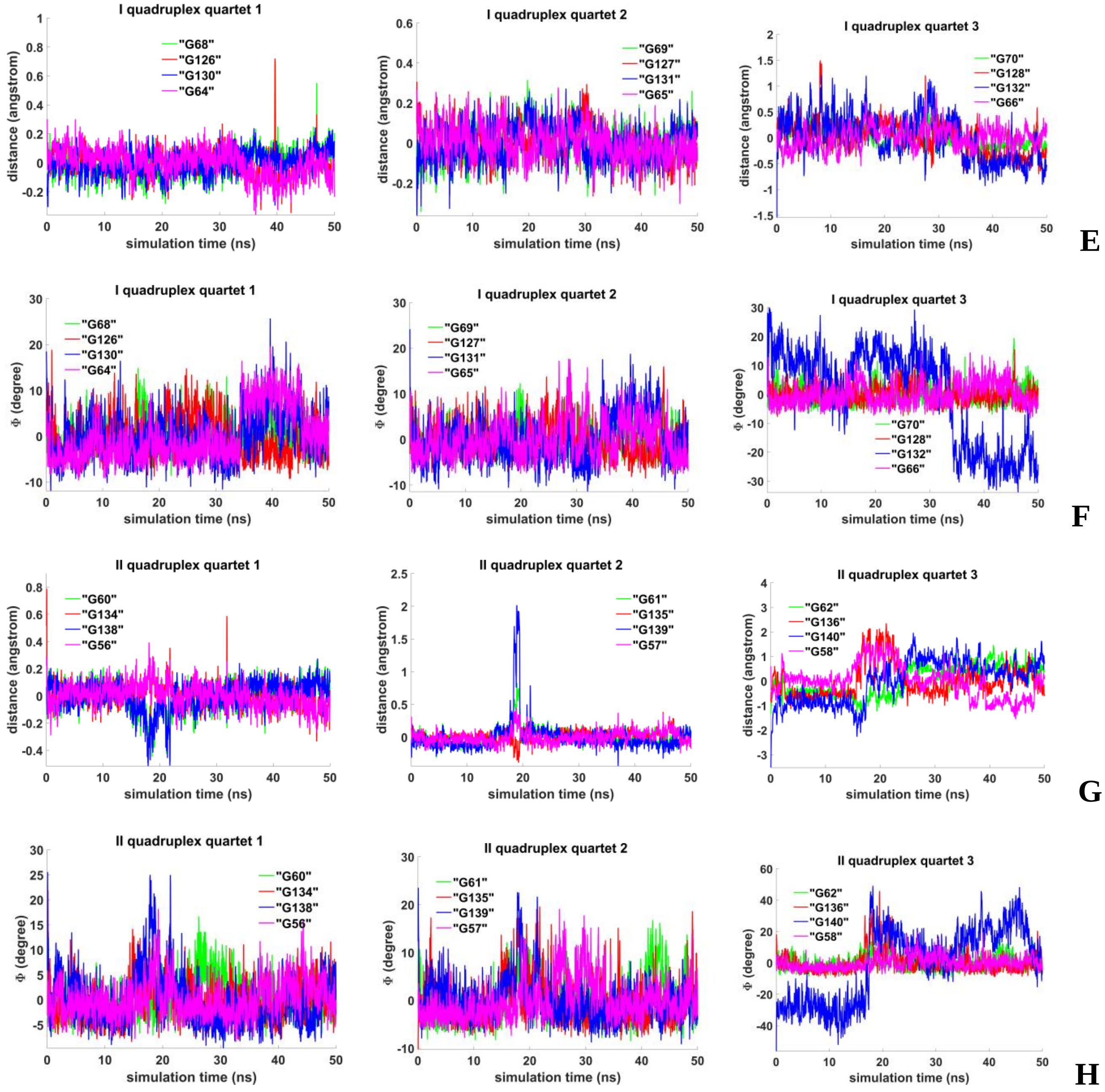


**Figure 7.2.B.S.3. Case 4: “Three parallel G4-dimers in the same plane and head-to-tail IM-dimer with the strands exchange”**: **E, G** - distances from COMs of the guanines’ bases to COMs of their containing tetrad; **F, H** - angles between normals to the guanines’ bases and vectors connecting COMs of the boundary tetrads.

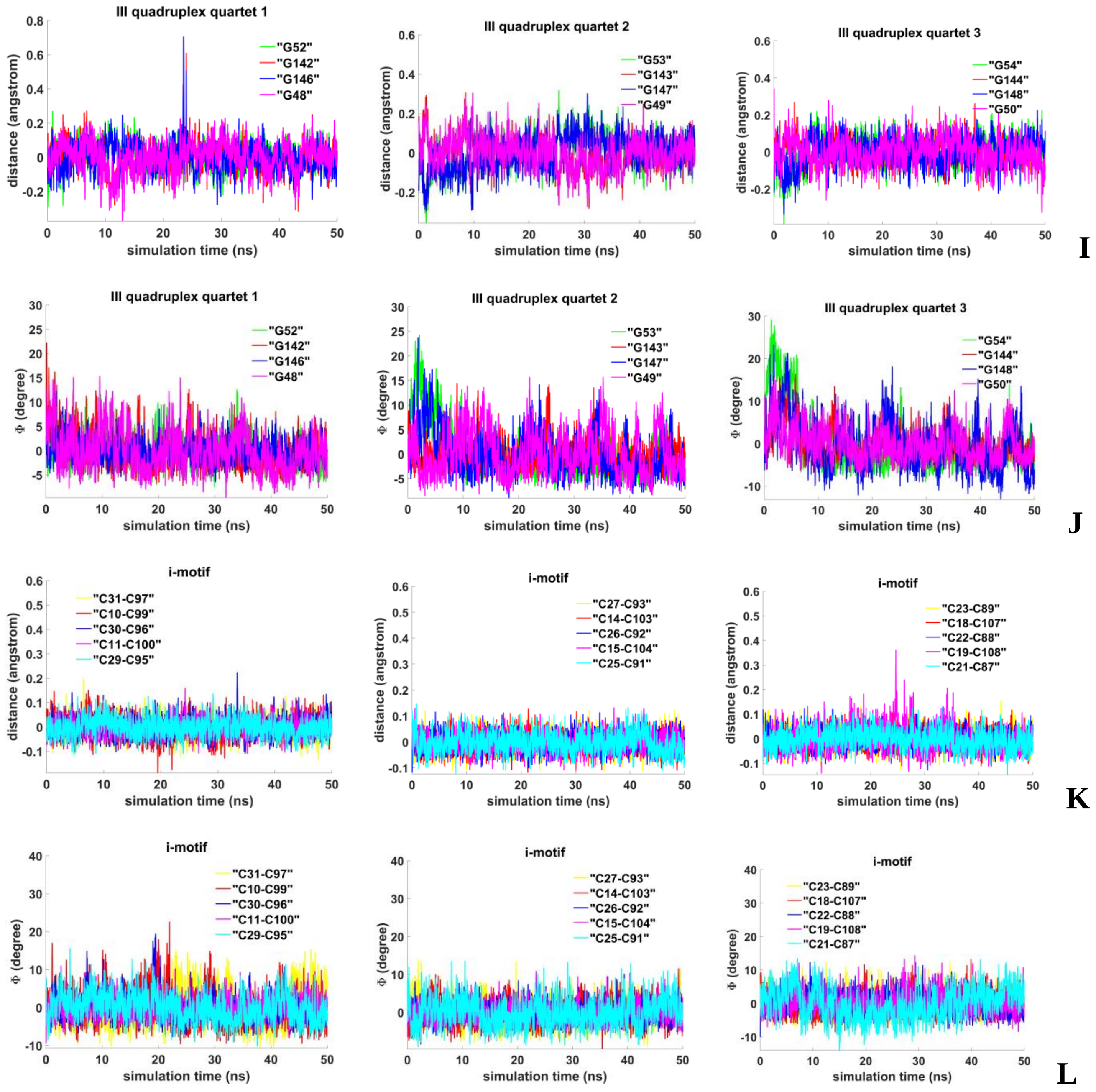


**Figure 7.2.B.S.4. Case 4: "Three parallel G4-dimers in the same plane and head-to-tail IM-dimer with the strands exchange"**: **I** - distances from COMs of the guanines' bases to COMs of their containing tetrad, **J** - angles between normals to the guanines' bases and vectors connecting COMs of the boundary tetrads; **K** - distances between COMs of the cytosines' bases; **L** - angles between normals to the cytosines' bases.

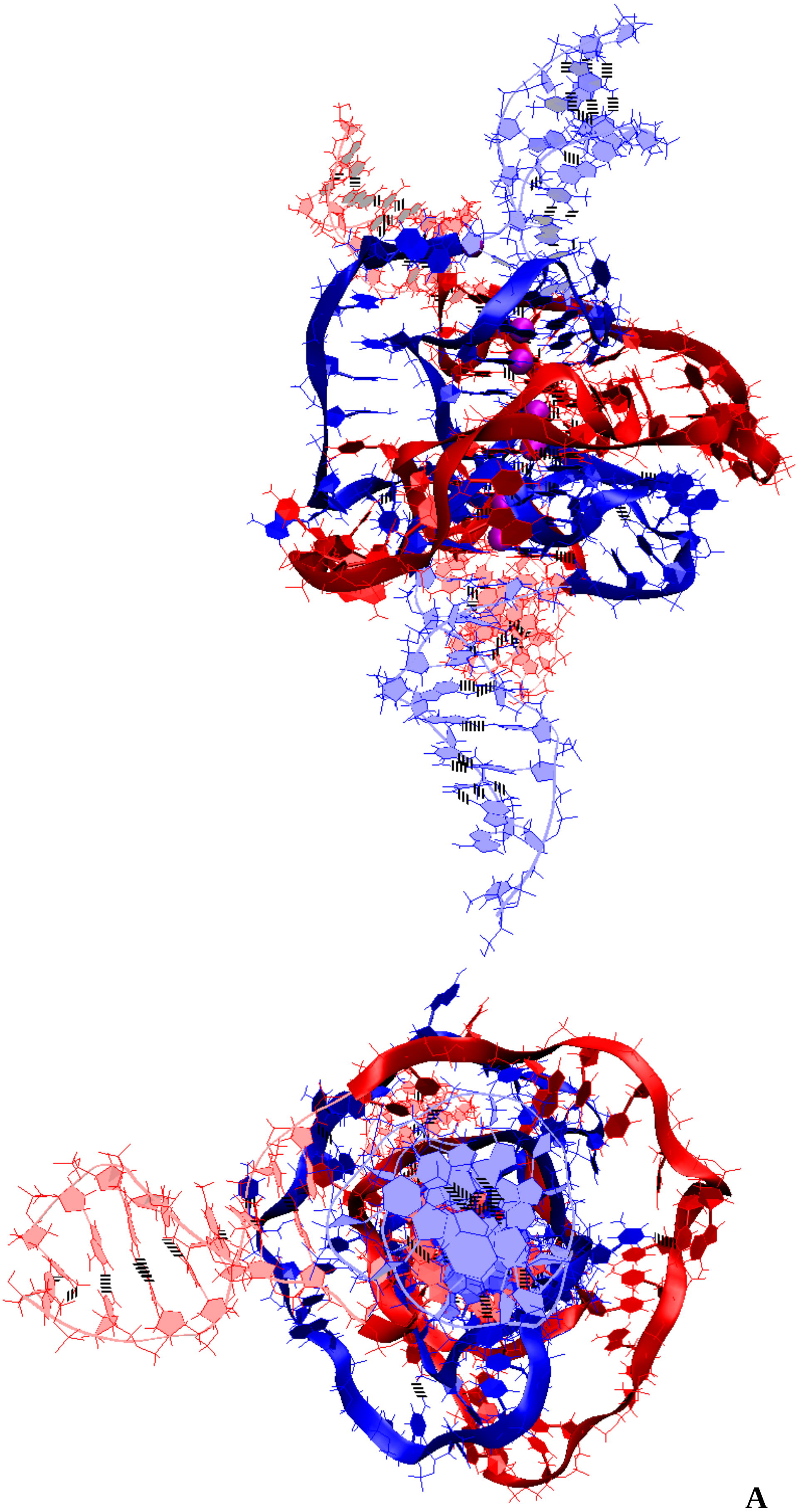

**A**

**Figure 7.3.A.S.1. Case 5: “Stacking of three parallel G4-dimers”**: **A** – conformation obtained at the last step of the MD trajectory (side and top view).

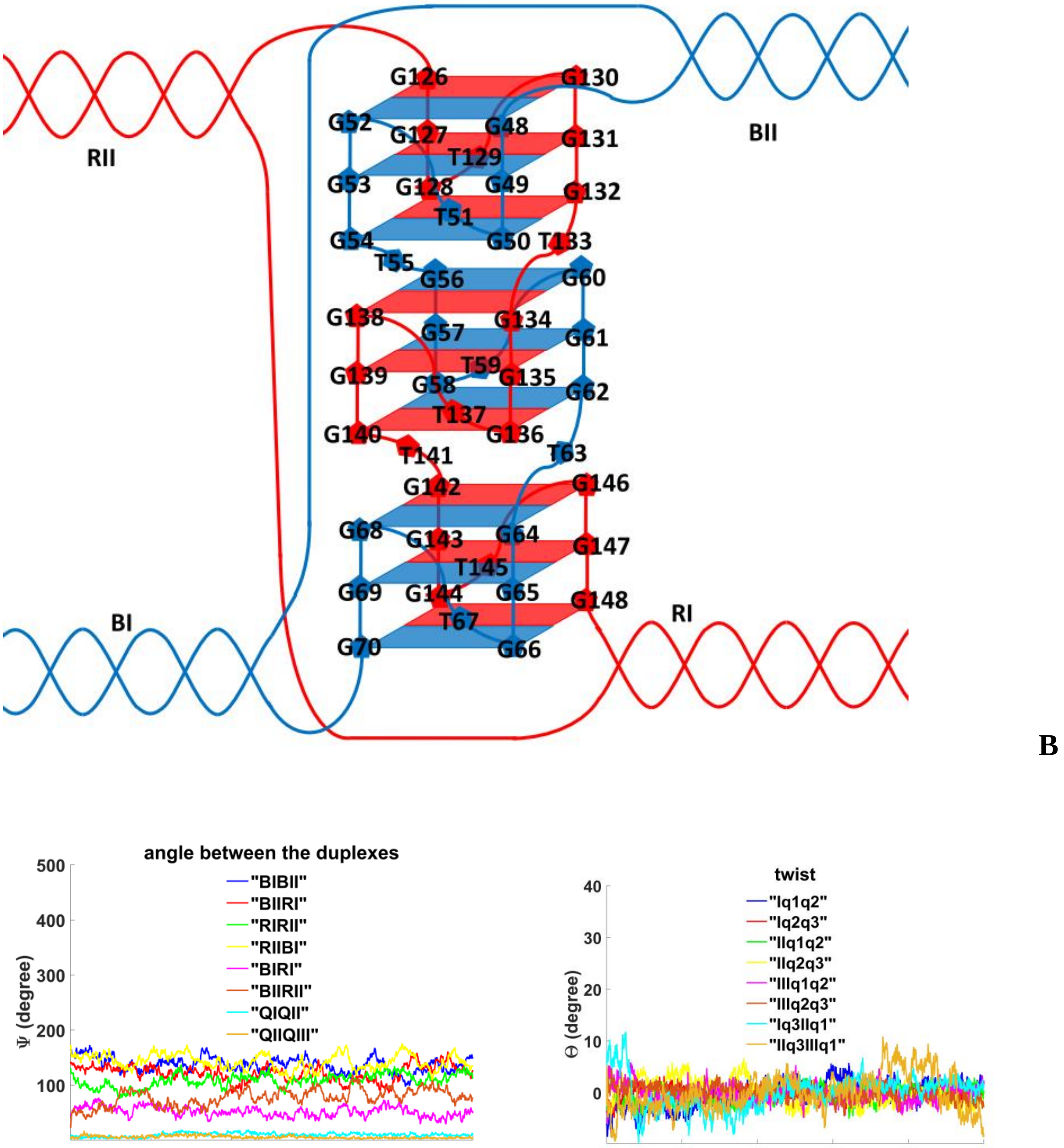


**Figure 7.3.A.S.2. Case 5: "Stacking of three parallel G4-dimers"**: **B** – the complex scheme; **C** – angles between unmelted fragments of the duplexes and axes passing through COMs of the tetrads, angle between axes passing through COMs of the boundary tetrads (**QIQII, QIIQIII**); **D** – angles of rotation of the tetrads relative to each other.

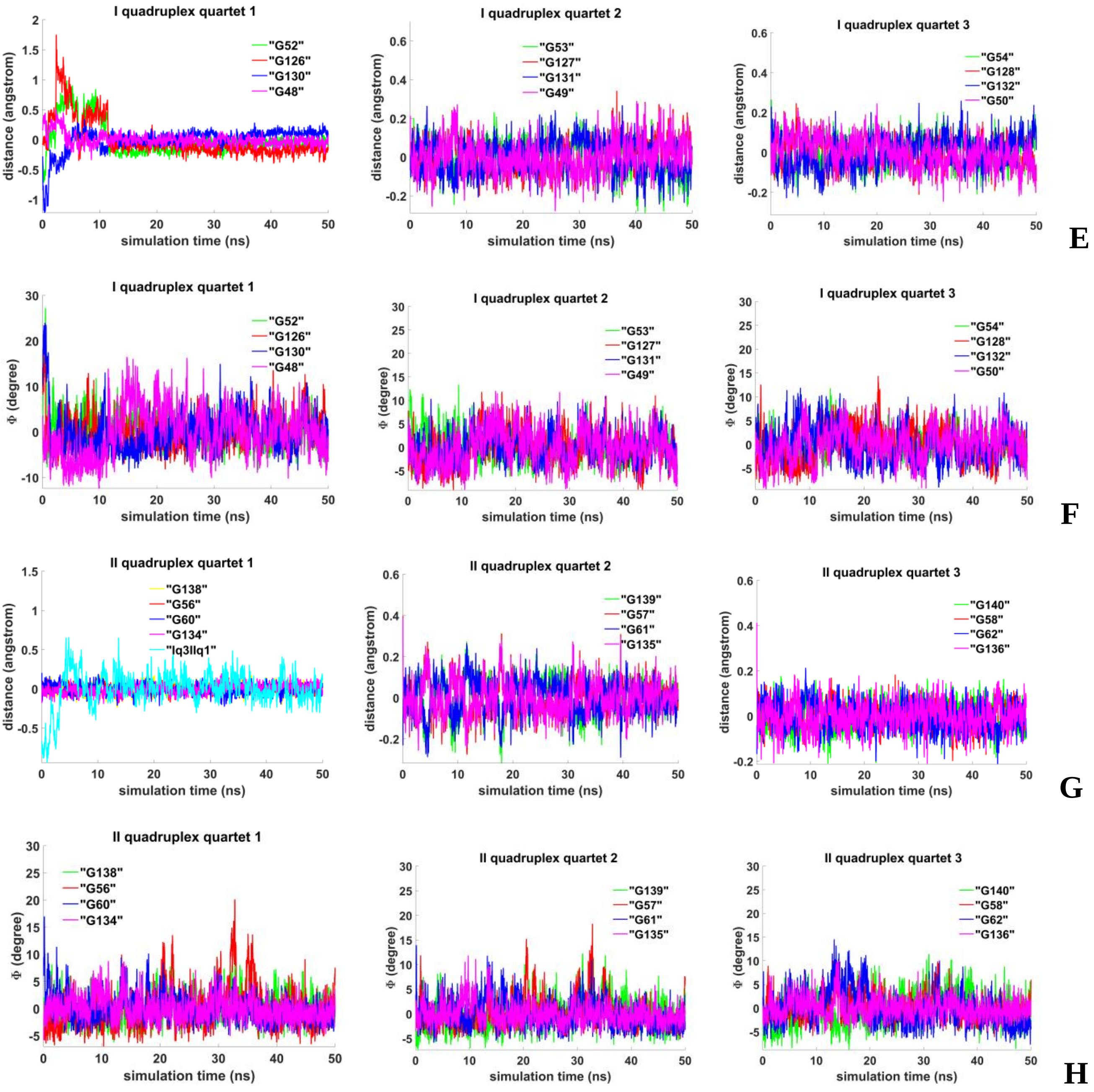


**Figure 7.3.A.S.3. Case 5: "Stacking of three parallel G4-dimers"**: **E, G** - distances from COMs of the guanines' bases to COMs of their containing tetrad, distance between COMs of the boundary tetrads (**Iq3 IIq1**); **F, H** - angles between normals to the guanines' bases and vectors connecting COMs of the boundary tetrads.

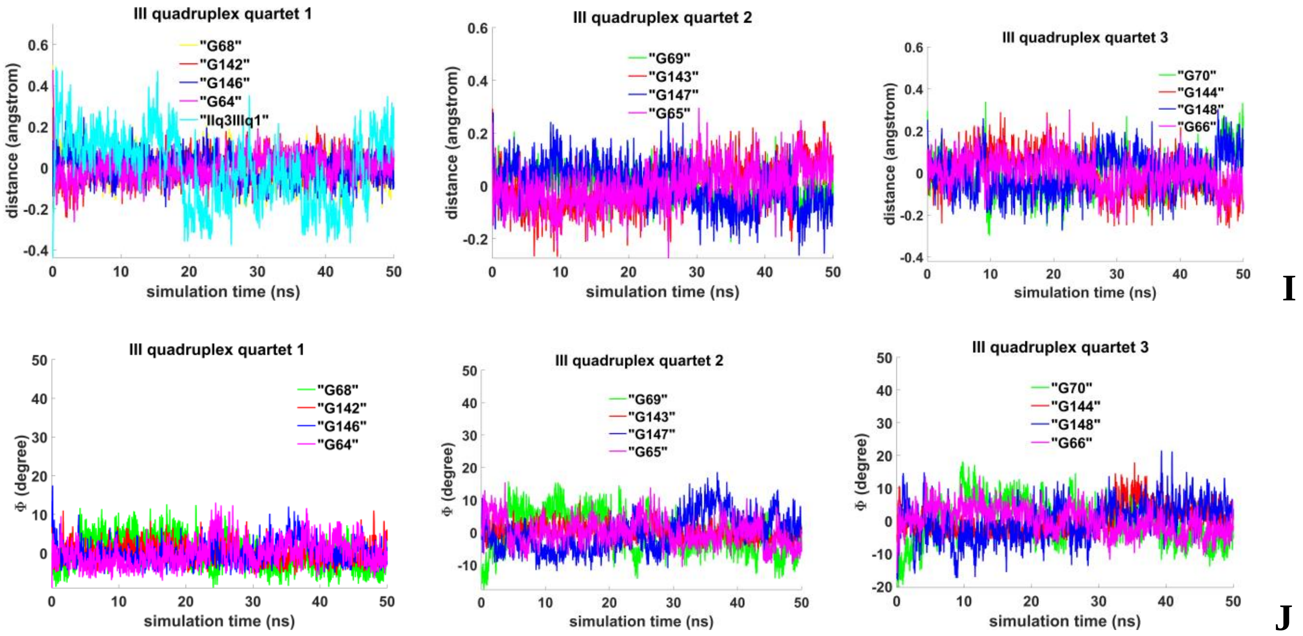


**Figure 7.3.A.S.4. Case 5: "Stacking of three parallel G4-dimers"**: **I** - distances from COMs of the guanines' bases to COMs of their containing tetrad, distance between COMs of the boundary tetrads (**IIq3 IIIq1**); **J** - angles between normals to the guanines' bases and vectors connecting COMs of the boundary tetrads.

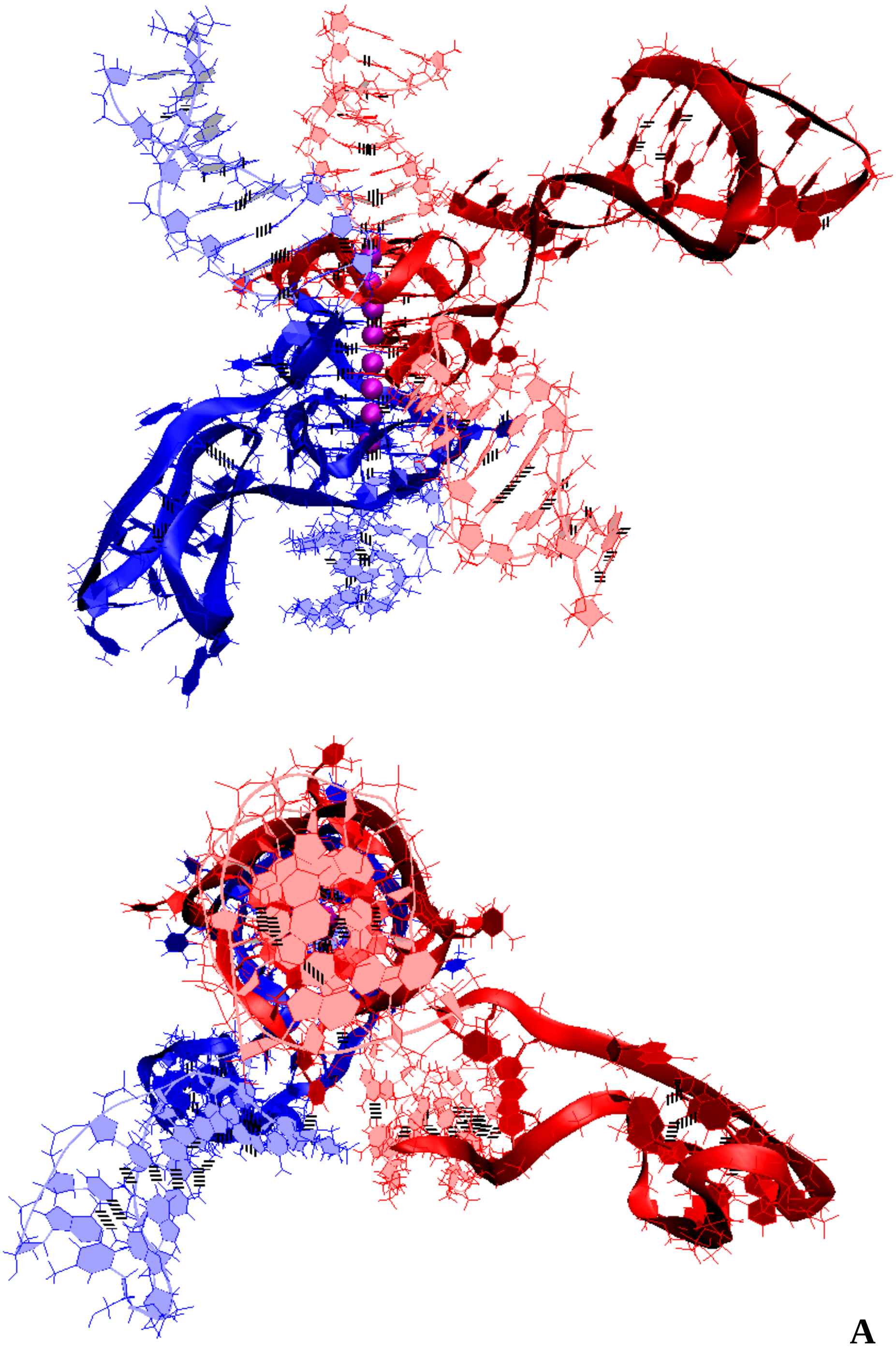

**A**

**Figure 7.3.B.S.1. Case 6: "Stacking of two parallel G4-monomers and G4-dimer, and two IM-monomers"**: **A** – conformation obtained at the last step of the MD trajectory (side and top view).

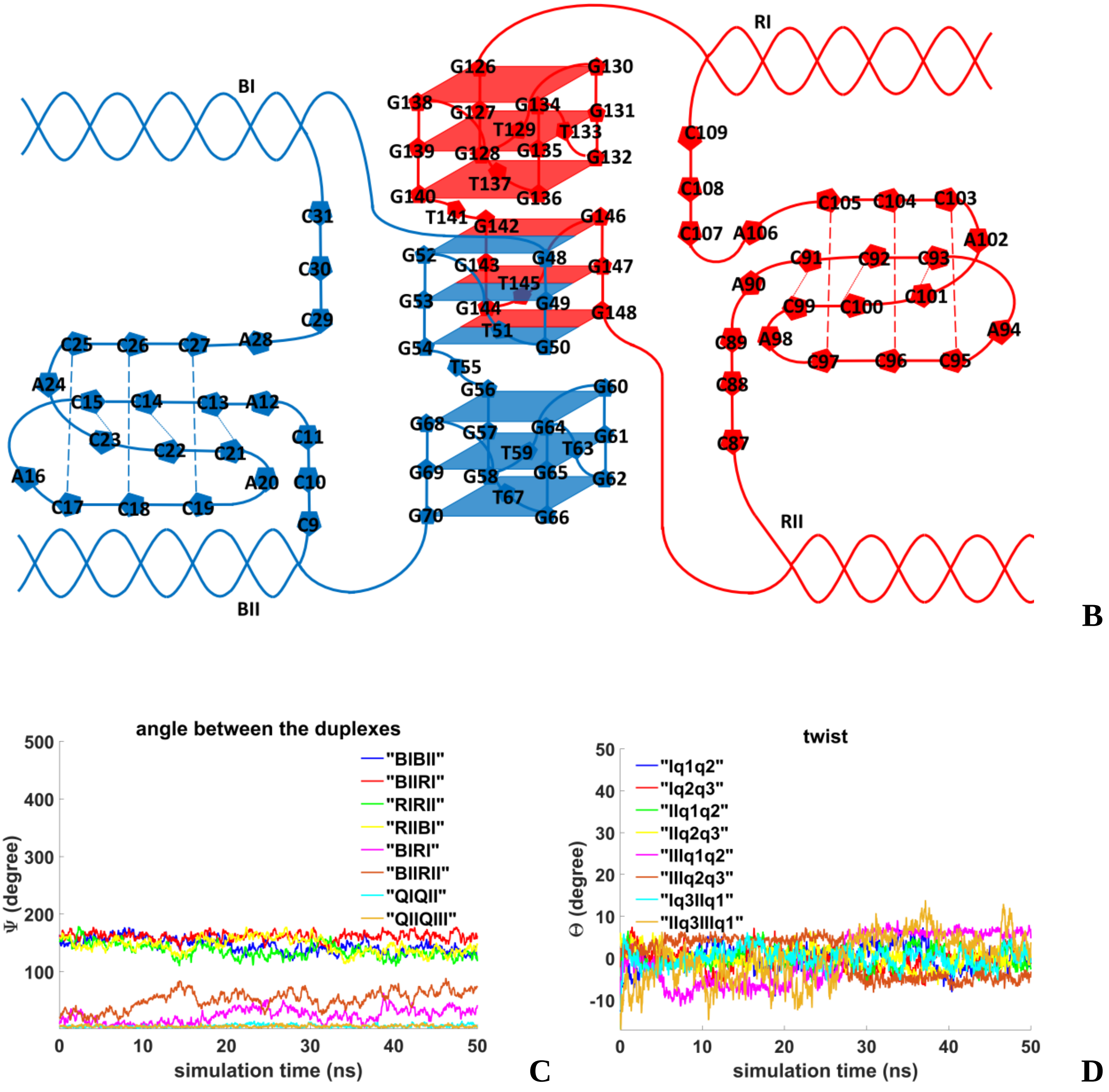


**Figure 7.3.B.S.2. Case 6: "Stacking of two parallel G4-monomers and G4-dimer, and two IM-monomers"**: **B** – the complex scheme; **C** – angles between unmelted fragments of the duplexes, angles between axes passing through the COMs of boundary tetrads of the G4s (**QIQII**, **QIIQIII**); **D** – angles of rotation of the tetrads relative to each other.

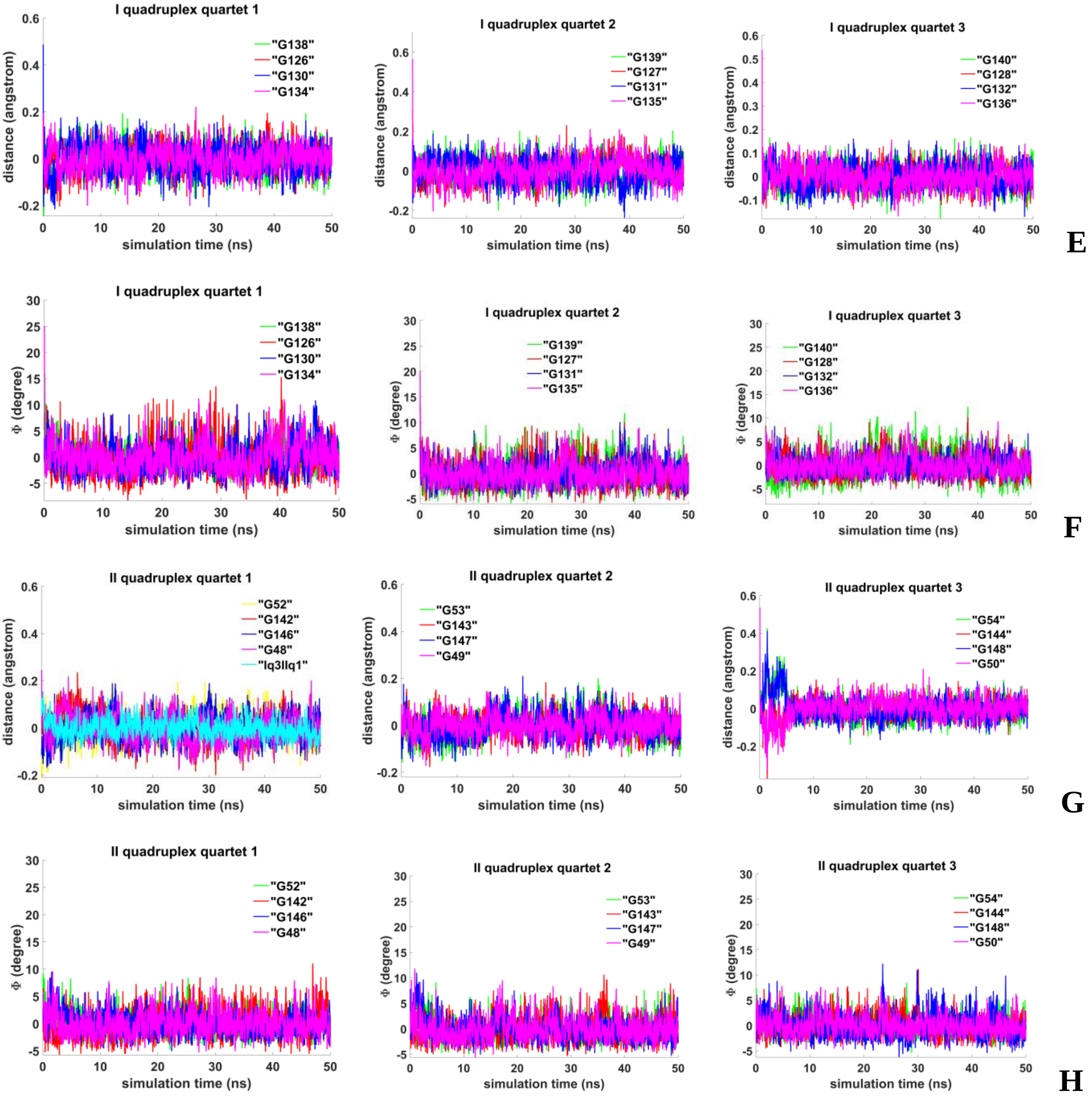


**Figure 7.3.B.S.3. Case 6: "Stacking of two parallel G4-monomers and G4-dimer, and two IM-monomers"**: **E, G** - distances from COMs of the guanines' bases to COMs of their containing tetrad, distance between COMs of the boundary tetrads (**Iq3 IIq1**); **F, H** - angles between normals to the guanines' bases and vectors connecting COMs of the boundary tetrads.

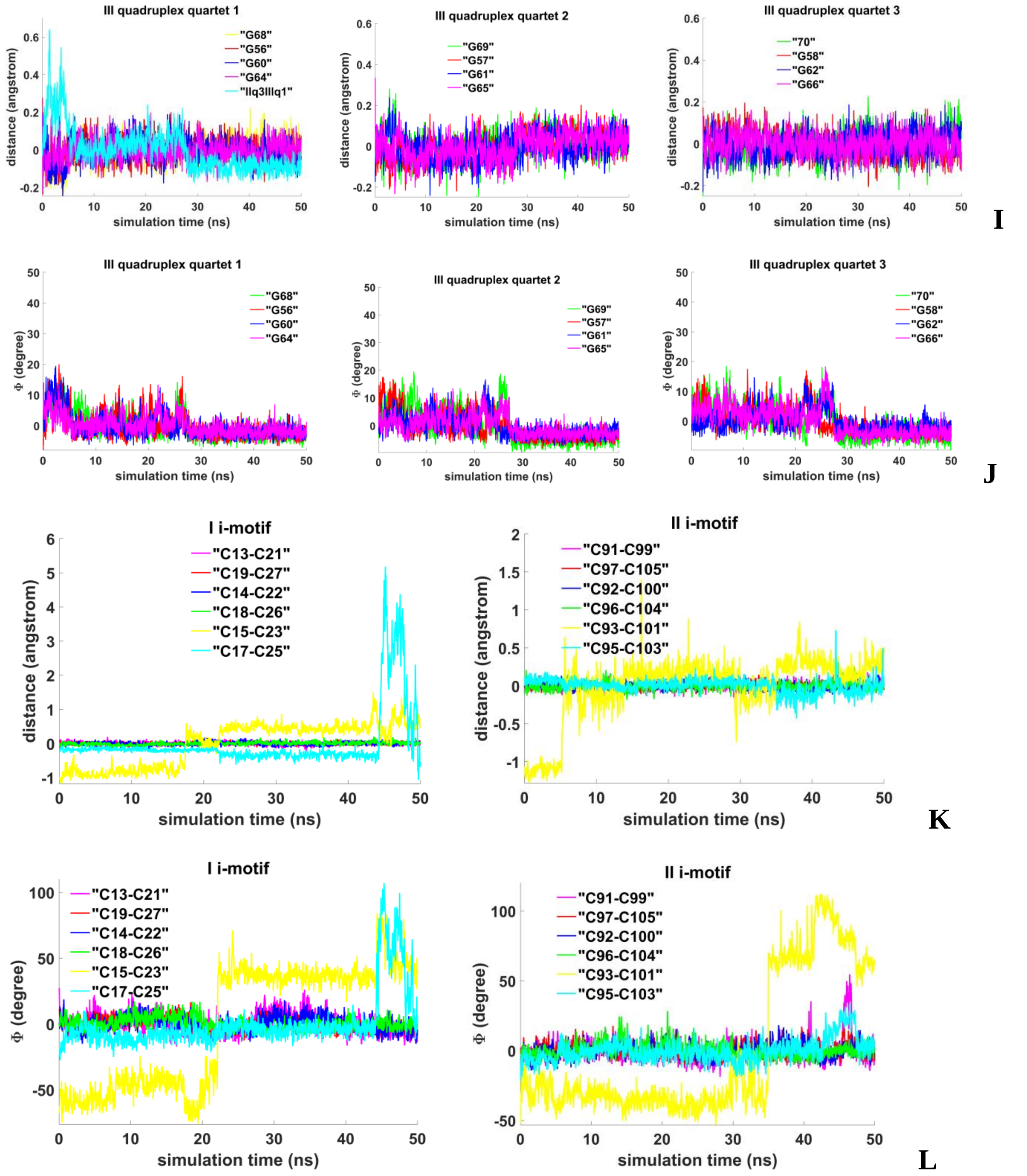


**Figure 7.3.B.S.4. Case 6: "Stacking of two parallel G4-monomers and G4-dimer, and two IM-monomers"**: **I** - distances from COMs of the guanines' bases to COMs of their containing tetrad, distance between COMs of the boundary tetrads (**IIq3 IIIq1**); **J** - angles between normals to the guanines' bases and vectors connecting COMs of the boundary tetrads; **K** - distances between COMs of the cytosines' bases; **L** - angles between normals to the cytosines' bases.

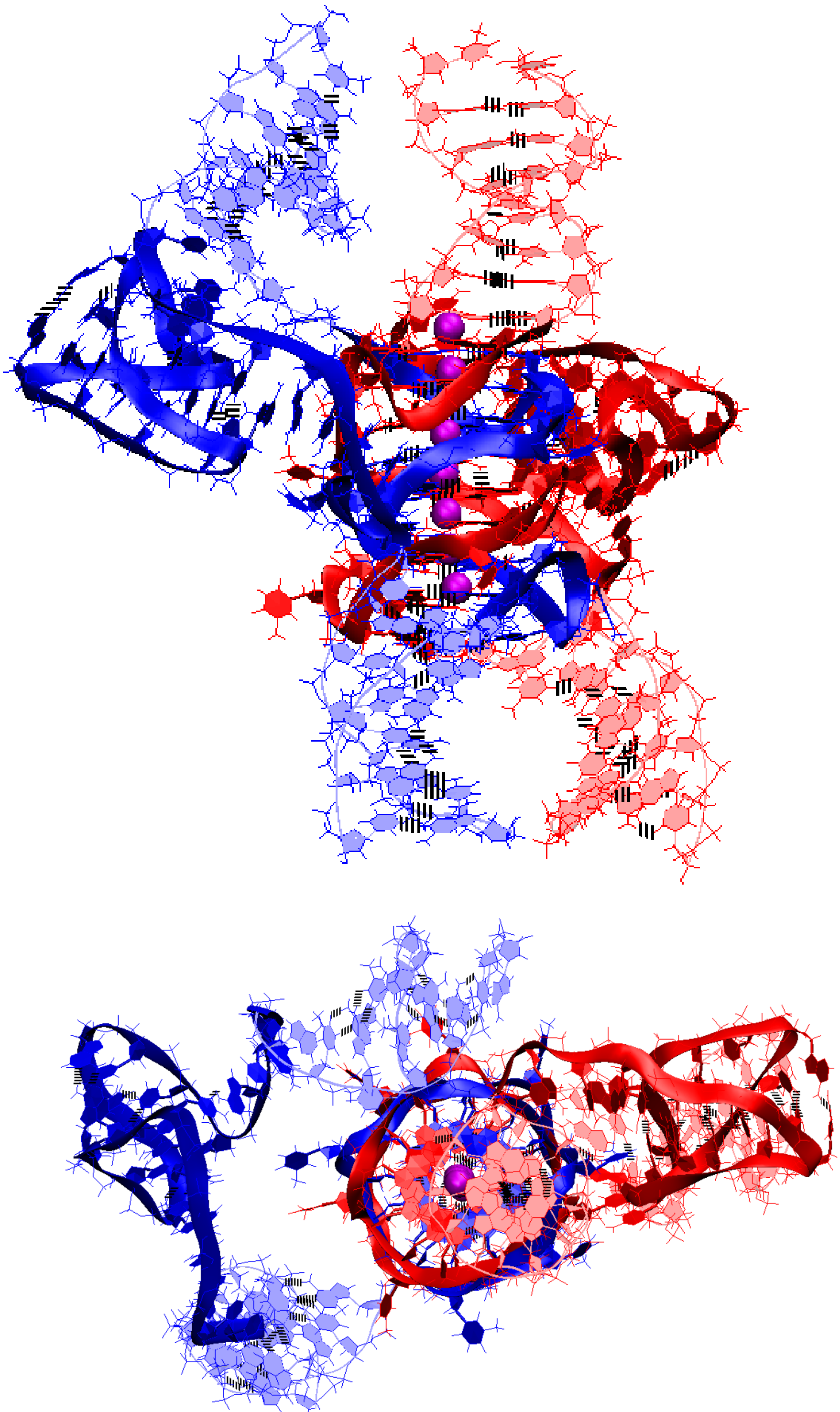

**A**

**Figure 7.4.A.S.1. Case 7: “Stacking of three parallel G4-dimer and two IM-monomers with mutual girth of the strands”: A** – conformation obtained at the last step of the MD trajectory (side and top view).

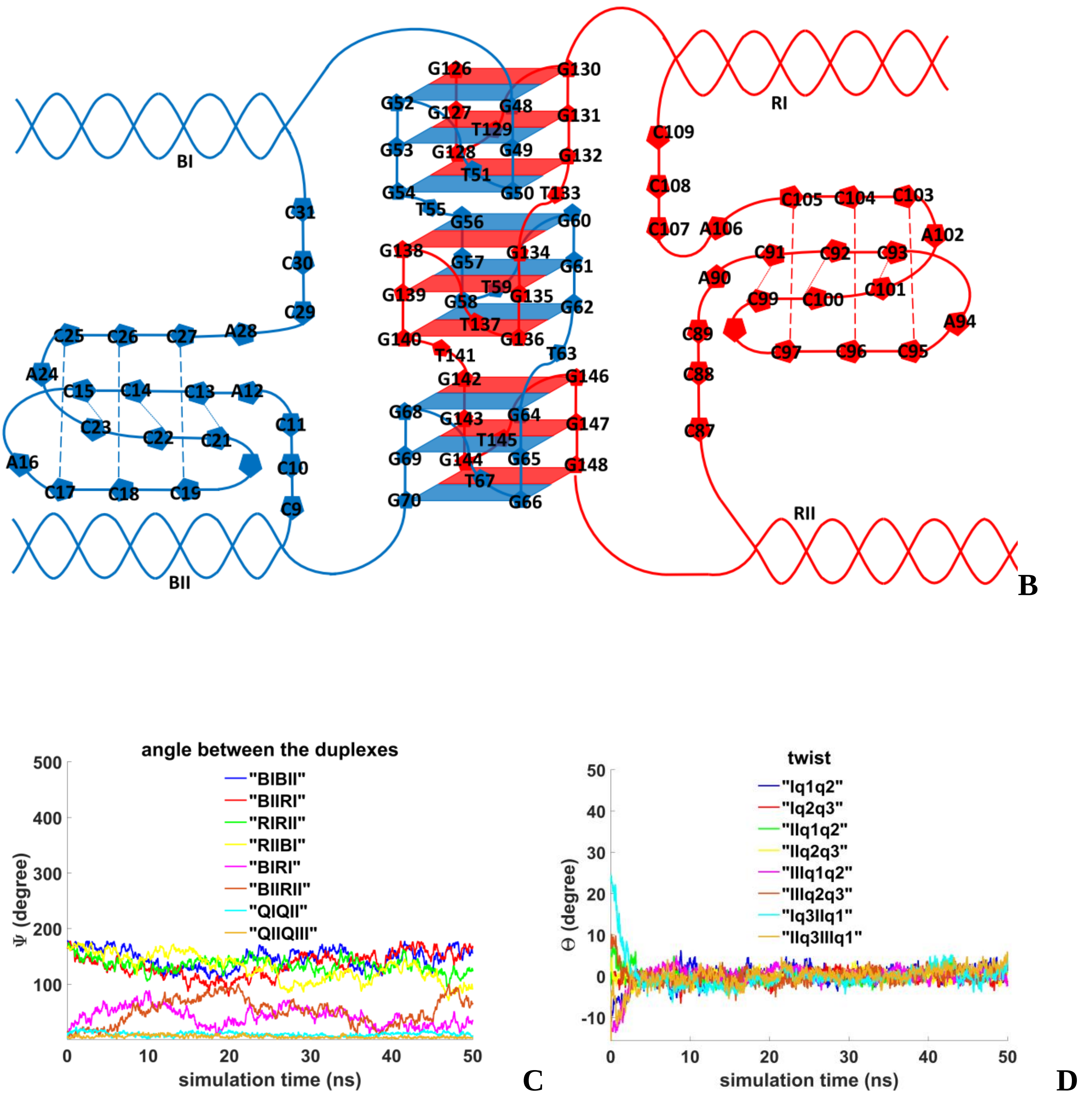


**Figure 7.4.A.S.2. Case 7: "Stacking of three parallel G4-dimer and two IM-monomers with mutual girth of the strands": B** – the complex scheme; **C** – angles between unmelted fragments of the duplexes, angles between axes passing through the COMs of boundary tetrads of the G4s (**QIQII**, **QIIQIII**); **D**– angles of rotation of the tetrads relative to each other.

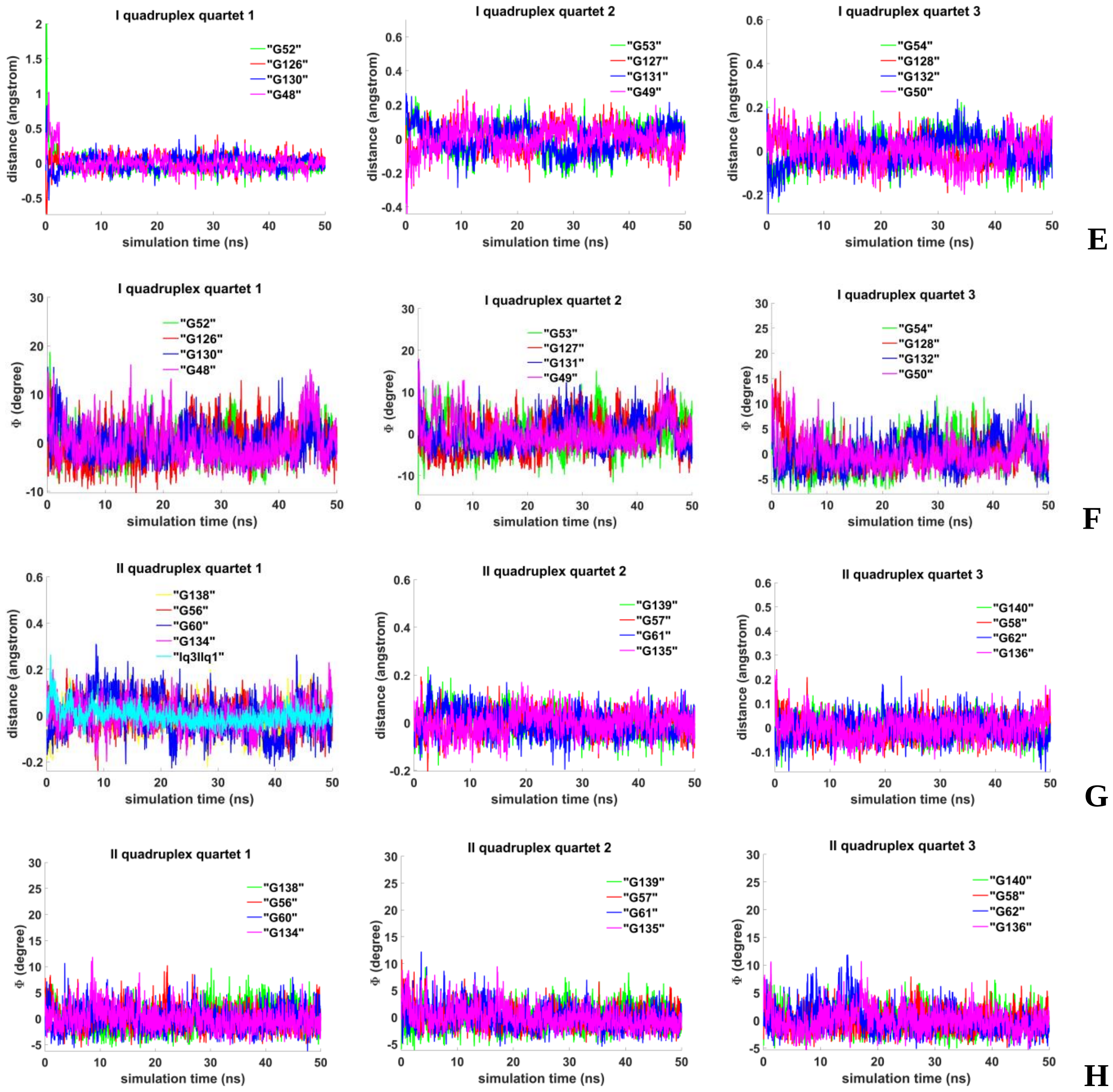


**Figure 7.4.A.S.3. Case 7: "Stacking of three parallel G4-dimer and two IM-monomers with mutual girth of the strands": E, G** - distances from COMs of the guanines' bases to COMs of their containing tetrad, distance between COMs of the boundary tetrads (**Iq3 IIq1**); **F, H** - angles between normals to the guanines' bases and vectors connecting COMs of the boundary tetrads.

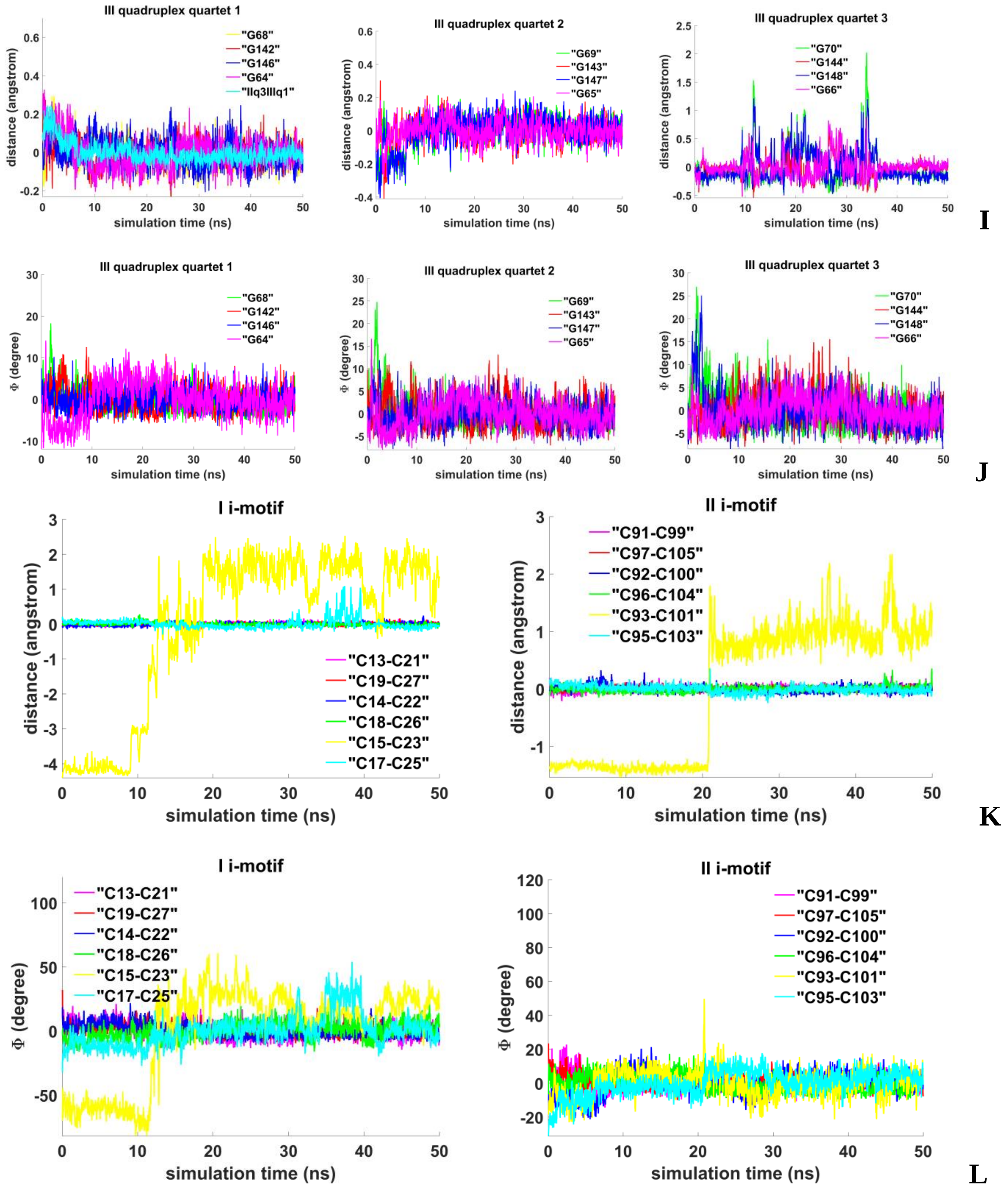


**Figure 7.4.A.S.3. Case 7: "Stacking of three parallel G4-dimer and two IM-monomers with mutual girth of the strands": I** - distances from COMs of the guanines' bases to COMs of their containing tetrad, distance between COMs of the boundary tetrads (**IIq3 IIIq1**); **J** - angles between normals to the guanines' bases and vectors connecting COMs of the boundary tetrads; **K**- distances between COMs of the cytosines' bases; **L** - angles between normals to the cytosines' bases.

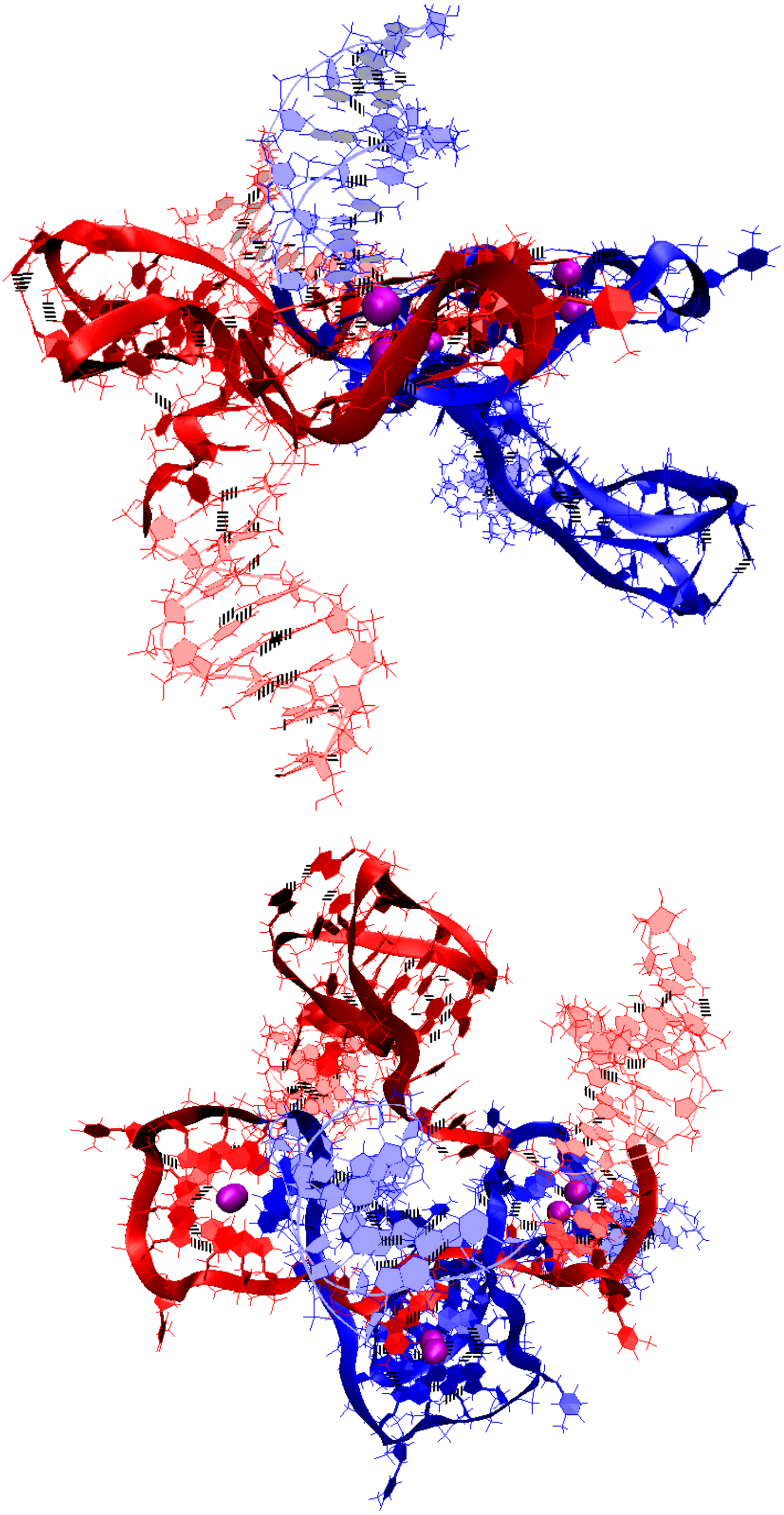

A

**Figure 7.4.B.S.1. Case 8: "Three parallel G4-dimers in the same plane with and two IM-monomers with mutual girth of the strands"**: **A** – conformation obtained at the last step of the MD trajectory (side and top view).

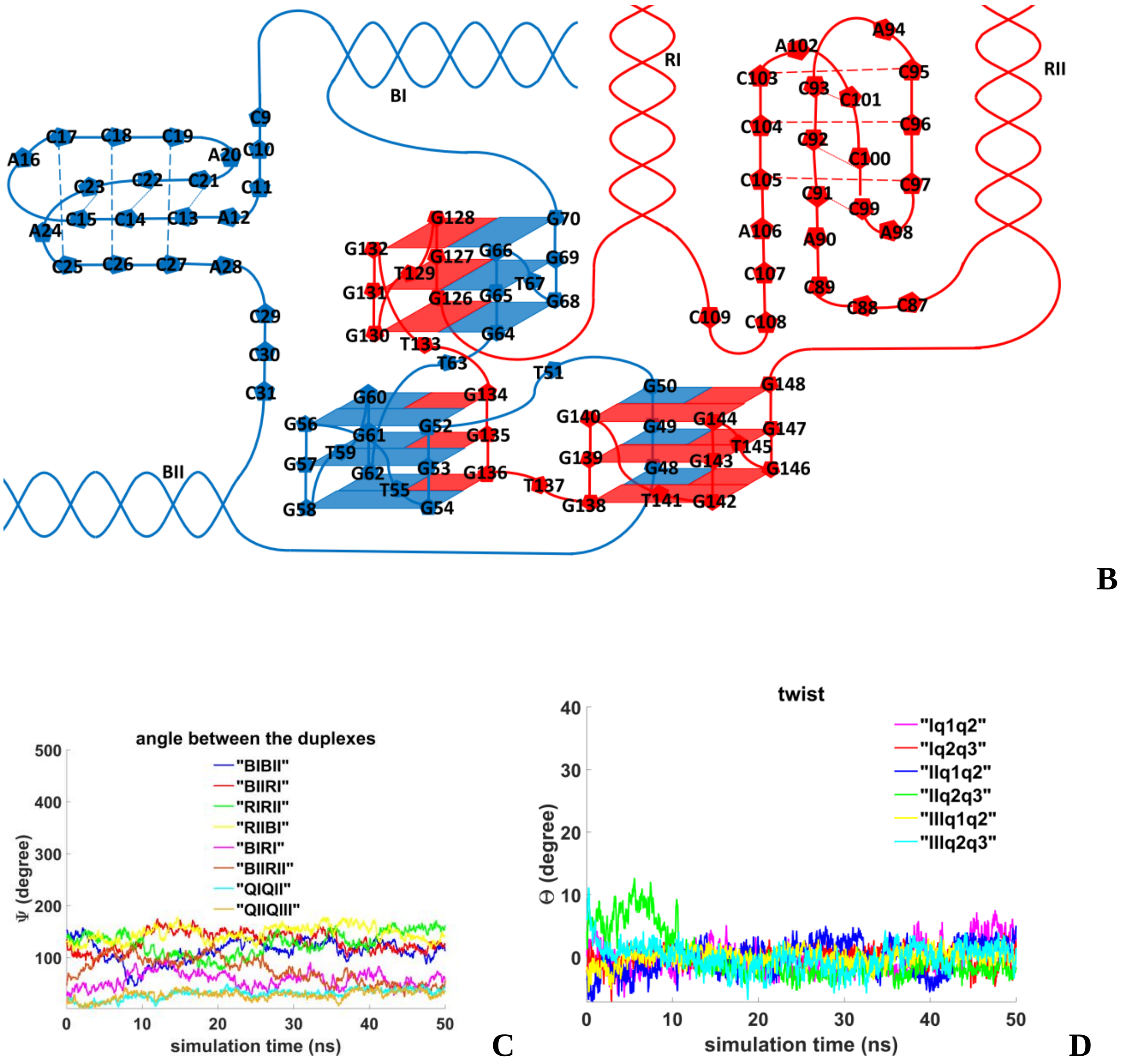


**Figure 7.4.B.S.2. Case 8: "Three parallel G4-dimers in the same plane with and two IM-monomers with mutual girth of the strands"**: **B** – the complex scheme; **C** – angles between unmelted fragments of the duplexes and the axes passing through the COMs of the tetrads, the angle values between the G4s (**QIQII**, **QIIQIII**); **D** – angles of rotation of the tetrads relative to each other.

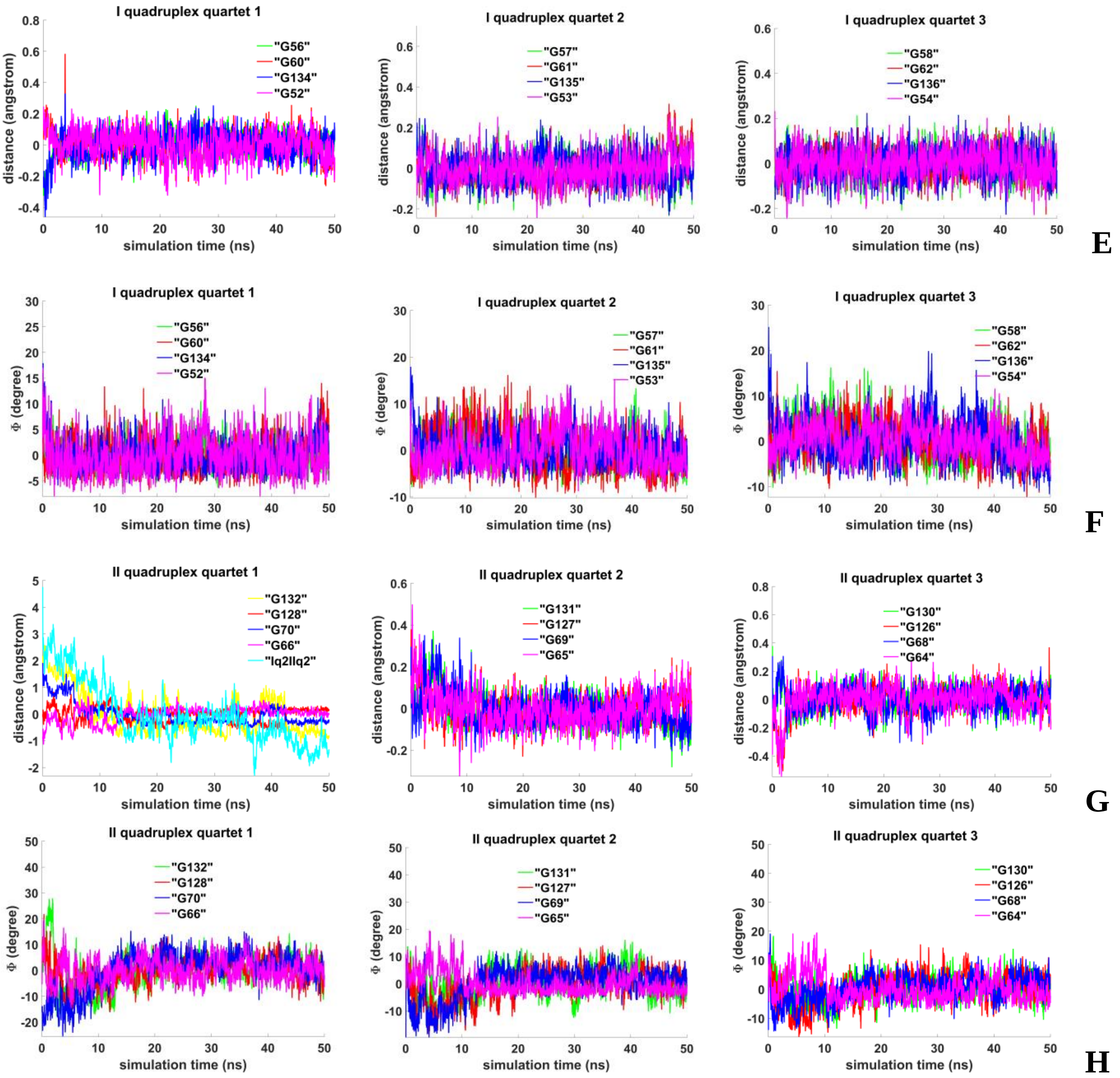


**Figure 7.4.B.S.3. Case 8: “Three parallel G4-dimers in the same plane with and two IM-monomers with mutual girth of the strands”**: **E, G** - distances from COMs of the guanines’ bases to COMs of their containing tetrad, distance between COMs of the G4s (**Iq2 Iiq2**); **F, H** - angles between normals to the guanines’ bases and vectors connecting COMs of the boundary tetrads.

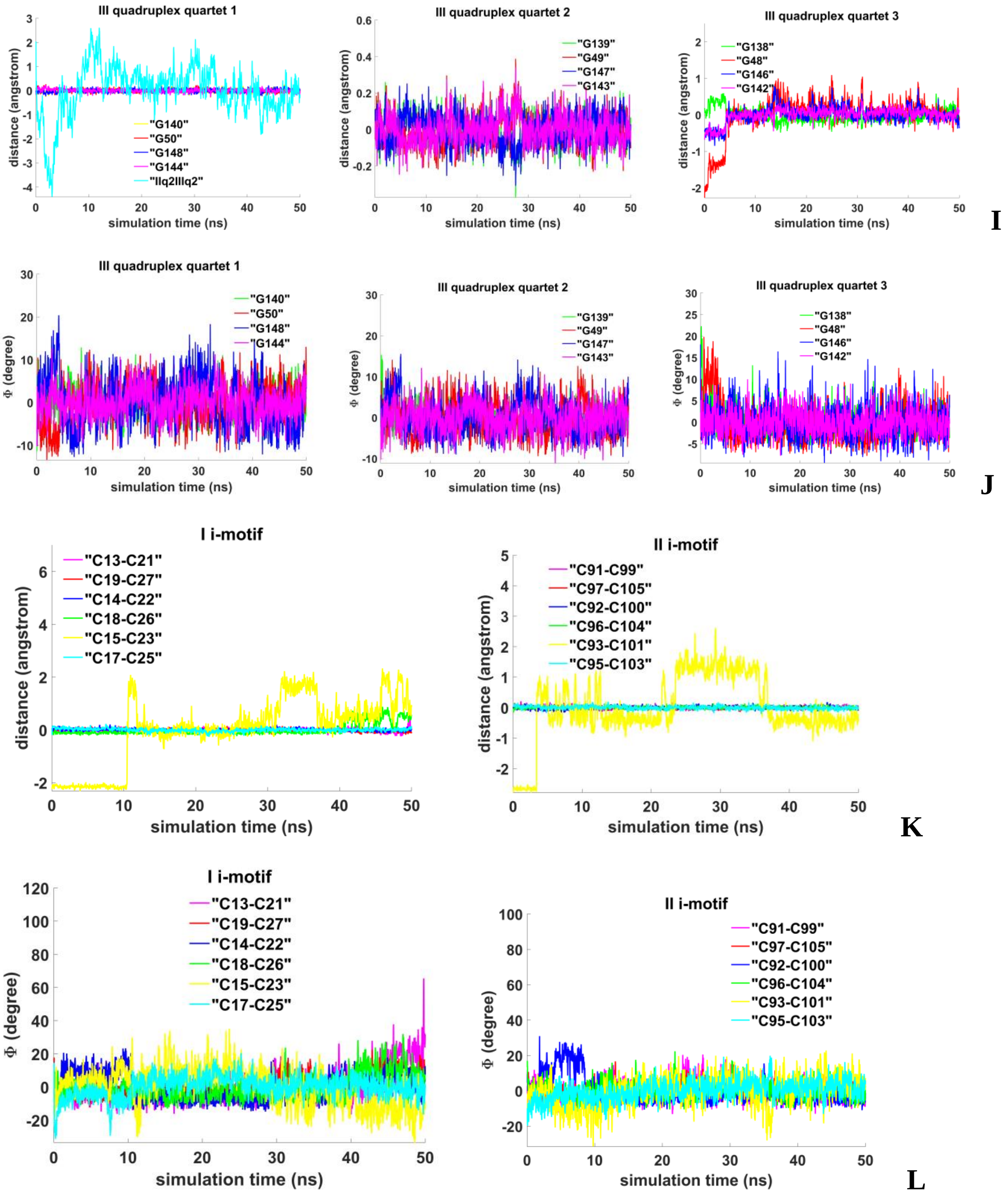


**Figure 7.4.B.S.4. Case 8: “Three parallel G4-dimers in the same plane with and two IM-monomers with mutual girth of the strands”**: **I** - distances from COMs of the guanines’ bases to COMs of their containing tetrad, distance between COMs of the boundary tetrads (**Iiq2 IIIq2**); **J** - angles between normals to the guanines’ bases and vectors connecting COMs of the boundary tetrads; **K** - distances between COMs of the cytosines’ bases; **L** - angles between normals to the cytosines’ bases.

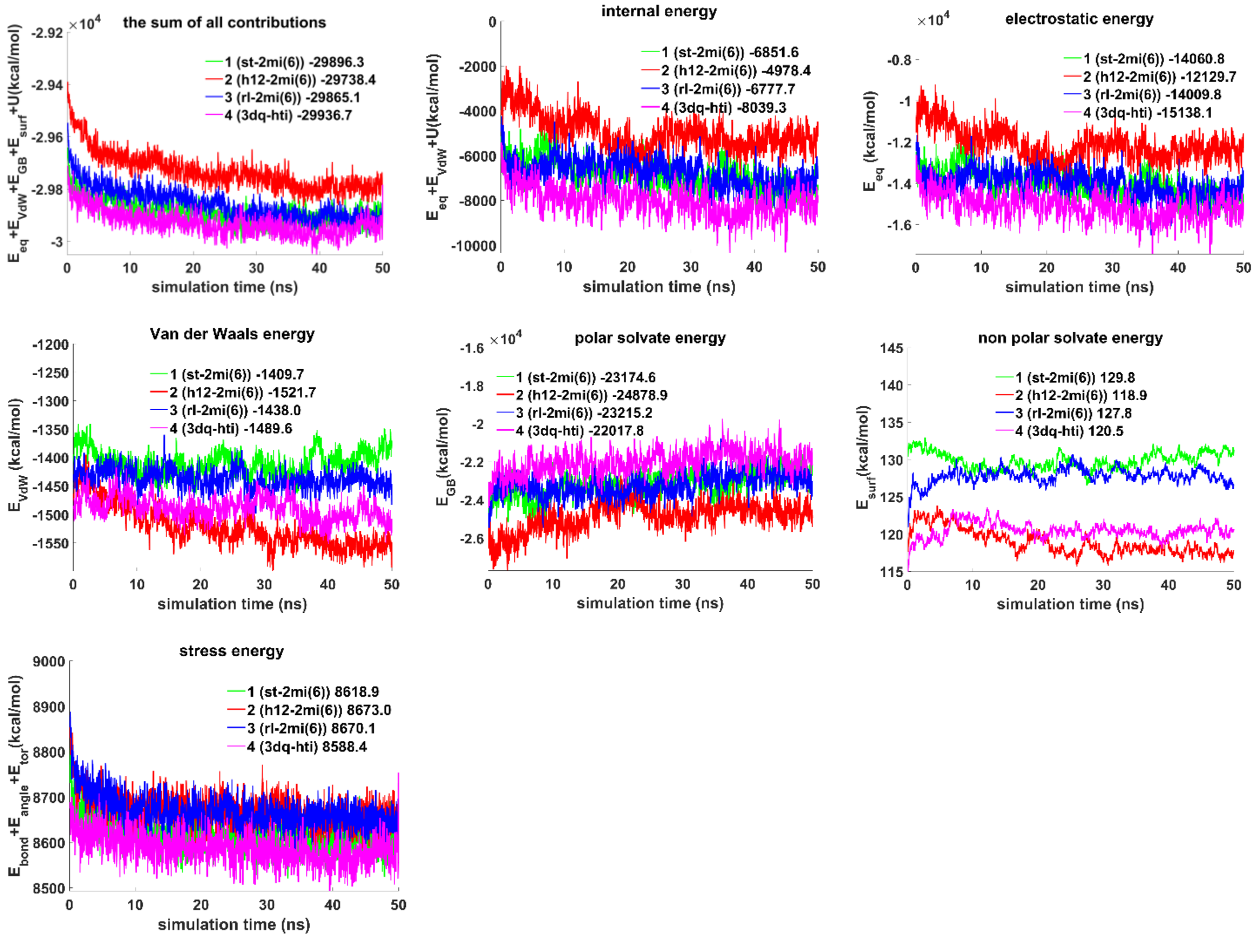


**Figure 7.S.E.1. The contributions to free energy during MD calculations for the variants of bimolecular complex of duplexes containing $(G_3T)_5G_3$ and $(C_3A)_5C_3$ sequences with G4/IM in cases from 1 to 4.** $E_{eq}$ – electrostatic, $E_{vdw}$ - Van der Waals, $E_{GB}$ - polar energy of solvation, $E_{surf}$ - non-polar energy of solvation due to the hydrophobic surface available to the solvent, $U = E_{bond} + E_{angle} + E_{tor}$ , e.g. $E_{bond}$, $E_{angle}$ and $E_{tor}$ –bond, angle and torsion stress energies. The energy plots were smoothed using moving average method (span = 5). Average energy values are indicated in the figure legends.

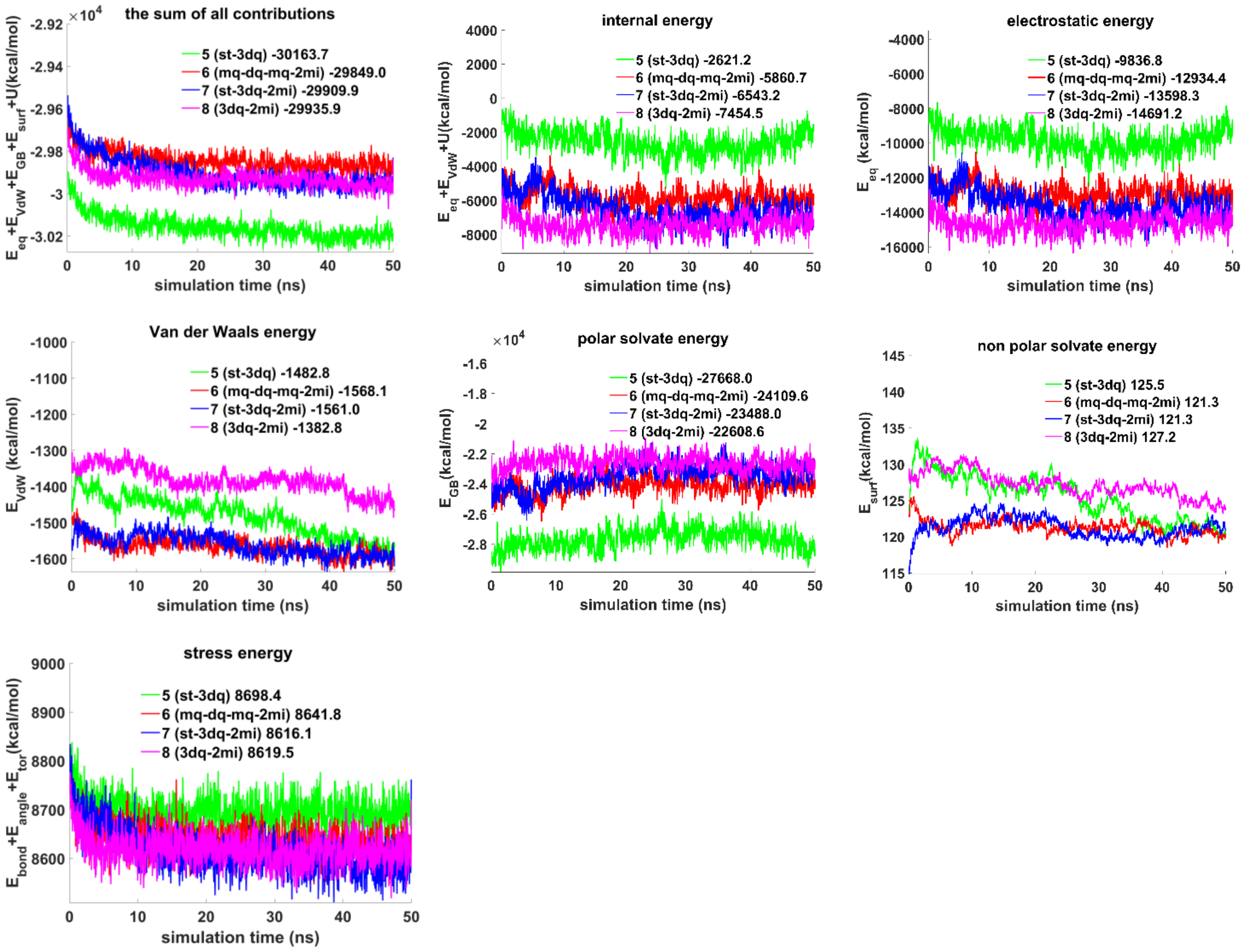


**Figure 7.S.E.2. The contributions to free energy during MD calculations for the variants of bimolecular complex of duplexes containing $(G_3T)_5G_3$ and $(C_3A)_5C_3$ sequences with G4/IM in cases from 5 to 8.** $E_{eq}$ – electrostatic, $E_{vdw}$ - Van der Waals, $E_{GB}$ - polar energy of solvation, $E_{surf}$ - non-polar energy of solvation due to the hydrophobic surface available to the solvent, $U = E_{bond} + E_{angle} + E_{tor}$ , e.g. $E_{bond}$, $E_{angle}$ and $E_{tor}$ –bond, angle and torsion stress energies. The energy plots were smoothed using moving average method (span = 5). Average energy values are indicated in the figure legends.
